# Movement strategies reveal the success of mammals in urban areas

**DOI:** 10.64898/2026.07.13.738202

**Authors:** Marius Grabow, Carolin Scholz, Manuel Roeleke, Milena Stillfried, Sophia Kimmig, Carolin Weh, Konstantin Börner, Niels Blaum, Florian Jeltsch, Sylvia Ortmann, Stephanie Kramer-Schadt

**Author notes:** Corresponding author: Marius Grabow.

## Abstract

Various hypotheses have been proposed to explain why some species persist or even flourish in urban areas. Yet, despite its central role in determining when and where animals encounter resources, disturbance, and risk, movement behaviour remains an overlooked mechanism of urban success. In urban areas, human activities are strongly periodic, i.e. predictable in space and time. This may favour species able to adjust their behaviour to predictable cycles of resources and risks in space and time. Here, we tested this hypothesis and tracked movement behaviour along an urbanisation gradient in three mammal species with different urban success: red fox (*Vulpes vulpes*), an urban dweller; raccoon (*Procyon lotor*), an invasive urban dweller; and wild boar (*Sus scrofa*), an urban utiliser. We analysed periodicity in movement behaviour and investigated whether increasing urbanisation is associated with periodic reorganisation of activity timing, space use, and further analysed alterations in habitat selection along the urbanisation gradient. Our results show that foxes aligned their movement behaviour with human activity, having stronger day-night contrasts and more repeatable space use than their rural counterparts. Urban raccoons showed a contrasting strategy; they were more active during the day, without changes in their movement routines under increasing urbanisation, suggesting a flexible strategy that explains their urban success. In contrast, wild boars reduced routine movement behaviours with increasing urbanisation, consistent with their occurrence in less predictable suburban environments and avoidance of city centres. In summary, our results suggest that movement behaviour may be a key mechanism enabling animals to persist in cities, revealing distinct behavioural strategies for coping with urban environments.

## 1 Introduction

Urbanisation is one of the most pervasive pressures on wildlife. Yet, some species persist or even flourish in these novel environments (Hansen et al., 2024). While it is generally accepted that animals avoid human presence and roads (Franke et al., 2026), identifying the mechanisms that allow some species to thrive in these novel habitats while others decline remains a key question. To inhabit urban habitats, animals must navigate landscapes where human activity dominates and structures resources and risks more strongly than in most non-urban systems. Because human-driven dynamics are often predictable, they may favour species capable of tracking recurring resource-risk cycles (Bautista et al., 2004; Péron et al., 2016). Many species shift their activity timing to reduce direct encounters with humans (Gaynor et al., 2018), a pattern that is often more pronounced in urban habitats, where temporal activity patterns tightly link to human routines (Drenske et al., 2024; Louvrier et al., 2022). Yet, analysing temporal niche shifts alone remains inconclusive because they do not capture how individuals restructure space use in urban areas. By focusing only on when animals are active, we risk missing the mechanisms through which urban species adjust where they move, which resources they access, and how they persist under novel urban conditions (Péron et al., 2017). In this context, examining individual behaviour becomes essential (Grimm et al., 2025) because it reveals whether wildlife responds to urban conditions through a shared restructuring of movement and space use or via distinct strategies among individuals and species, thereby helping explain how behavioural variation shapes urban community assembly (Schlägel et al., 2020).

Numerous traits have been proposed to explain why some species persist or even flourish in cities (Weiss et al., 2023), including broader diets (e.g. Palacio, 2020), behavioural flexibility (Lee & Thornton, 2021), increased reproductive output (Takahata & Kutsukake, 2025), or reduced sensitivity to human disturbance (Ritzel & Gallo, 2020). Many of these traits will ultimately influence how individuals move through urban landscapes. For example, species exploiting anthropogenic food sources may repeatedly return to predictable resource patches, whereas species tolerant of human presence may expand their movements into otherwise risky areas. Conversely, species that persist by avoiding disturbance may restrict their movements to quieter areas or adjust the timing of their visits to times of lower human activity (e.g. Tolon et al., 2009).

Movement behaviour links individuals to their environment (Nathan et al., 2008). Therefore, studying movement behaviour of animals can reveal the mechanisms how animals adjust to urban environments (Gomez et al., 2025). For example, during COVID-19 stay-at-home orders, mountain lions (*Puma concolor*) increased trail use in response to reduced human activity, indicating rapid adjustment to shifts in perceived risk (Benson et al., 2021). Similarly, along an urbanisation gradient, grey wolves *(Canis lupus)* balanced resource exploration and risk perception that may enable persistence in more urban habitats (Lazzaroni et al., 2026). In home range-bound species, such behavioural adjustments may be expressed through how individuals repeatedly structure space use (Riotte-Lambert et al., 2017; Riotte-Lambert & Matthiopoulos, 2020), i.e. rely on reusing travel routeways and structured travel routes, as detected in canids (Fagan et al., 2025). Such revisitation of certain locations is common within home ranges (Bar-David et al., 2009; Berger-Tal & Bar-David, 2015; Bracis et al., 2018). However, in human dominated environments, little is known whether such revisitation follows consistent temporal patterns that reflect tracking of predictable resource–risk dynamics (Péron et al., 2016, 2017, 2018; Riotte-Lambert et al., 2017).

Mammal communities comprise species that vary systematically in their response to urbanisation, ranging from ‘urban avoiders’, which are largely absent from cities, to ‘urban utilisers’ and ‘urban dwellers’ (sensu Fischer et al., 2015), which increasingly benefit from urban conditions.

We hypothesize that species that differ in their degree of adjustment to urbanisation also differ, above all, in the extent to which periodic movement is evident along the urbanisation gradient. We predict movement routines reflecting the spatial distribution and persistence in urban habitats, where species occurring in the urban core will be more aligned with human routines. Here, we compiled movement data from two urban dwellewow, that’s way too longrs and an urban utiliser along a rural-to-urban gradient in and around the city of Berlin, Germany. More specifically, we investigated species-specific signatures of space use and activity along the urbanisation gradient, focusing on recurrent, periodic movement behaviour, expressed as temporal niche shifts and repeated habitat selection patterns. We compared red fox (*Vulpes vulpes,* hereafter: fox), a generalist with an omnivorous diet and urban dweller (Scholz et al., 2020) with raccoon (*Procyon lotor*), an invasive urban dweller with a broad dietary niche and high flexibility in space use due to its ability to climb (M. Fischer et al., 2026). Both species cover the entire urban area, including the city centre (e.g. Louvrier et al., 2022; Scholz et al., 2024). As a reference species, we included wild boar (*Sus scrofa*), an urban utiliser of suburbs, typically avoiding urban core areas but persisting at urban outskirt areas (Stillfried, Gras, Börner, et al., 2017; Stillfried, Gras, Busch, et al., 2017).

We expected species to differ in periodic movement behaviour along the urbanisation gradient (see next paragraph) because urban success, here defines as persisting in the dense urban matrix, should depend on how animals organise movement around the recurring risks and resources of human-dominated landscapes (Figure 1a). Periodicity may emerge not only from repeated use of spatially predictable resources, as for example present in trash bins of restaurants and around other points of interest (POIs) for humans or tree cover as refuge habitat, but also from consistent spatiotemporal avoidance of predictable disturbance, like roads. In this context, urban responses need to be assessed across complementary dimensions of behaviour. Periodic movement patterns indicate whether individuals repeatedly organise space use in time, consistent with tracking recurring environmental conditions. However, adjustments to predictable human activity may also be expressed through shifts in temporal niche use, i.e. when animals are active, and spatial niche use, i.e. where animals select resources and avoid risks. Together, these metrics capture how individuals coordinate movement, timing, and space use in response to structured urban environments, providing a mechanistic framework to assess species-specific strategies along urbanisation gradients.

**Figure 1.**
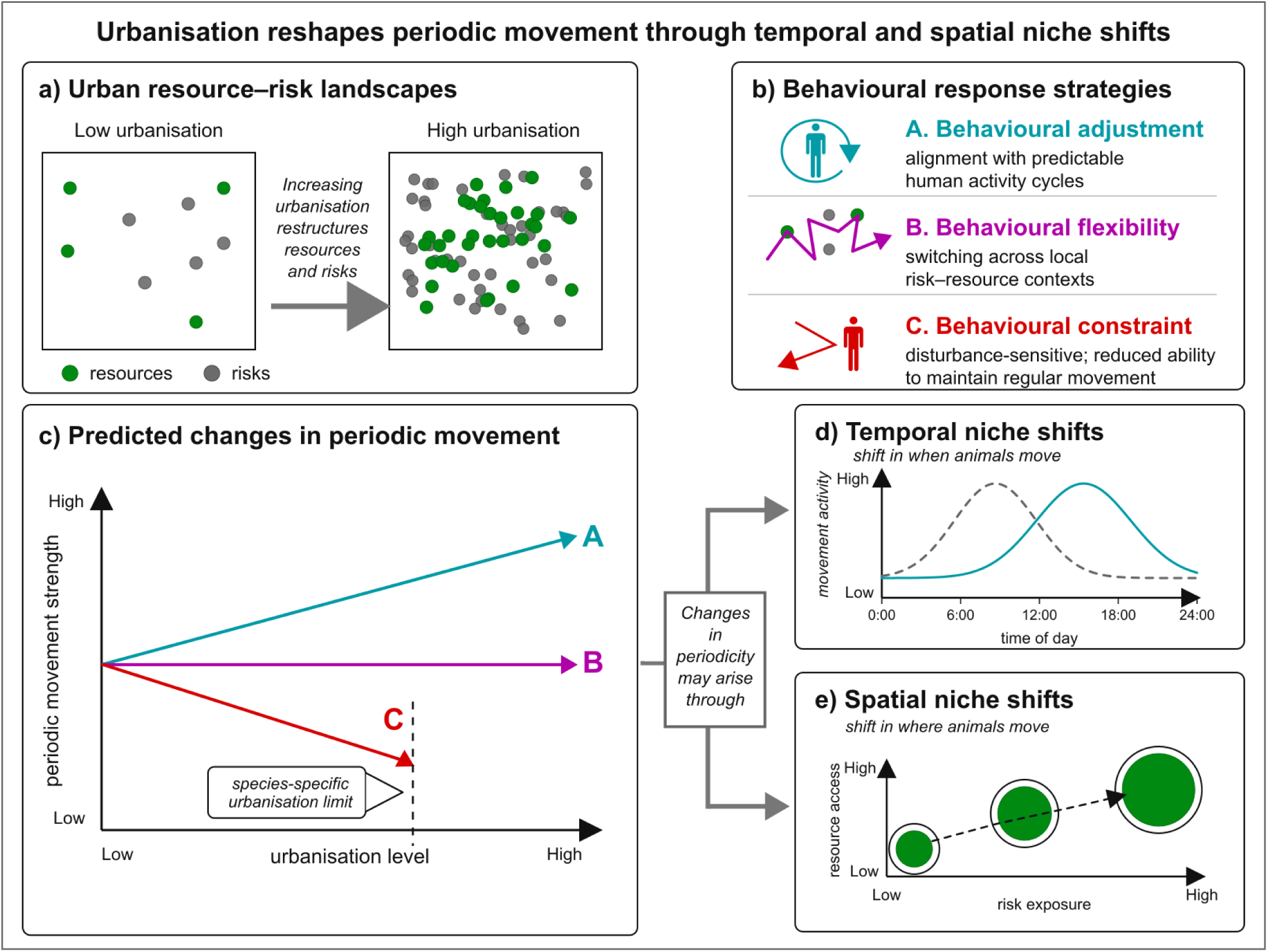
Conceptual overview of predicted movement responses to urbanisation. a) Urbanisation restructures resource–risk landscapes from low-intensity, spatially dispersed conditions to high-intensity urban environments where resources and risks are more concentrated and linked to predictable human activity cycles. b) Species may respond through distinct behavioural strategies: adjustment to predictable human dynamics, flexible switching among local risk–resource contexts, or constraint under recurrent disturbance. c) These strategies are expected to alter periodic movement differently: behavioural adjustment should increase routine, repeatable movement; behavioural flexibility may allow persistence without stronger periodicity; and behavioural constraint should reduce routine movement and limit persistence at high urbanisation. Changes in periodicity may arise through d) temporal niche shifts, altering when animals move, and e) spatial niche shifts, altering where animals move across risk–resource gradients.

For foxes, occurrence in dense urban cores suggests a capacity to exploit highly urbanised habitats, but this should require tight alignment of movement with daily cycles of human activity, risk, and anthropogenic resources. We therefore predict stronger periodicity with urbanisation, reflecting routine spatiotemporal alignment with human-induced risk–resource cycles and supporting persistence in urban-core habitats (Figure 1c; A). We expected this pathway to be expressed through strong temporal niche shifts, with marked diel contrasts in resource selection and risk avoidance driven by human activity times (Figure 1d–e). In contrast, we expected raccoons to persist through flexibility rather than routine, consistent with their invasive success in the study area and their occurrence even in dense urban habitats. Their flexible denning behaviour, climbing ability, dexterity, and broad diet should allow them to access resources and refuges opportunistically across the urban matrix, potentially reducing raccoons’s need for periodic movement (Figure 1c; B). Finally, we expected wild boars to face stronger constraints with increasing urbanisation. Their large body size, limited access to fine-scale refuges, and dependence on suitable foraging habitats should make dense urban areas harder to navigate, leading to a loss of structured periodic movement towards urban cores, with occasional use of food sources and higher use of refuge habitat, and restricting persistence to less intensely urbanised areas like suburbs (Figure 1c; C).

## 2 Methods

### 2.1 Study area and animal tracking data

We studied movement behaviour of foxes (*Vulpes vulpes*), raccoons (*Procyon lotor*), and wild boar (Sus scrofa) along an urbanisation gradient in Berlin (52.5200° N, 13.4050° E) and surrounding Brandenburg (Figure 2a-b). Berlin contains dense urban cores, around 3.8 million inhabitants, 63% built-up area, and 26.8% green features, including parks and forested areas near the administrative borders (SenMVKU, 2025). Brandenburg extends this gradient from peri-urban landscapes near Berlin to rural areas dominated by agriculture, forests, and lakes. (Figure 2b).

**Figure 2.**
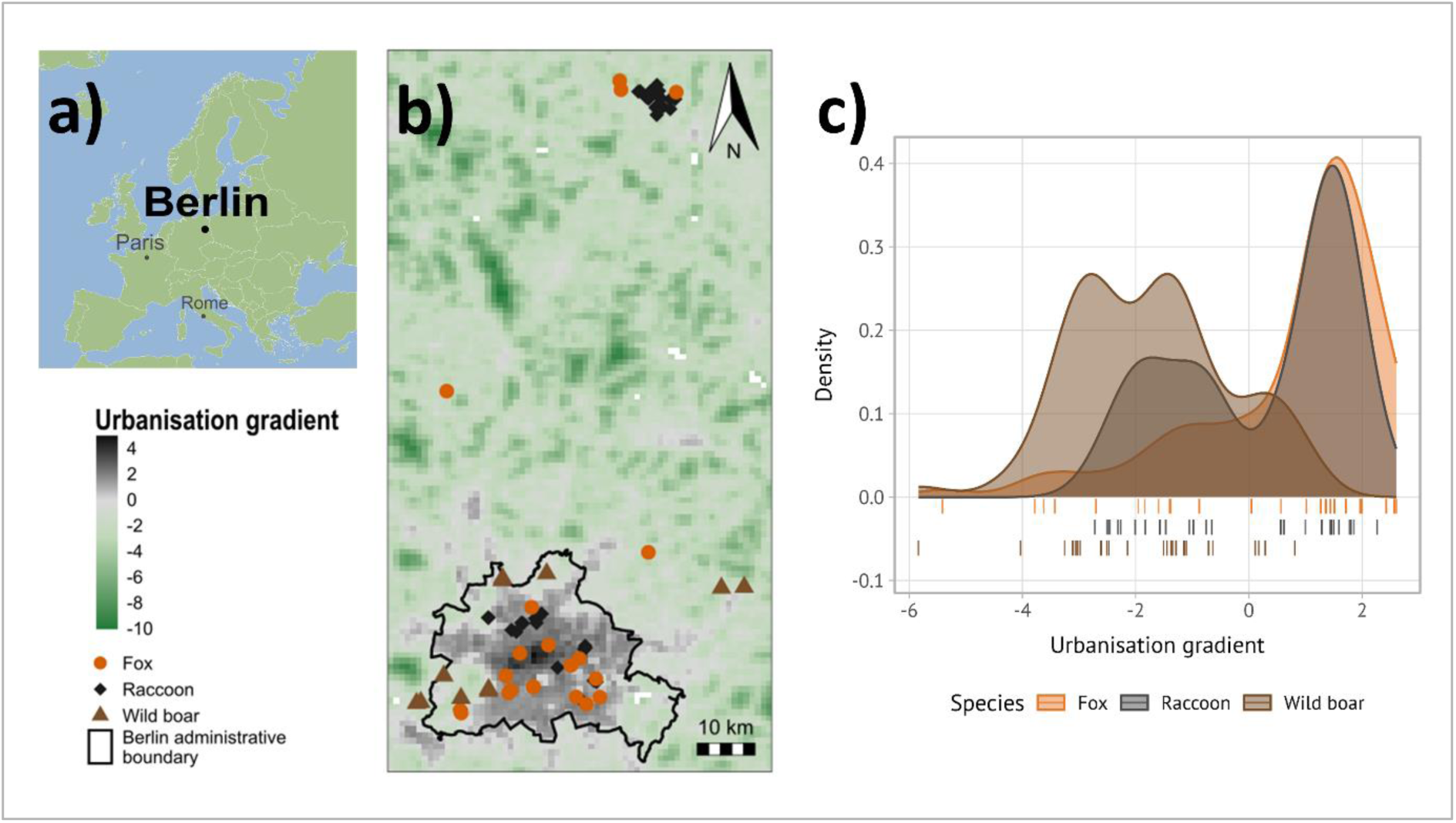
Species coverage along the urbanisation gradient. a) Centroids of tracked individuals in and around Berlin, from less urbanised areas in green to highly urbanised areas in grey to black. b) Species-specific coverage of the gradient.

We compiled movement data from studies conducted in Berlin and Brandenburg between 2013 and 2023 (S. Kimmig, 2021; Roeleke et al., 2025; Scholz, 2021; Stillfried, Gras, Börner, et al., 2017). Tracking protocols, durations, and species coverage varied among studies (Supplementary S1; Figure 2c): foxes and raccoons occurred across broad urbanisation ranges, including dense urban areas, whereas wild boars were mainly located toward less urbanised city outskirts. In total, we collared 80 individuals with GPS devices (21 foxes, 48 raccoons, 11 wild boars). We split movement data by calendar month to account for seasonality and retained only individual-month tracks with at least 14 days of monitoring, resulting in 295 observations from 18 foxes, 33 raccoons, and 11 wild boars.

We extracted environmental covariates from publicly available geospatial data to describe urbanisation across the study area (Supplementary S2). We performed a principal component analysis (PCA) on standardised raster layers representing human modification, light pollution, distance to roads, and tree cover density. The first principal component (PC1) captured the main urbanisation gradient, with higher values indicating higher human modification and light pollution, shorter distances to roads, and lower tree cover density. We projected PC1 back into geographic space using the PCA loadings (Figure 2b), generating a continuous raster from which we extracted urbanisation values for animal locations and model predictions.

### 2.2 Periodic movement behaviour

We tested whether individuals returned to similar positions at regular time intervals (Péron et al., 2016, 2016), as expected if movements were structured by diel human activity patterns. For each individual-month track, we used Lomb-Scargle periodograms with fast Fourier transformations in *ctmm* (Calabrese et al., 2016) to identify peaks at natural periods, including the 24-hour diel cycle and its harmonics. We compared these peaks with tag-specific sampling intervals and their multiples to ensure that detected periodicities did not reflect GPS schedules (Supplementary S3). We then fitted continuous-time periodic-mean movement models with pre-specified periods of 24 hours and its harmonics, with model selection implemented in *ctmm* (Calabrese et al., 2016).

We analysed periodicity using a two-part Bayesian regression approach in *brms* (Bürkner, 2017), similar to a hurdle model. First, we modelled whether periodicity was detected using a Bernoulli model; second, for tracks with detected periodicity, we modelled periodic strength using beta regression. Both components included species, urbanisation PC1, their interaction, linear and quadratic month effects, and individual identity as a random intercept. Models were fitted with four chains, 8,000 iterations per chain, and 2,000 burn-in samples. Inference was based on posterior summaries and contrasts using 89% credible intervals following Kruschke et al. (2015). Patterns found were subsequently disentangled into temporally and spatially recurring behaviour.

### 2.3 Temporal niche shifts

To decompose periodic movement behaviour into temporal activity periods, we fitted species-specific Generalised Additive Mixed Models (GAMMs; see Supplementary S4 for model formulas) using movement-derived speed as a proxy for movement activity in *mgcv* (Wood, 2011). Speed was calculated as step length per unit time, scaled before fitting, and interpreted as movement activity because it integrates low-movement periods and active displacement. We modelled 24-hour variation in speed using a cyclic cubic regression spline for time of day, included urbanisation as continuous PC1, and allowed diel activity curves to vary along this gradient through a smooth interaction between time of day and PC1. Models were fitted using Restricted Maximum Likelihood (REML) and included individual identity as a random intercept. For visualisation, we derived marginal predictions at low, intermediate, and high urbanisation, defined as the 10th, 50th, and 90^th^ percentiles within each species’ observed PC1 range. We additionally tested whether solar time or fixed-clock time better described the observed activity patterns, and whether their relative performance changed with increasing urbanisation (Supplementary S4).

### 2.4 Spatial niche shifts

To quantify habitat selection while accounting for movement capacity, we used integrated Step Selection Functions (iSSFs), modelling habitat use at the step level conditional on observed movements (Avgar et al., 2016). Tracks were resampled to regular intervals of 20, 36, and 30 minutes for foxes, raccoons, and wild boars, respectively, corresponding to predominant sampling intervals. We retained bursts with at least five observed locations, generated 10 random steps per observed step, and extracted covariates using *amt* (Signer et al., 2019). Observed and random steps were contrasted using conditional Poisson Generalised Linear Mixed Models (GLMMs) in glmmTMB, following Muff et al. (2020). We modelled step selection as a function of environmental covariates and their interaction with time of day along the urbanisation gradient.

For the iSSF analyses, we used human modification as a fine-scale proxy for local urbanisation because the composite PC1 was available only at 1 × 1 km resolution and was therefore reserved for broad-scale analyses. Human modification was the strongest contributor to PC1 (Supplementary S2) and provided a spatially appropriate measure for step-level habitat selection. We modelled selection as a function of proximity to roads, proximity to points of interest (POIs), tree-cover density, and their interactions with time of day and human modification. Proximity variables were calculated as flipped distances, such that larger values indicate greater proximity. Roads represented linear infrastructure associated with disturbance and movement constraints; POIs represented concentrations of human activity and potential anthropogenic resources; and tree cover represented vegetated refuge habitat. We contrasted times of high human activity (6 am–10 pm) and low human activity (10 pm–6 am) and report results as relative selection strength (RSS).

These covariates were selected to capture three ecologically distinct components of the urban landscape. Roads represent linear infrastructure associated with disturbance, mortality risk, habitat edges, and potential movement barriers or corridors. POIs represent spatial concentrations of human activity and associated anthropogenic resource opportunities, such as buildings of public interest, facilities, recreational areas, or food-related infrastructure. Tree-cover density was included as a measure of vegetated cover and potential refuge habitat within the urban matrix. Together, these covariates allowed us to test whether animals responded to urbanisation by shifting their space use towards human-associated features, avoiding concentrated human activity, or increasing use of vegetated refuge habitats.

## 3 Results

### 3.1 periodic movement behaviour

We found species-specific, urbanisation-mediated differences in the probability that periodic movement was detected (Figure 3a; Supplementary S5). Periodicity tended to decline with urbanisation in wild boar, the reference species (β = −0.42 [−0.95, 0.05], P(β < 0) = 0.92), but tended to increase in foxes (β = 0.30 [−0.08, 0.68], P(β > 0) = 0.90). Accordingly, the fox urbanisation slope was more positive than the wild boar slope (fox − wild boar: β = 0.72 [0.11, 1.38], P(β > 0) = 0.97). Raccoons showed little directional response (β = 0.03 [−0.29, 0.35], P(β > 0) = 0.57), and their slope did not clearly differ from either species. Baseline probabilities differed mainly because foxes showed a higher baseline probability of periodicity than wild boar (β = 1.66 [0.30, 3.10]).

**Figure 3:**
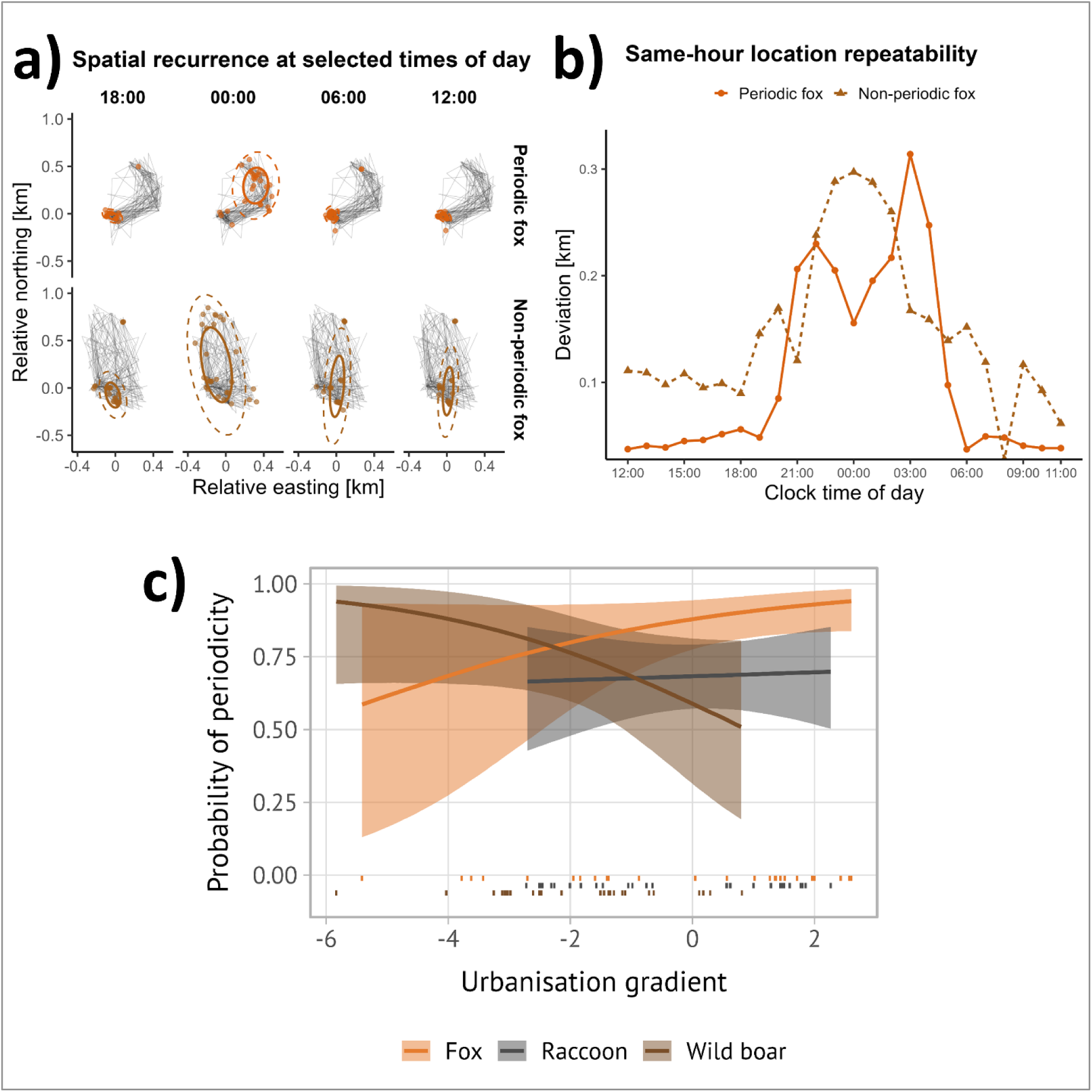
Urbanisation is associated with species-specific changes in periodic movement behaviour, illustrated by recurrent and non-recurrent fox trajectories. a) Daily trajectories of two representative foxes across 30 consecutive days; coloured points show locations at selected times of day. Solid and dashed ellipses indicate the 50% and 90% spatial spread of same-hour locations. b) Same-hour location repeatability, quantified as the median distance between each location and the individual’s usual location at that hour. c) Model-predicted probability of periodic movement along the urbanisation gradient. Lines show species-specific predictions, ribbons show 89% credible intervals, and rug marks indicate observed individuals

Periodicity also varied seasonally and among individuals. The quadratic month effect was clearly negative (β = −0.04 [−0.07, −0.02]), indicating higher periodicity during summer and lower periodicity toward winter, whereas the linear effect was weak (β = −0.02 [−0.09, 0.05]). Among-individual variation was substantial (SD = 1.20 [0.66, 1.83]). Positive periodic strength varied little with urbanisation or species (Supplementary S5), indicating that species differences were driven mainly by whether a periodic component was detected rather than by its strength once present.

### 3.2 Temporal niche shifts

Fixed-clock time, reflecting human activity schedules, generally described activity patterns better than first-light time, suggesting that diel movement patterns were more closely aligned with human schedules than with natural light conditions, particularly in highly urbanised foxes (Supplementary S4). We therefore report clock-time models here and first-light models in the Supplementary.

In foxes and raccoons, movement-derived activity varied strongly across the diel cycle, but urban-associated changes differed markedly between species (Figure 4). In foxes, mean activity increased with urbanisation (estimate = 0.125, p < 0.001), and diel activity structure changed significantly along the gradient (time of day × urbanisation: F = 93.54, p < 0.001). Model predictions showed that foxes were most active at night, outside the main period of human activity, and that this nocturnal peak became stronger at higher urbanisation levels.

**Figure 4:**
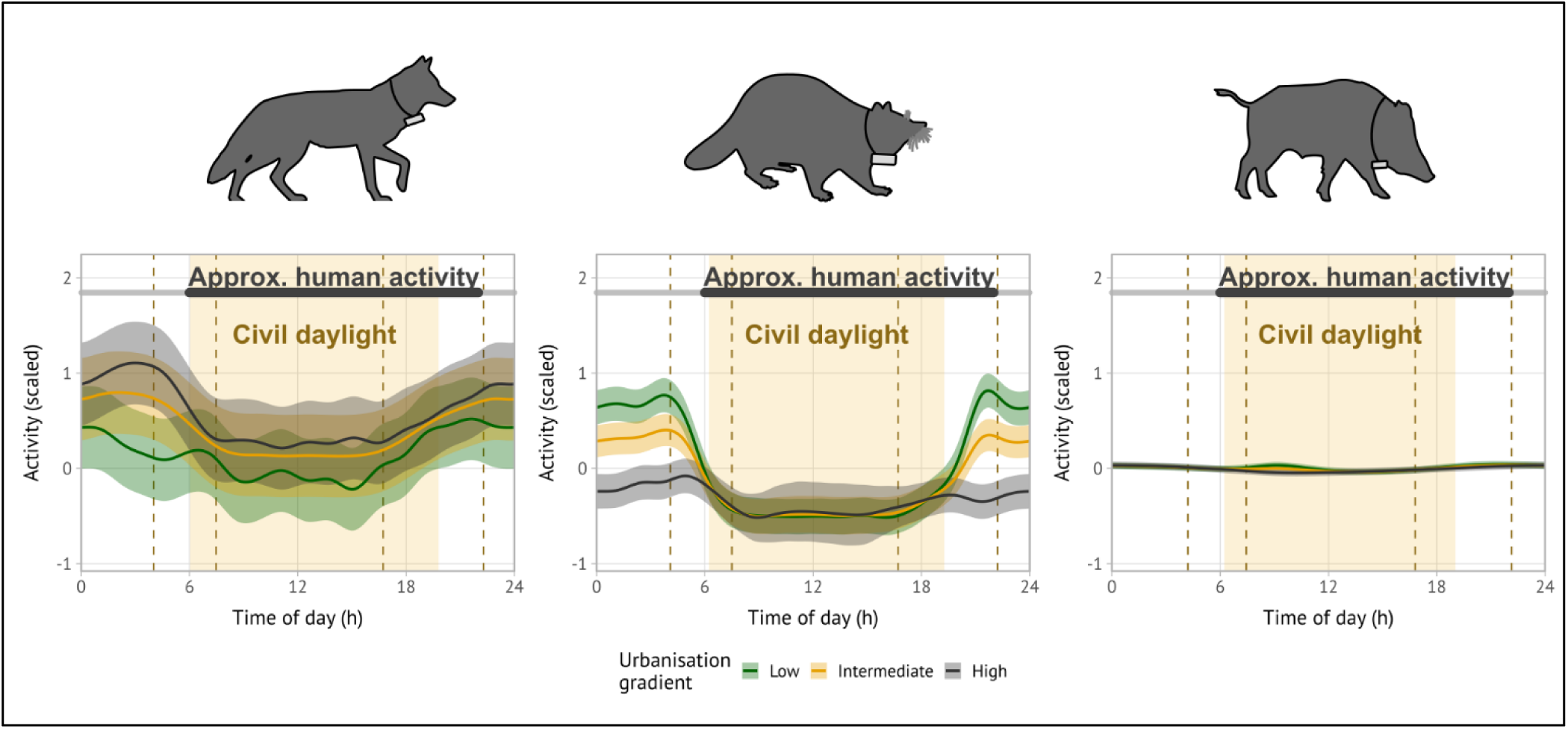
Predicted diel activity patterns from species-specific GAMMs. Activity is shown as scaled movement-derived speed across local time of day. Lines represent marginal model predictions and ribbons indicate 95% confidence intervals. Predictions are shown at low, intermediate, and high urbanisation, defined as the 10th, 50th, and 90th percentiles within each species. The yellow shading indicates the median civil daylight period, from median civil dawn to median civil dusk. Dashed yellow lines show the 10th–90th percentile range of civil dawn and civil dusk across sampling dates, illustrating seasonal variation in daylight timing. The line background indicates the approximate times of human activity, defined as 06:00–22:00 (see Methods).

Raccoons showed a contrasting pattern. Mean activity declined with urbanisation (estimate = −0.082, p < 0.001), while activity varied strongly across the diel cycle (F = 1356.95, p < 0.001). The diel activity curve also changed significantly along the gradient (time of day × urbanisation: F = 63.98, p < 0.001), but urban raccoons did not show increased nocturnal activity. Instead, the nocturnal peak was reduced at higher urbanisation levels, indicating lower overall activity, including at night.

In wild boar, we found little evidence for systematic diel variation or urban-associated restructuring of activity. Mean activity did not vary clearly with urbanisation (estimate = −0.004, p = 0.394), the main time-of-day smooth was not supported (F = 1.00, p = 0.887), and neither was the time-of-day × urbanisation interaction (F = 1.21, p = 0.237). This result indicates weak population-level variation in mean step-derived speed, not absence of rest or diel behaviour. Inference for highly urbanised conditions was also limited because wild boars occurred mainly at lower urbanisation levels (Figure 2b). Across species, individual identity explained significant variation in activity levels (Supplementary S4).

### 3.3 Spatial niche shifts

Step selection differed strongly among species, times of human activity, and local human modification (Figure 5; Supplementary S6). In foxes, selection for road proximity depended on human activity: during low-activity times, foxes selected road-adjacent areas, especially in highly modified habitats, whereas road proximity was weakly avoided or near neutral during high-activity times (Figure 5a). Foxes avoided POIs during both human activity times, with stronger avoidance under higher human modification (Figure 5b). Tree-cover selection shifted from avoidance in less modified habitats to weaker avoidance or selection in highly modified habitats, most clearly during high human activity (Figure 5c).

**Figure 5:**
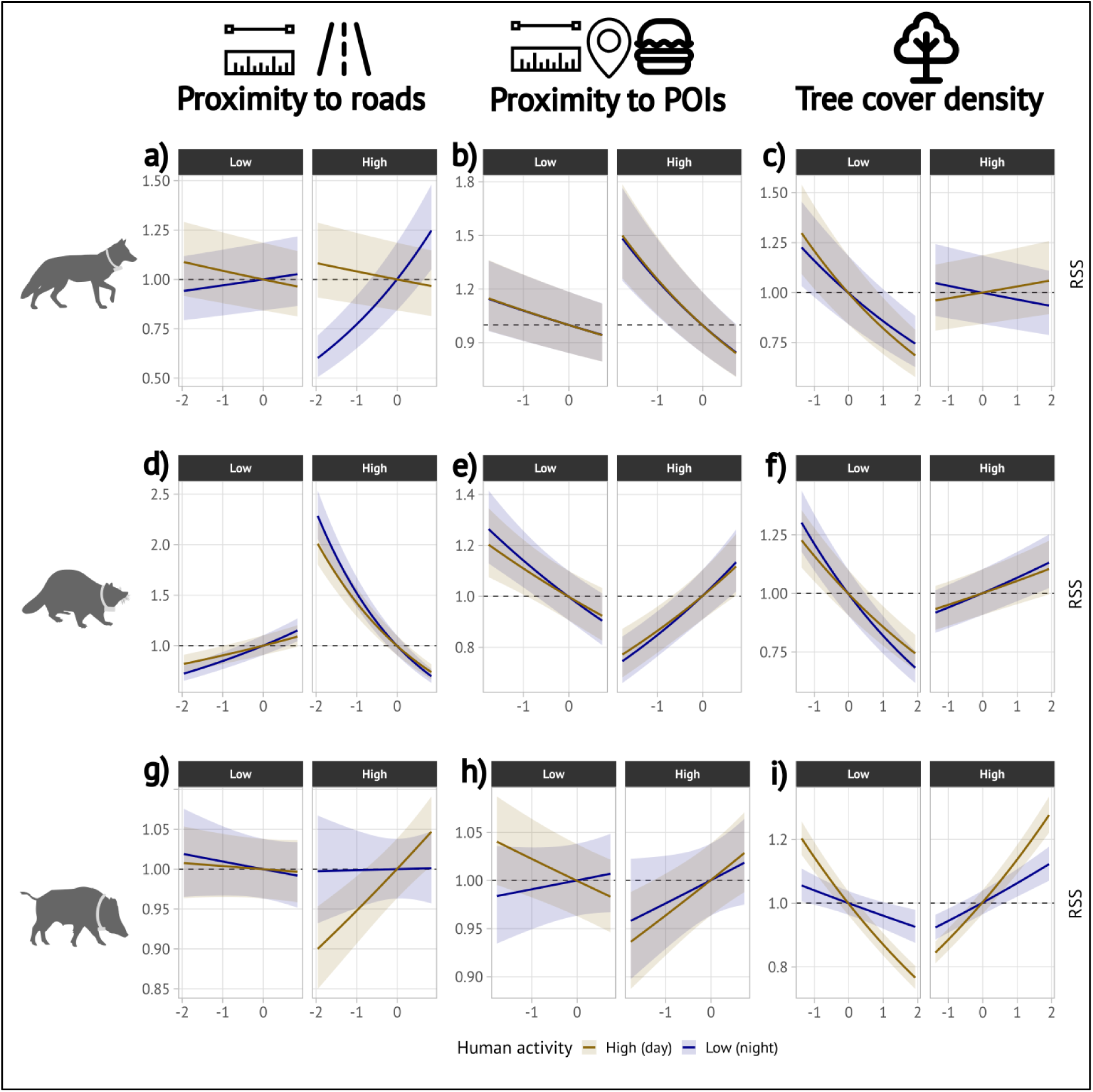
Species-specific spatial niche shifts along the human-modification gradient. Integrated step-selection results for proximity to roads, proximity to points of interest (POIs), and tree-cover density during high human activity (day; yellow lines) and low human activity (night; blue lines). Rows correspond to foxes, raccoons, and wild boars. Higher proximity values indicate closer proximity to the feature; higher tree-cover values indicate denser cover. Panels show relative selection strength (RSS) across the observed covariate range, separately for low and high local human modification. Values above 1 indicate selection, values below 1 indicate avoidance, ribbons indicate 95% confidence intervals, and dashed lines indicate neutral selection (RSS = 1). For example, raccoons (d-f) increasingly avoided roads but selected POIs and tree cover in highly urbanised habitats

In raccoons, road proximity and POIs showed opposite responses. Raccoons selected road-adjacent areas in less modified habitats but increasingly avoided them with higher human modification during both human activity times (Figure 5d). Conversely, raccoons shifted from weak avoidance or neutral selection of POIs in less modified habitats to strong selection in highly modified habitats (Figure 5e). Tree-cover selection similarly shifted from avoidance to selection with increasing human modification (Figure 5f).

In wild boars, road responses depended more strongly on human activity. Road proximity was neutral during low human activity times, whereas wild boars selected road-adjacent areas during high human activity in urbanised habitats (Figure 5g). POI responses were weaker and showed no general pattern (Figure 5h). Tree-cover selection showed the clearest shift, from avoidance in less modified habitats to strong selection in highly modified habitats, especially during high human activity (Figure 5i).

## 4 Discussion

Urbanisation is often expected to alter animal movement and temporal activity (e.g. Gallo et al., 2022), but our results suggest a more fundamental process: urban environments may filter species according to their capacity to restructure movement around human-imposed risk–resource dynamics. Across our three focal species, we found distinct behavioural pathways rather than a uniform urban response. Increased periodicity with urbanisation suggested temporal alignment with predictable cycles of disturbance and resource availability, potentially allowing animals to mitigate the costs of human presence. Flexible temporal and spatial restructuring without stronger periodicity indicated another form of adjustment, where movement remained responsive without becoming more scheduled and regular in space. By contrast, the loss of periodic routines and limited temporal restructuring in more urbanised habitats suggested that some species may be less able to maintain structured movement under intense human influence. These contrasts suggest that urban success may depend on the ability to reorganise movement in response to human pressure.

In our study, the observed species-specific responses closely reflected where each species occurs along the urbanisation gradient. Foxes and raccoons, which not only tolerate urban life but also live and reproduce in the most intensely built-up areas, showed stronger behavioural restructuring through routine movements, temporal niche shifts, and flexible habitat use. In contrast, wild boars, which are largely restricted to the urban periphery, showed little evidence of such reorganisation. This pattern matches previous work on the distribution and urban ecology of these species (Kimmig et al., 2020; Louvrier et al., 2022; Scholz et al., 2020, 2024; Stillfried, Gras, Börner, et al., 2017). Despite these differences, all three species relied more strongly on habitats with higher tree cover density in urban areas, suggesting that urban green spaces serve as refuges and may be important components of wildlife persistence not only in cities but in human-dominated landscapes in general (e.g. Oeser et al., 2023). More broadly, our findings highlight behavioural adjustment as a key mechanism through which wildlife can respond rapidly to environmental change (Gomez et al., 2025; Muhly et al., 2011; Potts et al., 2025; Russo et al., 2026).

### 4.1 Routine scheduling as one pathway of urban persistence

One pathway of urban persistence may involve increasingly routine behaviours, as exemplified by foxes, which structured their movements into regular travel routes visited at regular times as resources and risks became more predictable. Such scheduled movement routines were previously described in urban coyotes (*Canis latrans*) (Péron et al., 2017), and may enable repeated visits to profitable sites, consistent with memory-based foraging strategies that emerge when resources are spatially stable (Ranc et al., 2021; Robira et al., 2021). Anthropogenic food sources such as urban waste are often predictable in both space and time, favouring routine movements between known resource patches (Beck et al., 2026), as observed in urban gulls (e.g., Spelt et al., 2021) or brown bears (*Ursus arctos*) (Selva et al., 2017). Foxes are known to consume anthropogenic waste (Williams et al., 2025). In contrast, weaker periodicity in rural habitats may reflect the greater importance of hunting mobile prey, where successful foraging may require flexible search behaviour rather than repeated visits to fixed sites (Mueller & Fagan, 2008). This is also reflected in the foxes’ activity budget: rural foxes remained more active also during daytime hours. Hunting natural prey may require longer activity periods to meet energetic demands (Norberg, 2021; Pyke, 2019). At the same time, disturbances in rural landscapes – in contrast to predictable risks in urban home ranges – may be more closely tied to roads and commuting periods, which could allow foxes to remain active when away from these risks.

However, urban foxes also showed more pronounced diel structuring, i.e. foxes in urban areas were more nocturnal, and had more pronounced shifts in habitat selection across the diel cycle, particularly in highly urbanised areas, as corroborated by Alatawi (2025). In these environments, individuals strongly selected roads at night but avoided them during the day. Roads may provide efficient movement corridors that allow foxes to travel quickly between resource patches when traffic levels are lower and human disturbance is reduced (Bischof et al., 2019; S. E. Kimmig et al., 2020; Oehler et al., 2025). At the same time, urban streets often include sidewalks and adjacent infrastructure that may also provide access to anthropogenic food sources, meaning that road use may reflect both commuting behaviour and foraging opportunities. Roads may also serve as potential scavenging sites (Fielding et al., 2021; Planillo et al., 2015). Such behaviour likely comes with risks, as vehicle collisions represent one of the main sources of mortality and severe injury in urban fox populations (Börner, 2014; Harris, 1978), highlighting a trade-off between resource access and mortality risk. Avoidance of human disturbance agrees with previous work showing that foxes reduce diurnal activity where human disturbance is high (Díaz-Ruiz et al., 2016). Taken together, our results suggest that urban foxes organise movement and habitat use more consistently around predictable daily patterns of resources and risk, supporting routinisation as one pathway of urban persistence. This interpretation is further supported by the fact that models based on fixed-clock time gained in predictive performance in highly urbanised foxes, suggesting temporal alignment under increasing human pressures (see Supplementary S4).

### 4.2 Behavioural flexibility as an alternative pathway of urban persistence

Raccoons, in contrast to red foxes, showed higher flexibility and less periodic behaviour, more temporal activity shifts, and flexible habitat selection, all of which may allow them persist in urban environments without aligning spatiotemporal routines. Such increased flexibility and problem-solving skills have been observed before in wild raccoons (Stanton et al., 2024). While anthropogenic food availability and dietary flexibility are likely important, these factors alone may not explain the observed differences, as red foxes show similar opportunistic feeding strategies but different movement patterns. One possible explanation is that raccoons may exploit resources more flexibly due to their increased manual dexterity, allowing them to rely less on routine movements (Lazure & Weladji, 2024; Stanton et al., 2022). Their ability to manipulate objects with their forepaws allows them to access resources directly, for example, by opening waste containers, enabling exploitation of a wider range of anthropogenic food and natural sources (Knight, 2022). At the same time, raccoons may mitigate risks in other ways. For example, the raccoon’s ability to use vertical escape routes thanks to their climbing abilities may allow for more flexible risk avoidance also during the day. This third dimension in animal movement is often overlooked (Péron et al., 2020), but hypothesised to promote urban success in other species, such as red squirrels (Drenske et al., 2024; Grabow et al., 2022). Such behavioural flexibility has also been identified as an important predictor of invasion success in animals (Sol et al., 2002; Sol & Lefebvre, 2000), and may play an important role in raccoons.

Raccoons were also less strictly nocturnal in urban areas, indicating a relaxation of the typically nocturnal activity pattern reported for this species and contrasting with studies showing increased nocturnality in urban mammals (Burton et al., 2024; Gaynor et al., 2018). Habitat-selection patterns further support this interpretation. Unlike foxes, raccoons increasingly avoided roads in more human-modified habitats during both day and night, suggesting consistent avoidance of traffic-related risk (Barthelmess & Brooks, 2010). At the same time, selection for proximity POIs shifted from avoidance or weak use in less modified habitats to strong selection in highly modified habitats, indicating that raccoons increasingly oriented their movements towards human-associated resource hotspots where such resources are most concentrated. This combination of road avoidance, POI selection, and reduced nocturnality suggests flexible risk–resource tracking rather than stronger routinisation, allowing raccoons to persist in highly urbanised environments not by becoming more predictable, but by flexibly adjusting where and when they move.

### 4.3 Behavioural constraints limiting urban persistence

Wild boars showed a contrasting pattern to foxes and raccoons, with reduced periodicity towards more urbanised areas and no temporal restructuring, suggesting that their movement routines become disrupted as individuals approach human-dominated environments (Doherty et al., 2021) (Palmer et al., 2022). Wild boars primarily rely on rooting for below-ground resources, a foraging strategy that is difficult to perform in sealed or heavily modified urban substrates (Stillfried, Gras, Busch, et al., 2017). If present, these natural resources are spatially patchy and temporally unpredictable, which should favour exploratory foraging rather than repeated visits to fixed locations (Mueller & Fagan, 2008). In addition, wild boars rely less on fixed den or resting sites in the same way as red foxes. Although they use recurring resting locations, these can shift frequently, reducing the need for repeated returns to a central place (Fradin & Chamaillé-Jammes, 2023). However, our finding are in contrast with recent findings on wild boar resting site selection, where distinct resting sites were often revisited in urbanised areas (Fradin et al., 2025). Together, these ecological characteristics may already weaken the emergence of strong movement routines, particularly when individuals enter environments that are more disturbed.

This finding may reflect the situation of the study area, where wild boars primarily occur in semi-urban habitats at the city outskirts. Here, human activity is variable, and dogs frequently roam freely, explaining the counterintuitive daytime selection for road proximity: dog owners do not let their dogs unleashed near major roads, and no hunting occurs. In this context, this counterintuitive daytime selection for road proximity by wild boars in highly modified habitats may mark a threshold in their capacity for urban persistence: when refuge habitat becomes scarce, individuals may be forced into less predictable and riskier movements through the urban matrix, even during times of high human activity. Previous studies have also reported disrupted resting and sleep patterns in wild boar facing higher human activity pressure (Olejarz et al., 2023), suggesting that individuals remain in a heightened state of alertness (Ritzel & Gallo, 2020), which could further destabilise regular movement schedules.

To conclude, our results reveal species-specific pathways of urban persistence. Foxes appear to persist in urban environments through increasing routinisation of movement, raccoons through flexible adjustment of activity timing and habitat use, whereas wild boars showed weaker behavioural adjustment and remained largely confined to the urban periphery. Across all three species, urban habitats were associated with stronger reliance on refuge habitats, underscoring the importance of urban green spaces for wildlife persistence in cities (Parsons et al., 2019). While experimental and longitudinal studies are needed to distinguish behavioural plasticity from urban filtering and to test the fitness consequences of periodic movement, our findings suggest that urban wildlife communities are shaped by the capacity to reorganise movement across space and time while maintaining access to refuge habitat.

## Author contributions

MG and SKS designed the study and methodology. CS, MR, MS, SKi, CW and KB collected animal tracking data. NB, MR and SKS curated data. NB, FJ, SO, and SKS provided project administration and supervision. FJ, SO and SKS secured funding. MG analysed movement data. MG led writing of the manuscript. All authors edited the manuscript and made valuable scientific contributions throughout the writing process.

## Acknowledgements

We thank the veterinarians Frank Göritz, Johanna Painer, Kathleen Röllig, Janina Radwainski for support in animal capture and Anne Berger for support in e-obs collar programming. We thank Moritz Wenzler-Meya for preparing geodata.

## Animal ethics

All animal procedures were conducted under the following permits, issued by the Landesamt für Gesundheit und Soziales (LAGeSo) Berlin or Landesamt für Umwelt, Gesundheit und Verbraucherschutz (LfU) Brandenburg, Germany:

*Vulpes vulpes* (LfU Brandenburg): V3-2347 13-2011, V3-2347-13-2011Ä5, 2347-25-2015

*Vulpes vulpes* (LaGeSo Berlin): G 0211/15

*Procyon lotor* (LfU Brandenburg): 2347-7-2020

*Procyon lotor* (LaGeSo Berlin): G 0101/16

*Sus scrofa* (LfU Brandenburg): V3-2347-40-2012

*Sus scrofa* (LaGeSo Berlin): G 0383/12

