## Supplementary S1 for "Movement strategies reveal the success of mammals in urban areas"

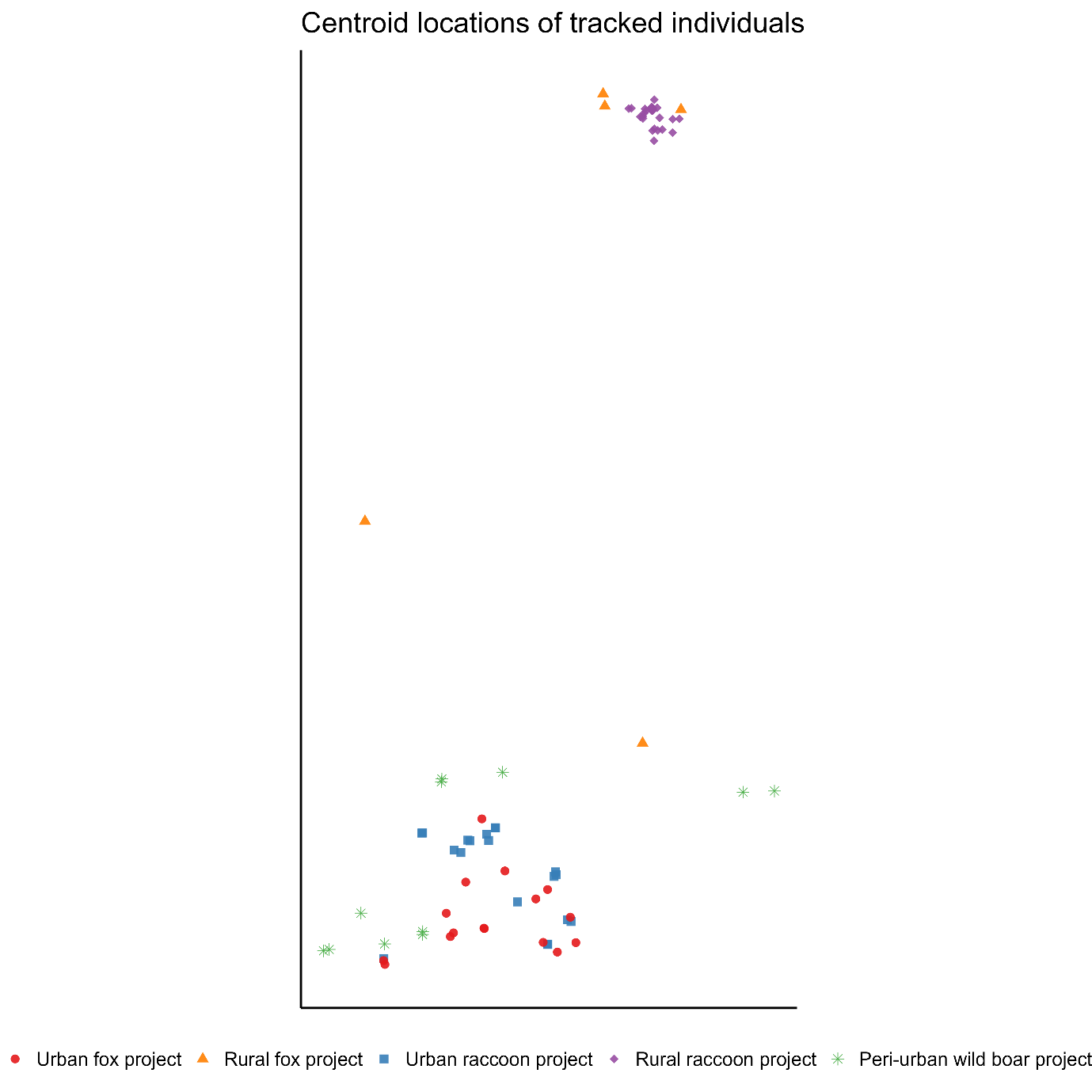

Figure S1.1: Centroid locations of tracked individuals by study project

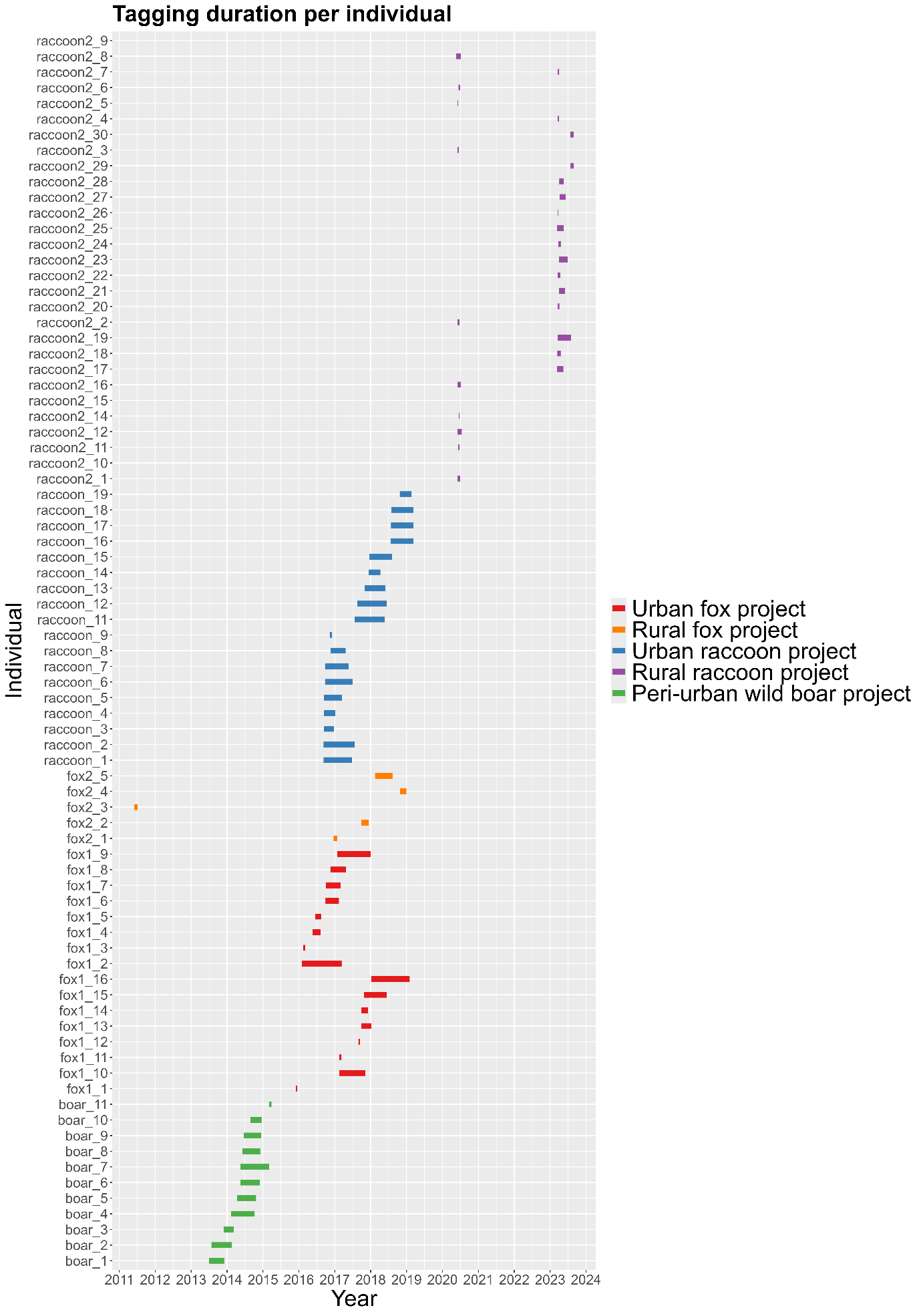

Figure S.1.2: Tagging duration of tracked individuals by study project
