## Supplementary S2 for "Movement strategies reveal the success of mammals in urban areas"

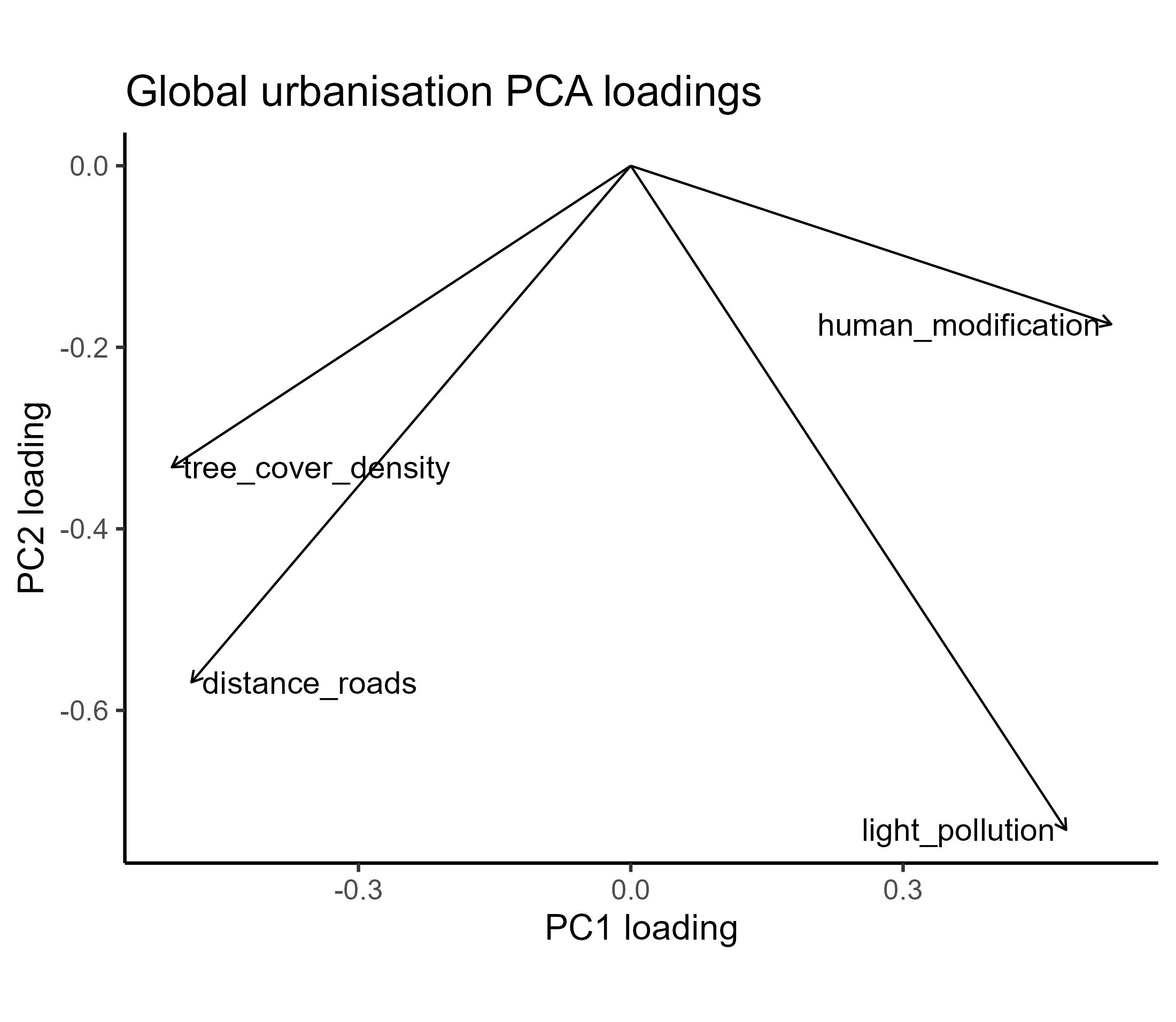

Figure S2.1: Plot of global urbanisation PCA loadings

Table S2.1: Table of global urbanisation PCA loadings

|  | PC1 | PC2 |
| --- | --- | --- |
| Tree_cover_density | -0.504 | 0.320 |
| Human_modification | 0.522 | 0.198 |
| Distance_roads | -0.488 | 0.590 |
| Light_pollution | 0.484 | 0.714 |
