## Supplementary S3 for "Movement strategies reveal the success of mammals in urban areas"

Supplementary diagnostic periodograms

Each figure shows the diagnostic periodogram for one individual-month time series.

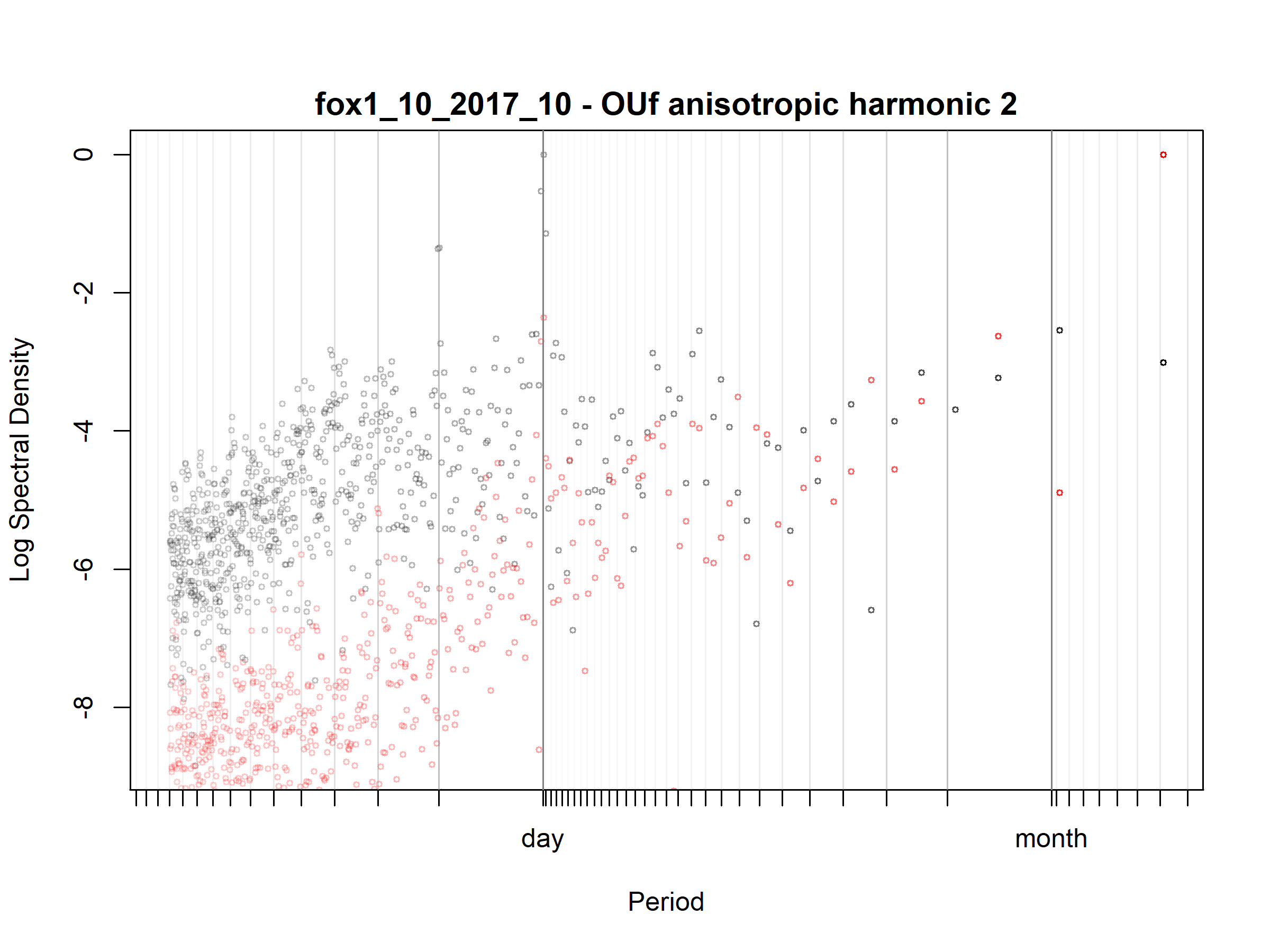

Figure S3.1. Fox; fox1_10_2017_10; diagnostic periodogram.

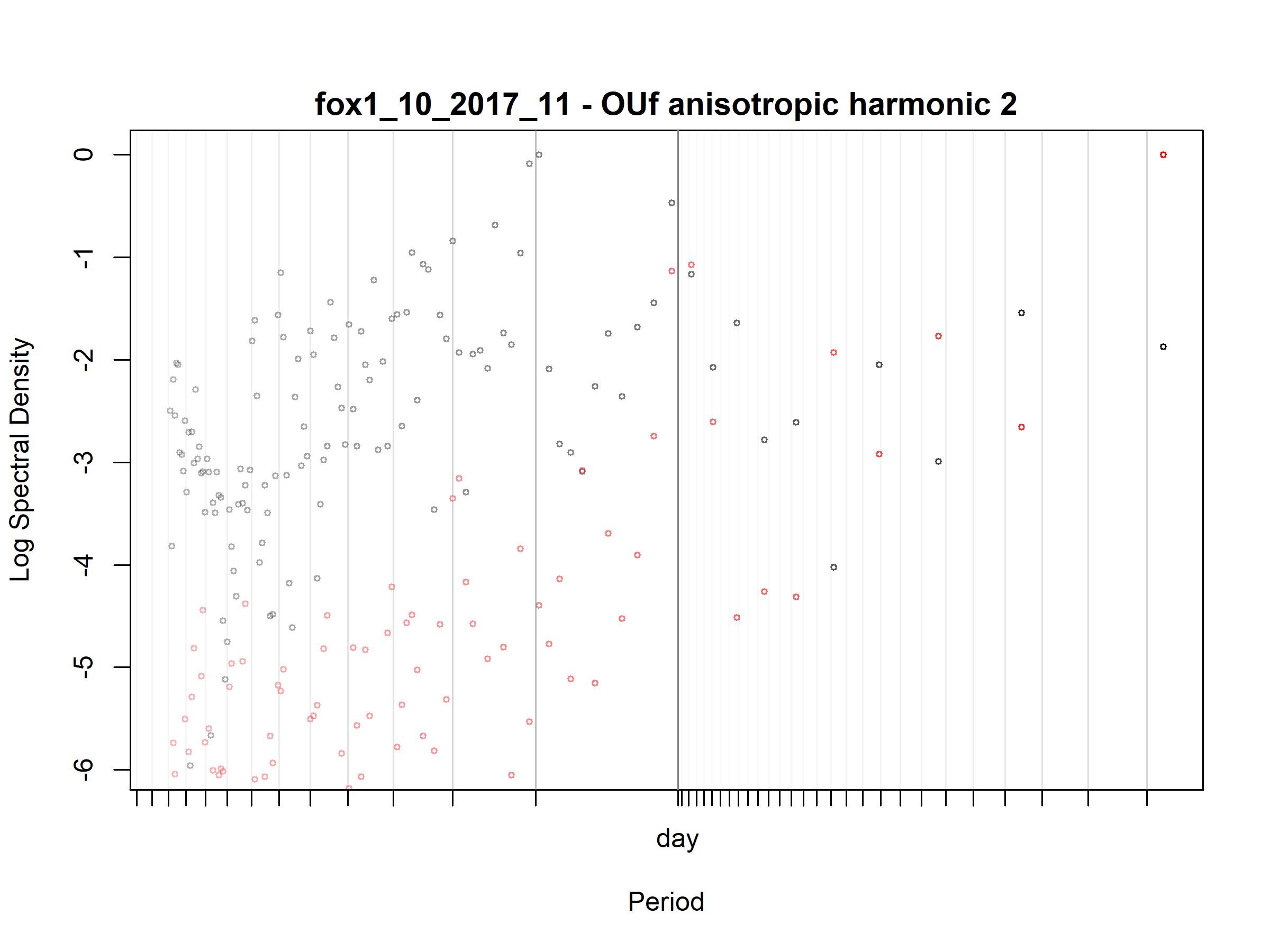

Figure S3.2. Fox; fox1_10_2017_11; diagnostic periodogram.

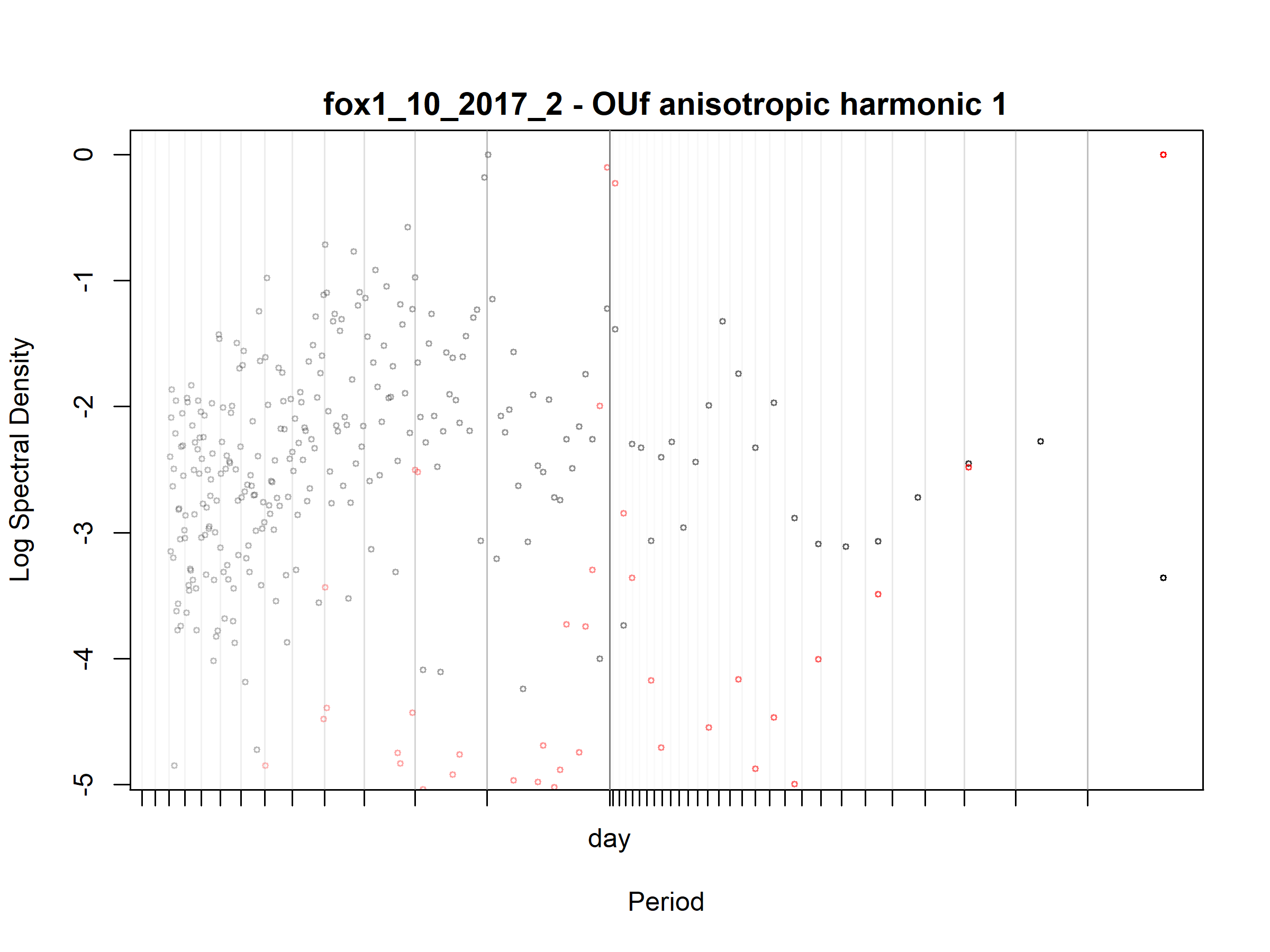

Figure S3.3. Fox; fox1_10_2017_2; diagnostic periodogram.

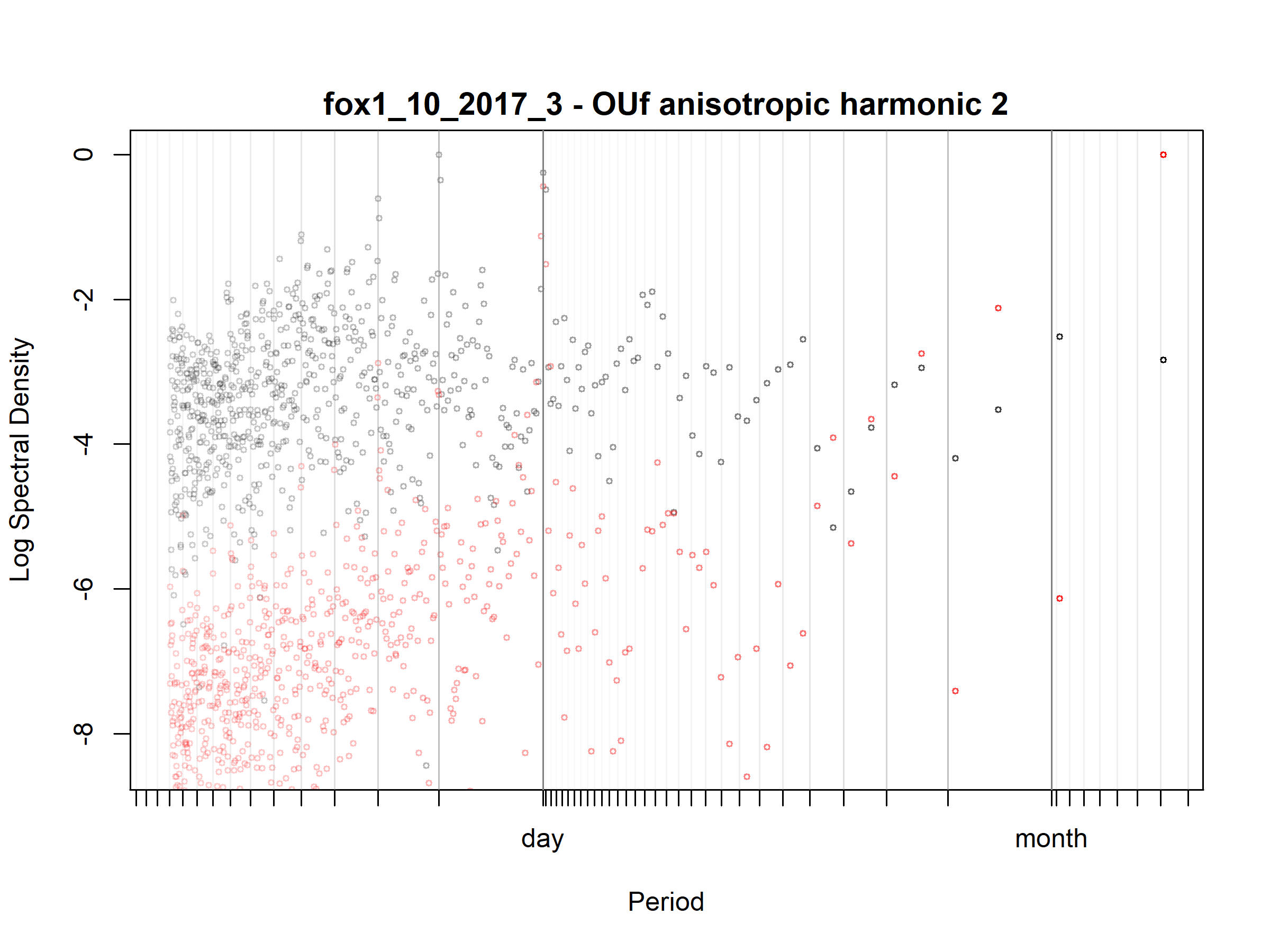

Figure S3.4. Fox; fox1_10_2017_3; diagnostic periodogram.

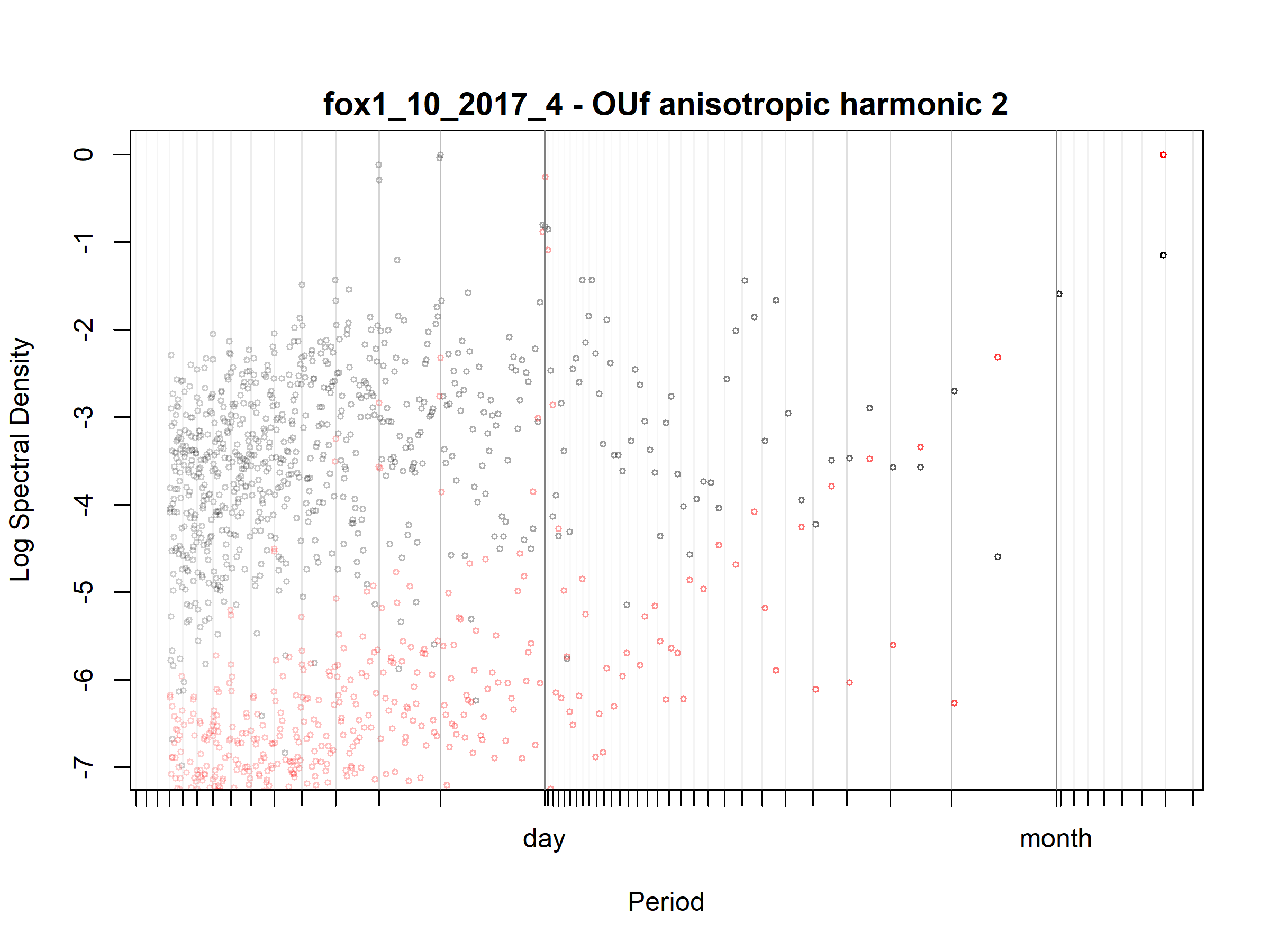

Figure S3.5. Fox; fox1_10_2017_4; diagnostic periodogram.

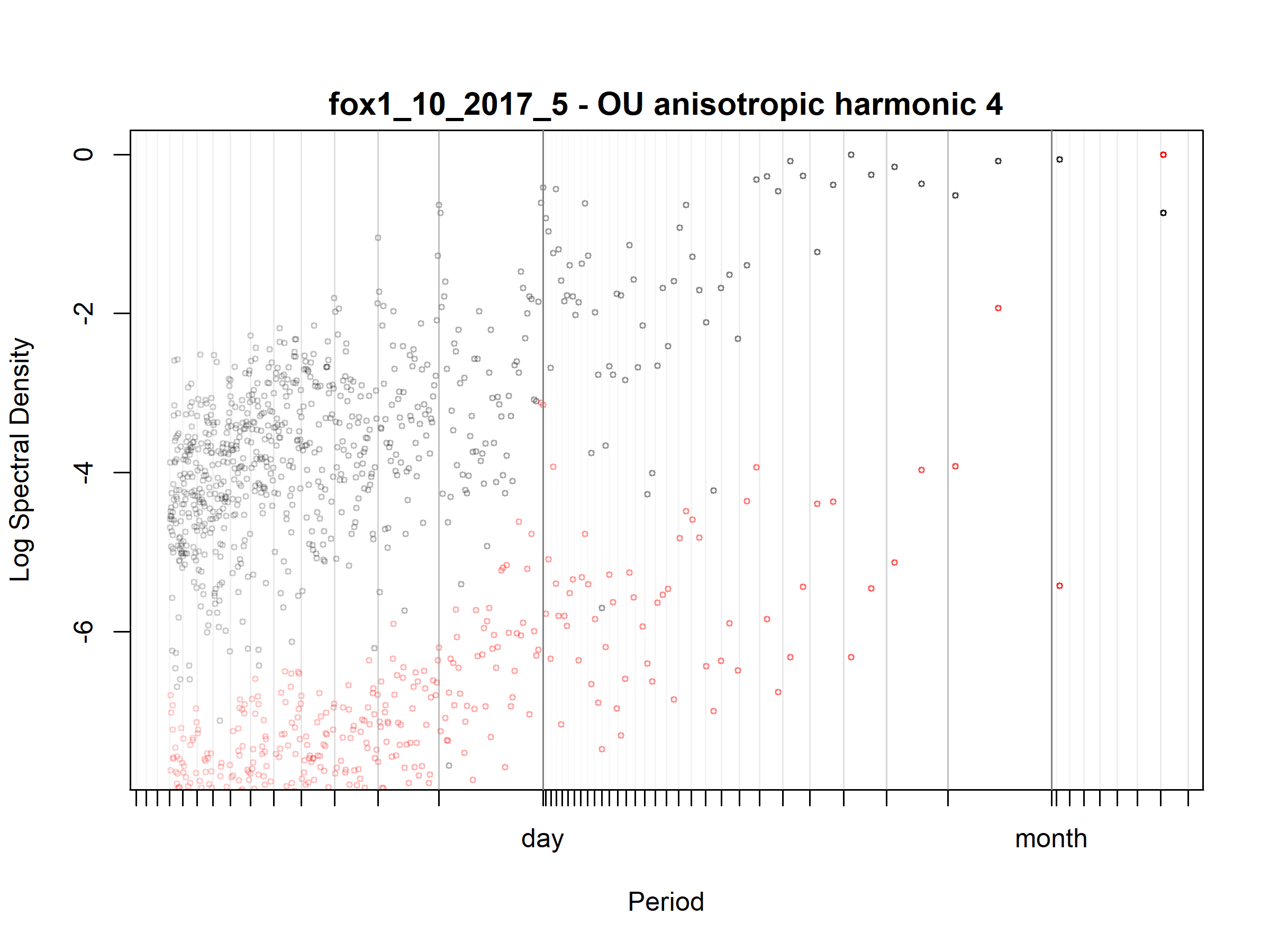

Figure S3.6. Fox; fox1_10_2017_5; diagnostic periodogram.

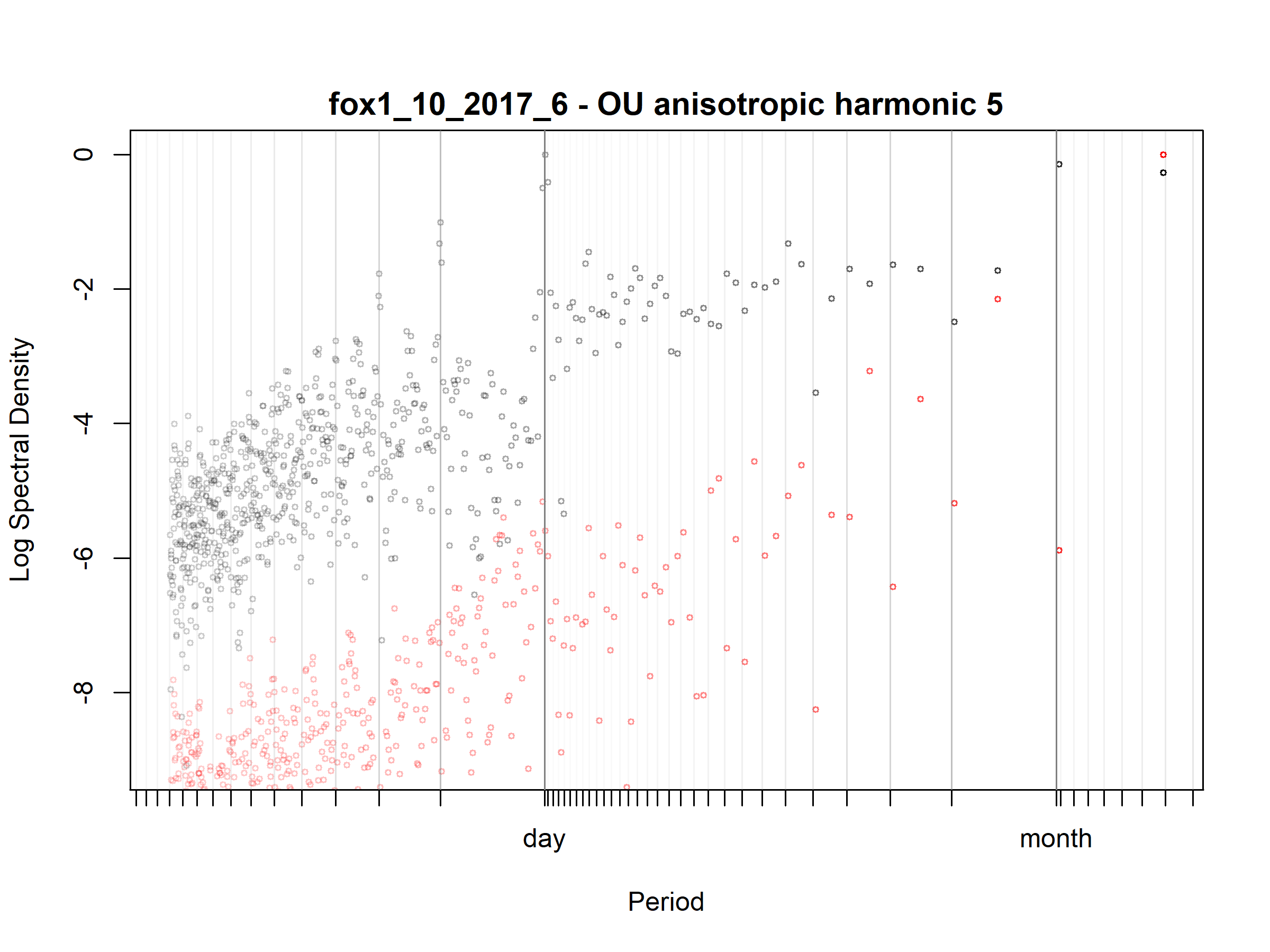

Figure S3.7. Fox; fox1_10_2017_6; diagnostic periodogram.

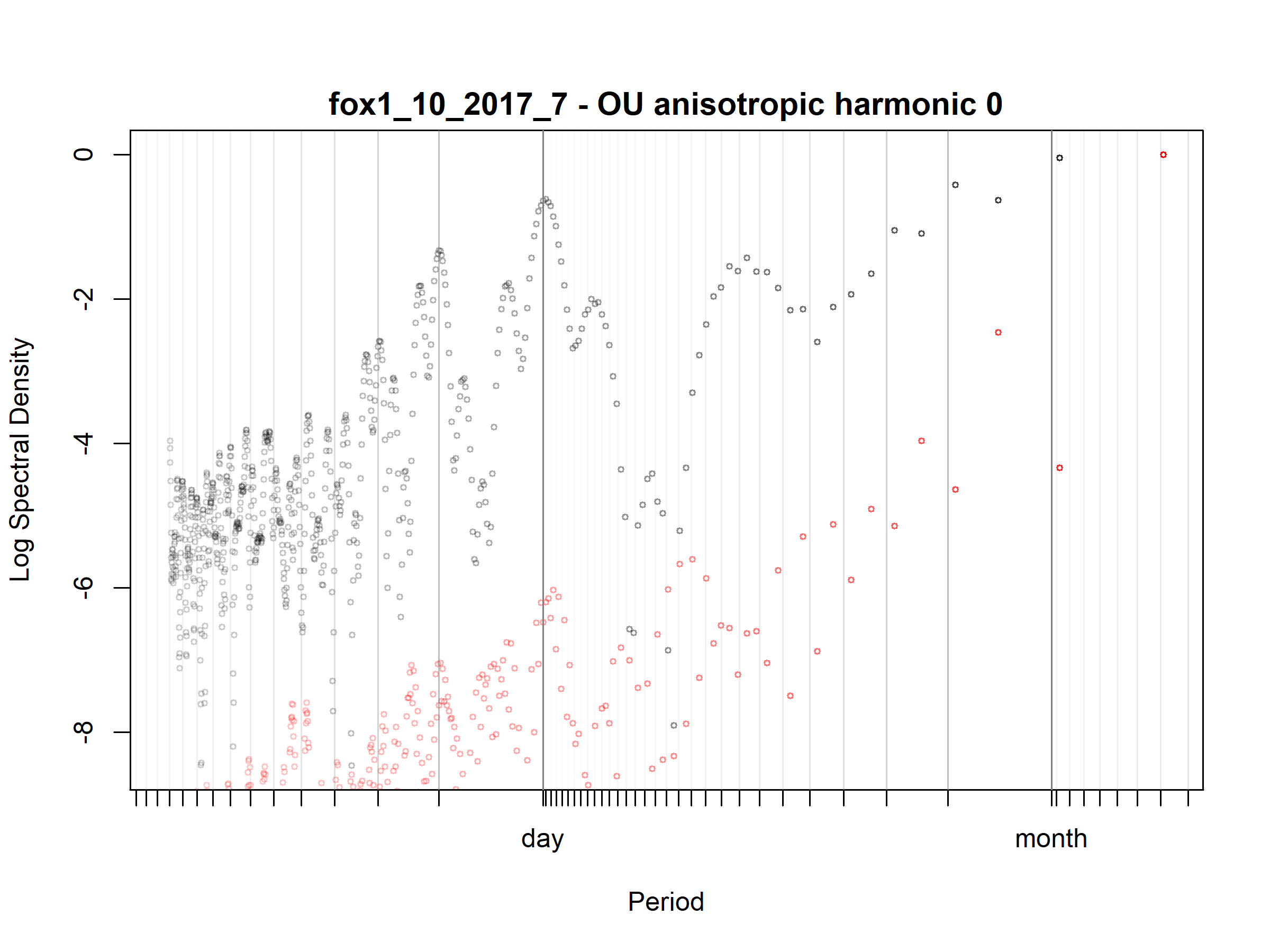

Figure S3.8. Fox; fox1_10_2017_7; diagnostic periodogram.

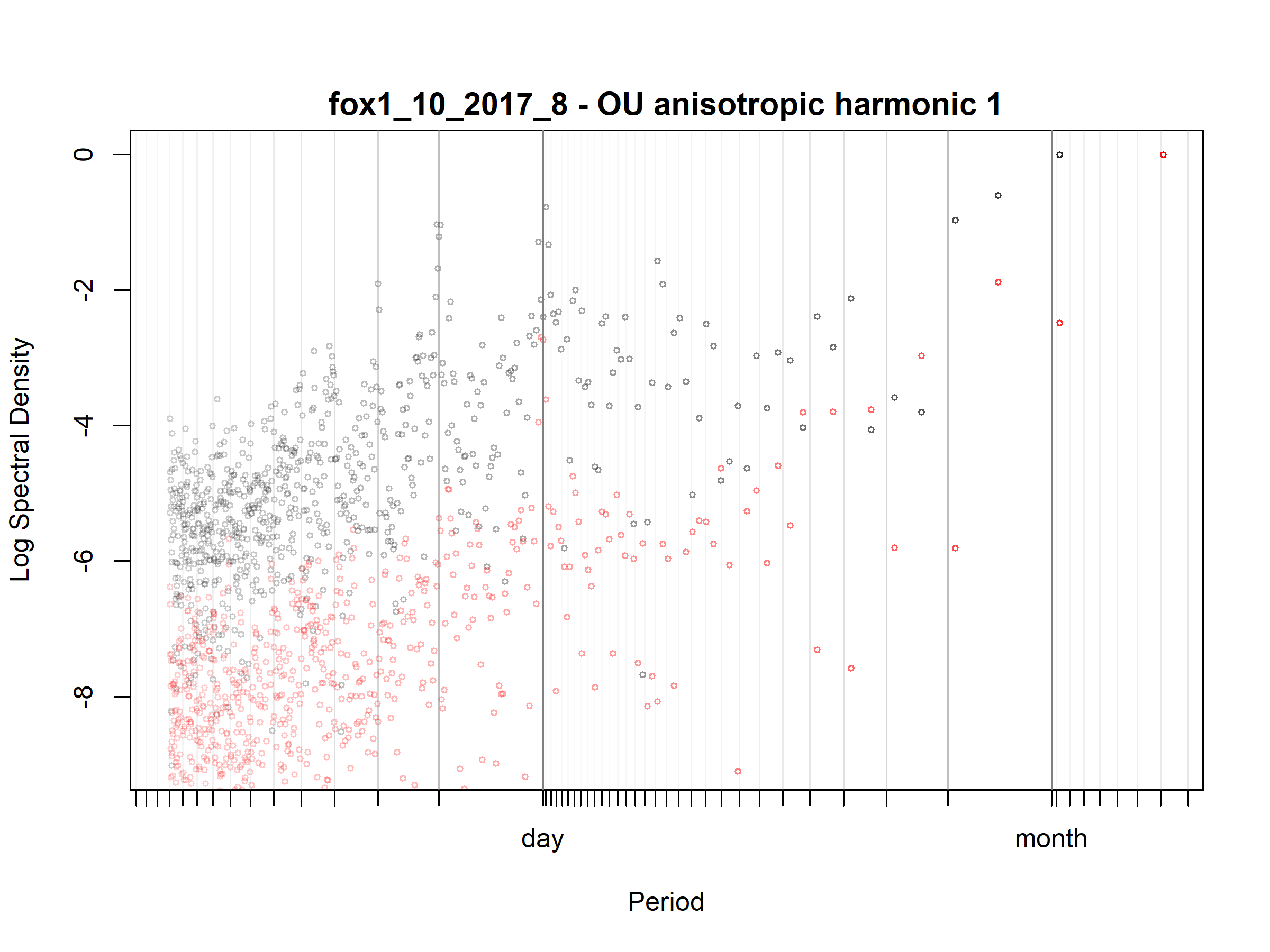

Figure S3.9. Fox; fox1_10_2017_8; diagnostic periodogram.

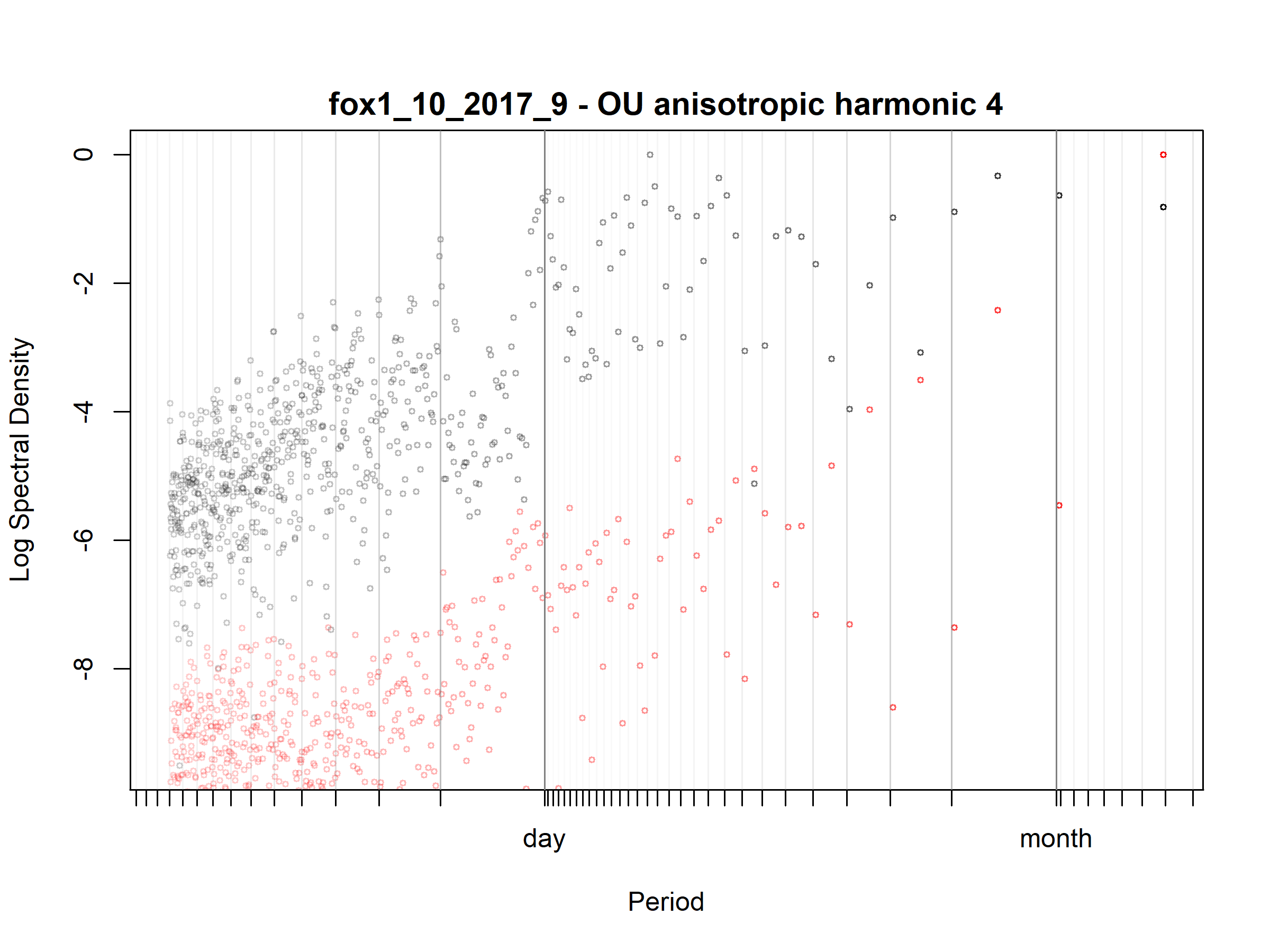

Figure S3.10. Fox; fox1_10_2017_9; diagnostic periodogram.

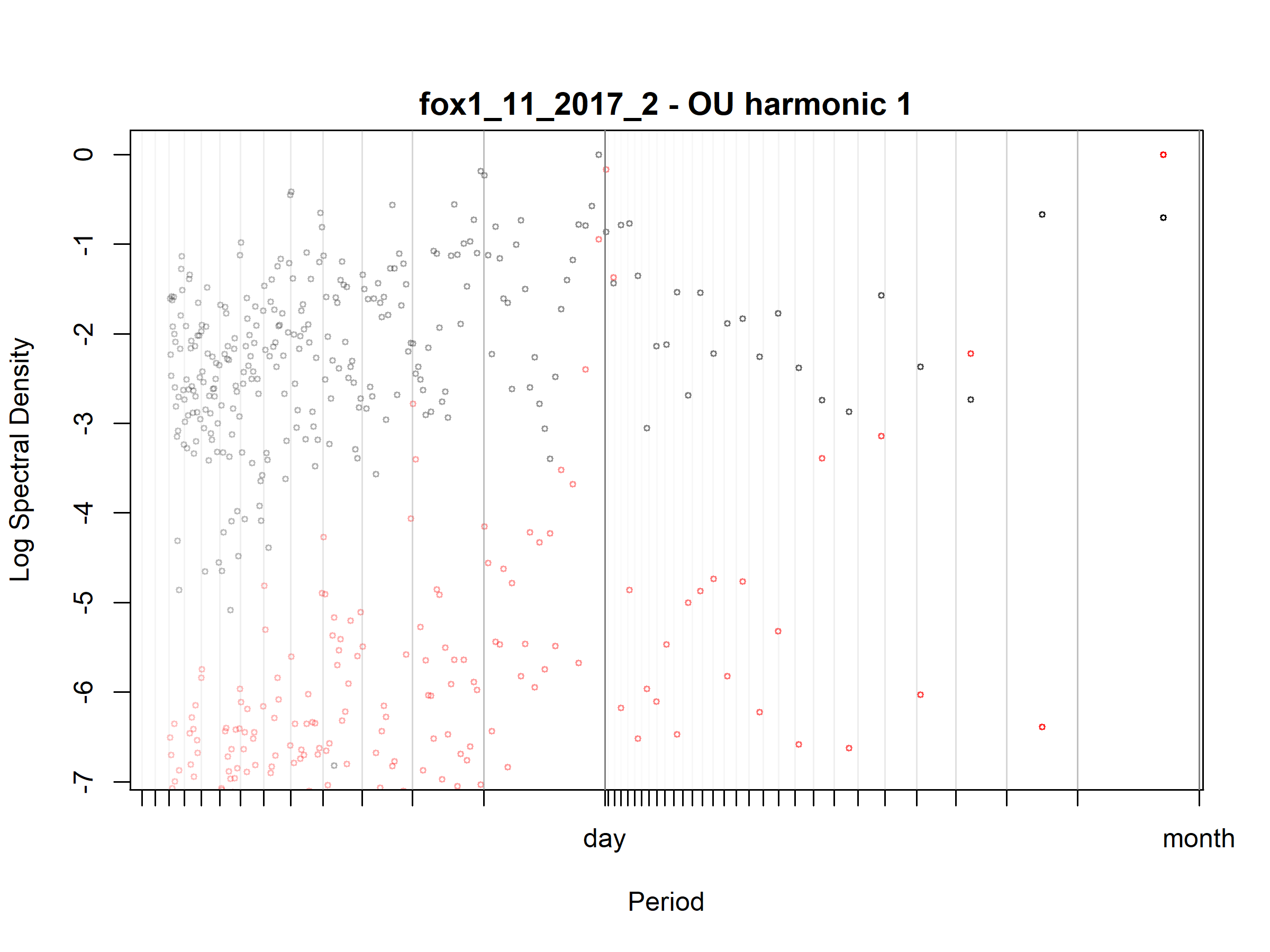

Figure S3.11. Fox; fox1_11_2017_2; diagnostic periodogram.

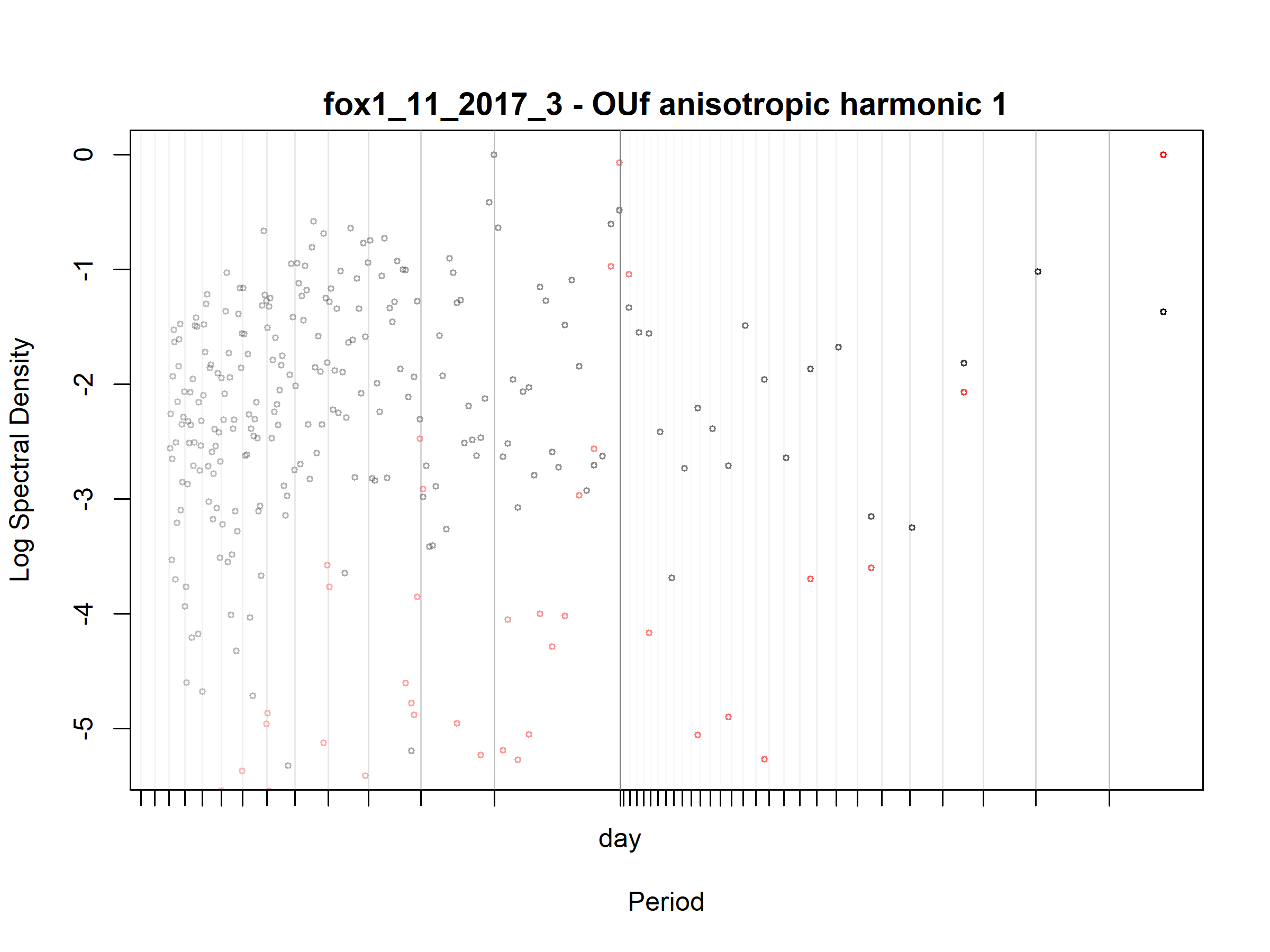

Figure S3.12. Fox; fox1_11_2017_3; diagnostic periodogram.

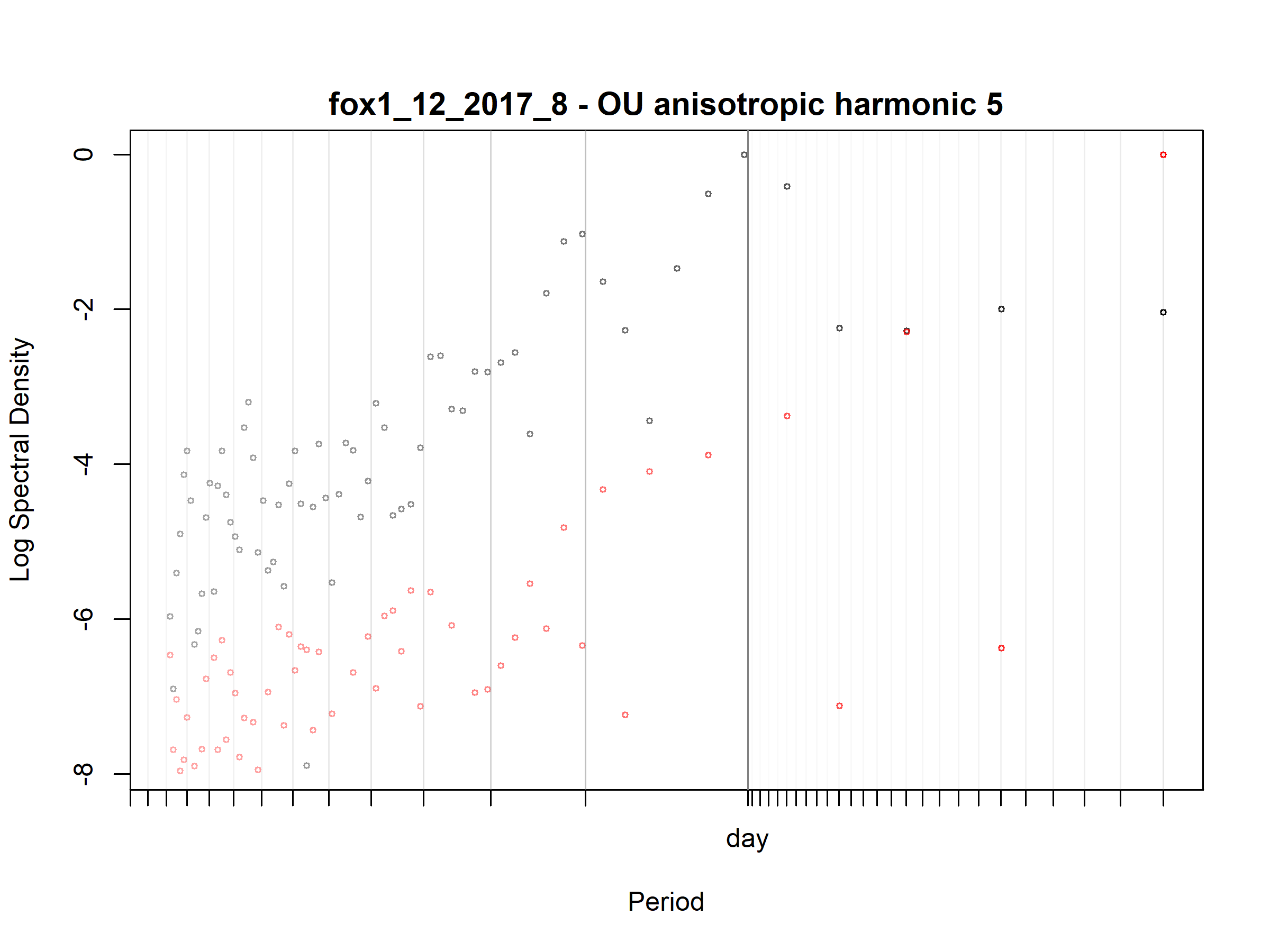

Figure S3.13. Fox; fox1_12_2017_8; diagnostic periodogram.

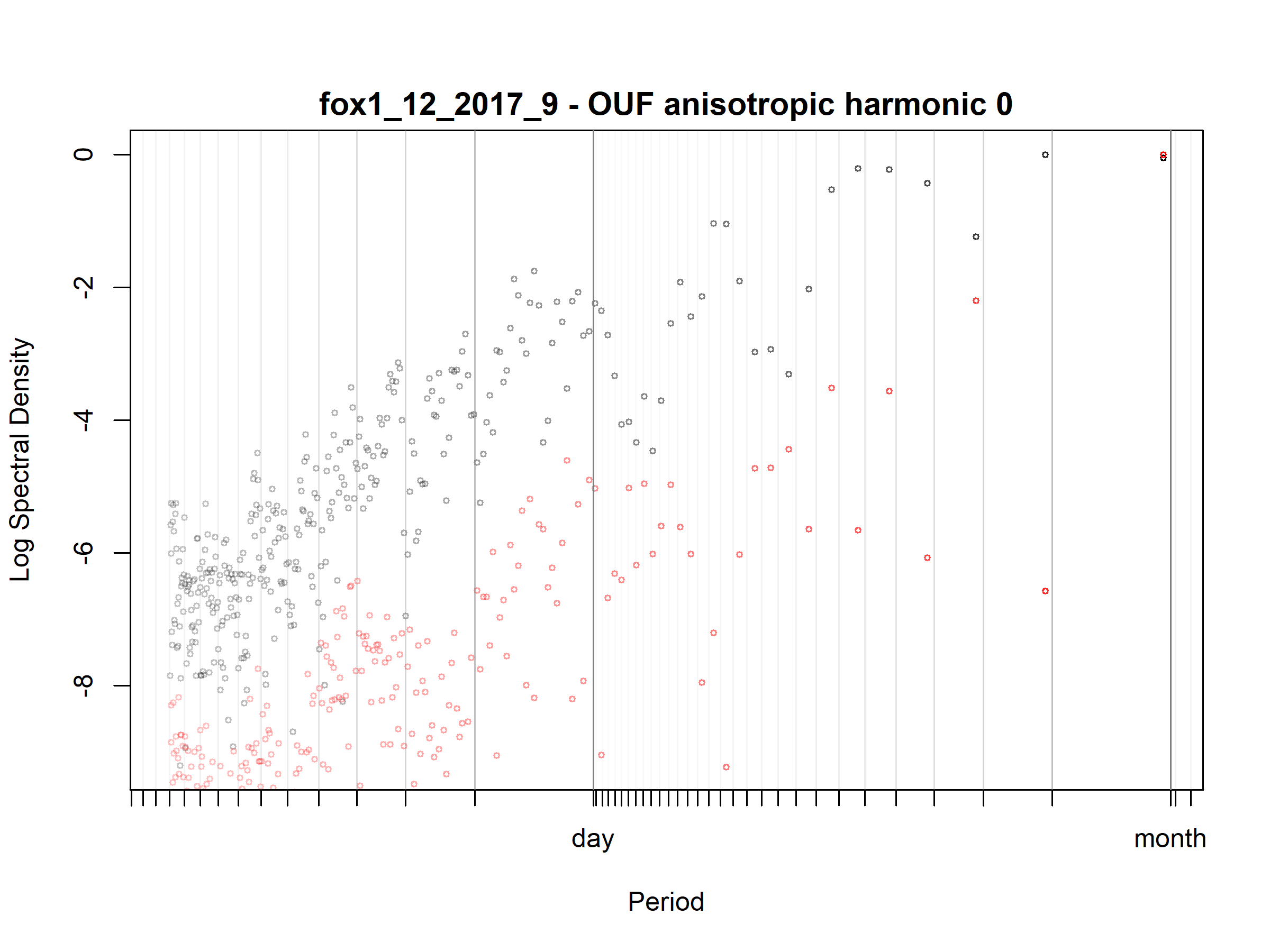

Figure S3.14. Fox; fox1_12_2017_9; diagnostic periodogram.

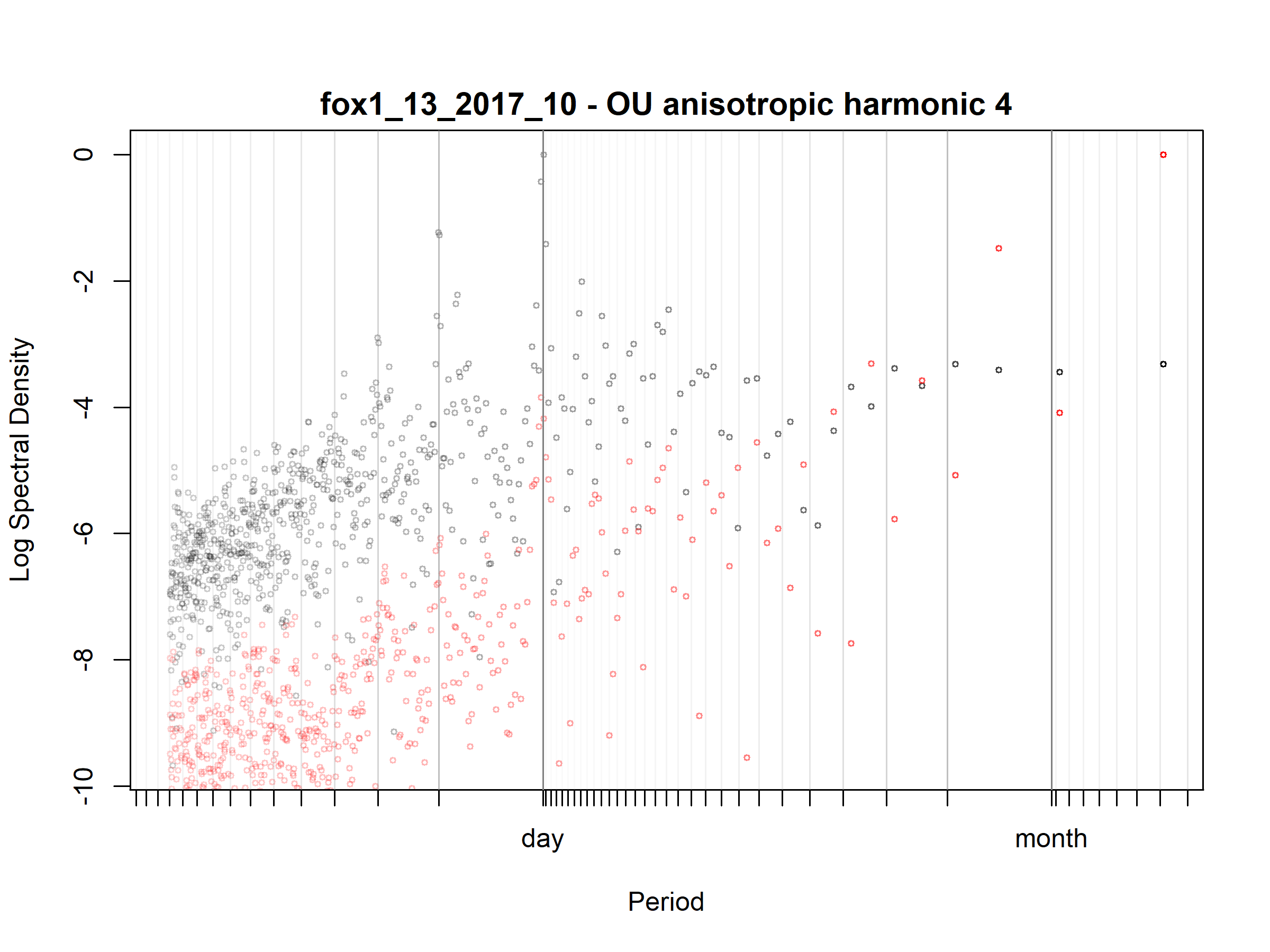

Figure S3.15. Fox; fox1_13_2017_10; diagnostic periodogram.

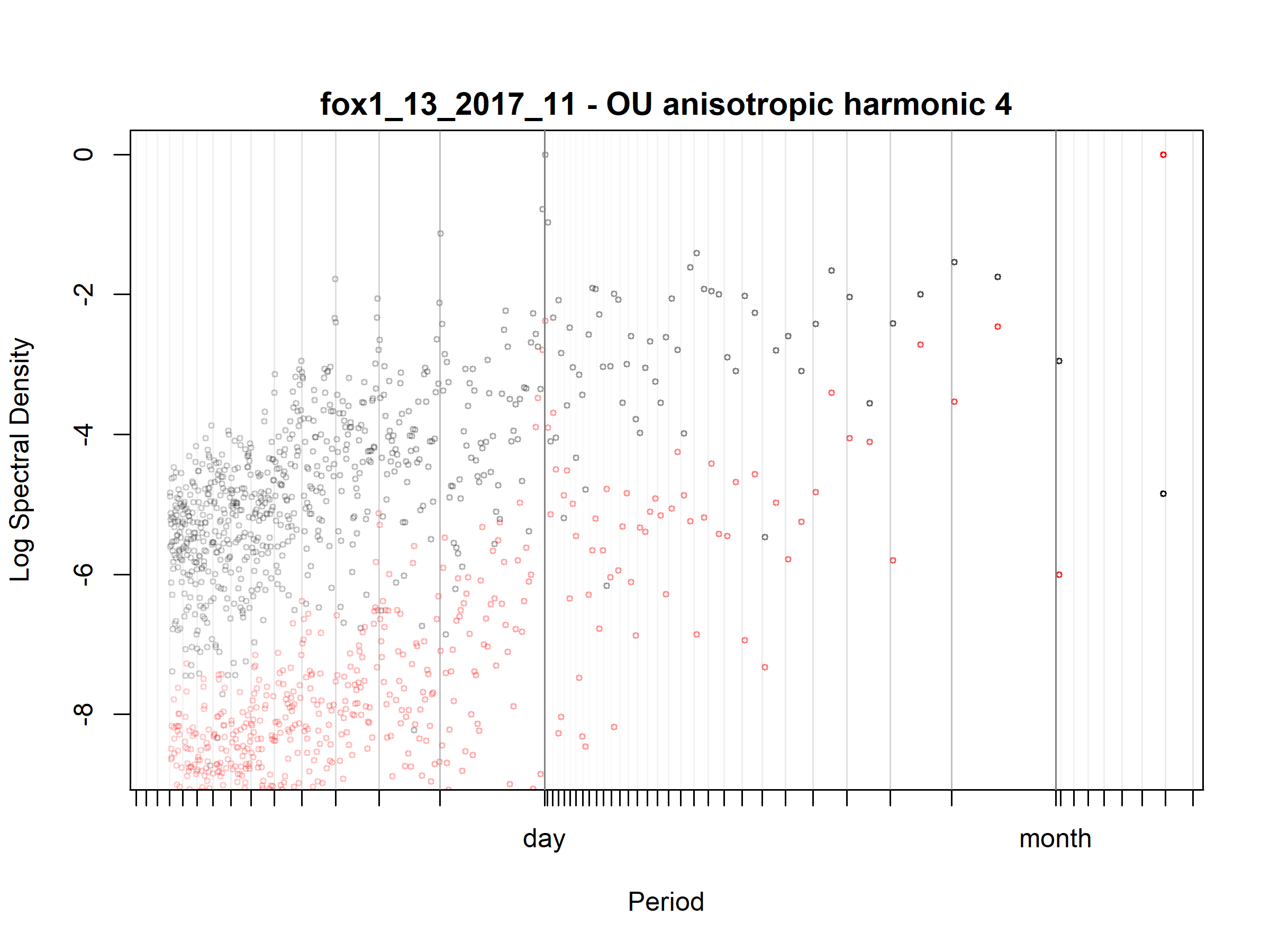

Figure S3.16. Fox; fox1_13_2017_11; diagnostic periodogram.

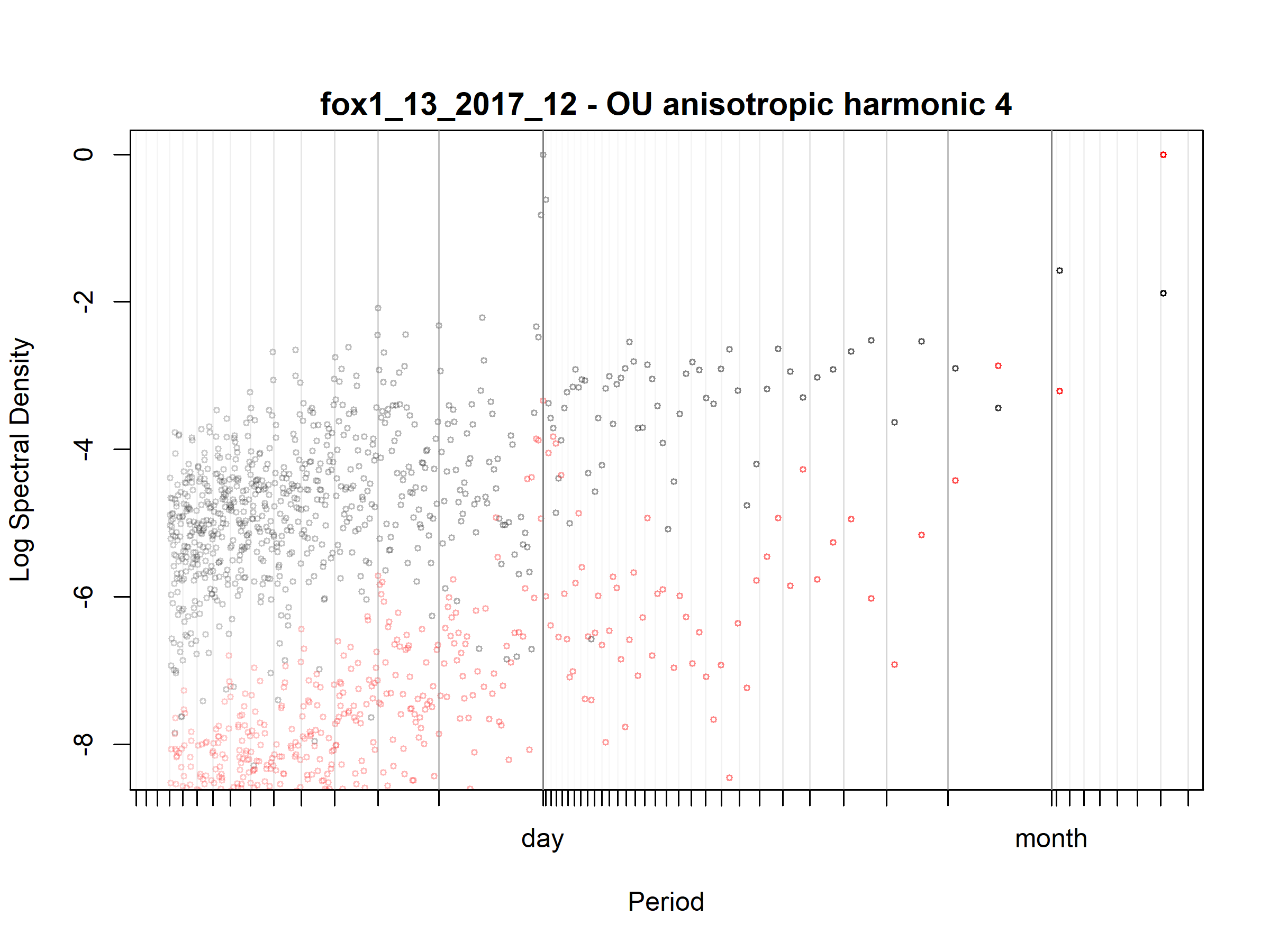

Figure S3.17. Fox; fox1_13_2017_12; diagnostic periodogram.

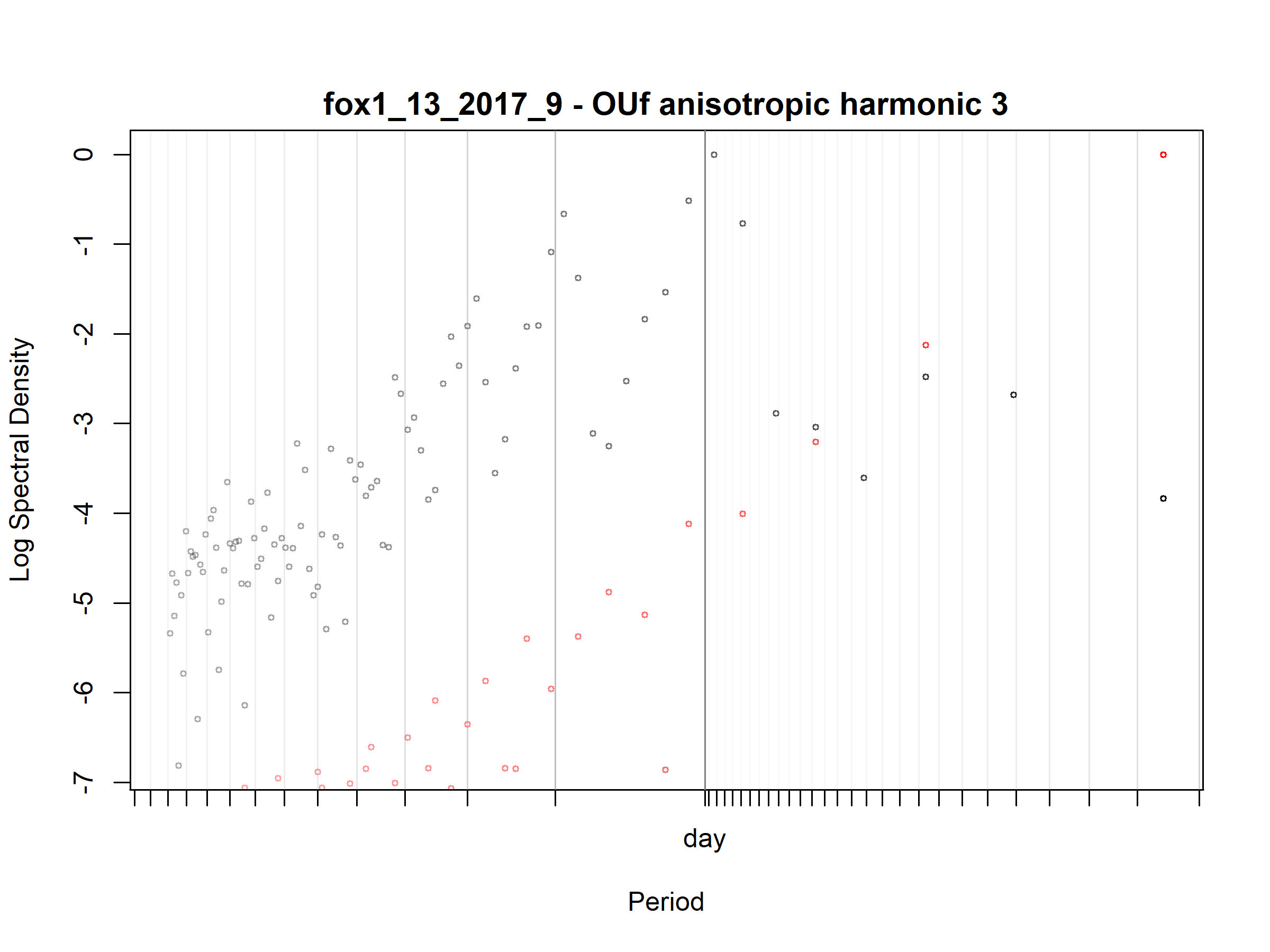

Figure S3.18. Fox; fox1_13_2017_9; diagnostic periodogram.

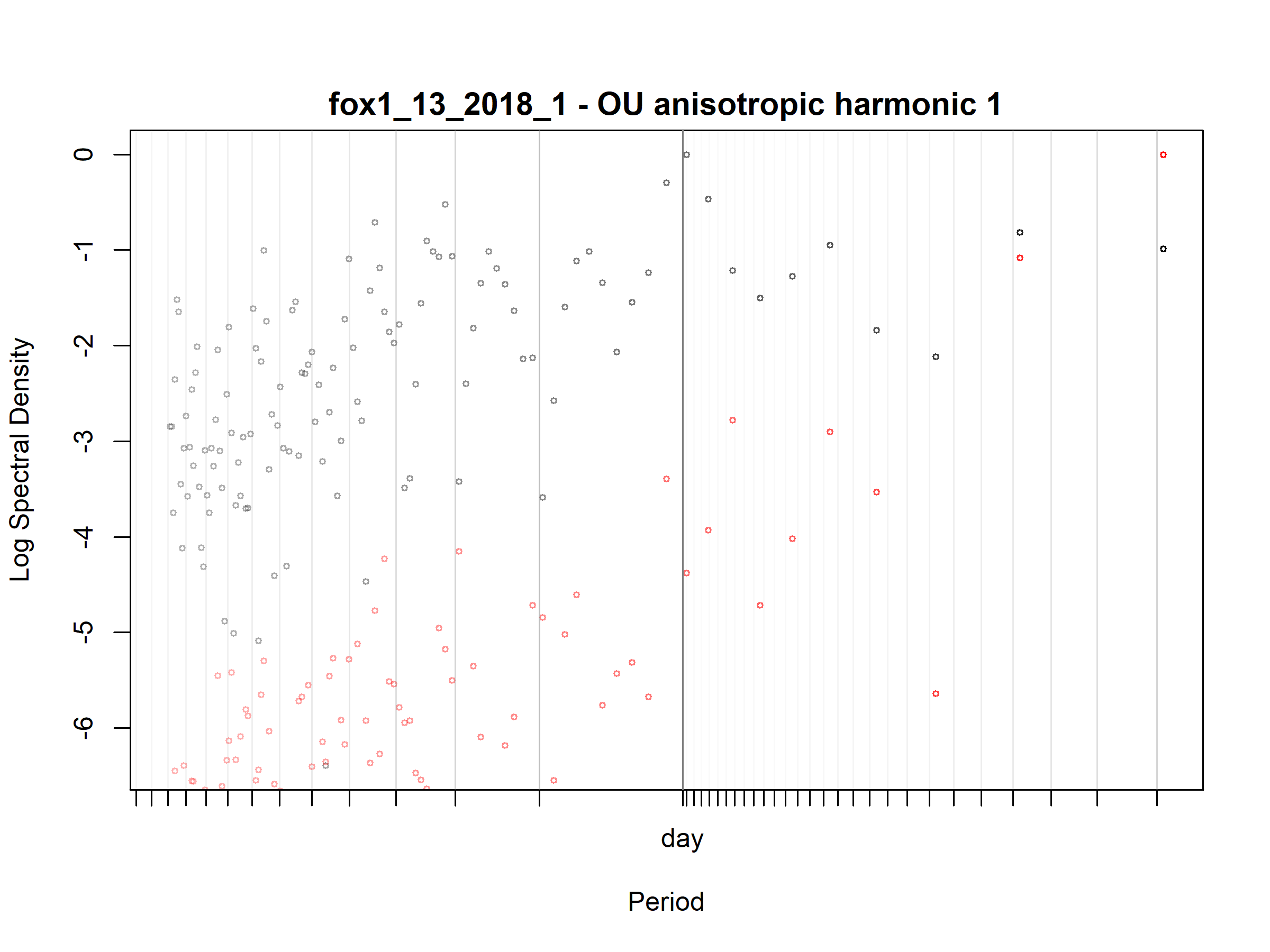

Figure S3.19. Fox; fox1_13_2018_1; diagnostic periodogram.

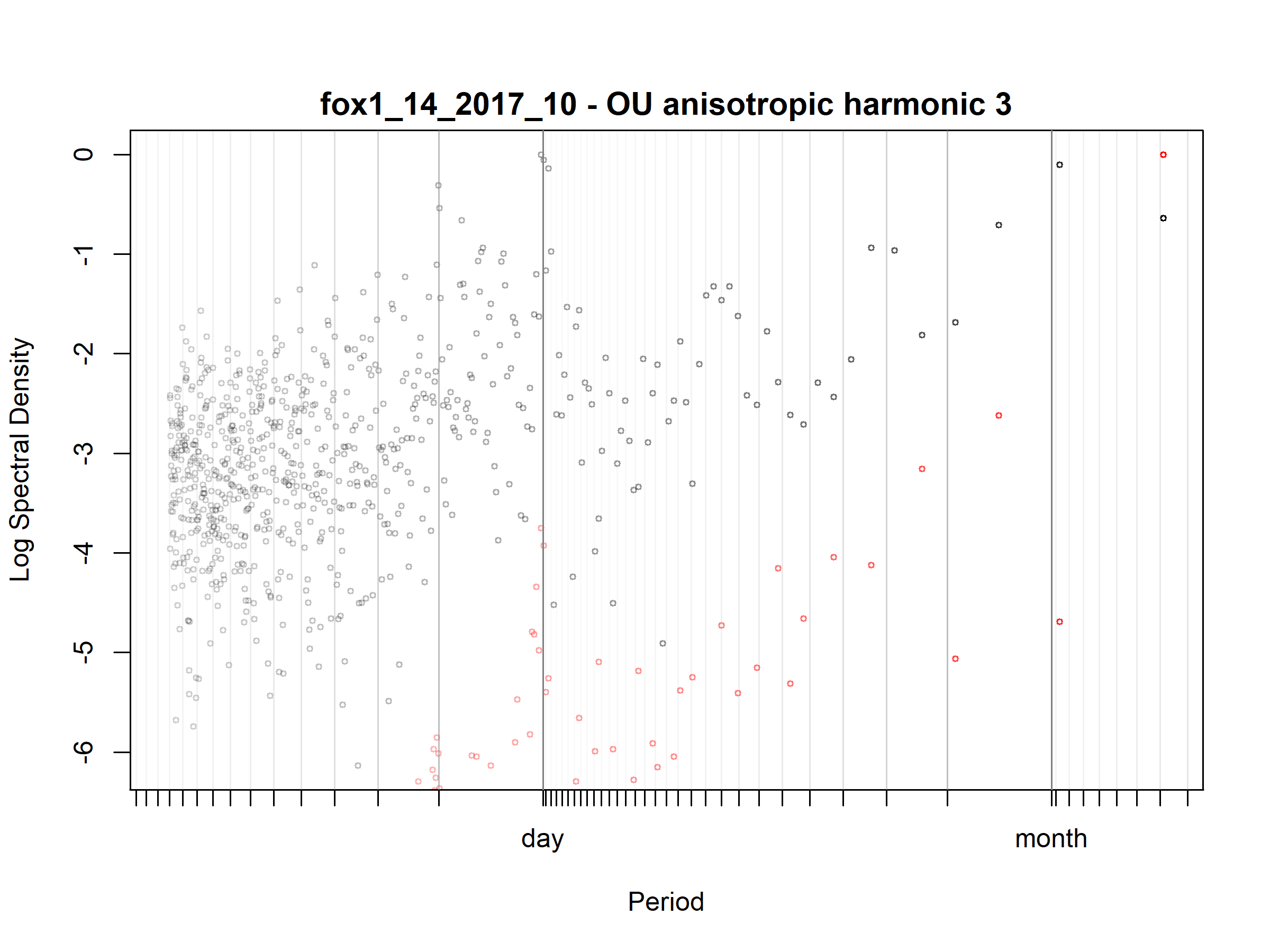

Figure S3.20. Fox; fox1_14_2017_10; diagnostic periodogram.

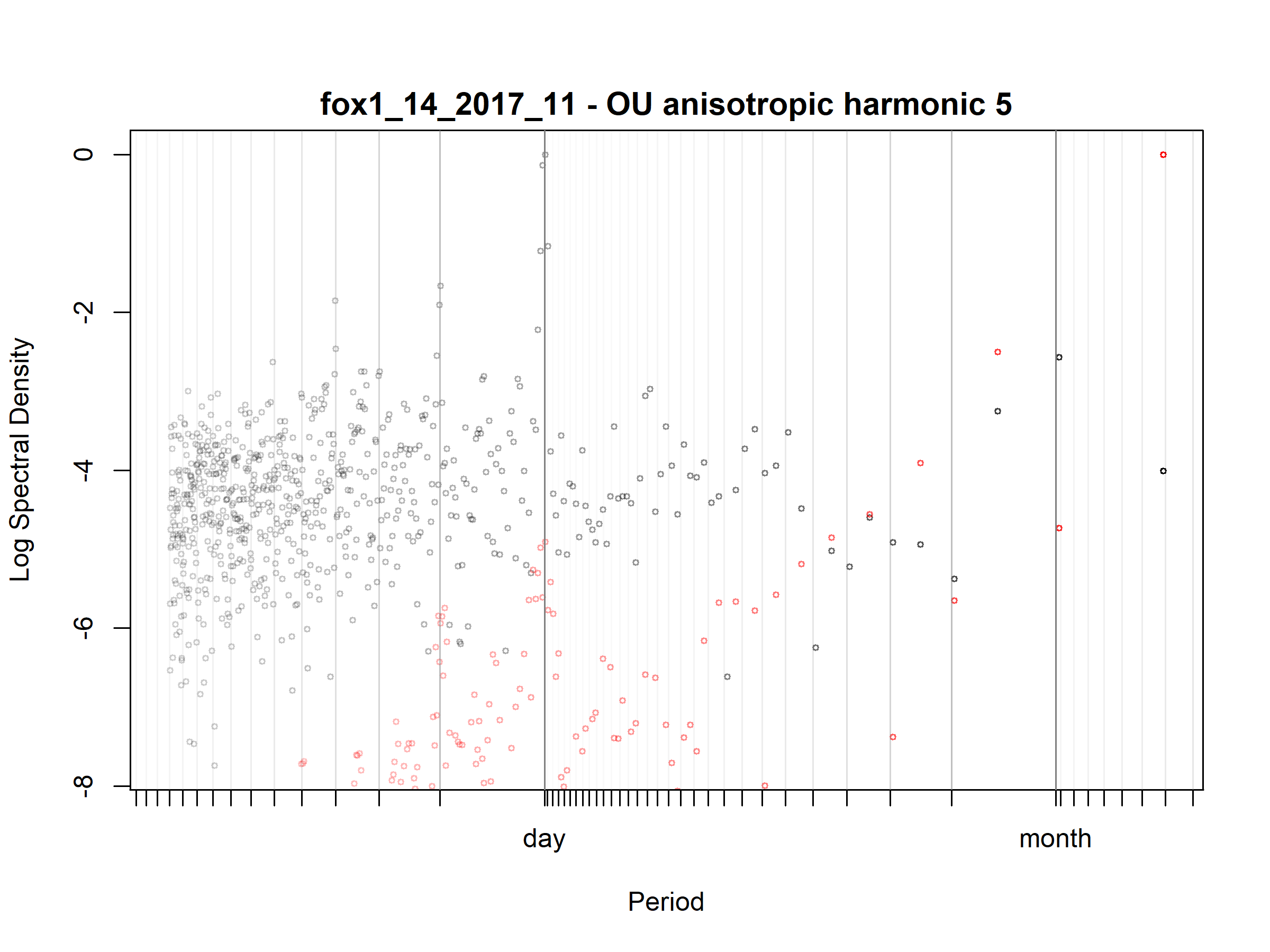

Figure S3.21. Fox; fox1_14_2017_11; diagnostic periodogram.

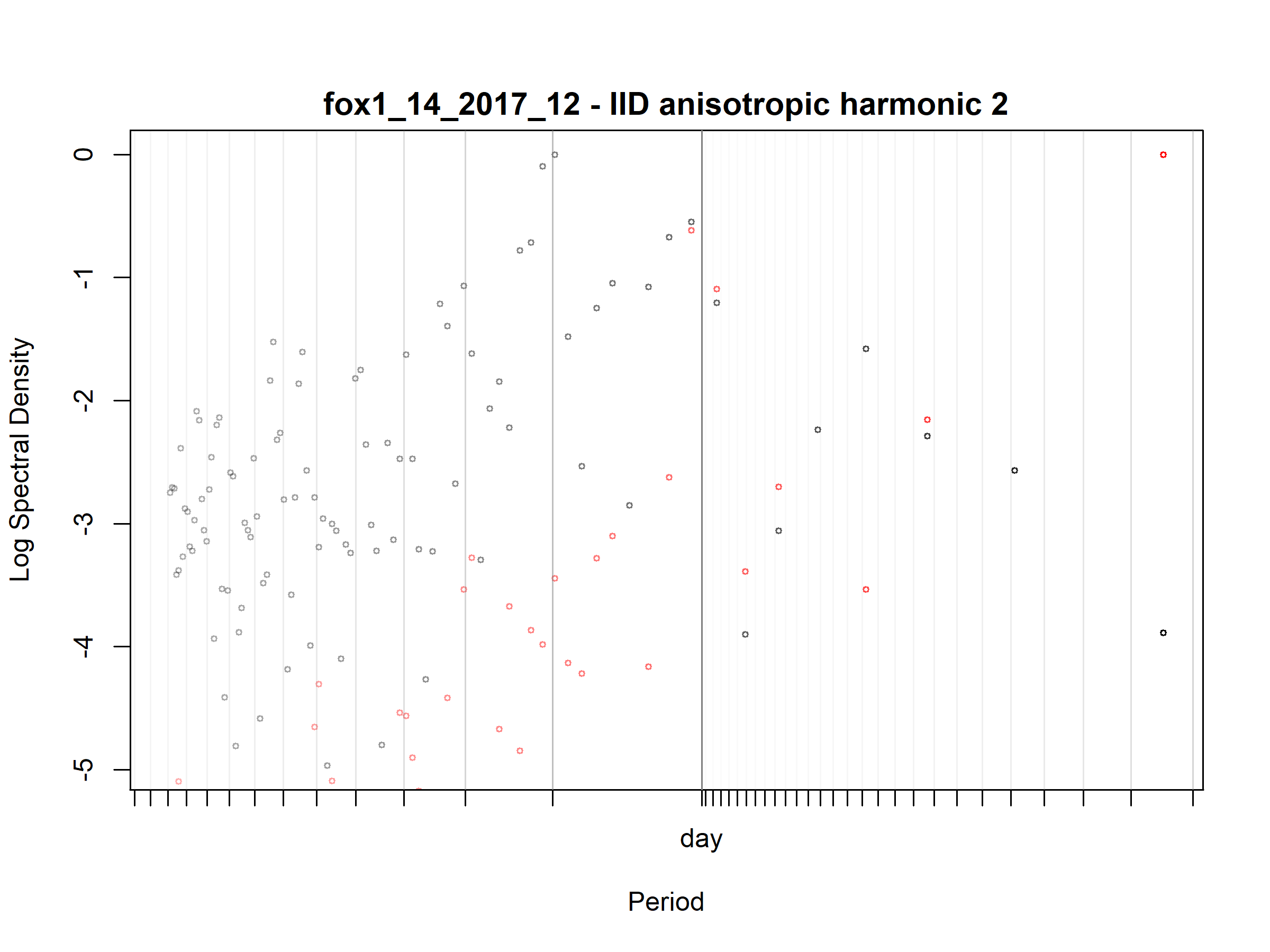

Figure S3.22. Fox; fox1_14_2017_12; diagnostic periodogram.

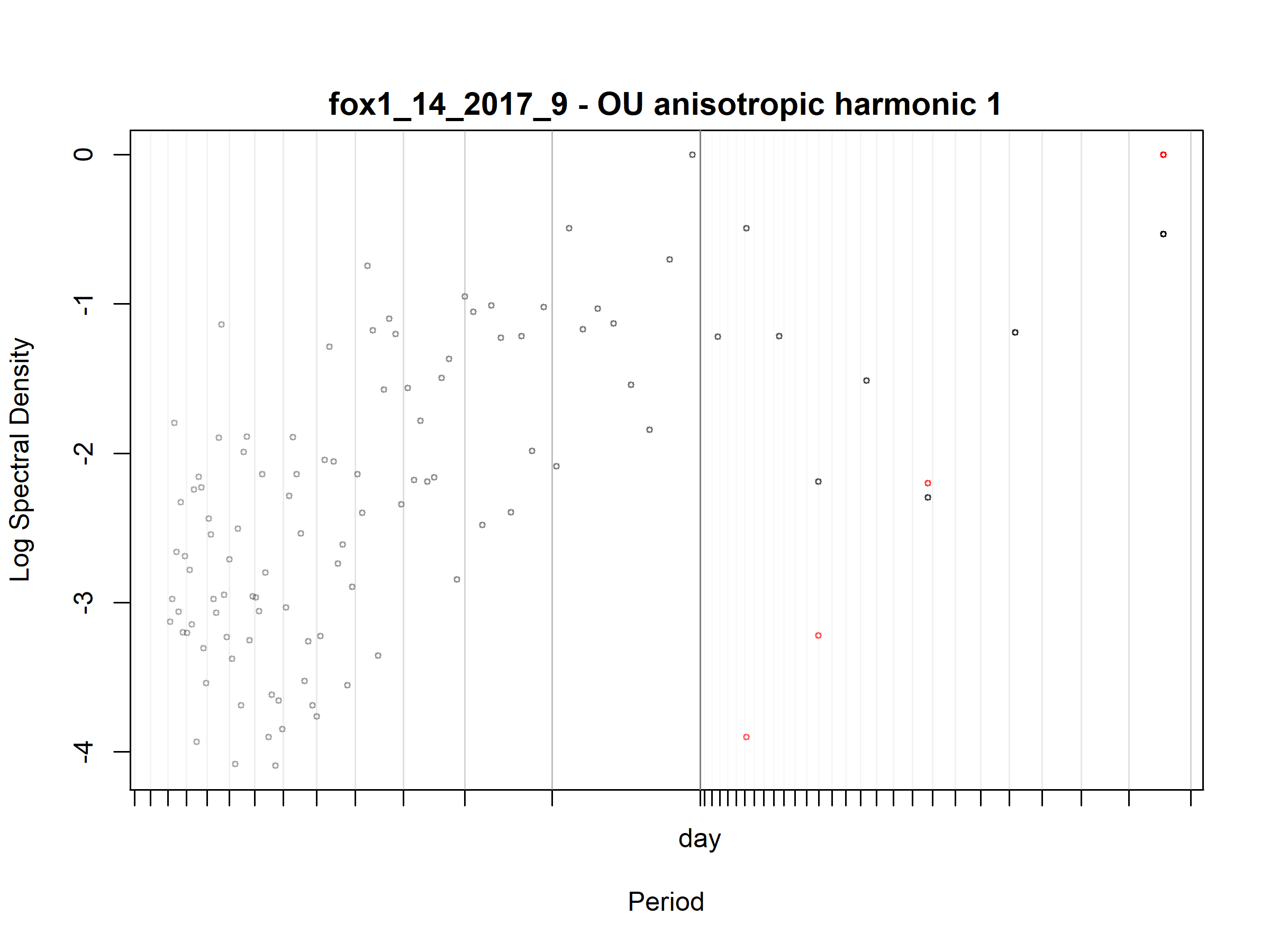

Figure S3.23. Fox; fox1_14_2017_9; diagnostic periodogram.

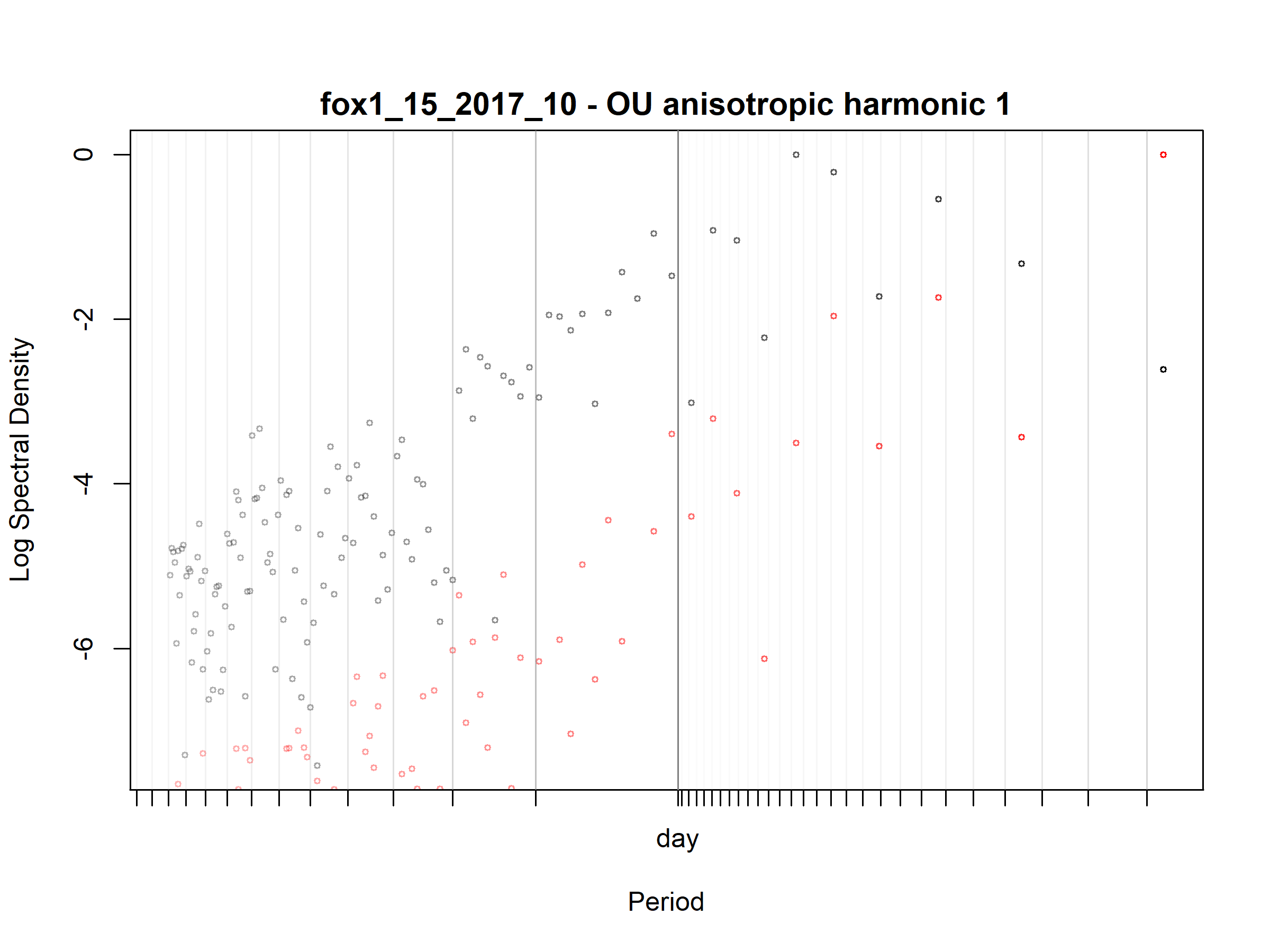

Figure S3.24. Fox; fox1_15_2017_10; diagnostic periodogram.

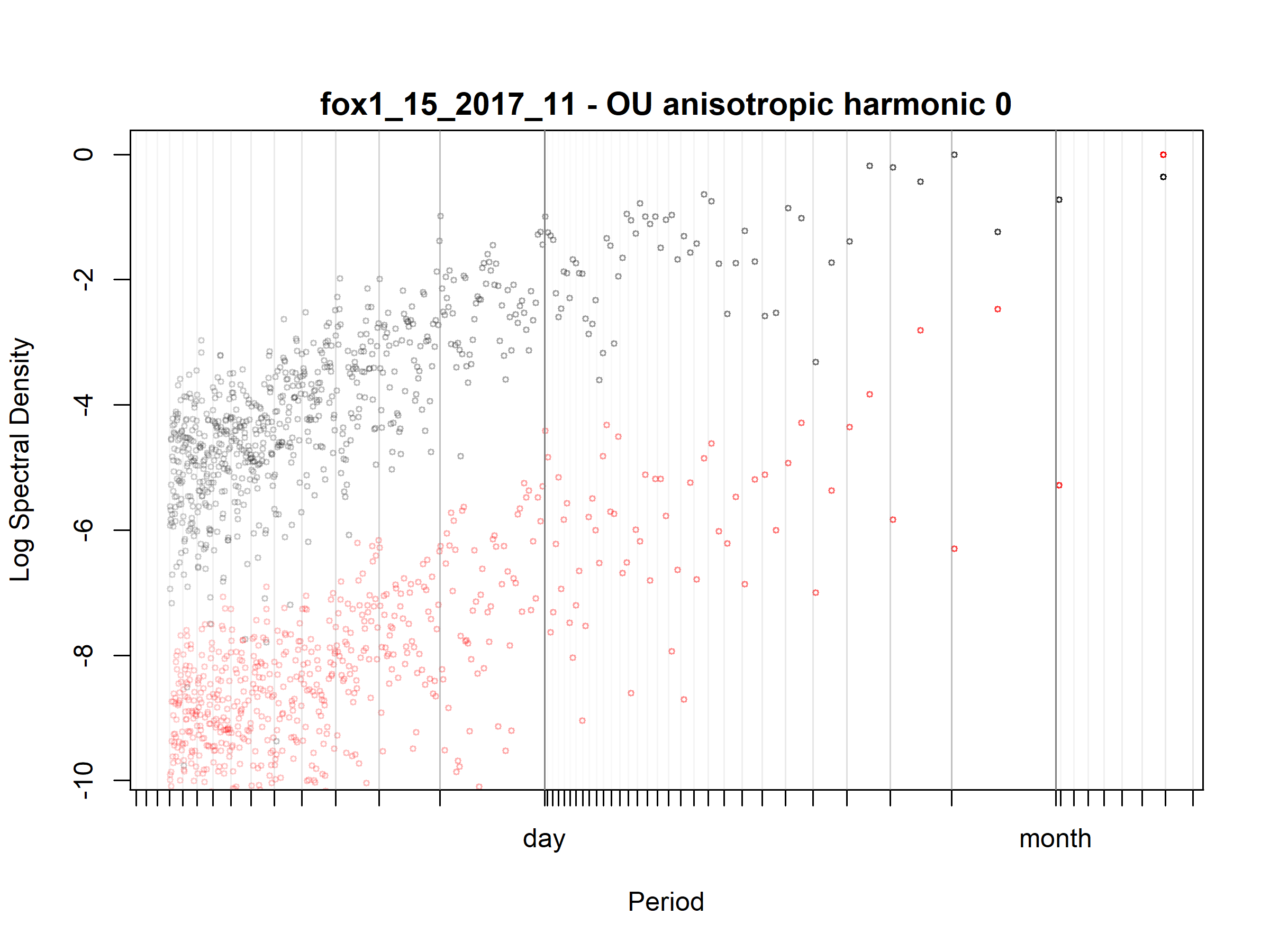

Figure S3.25. Fox; fox1_15_2017_11; diagnostic periodogram.

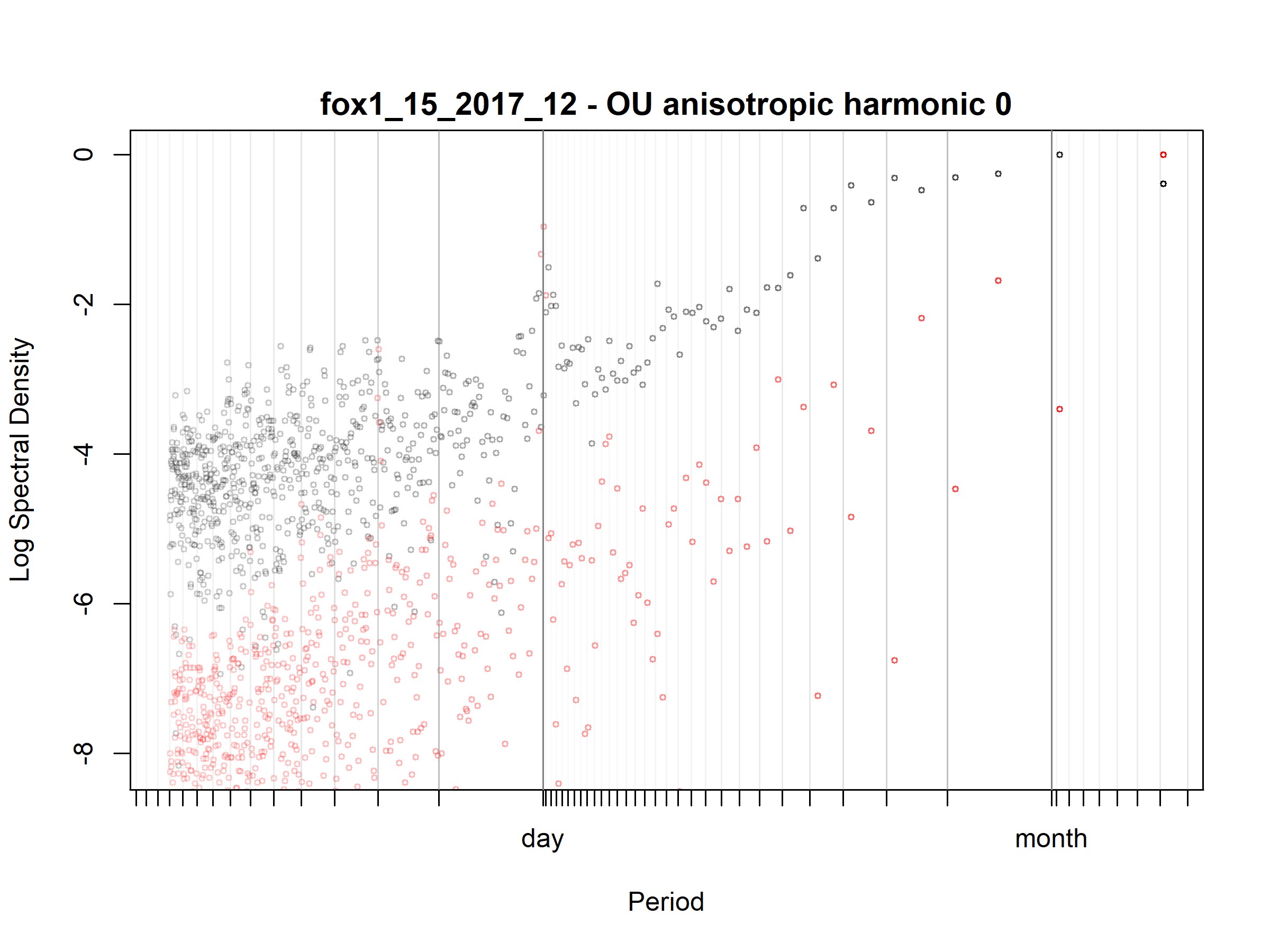

Figure S3.26. Fox; fox1_15_2017_12; diagnostic periodogram.

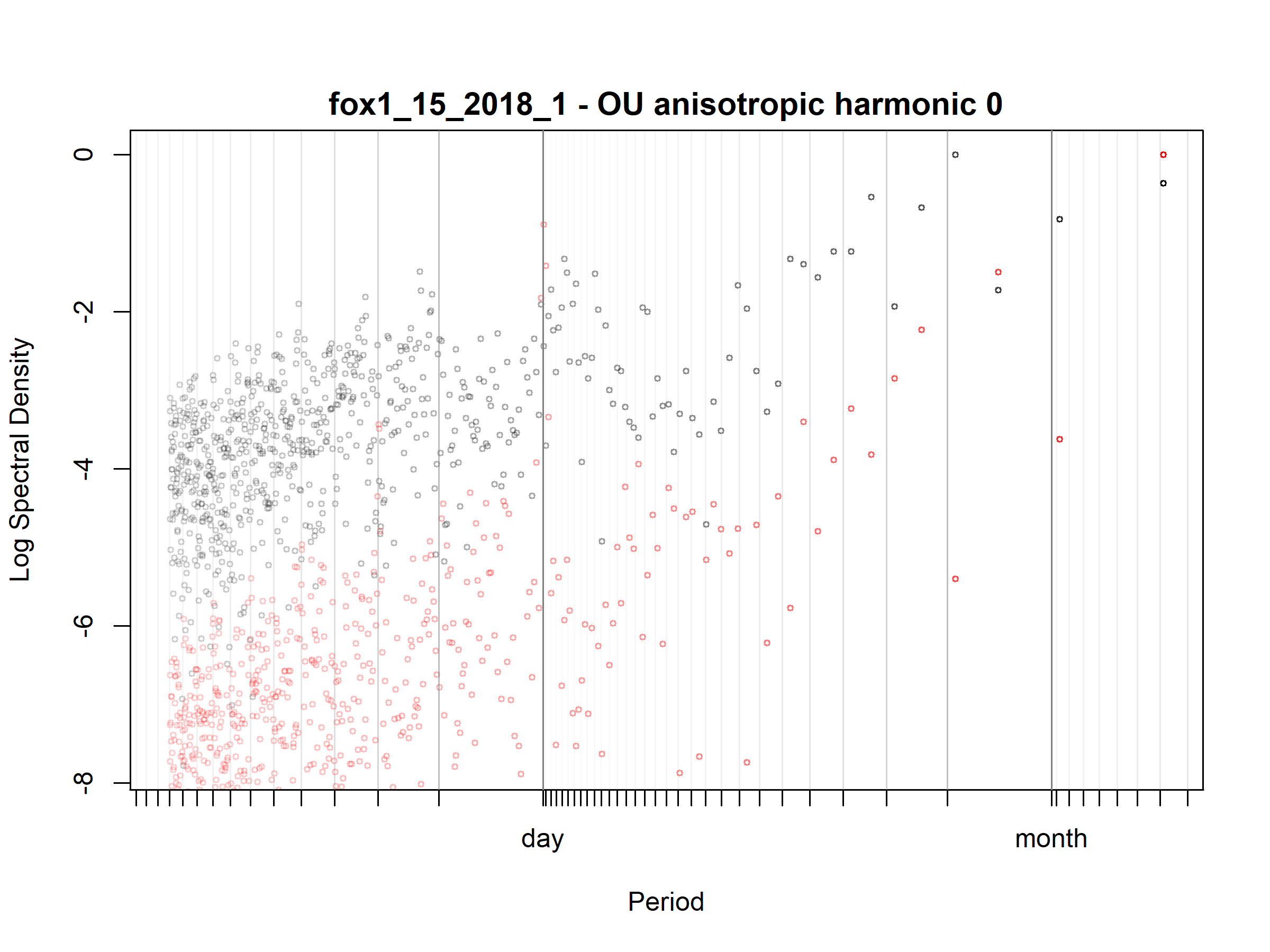

Figure S3.27. Fox; fox1_15_2018_1; diagnostic periodogram.

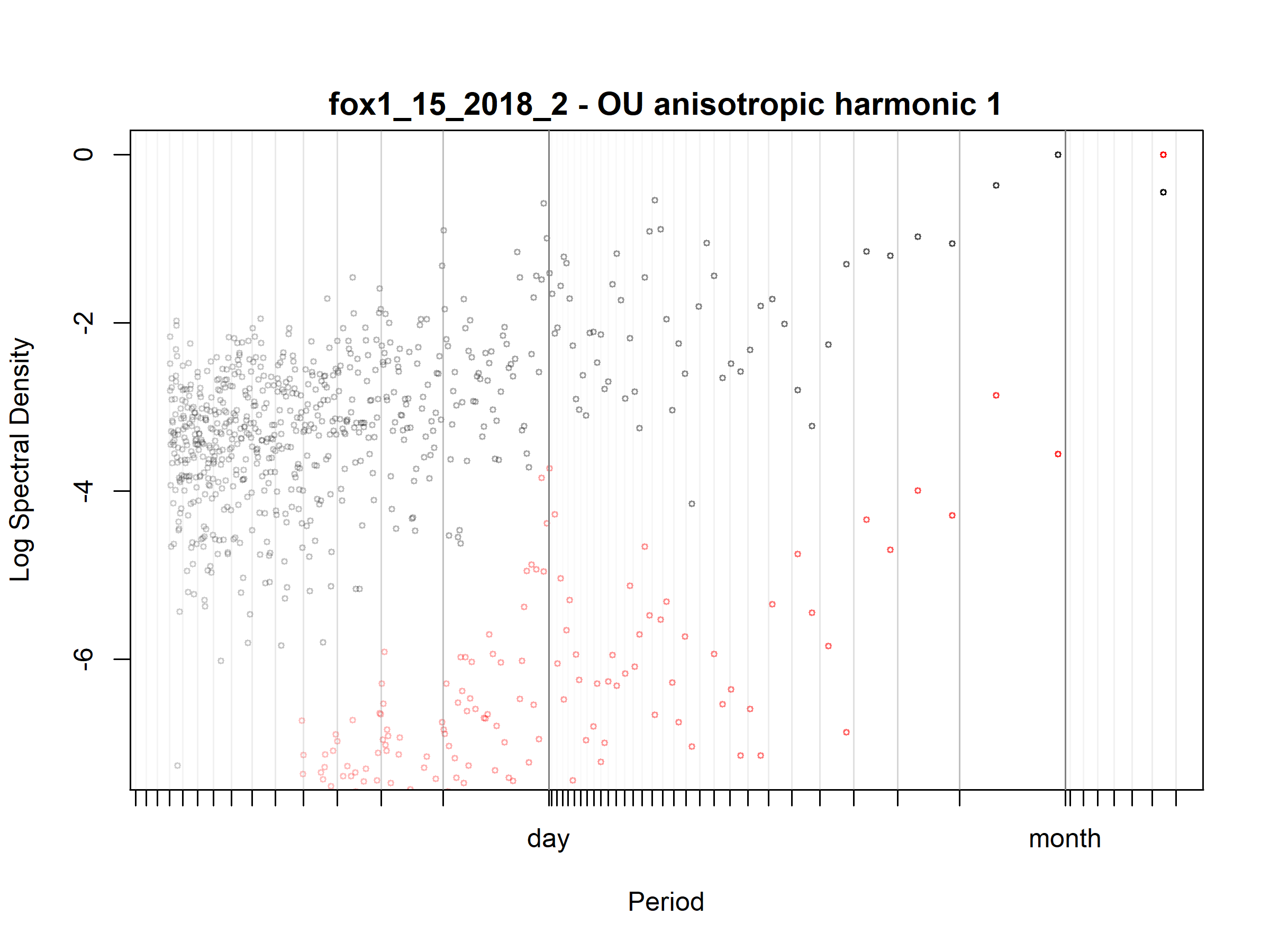

Figure S3.28. Fox; fox1_15_2018_2; diagnostic periodogram.

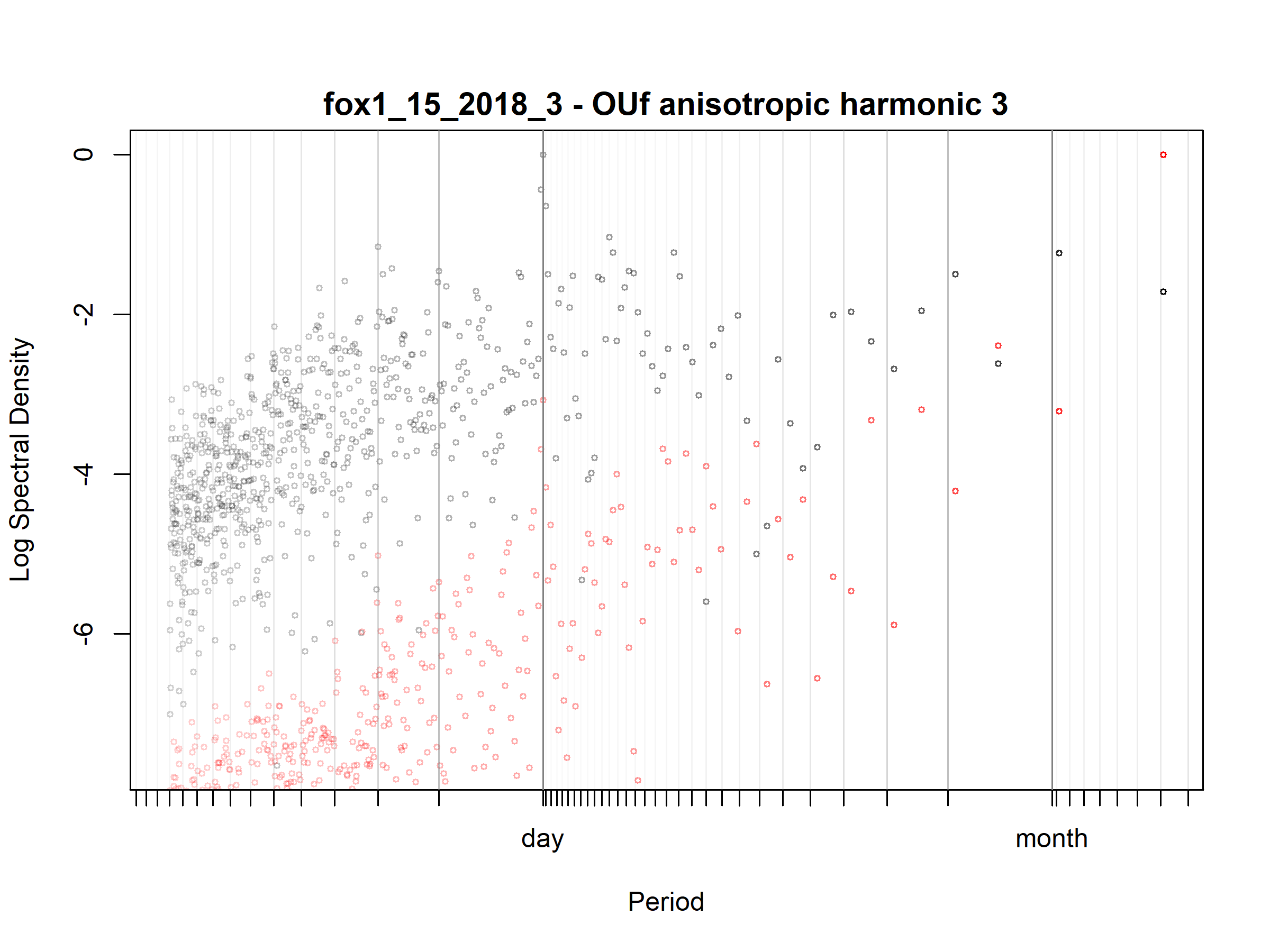

Figure S3.29. Fox; fox1_15_2018_3; diagnostic periodogram.

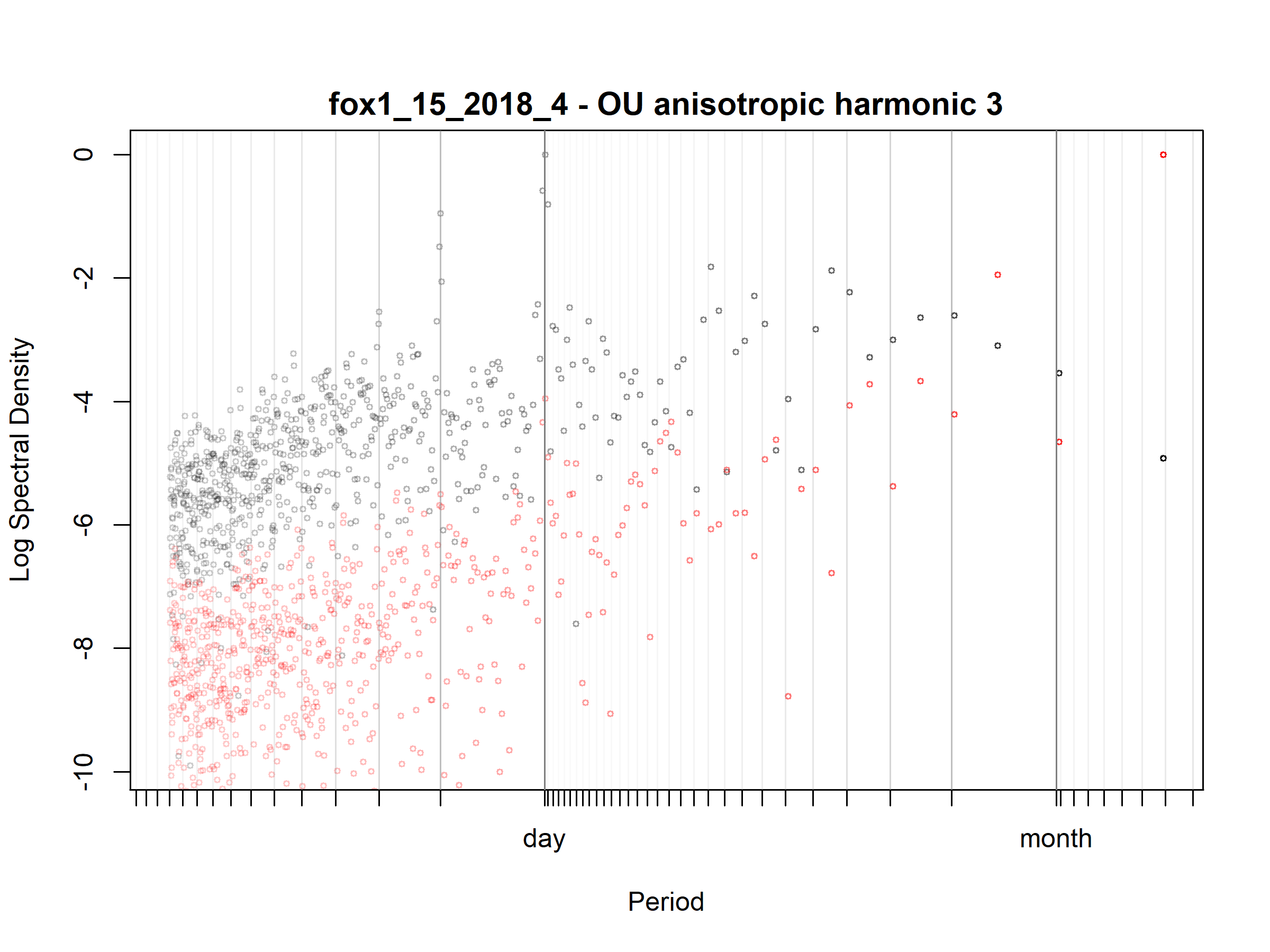

Figure S3.30. Fox; fox1_15_2018_4; diagnostic periodogram.

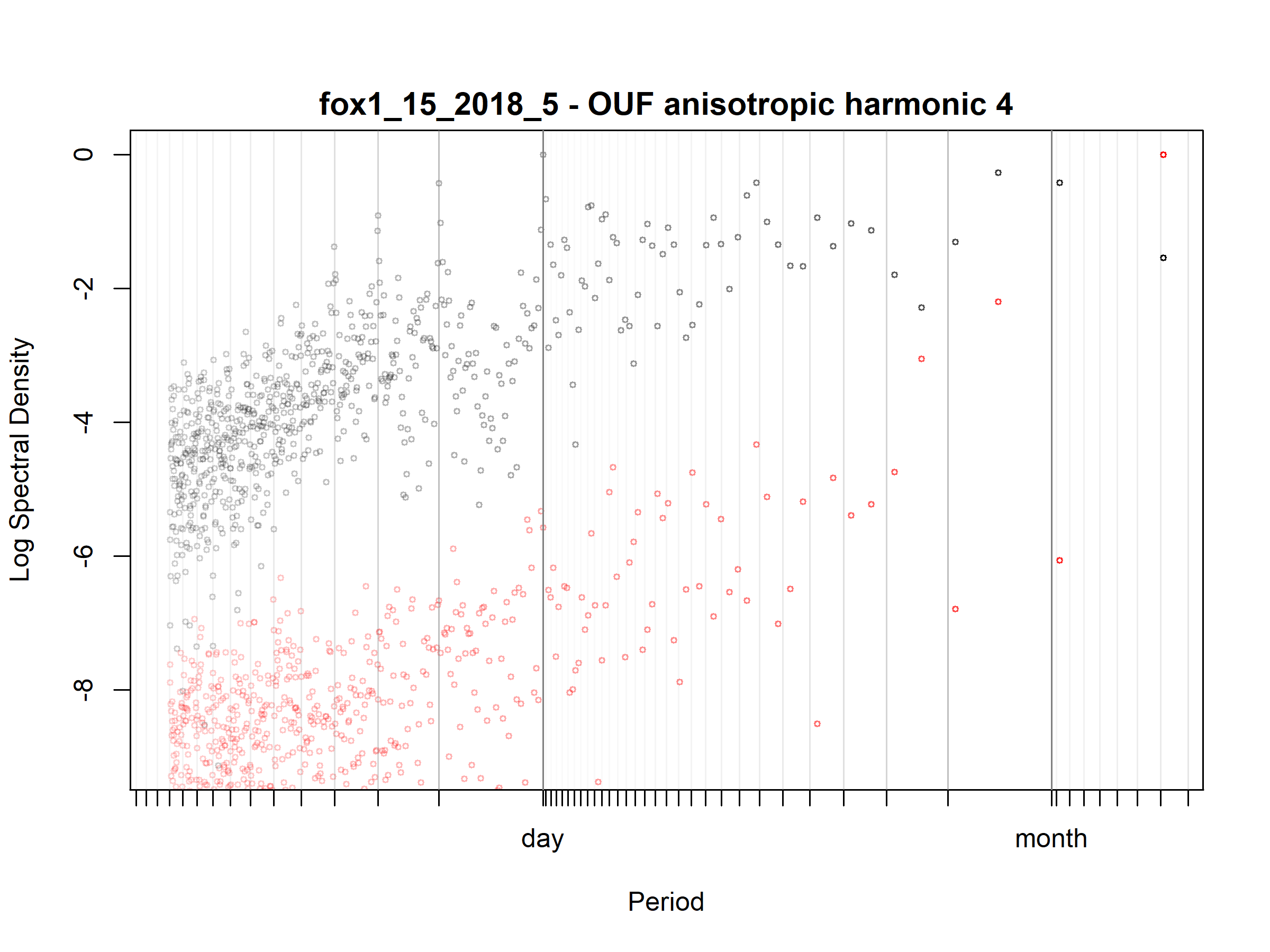

Figure S3.31. Fox; fox1_15_2018_5; diagnostic periodogram.

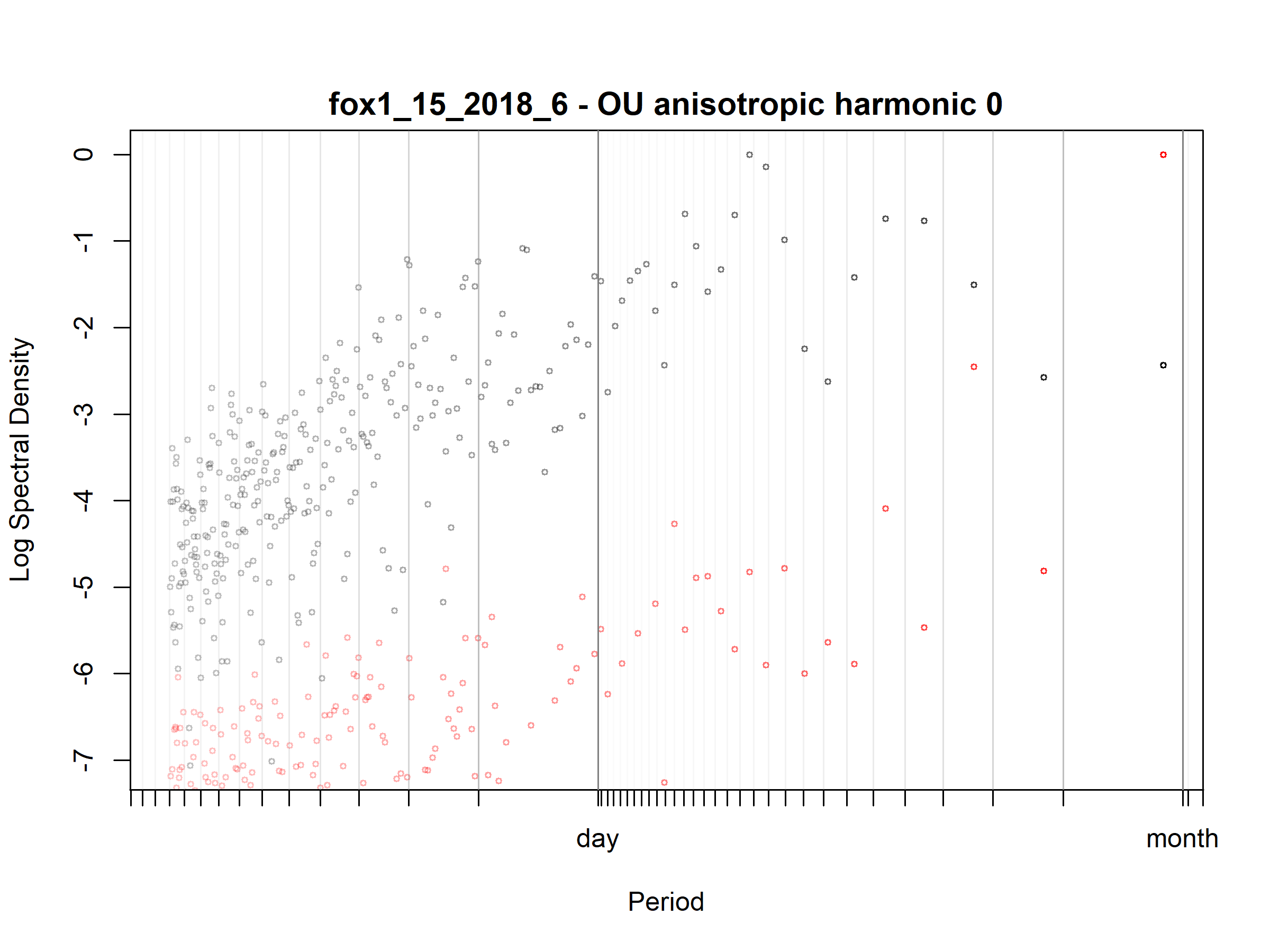

Figure S3.32. Fox; fox1_15_2018_6; diagnostic periodogram.

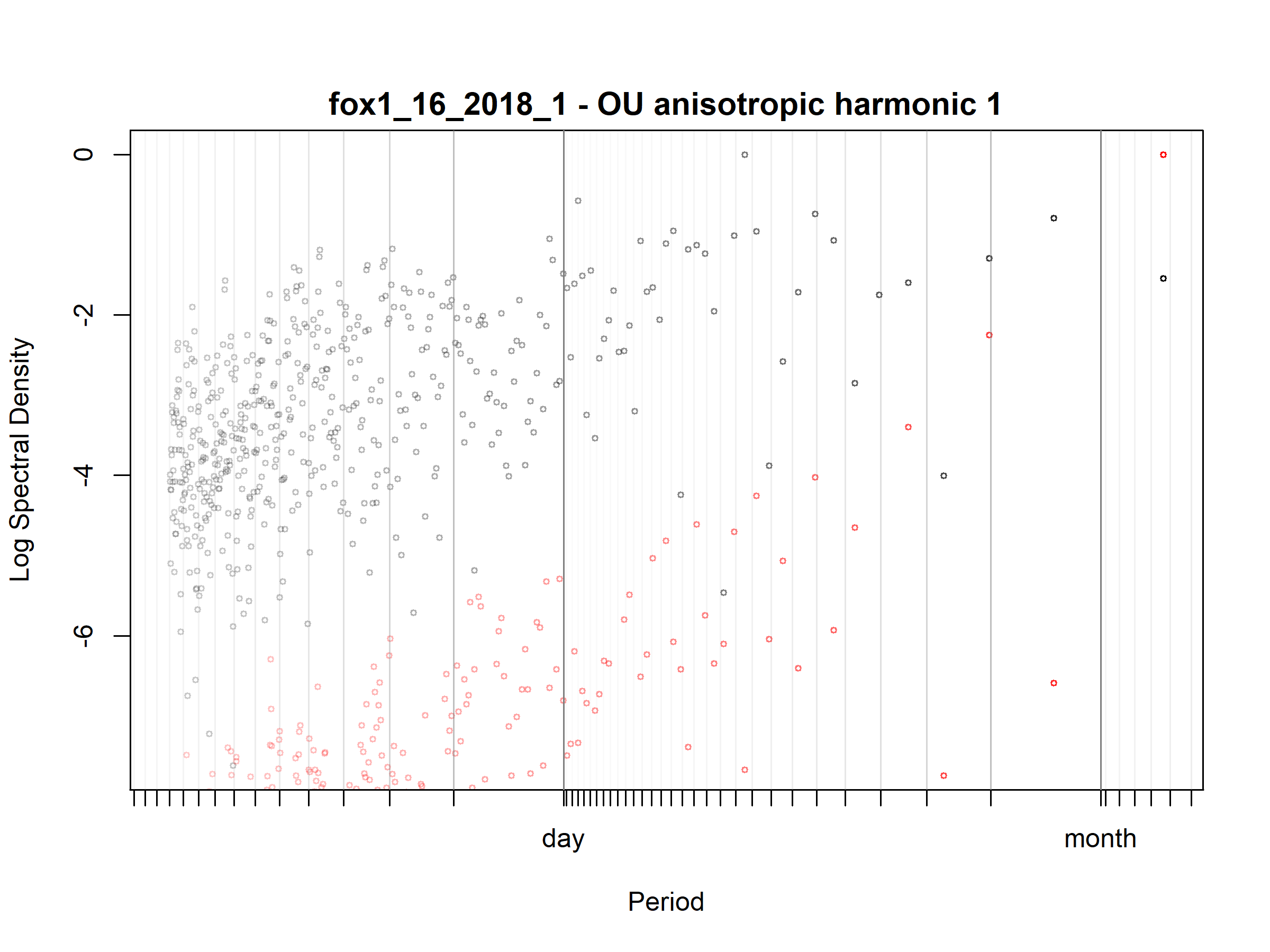

Figure S3.33. Fox; fox1_16_2018_1; diagnostic periodogram.

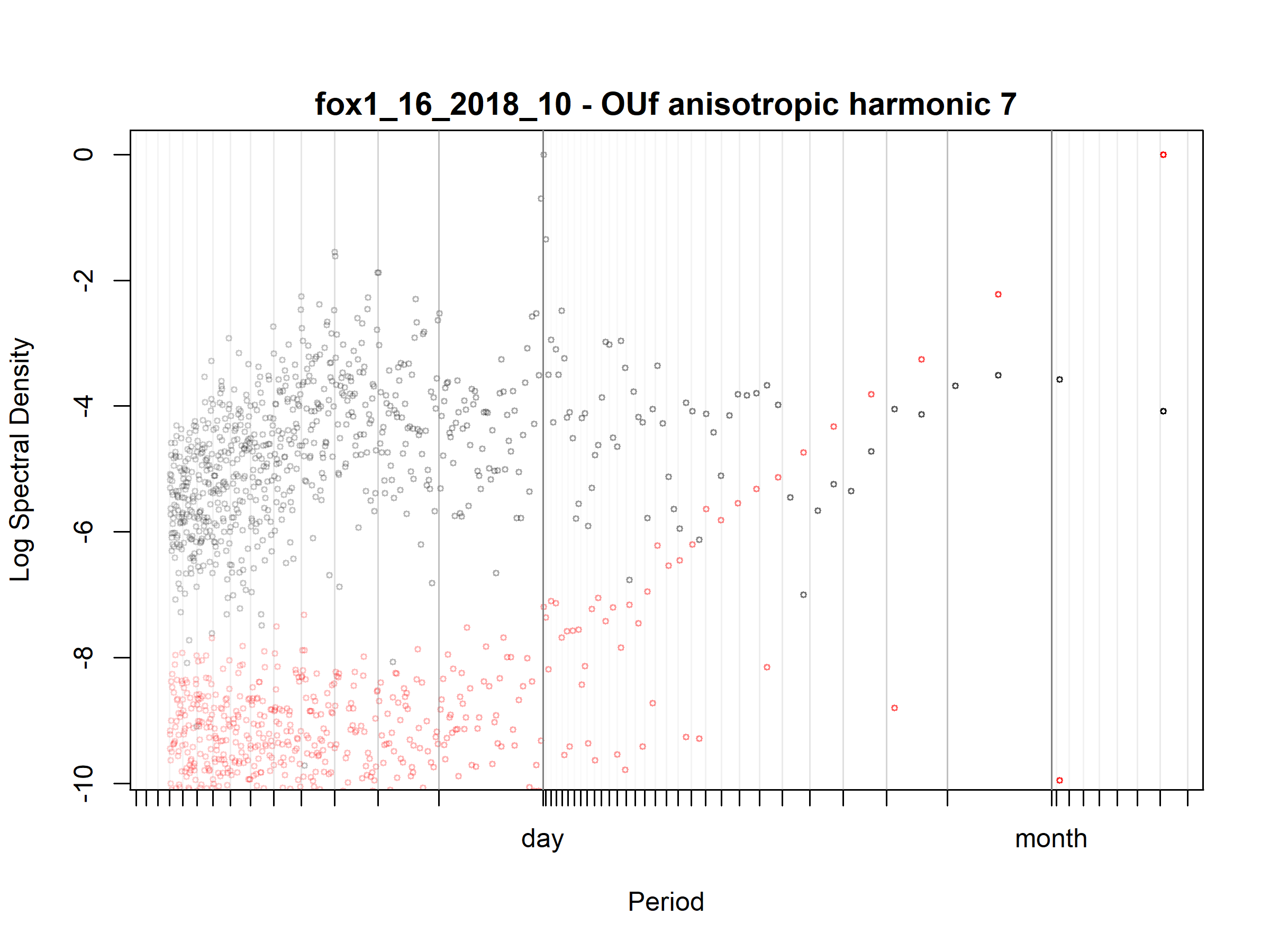

Figure S3.34. Fox; fox1_16_2018_10; diagnostic periodogram.

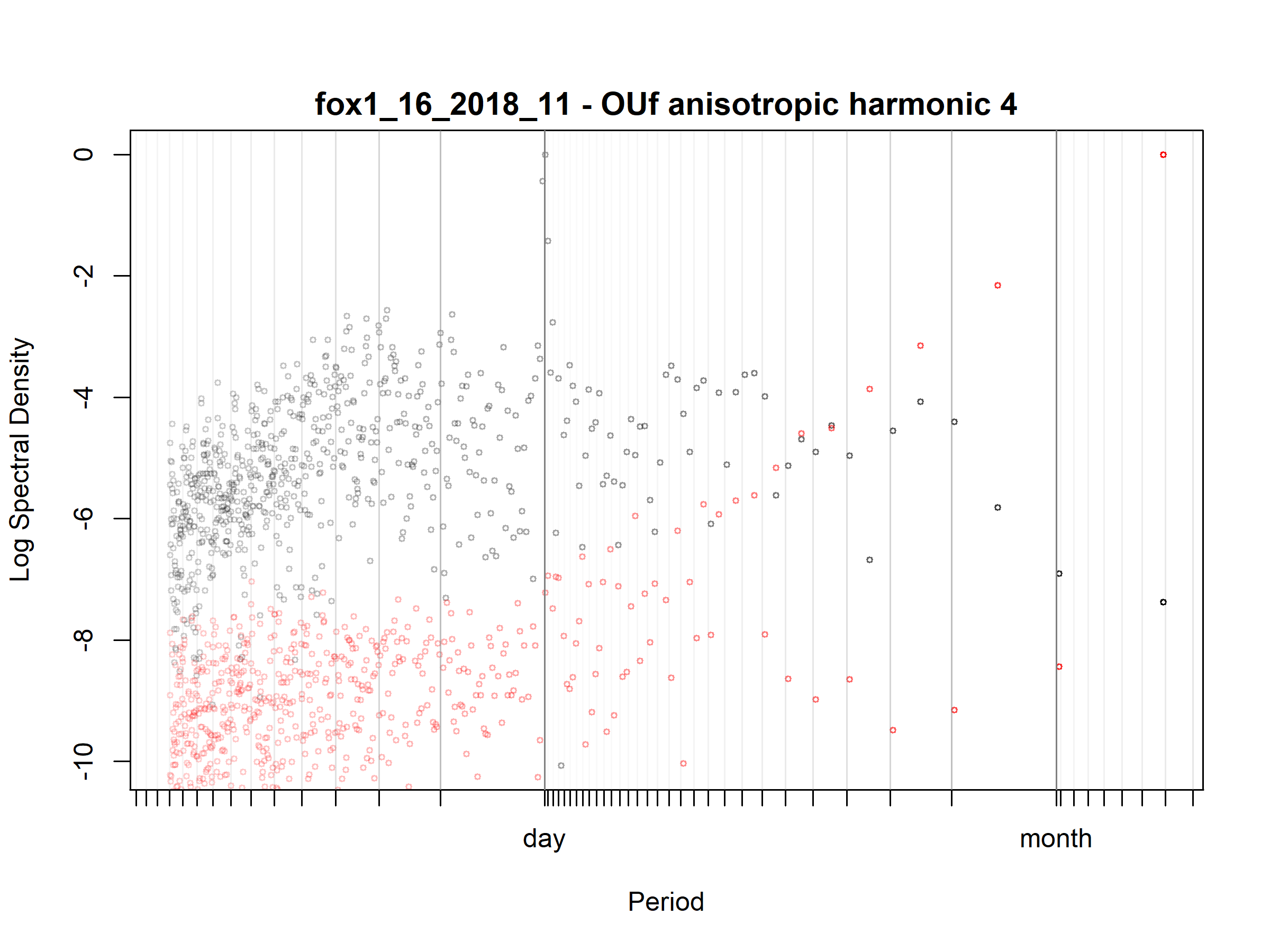

Figure S3.35. Fox; fox1_16_2018_11; diagnostic periodogram.

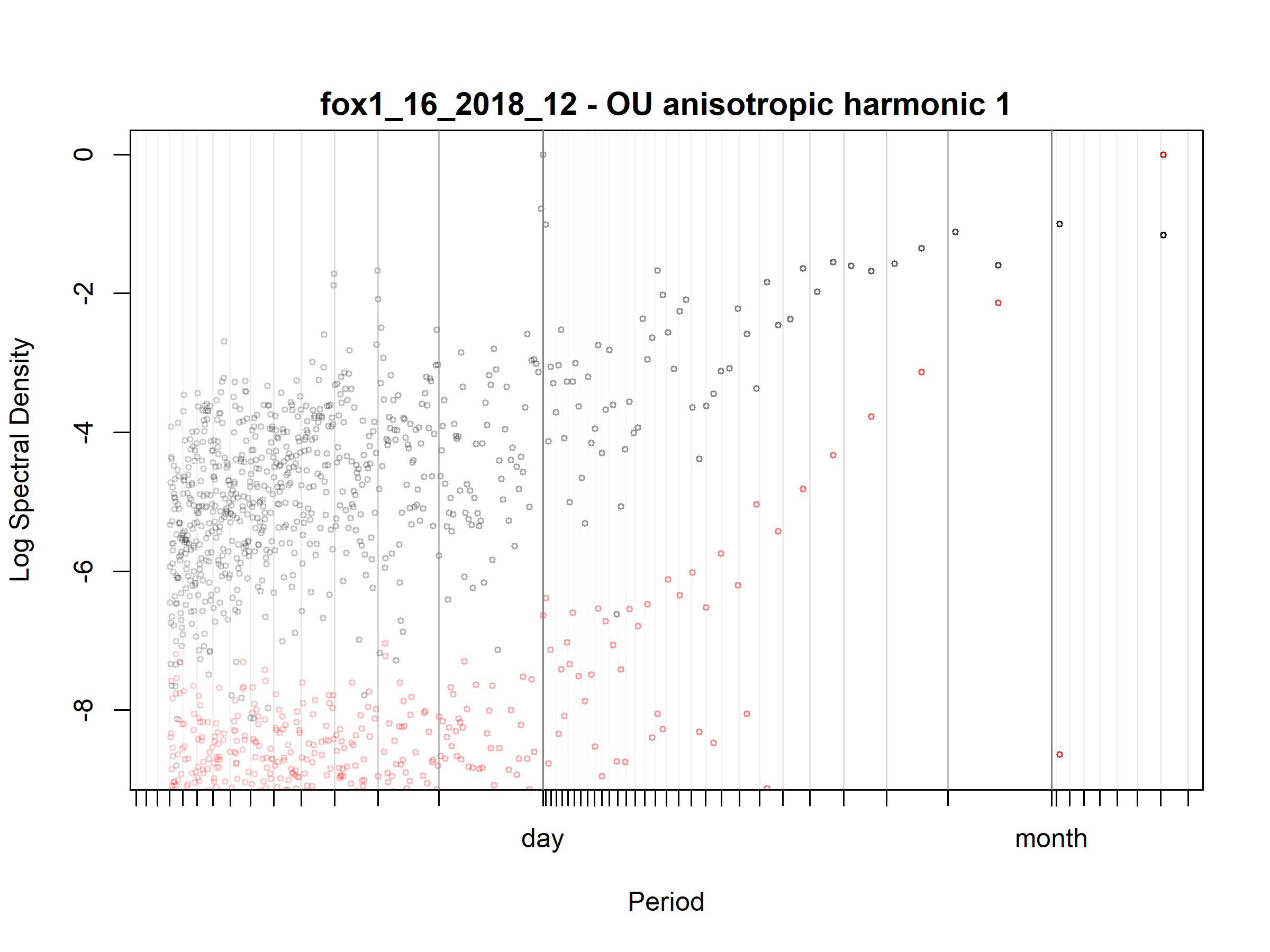

Figure S3.36. Fox; fox1_16_2018_12; diagnostic periodogram.

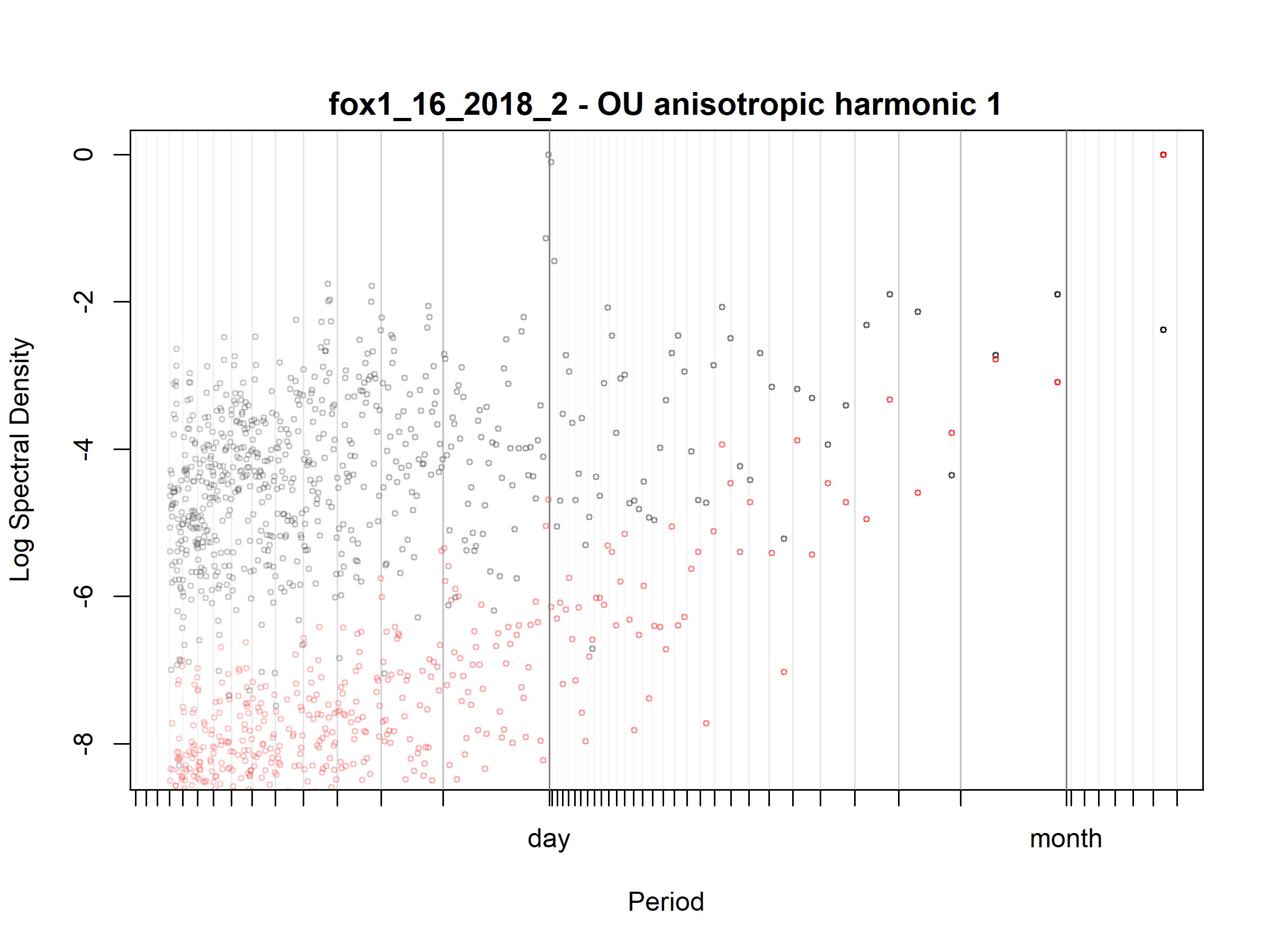

Figure S3.37. Fox; fox1_16_2018_2; diagnostic periodogram.

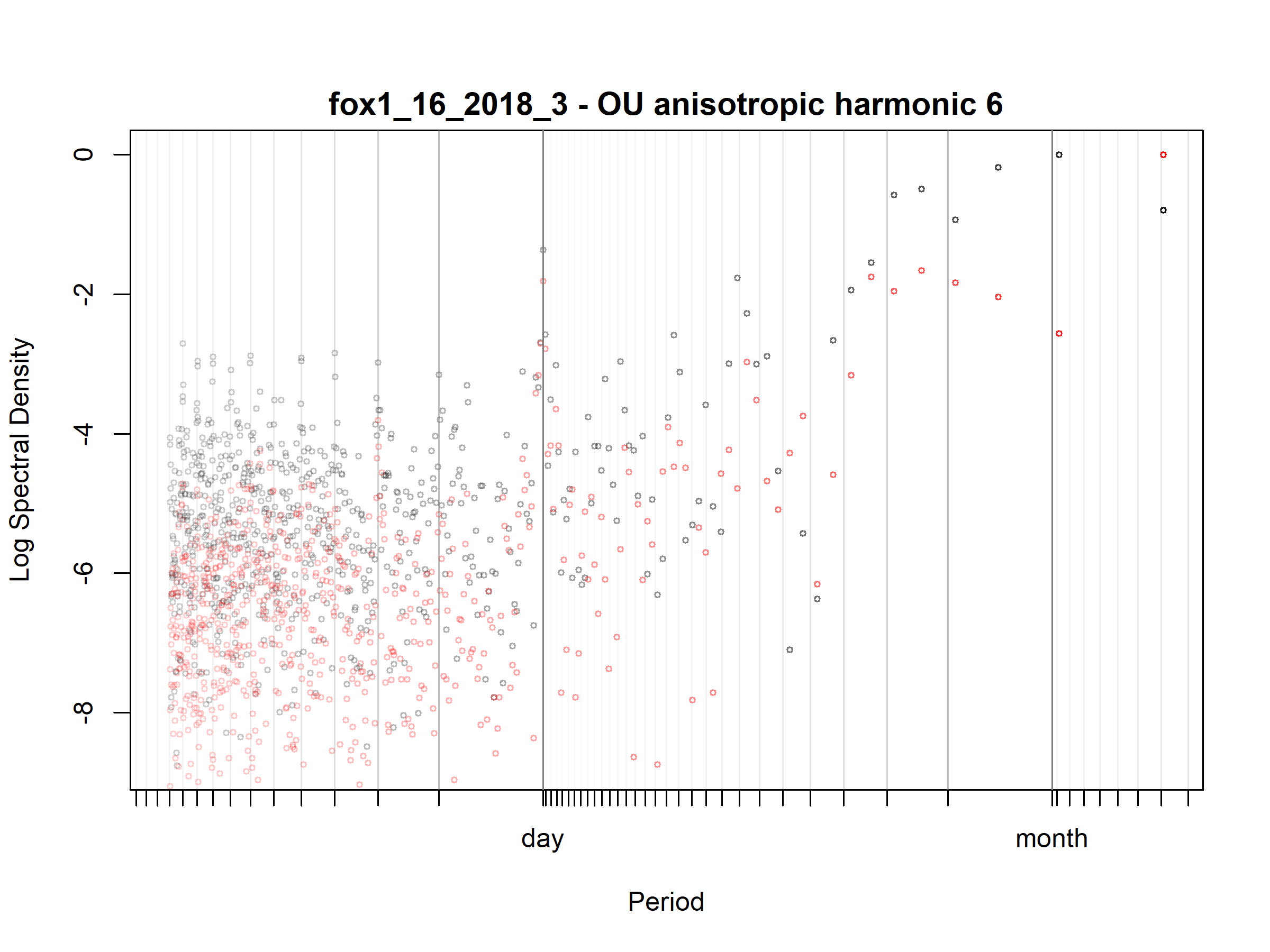

Figure S3.38. Fox; fox1_16_2018_3; diagnostic periodogram.

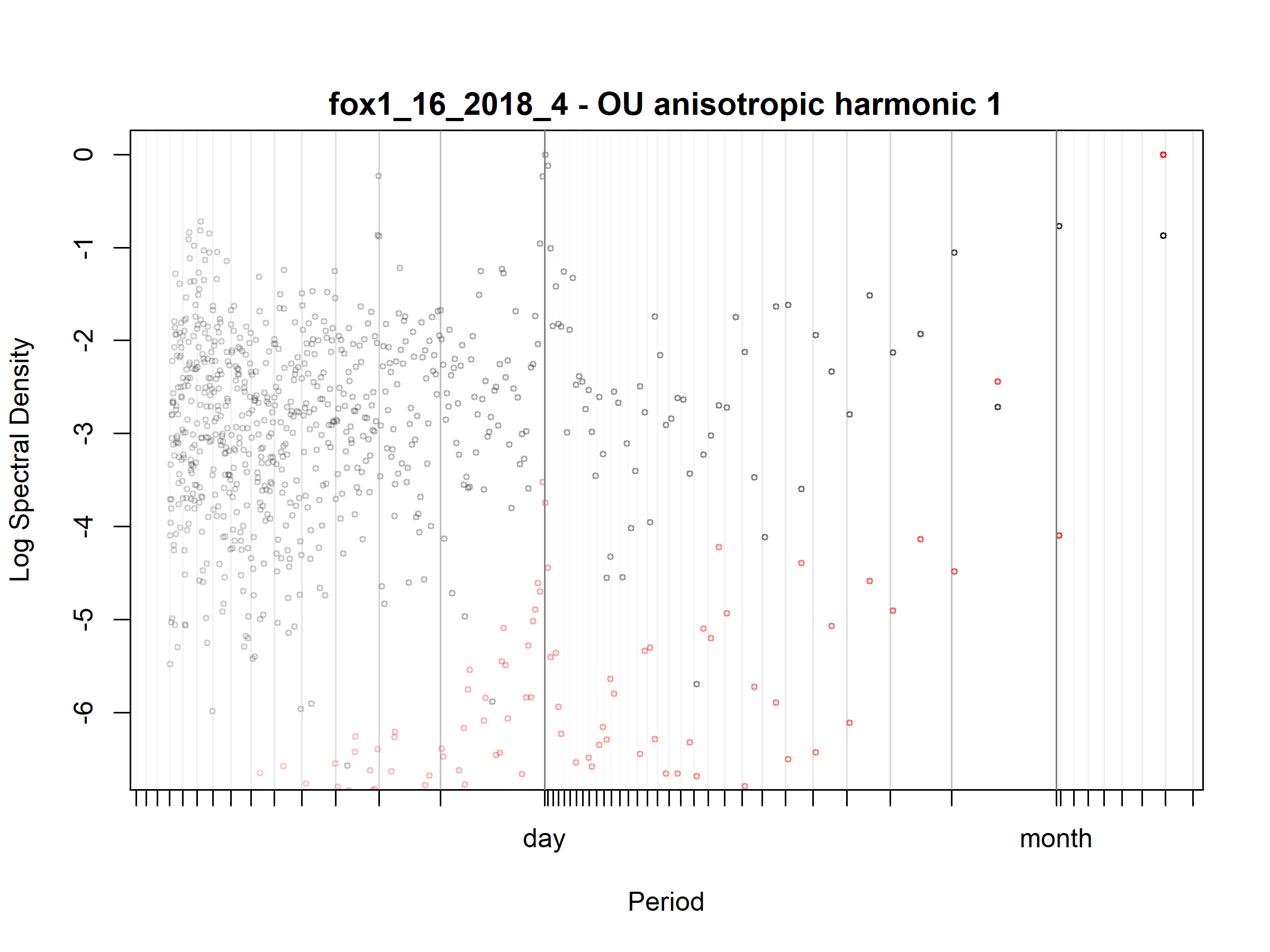

Figure S3.39. Fox; fox1_16_2018_4; diagnostic periodogram.

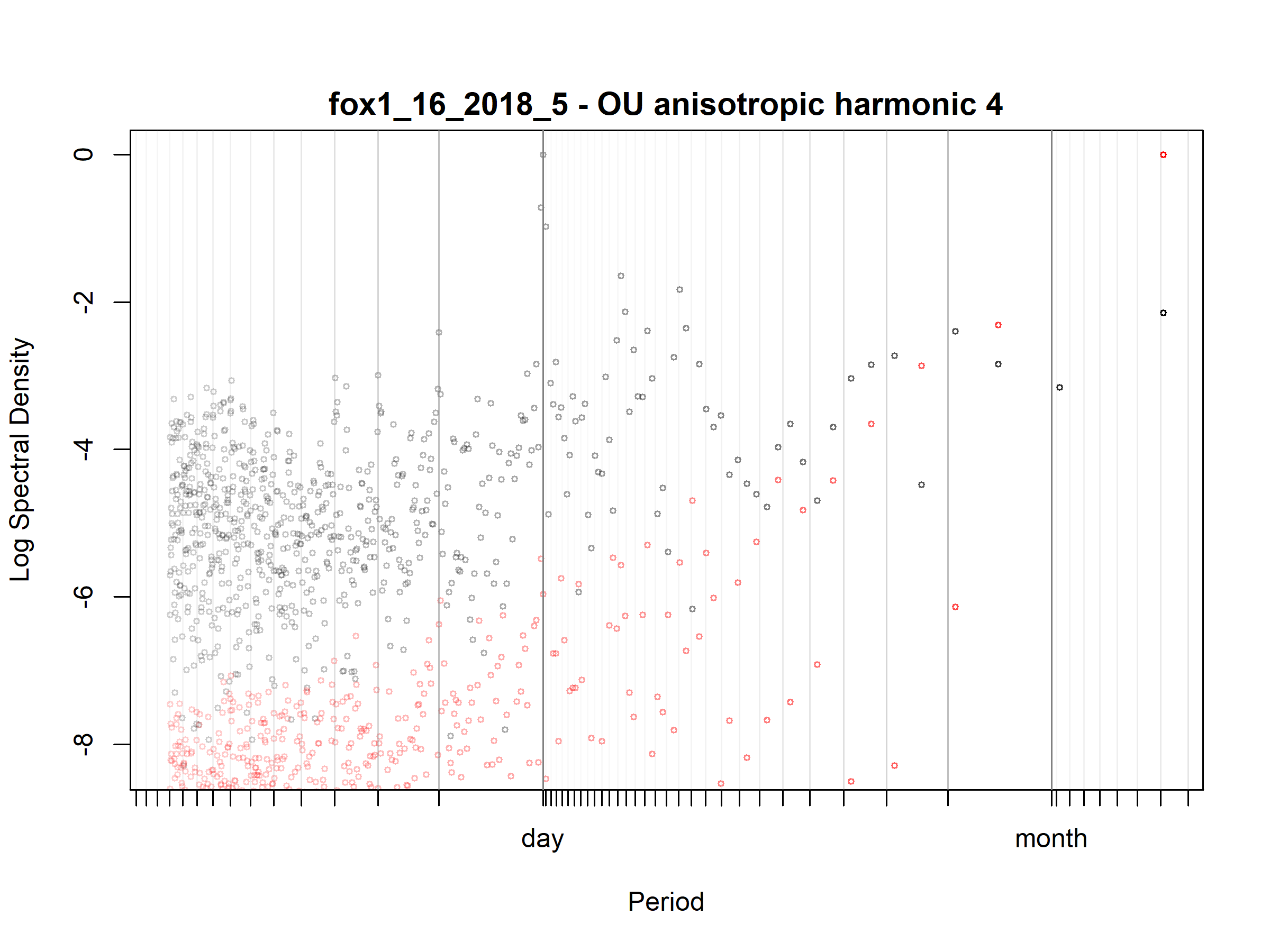

Figure S3.40. Fox; fox1_16_2018_5; diagnostic periodogram.

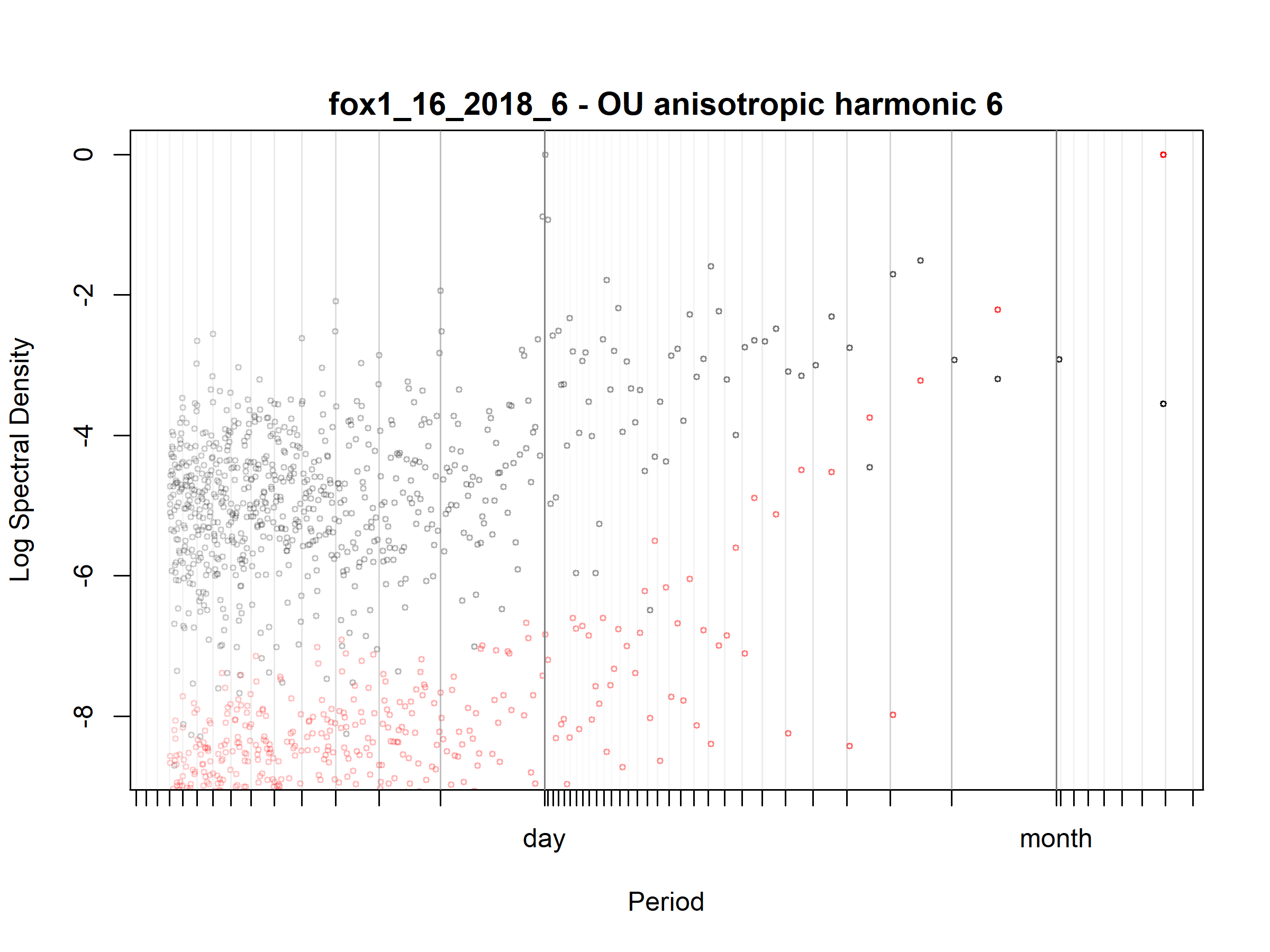

Figure S3.41. Fox; fox1_16_2018_6; diagnostic periodogram.

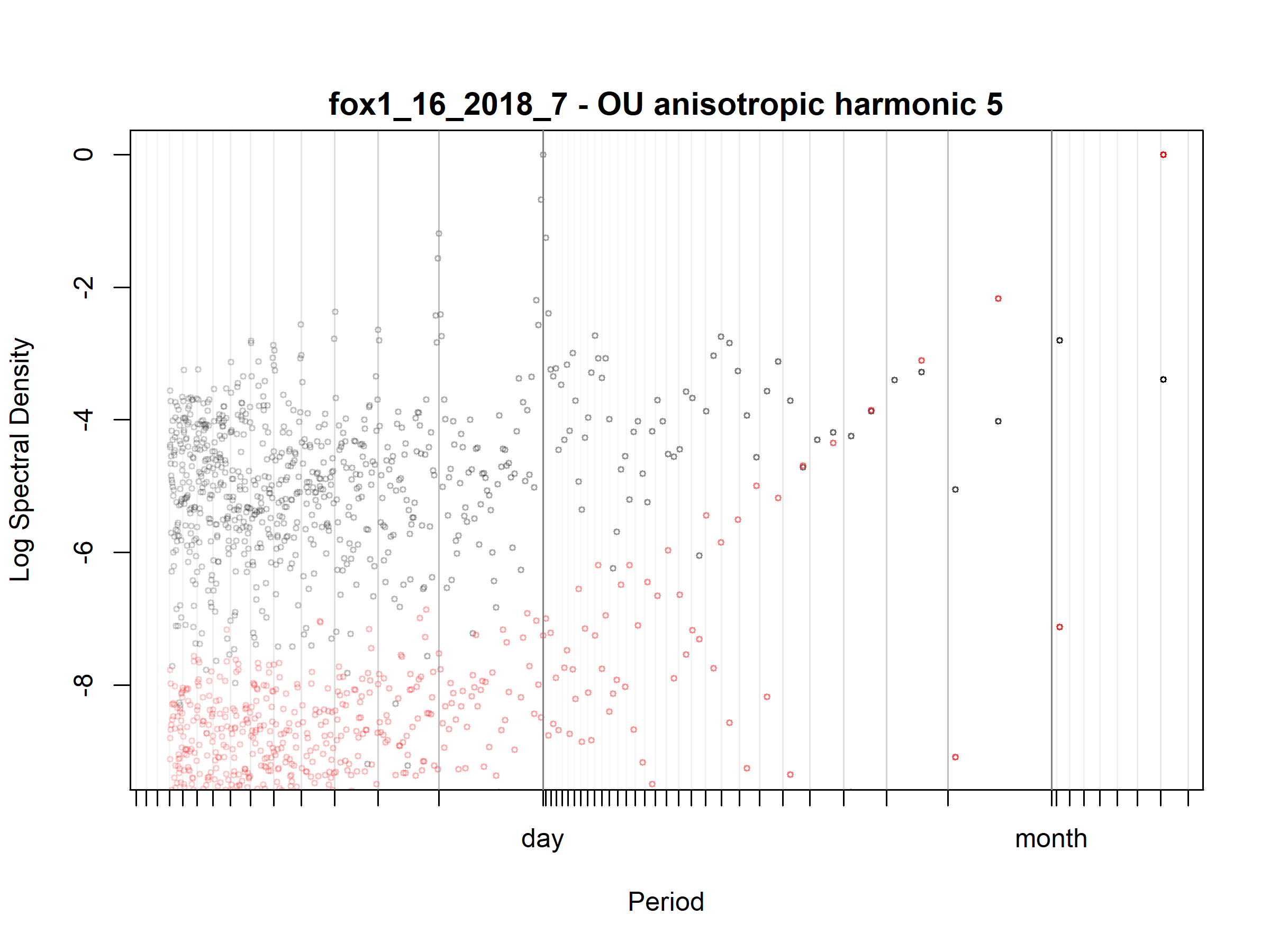

Figure S3.42. Fox; fox1_16_2018_7; diagnostic periodogram.

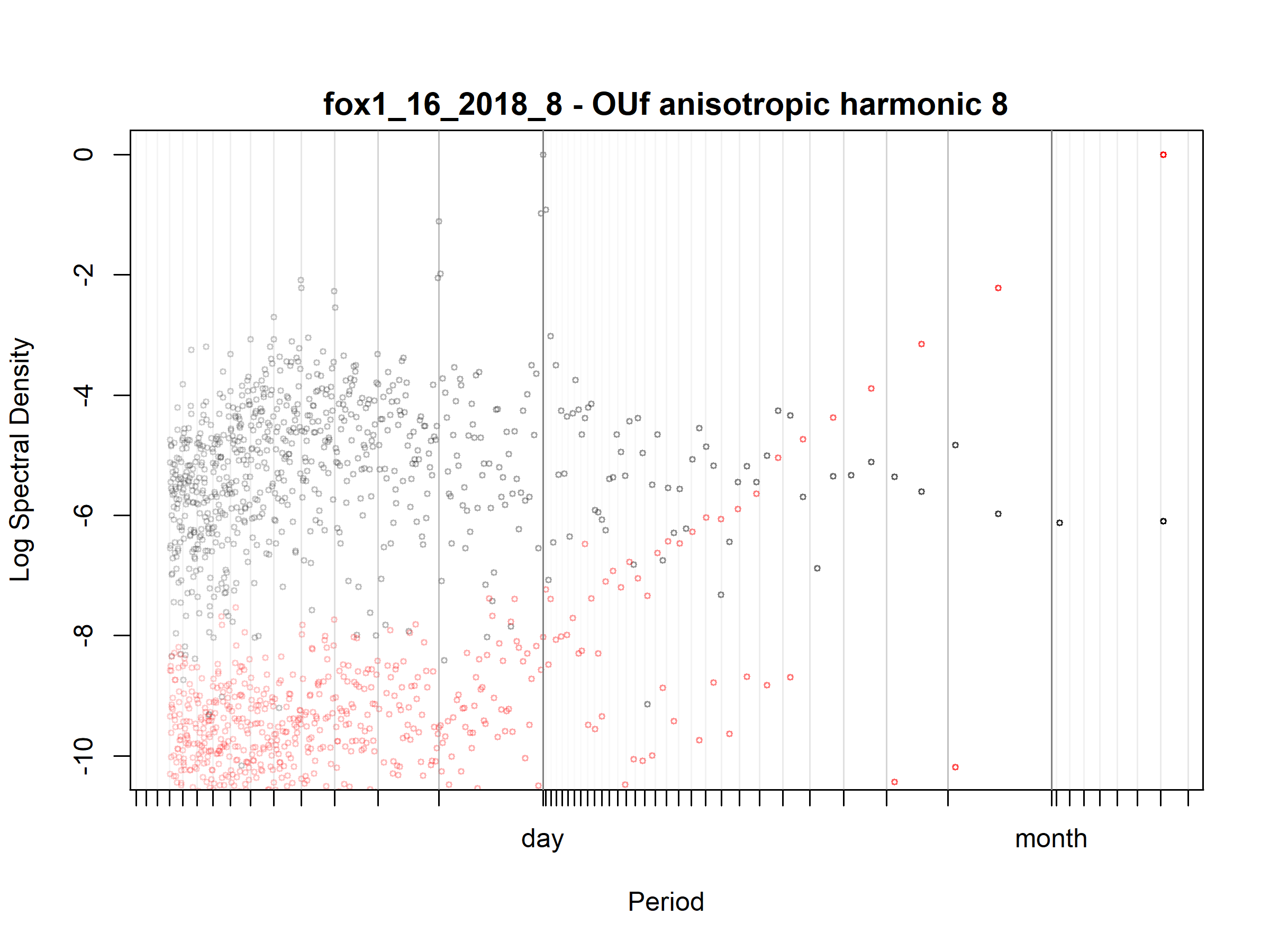

Figure S3.43. Fox; fox1_16_2018_8; diagnostic periodogram.

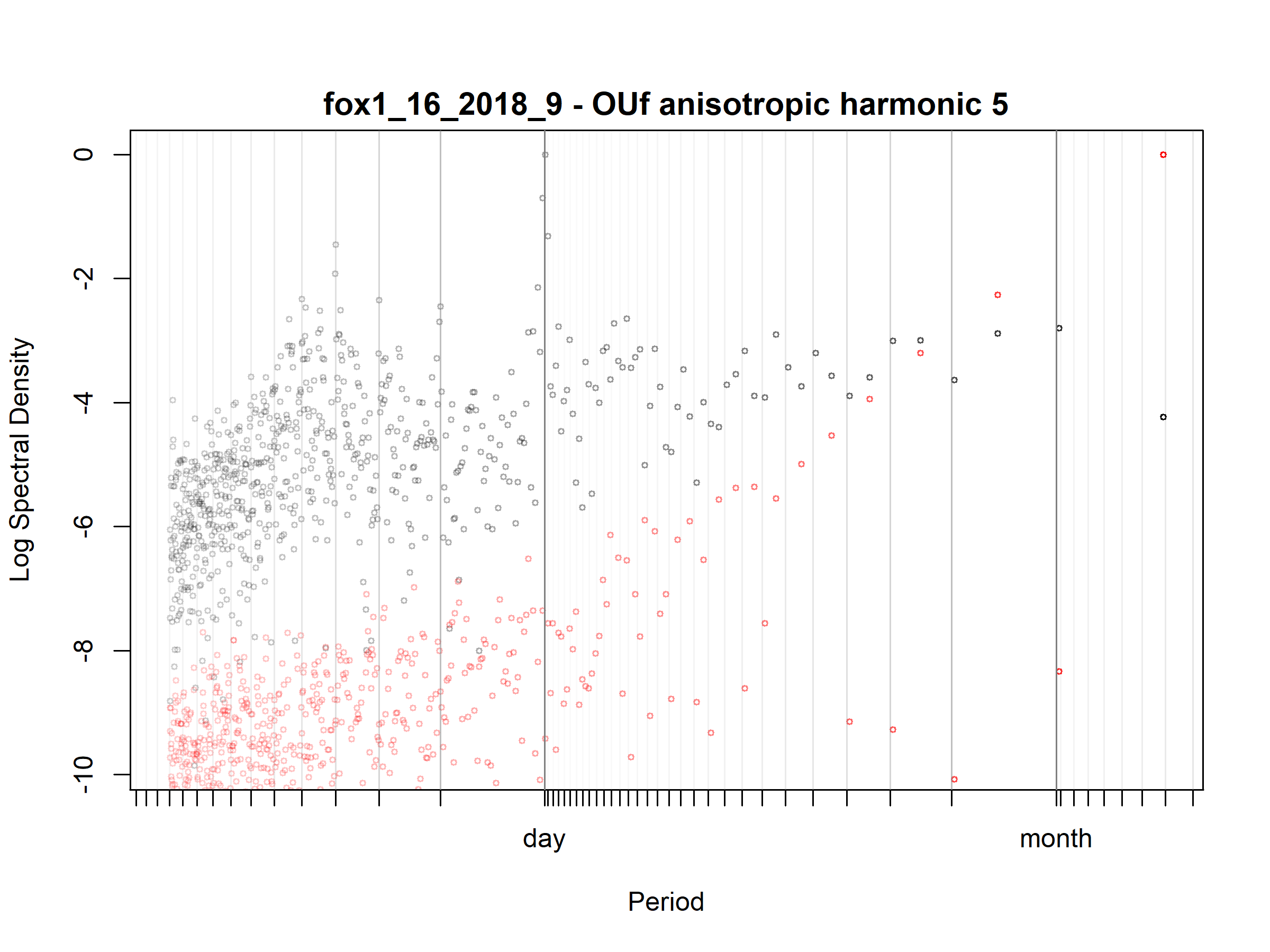

Figure S3.44. Fox; fox1_16_2018_9; diagnostic periodogram.

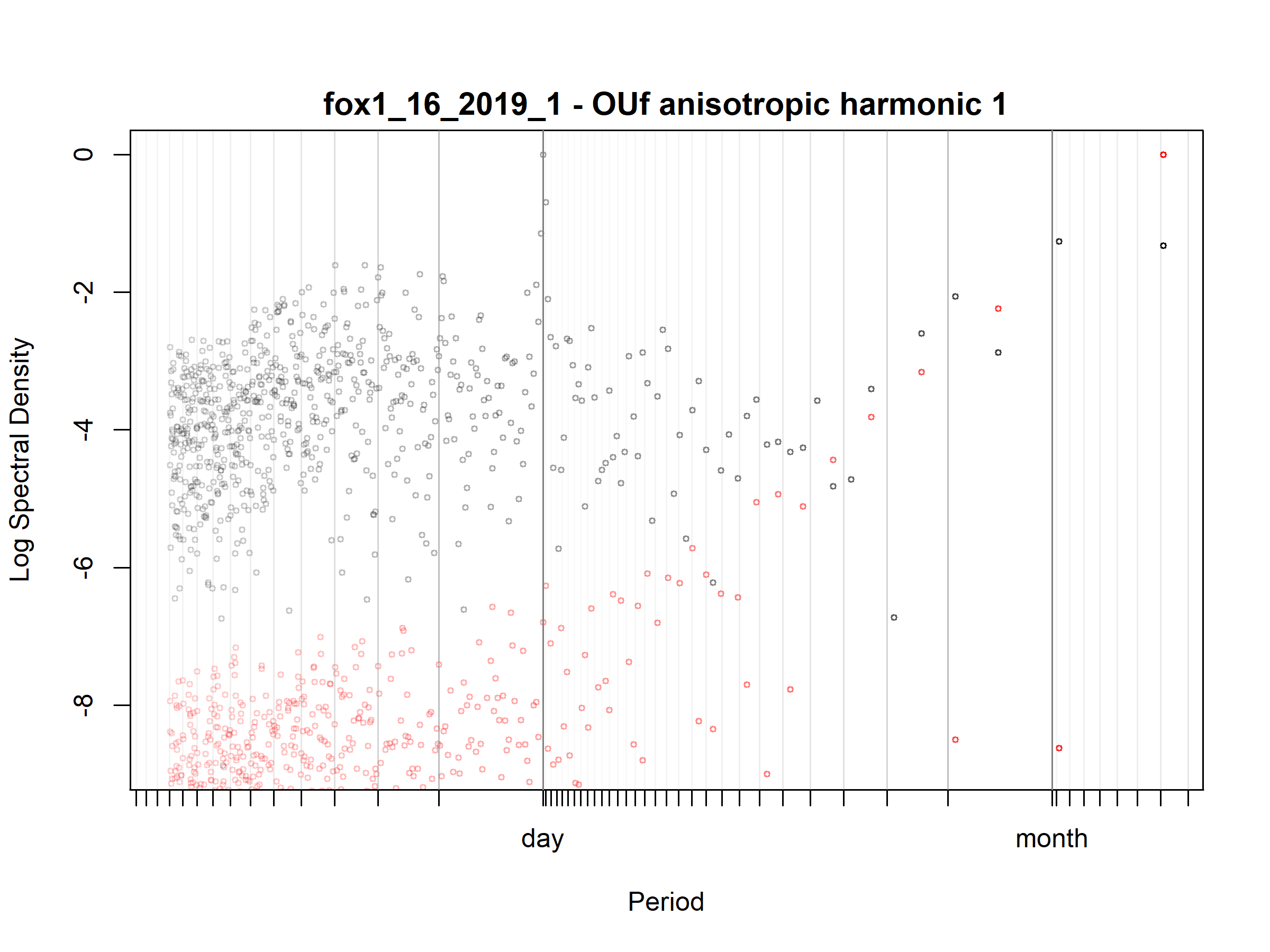

Figure S3.45. Fox; fox1_16_2019_1; diagnostic periodogram.

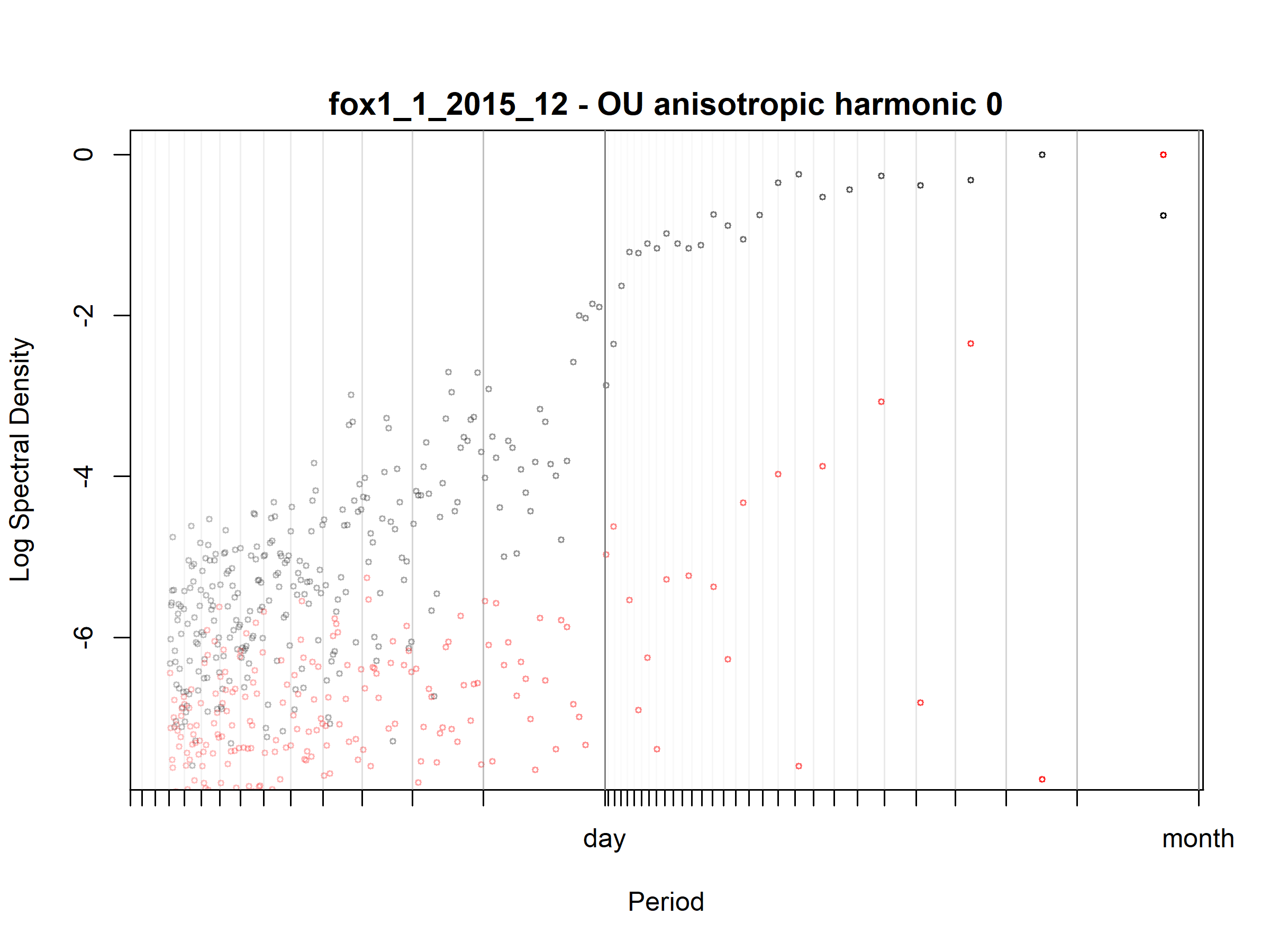

Figure S3.46. Fox; fox1_1_2015_12; diagnostic periodogram.

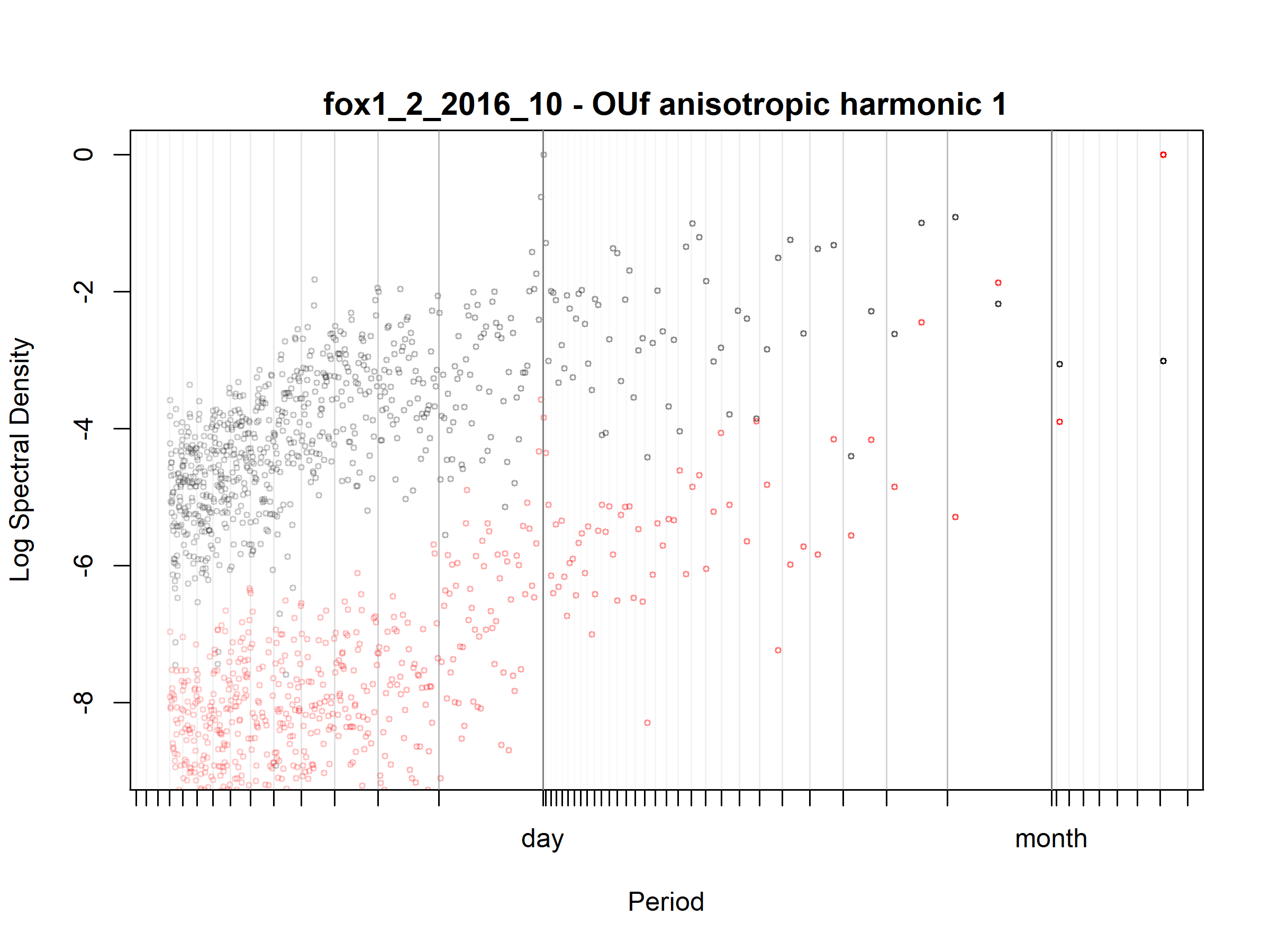

Figure S3.47. Fox; fox1_2_2016_10; diagnostic periodogram.

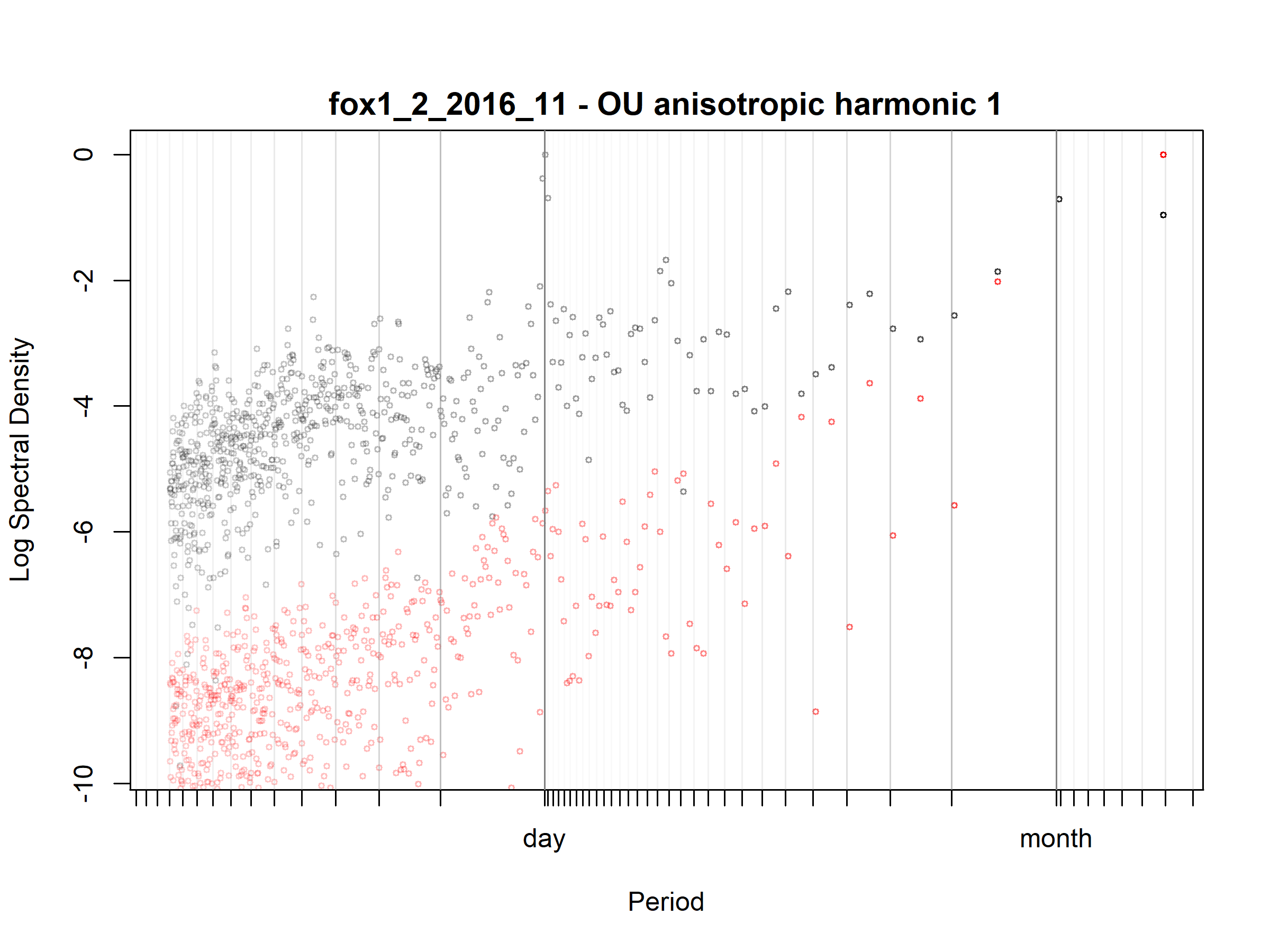

Figure S3.48. Fox; fox1_2_2016_11; diagnostic periodogram.

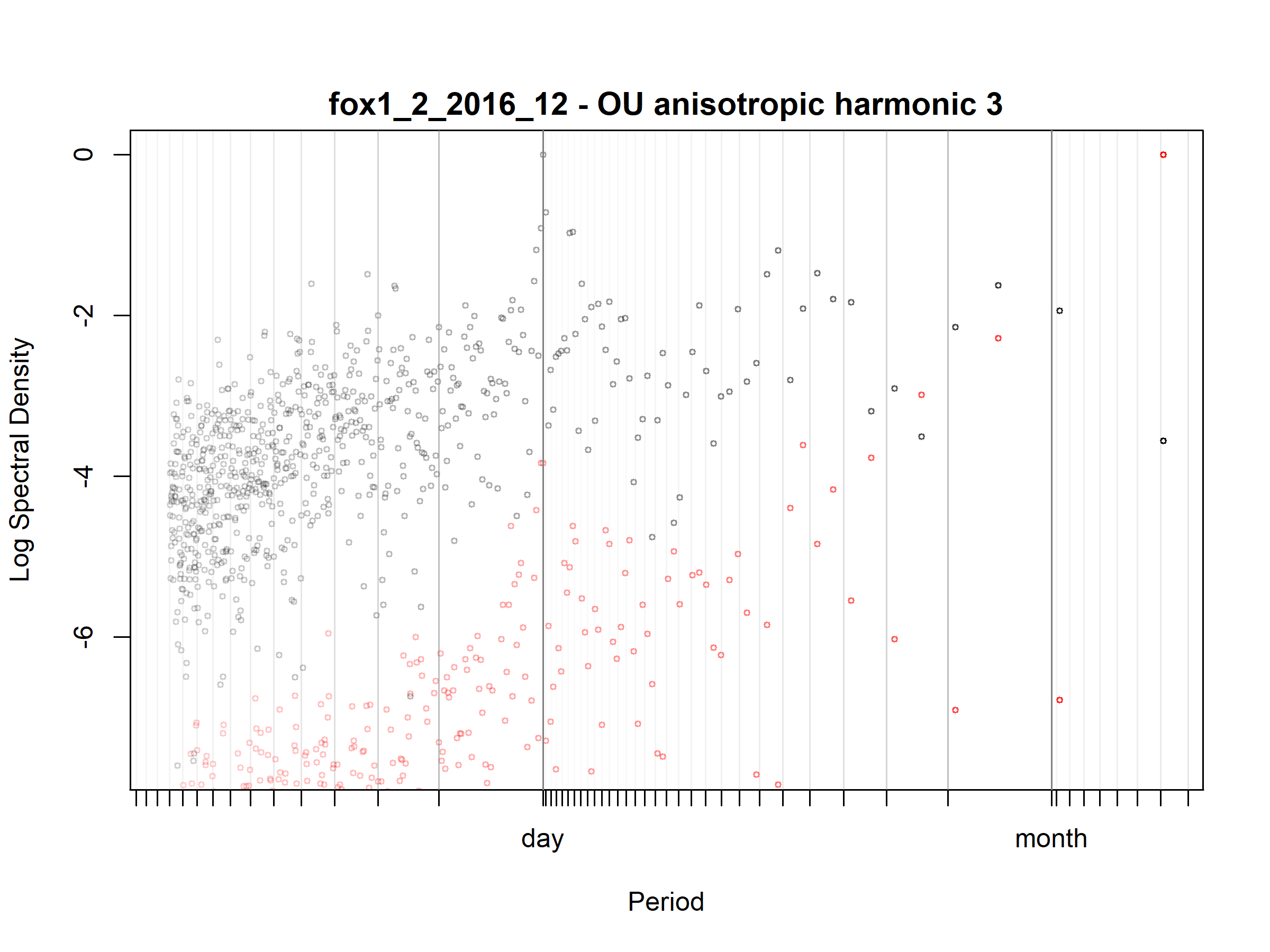

Figure S3.49. Fox; fox1_2_2016_12; diagnostic periodogram.

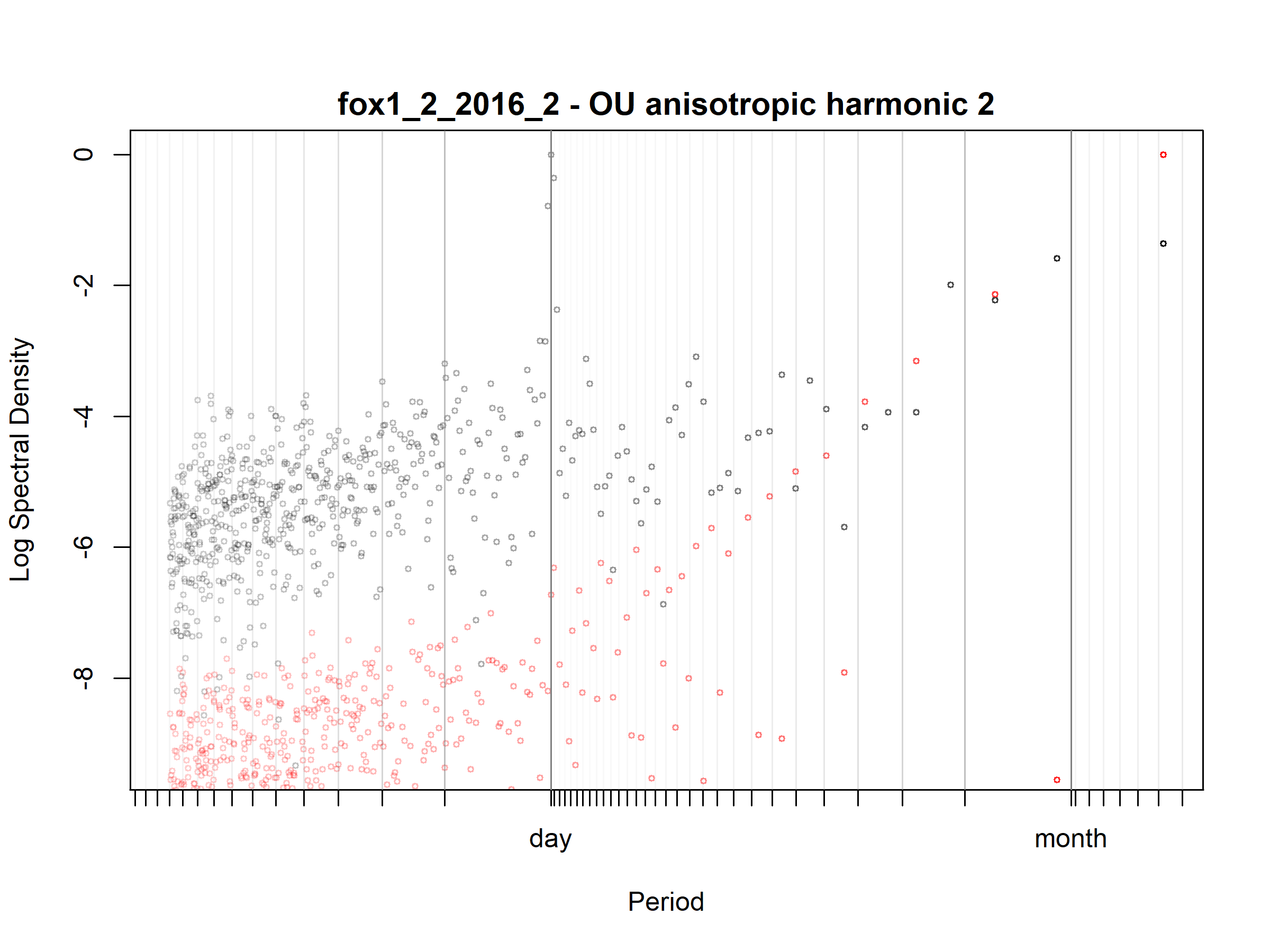

Figure S3.50. Fox; fox1_2_2016_2; diagnostic periodogram.

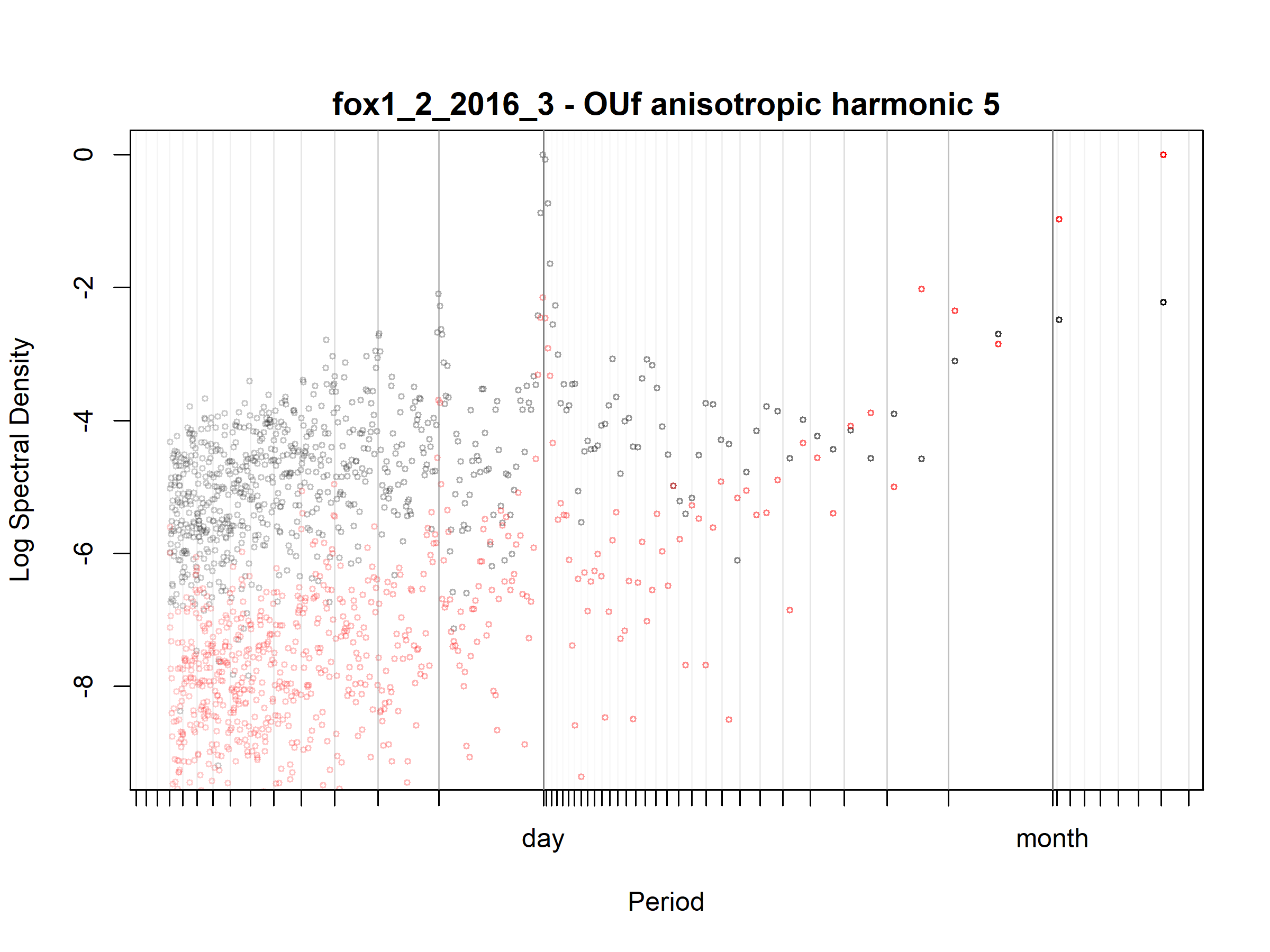

Figure S3.51. Fox; fox1_2_2016_3; diagnostic periodogram.

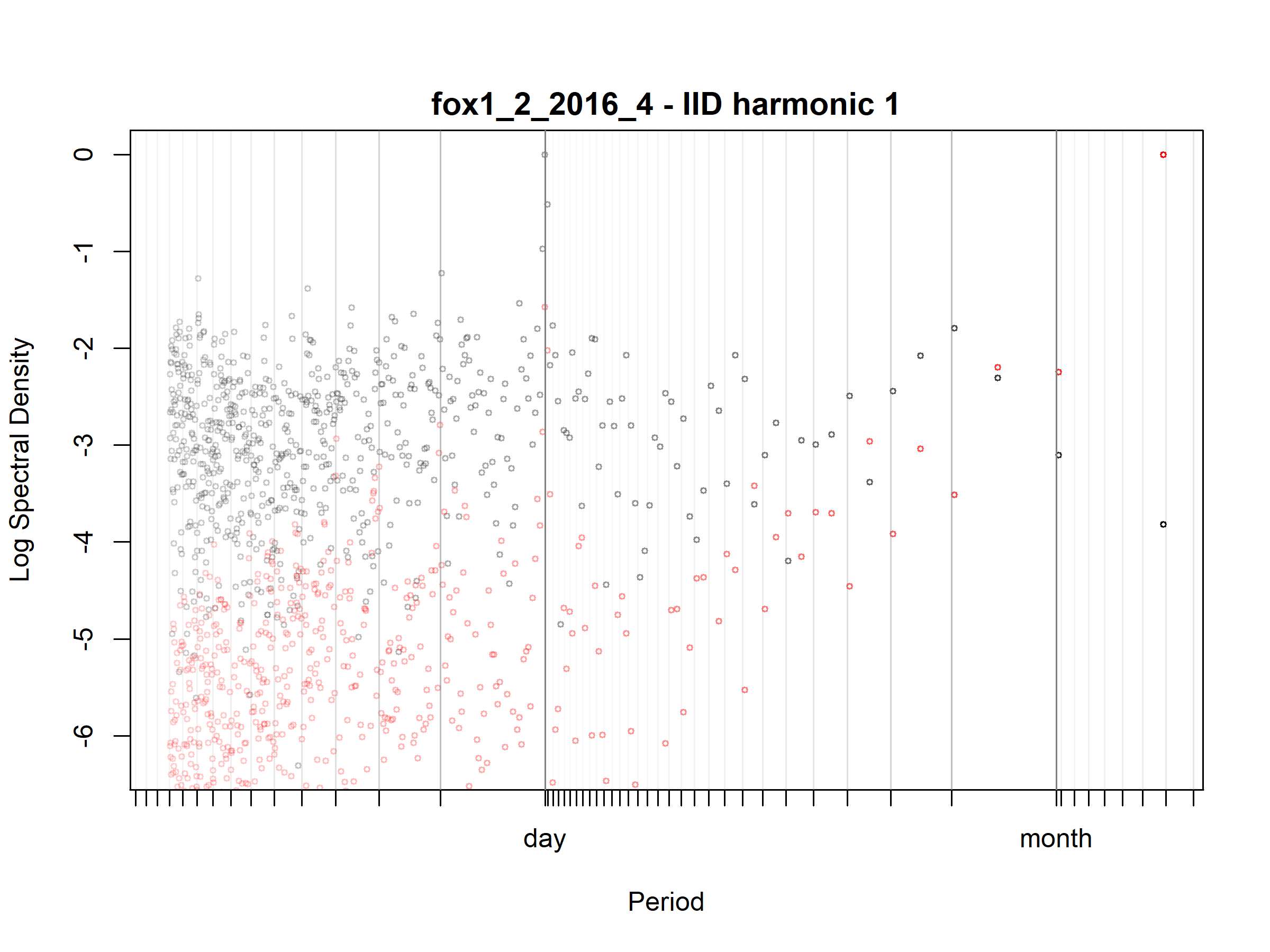

Figure S3.52. Fox; fox1_2_2016_4; diagnostic periodogram.

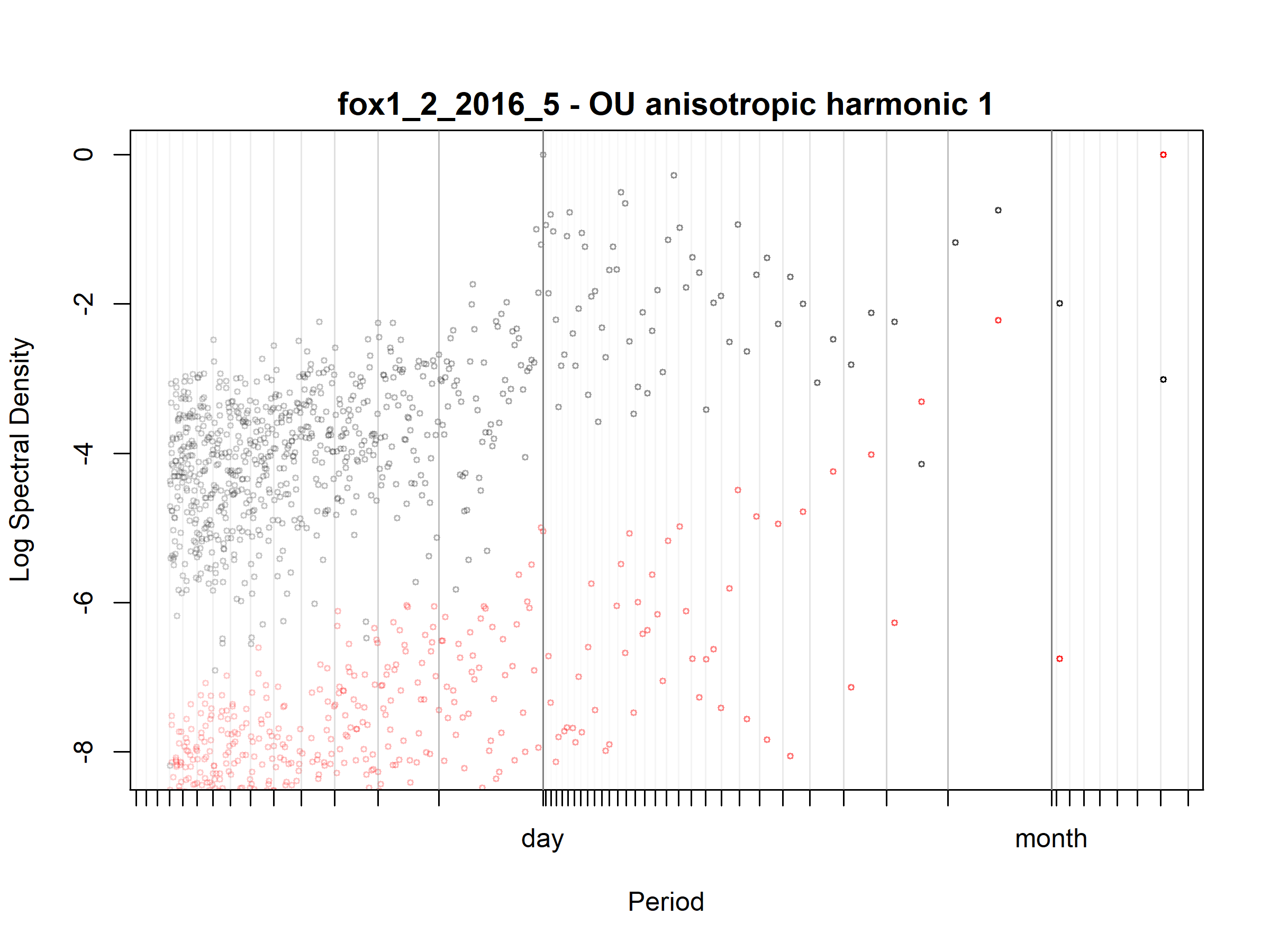

Figure S3.53. Fox; fox1_2_2016_5; diagnostic periodogram.

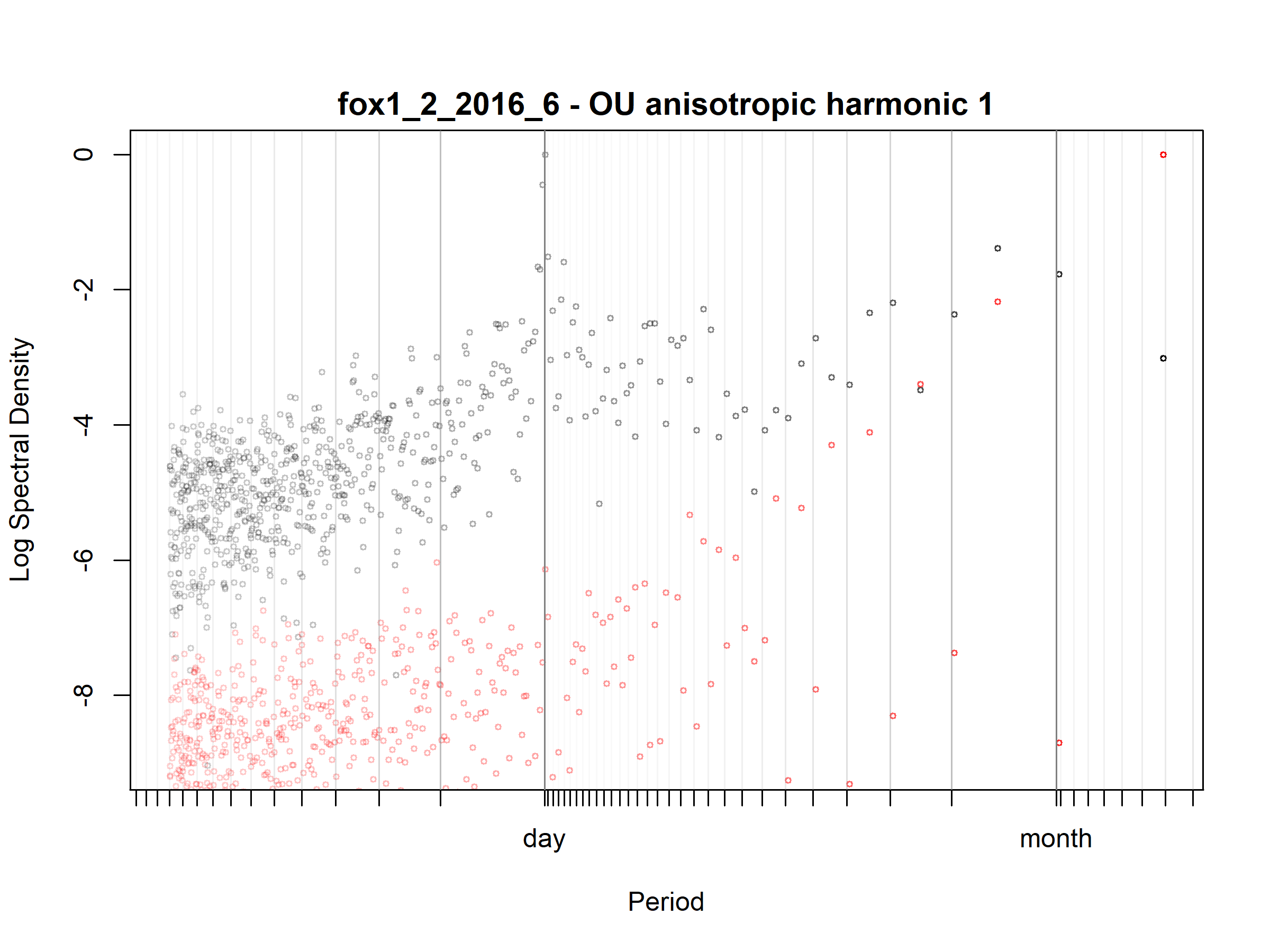

Figure S3.54. Fox; fox1_2_2016_6; diagnostic periodogram.

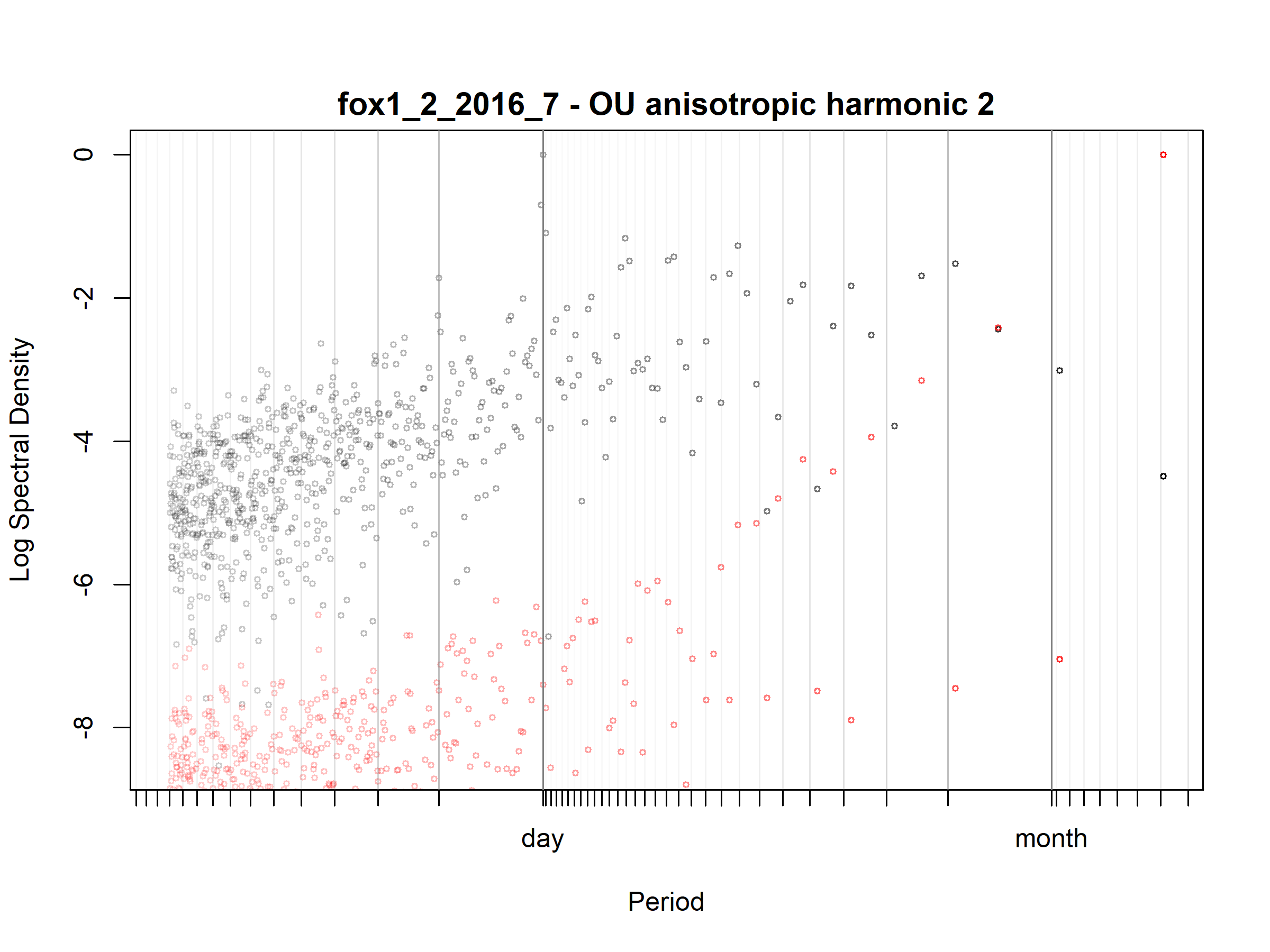

Figure S3.55. Fox; fox1_2_2016_7; diagnostic periodogram.

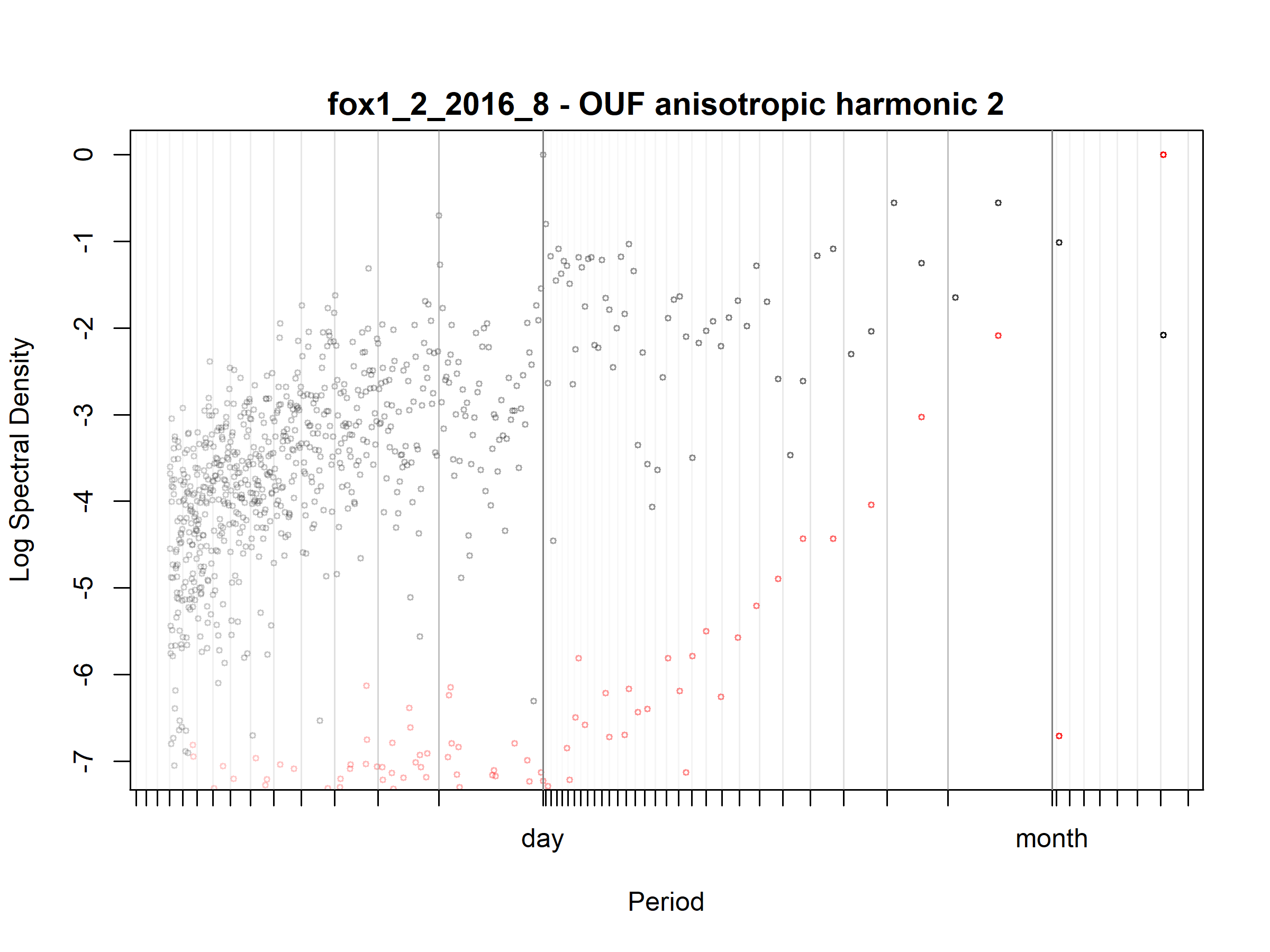

Figure S3.56. Fox; fox1_2_2016_8; diagnostic periodogram.

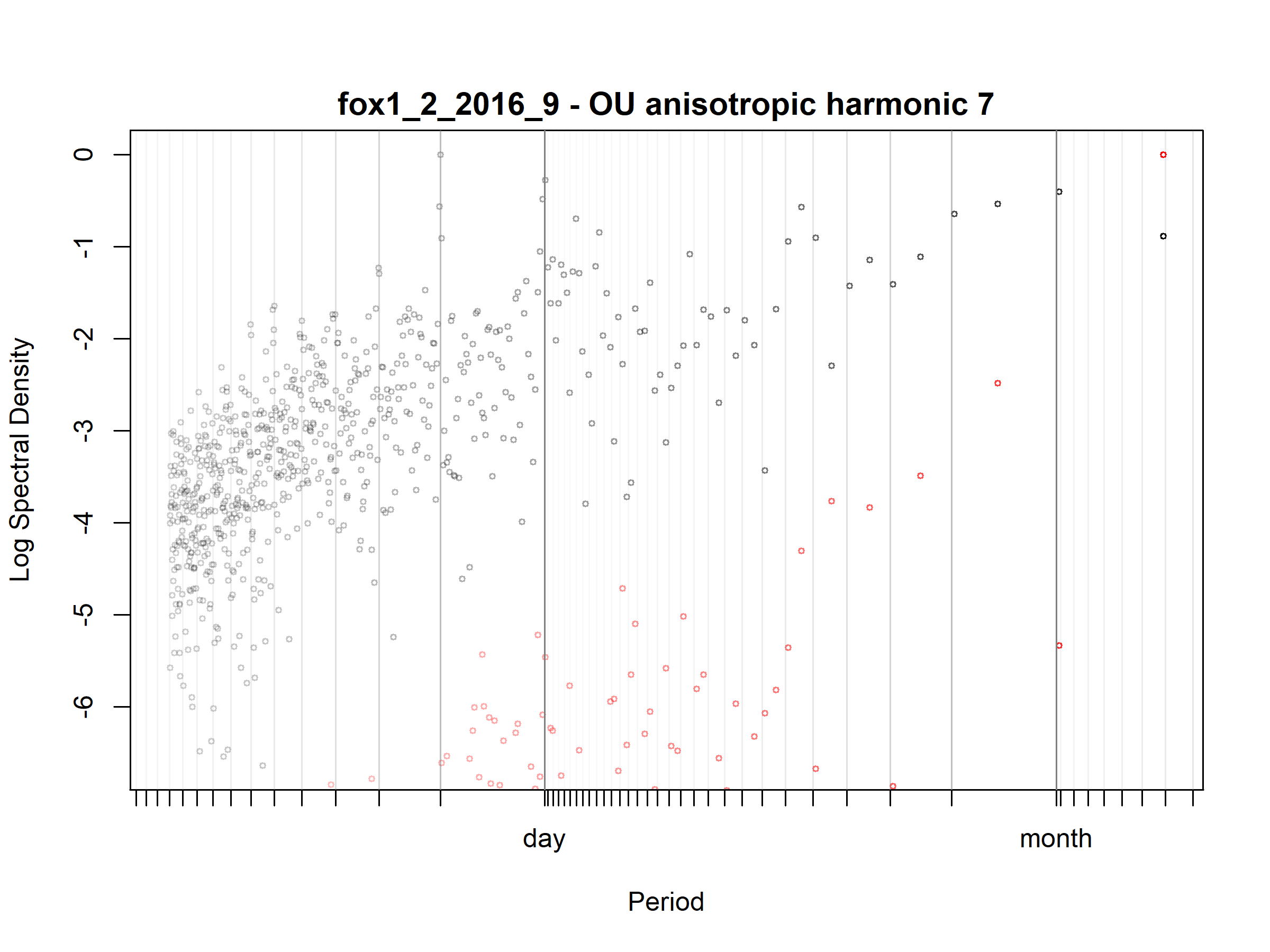

Figure S3.57. Fox; fox1_2_2016_9; diagnostic periodogram.

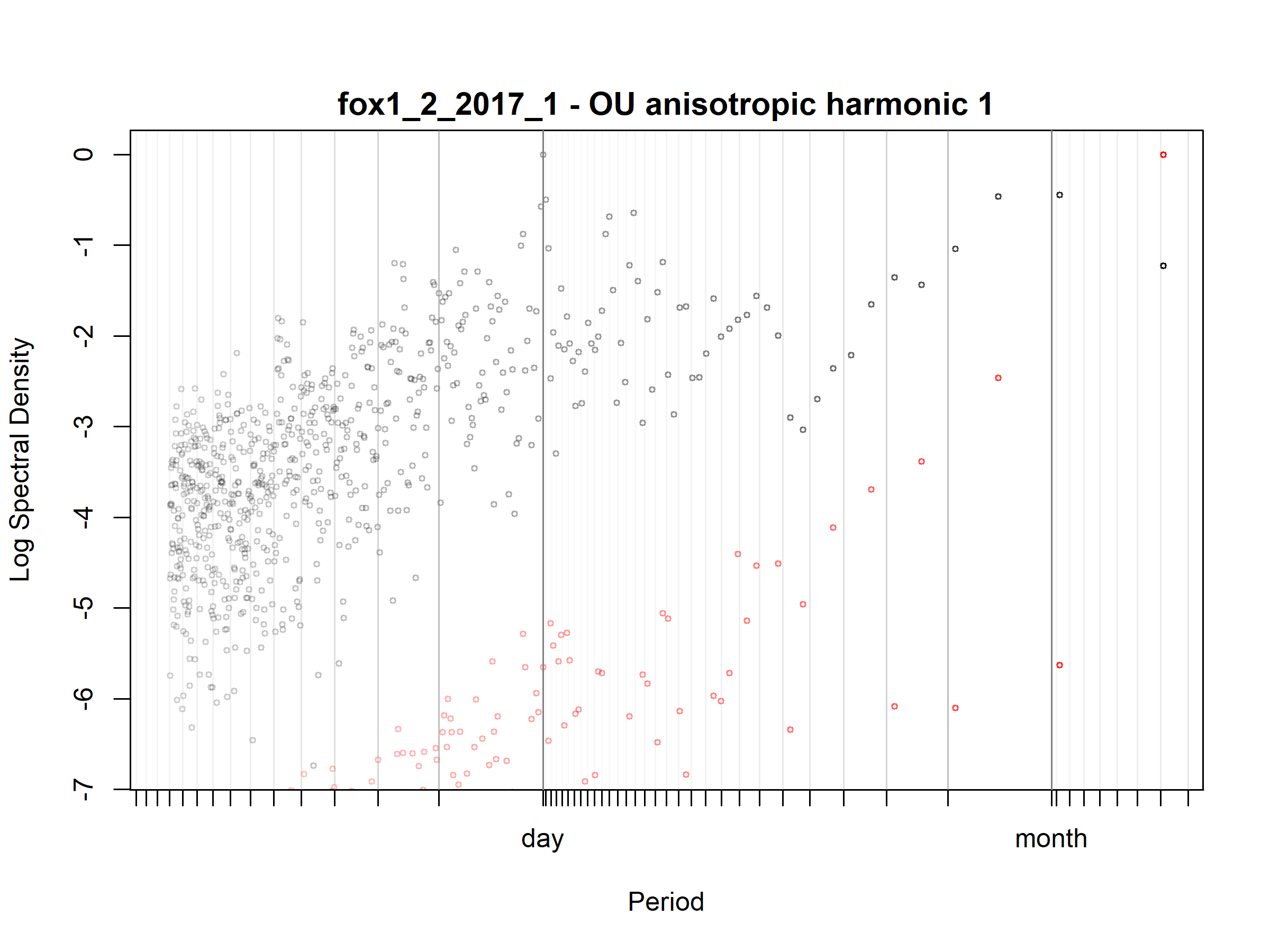

Figure S3.58. Fox; fox1_2_2017_1; diagnostic periodogram.

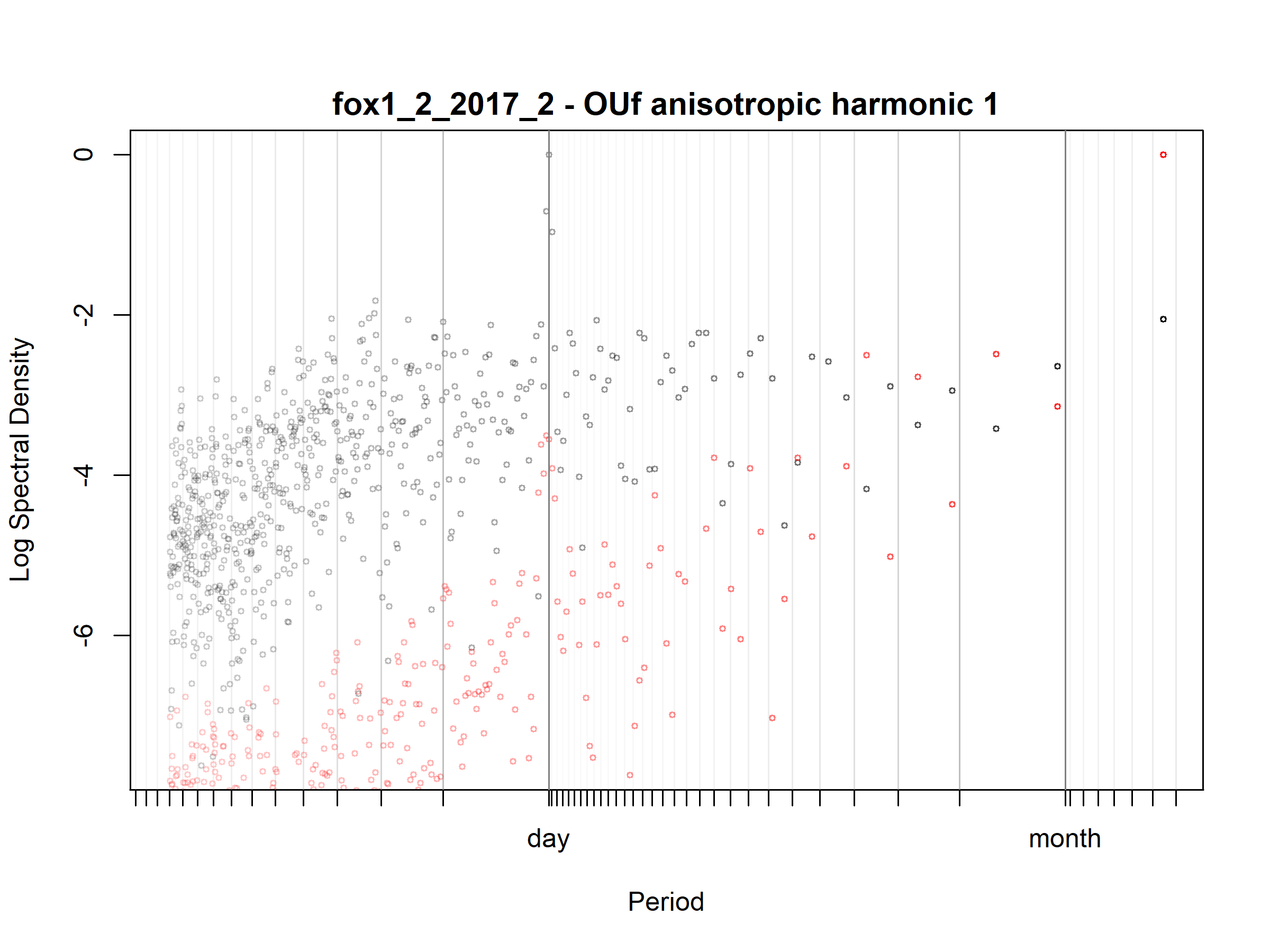

Figure S3.59. Fox; fox1_2_2017_2; diagnostic periodogram.

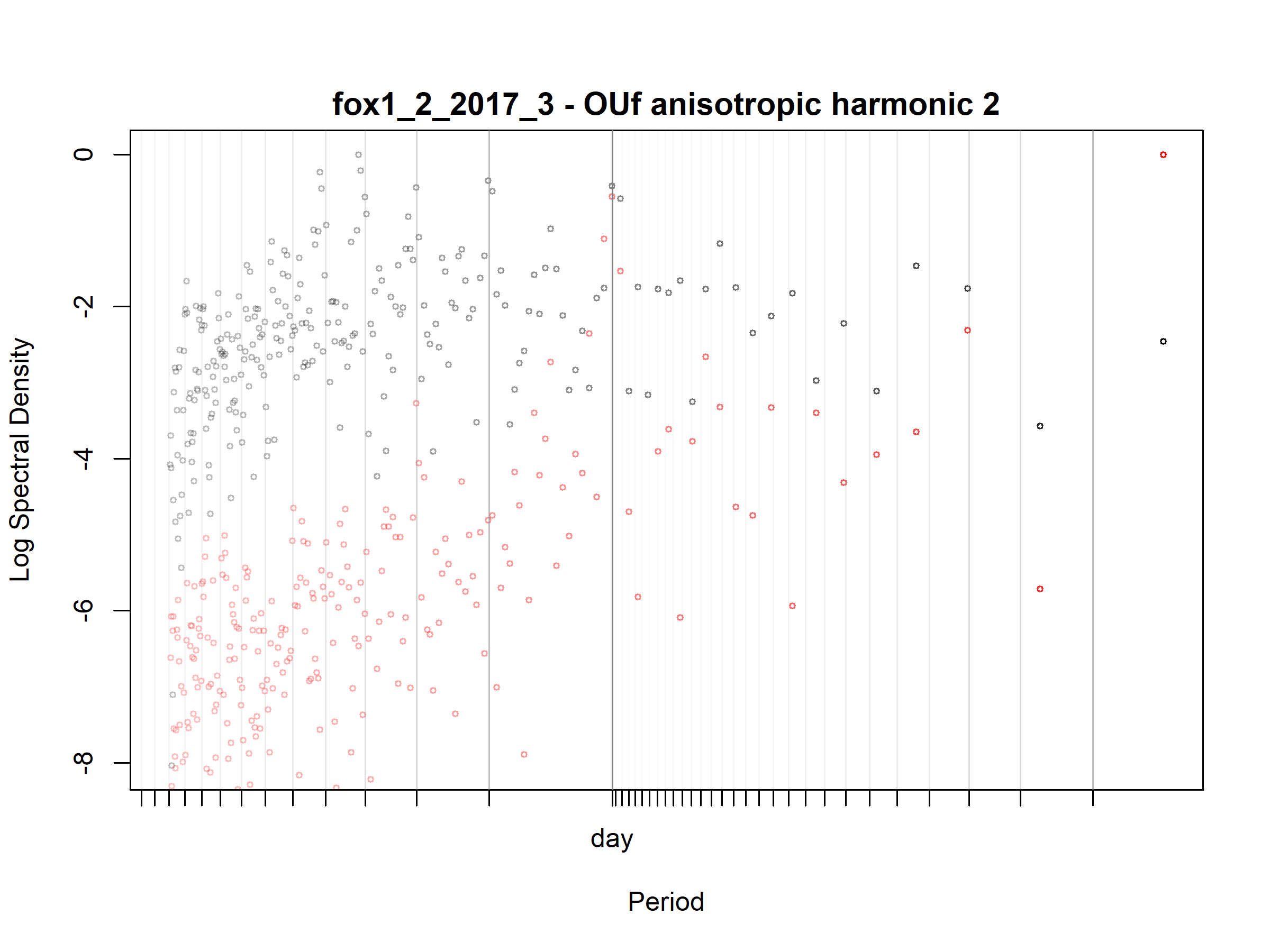

Figure S3.60. Fox; fox1_2_2017_3; diagnostic periodogram.

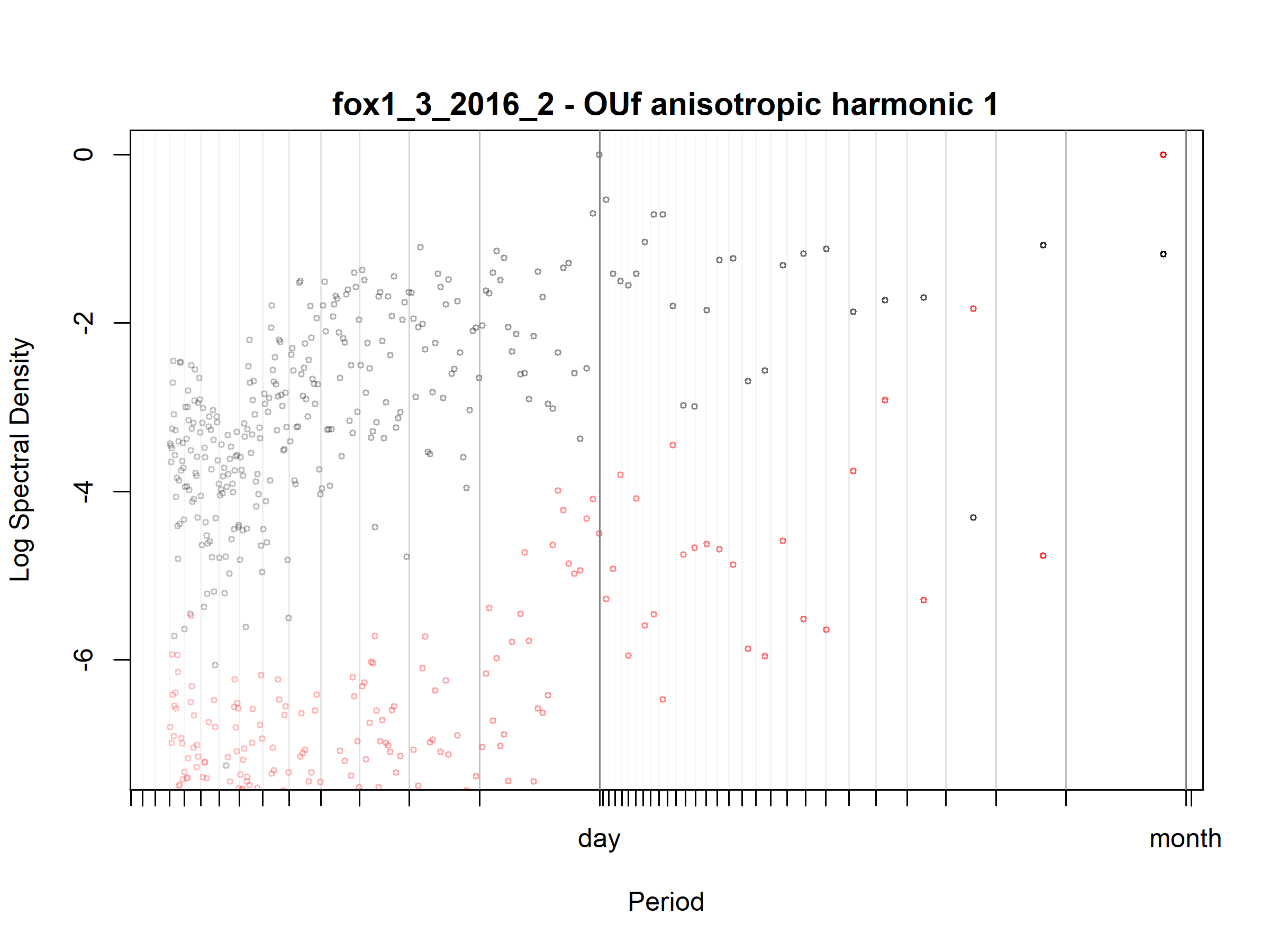

Figure S3.61. Fox; fox1_3_2016_2; diagnostic periodogram.

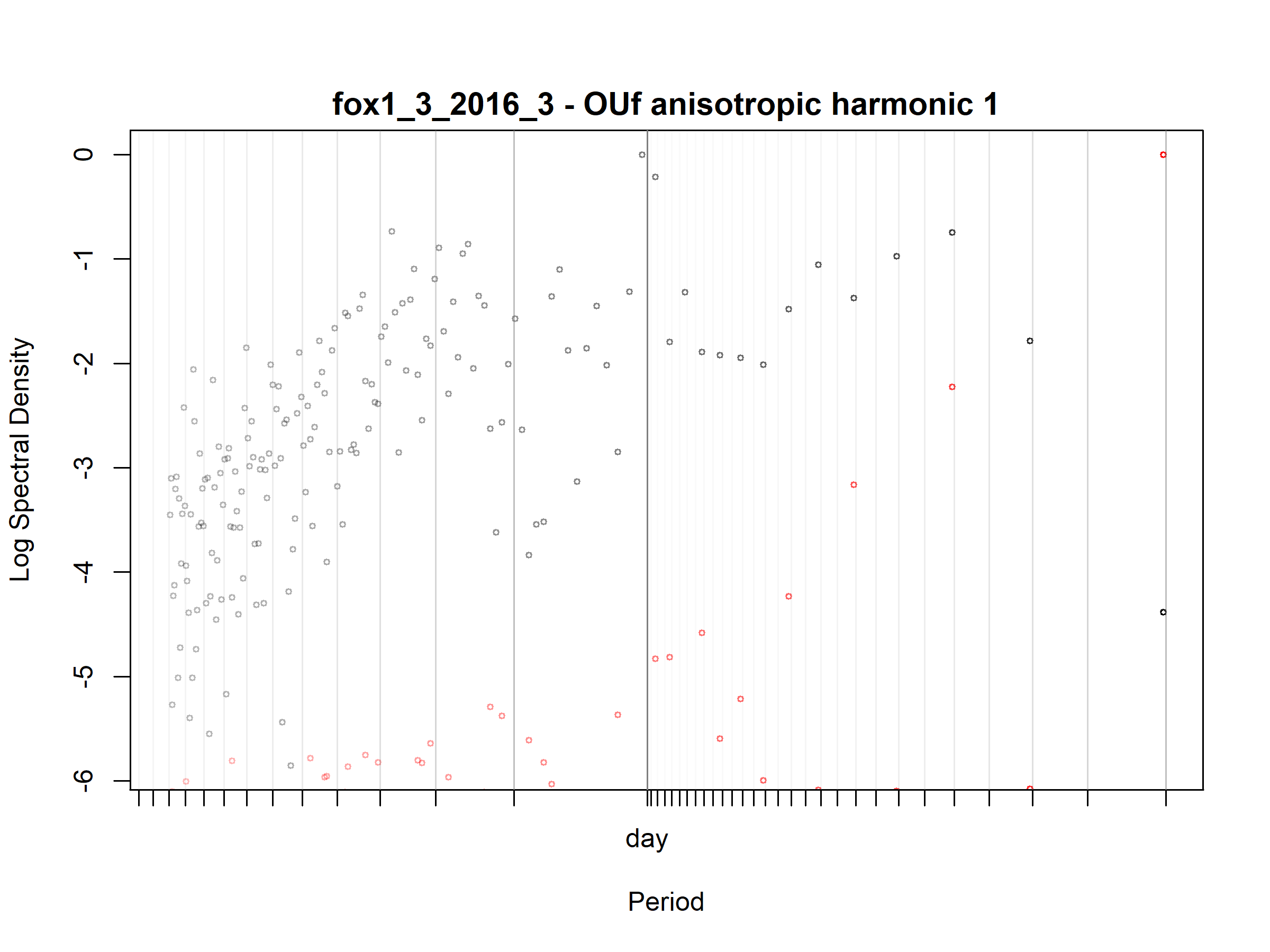

Figure S3.62. Fox; fox1_3_2016_3; diagnostic periodogram.

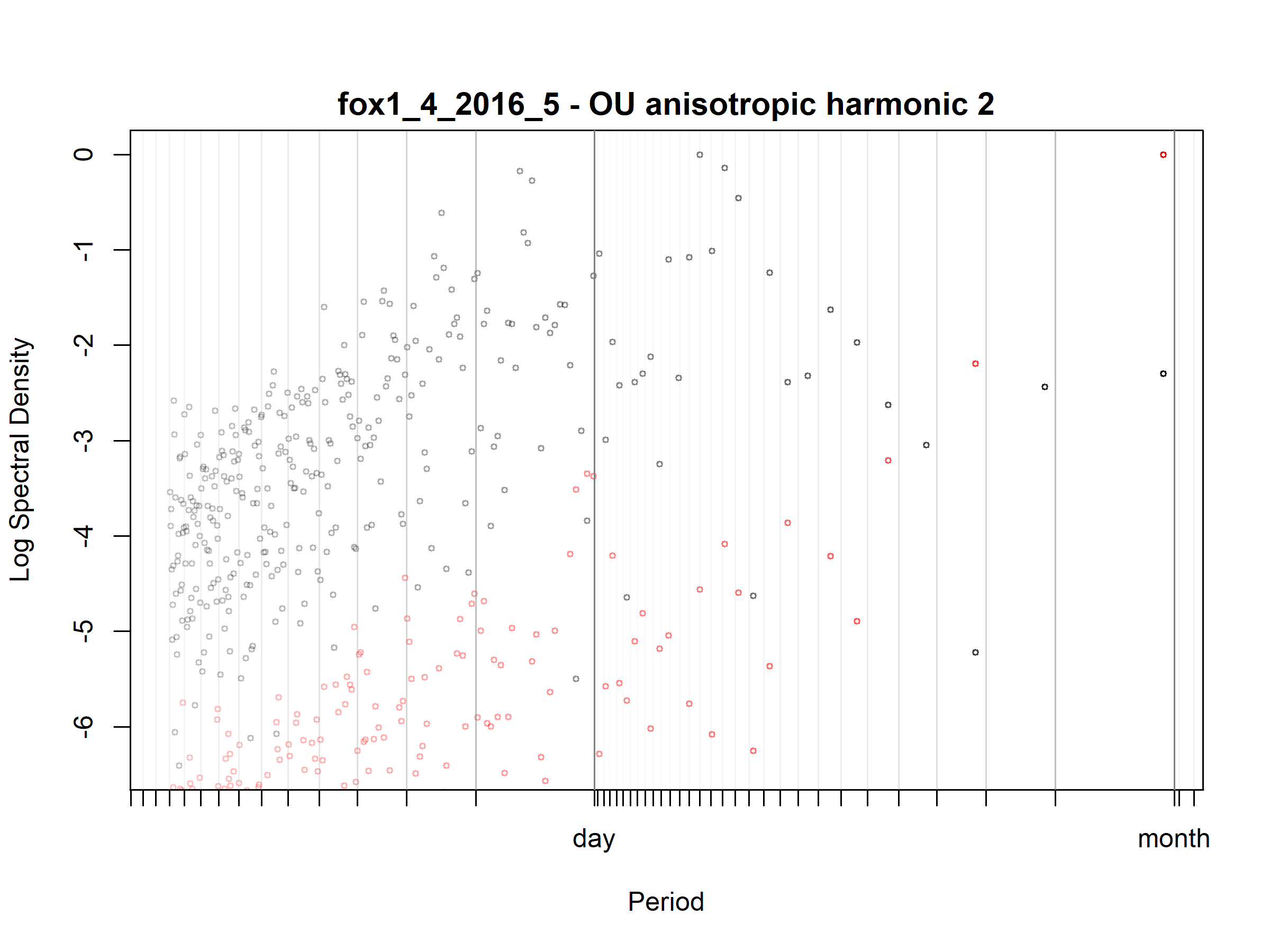

Figure S3.63. Fox; fox1_4_2016_5; diagnostic periodogram.

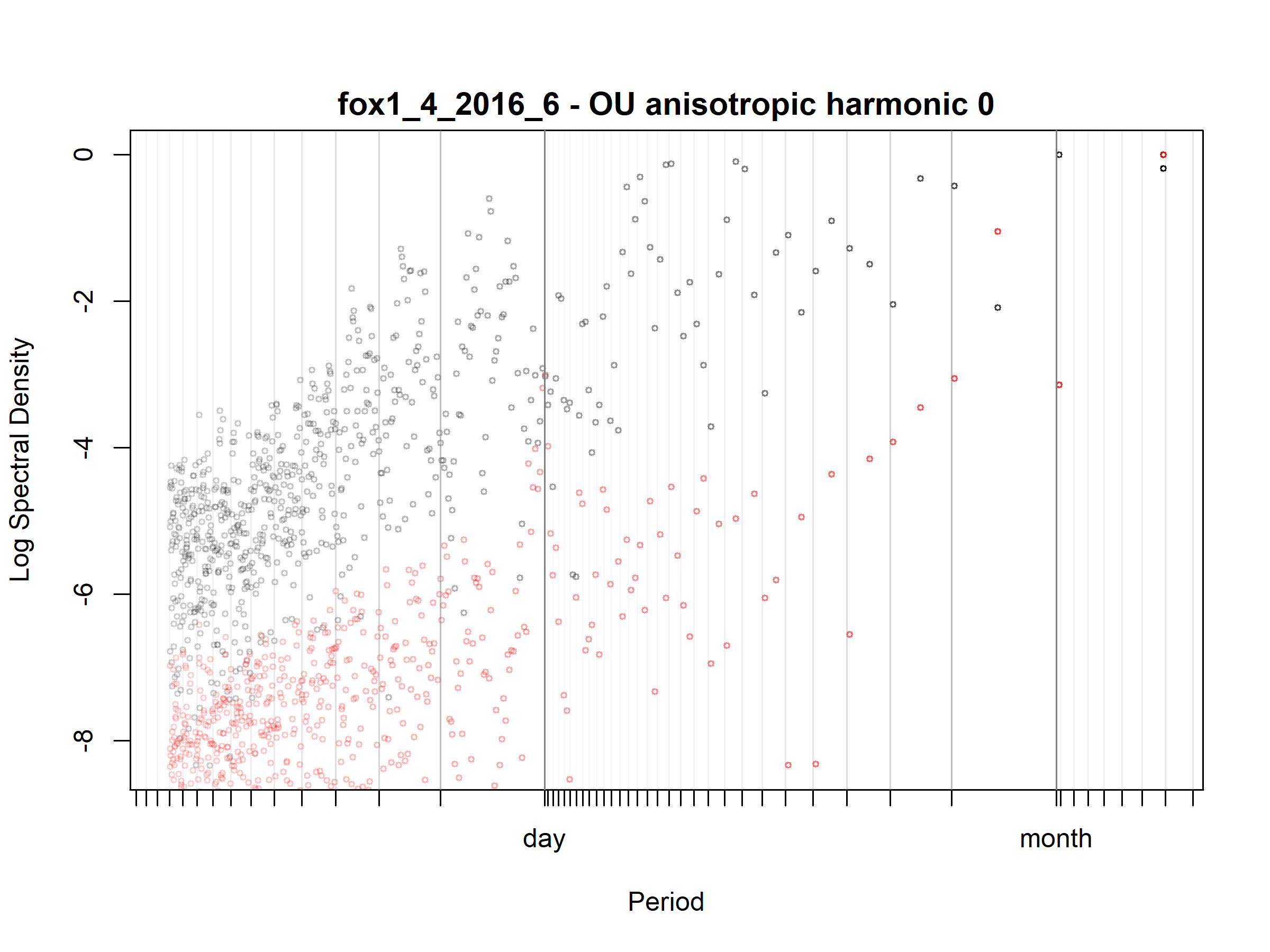

Figure S3.64. Fox; fox1_4_2016_6; diagnostic periodogram.

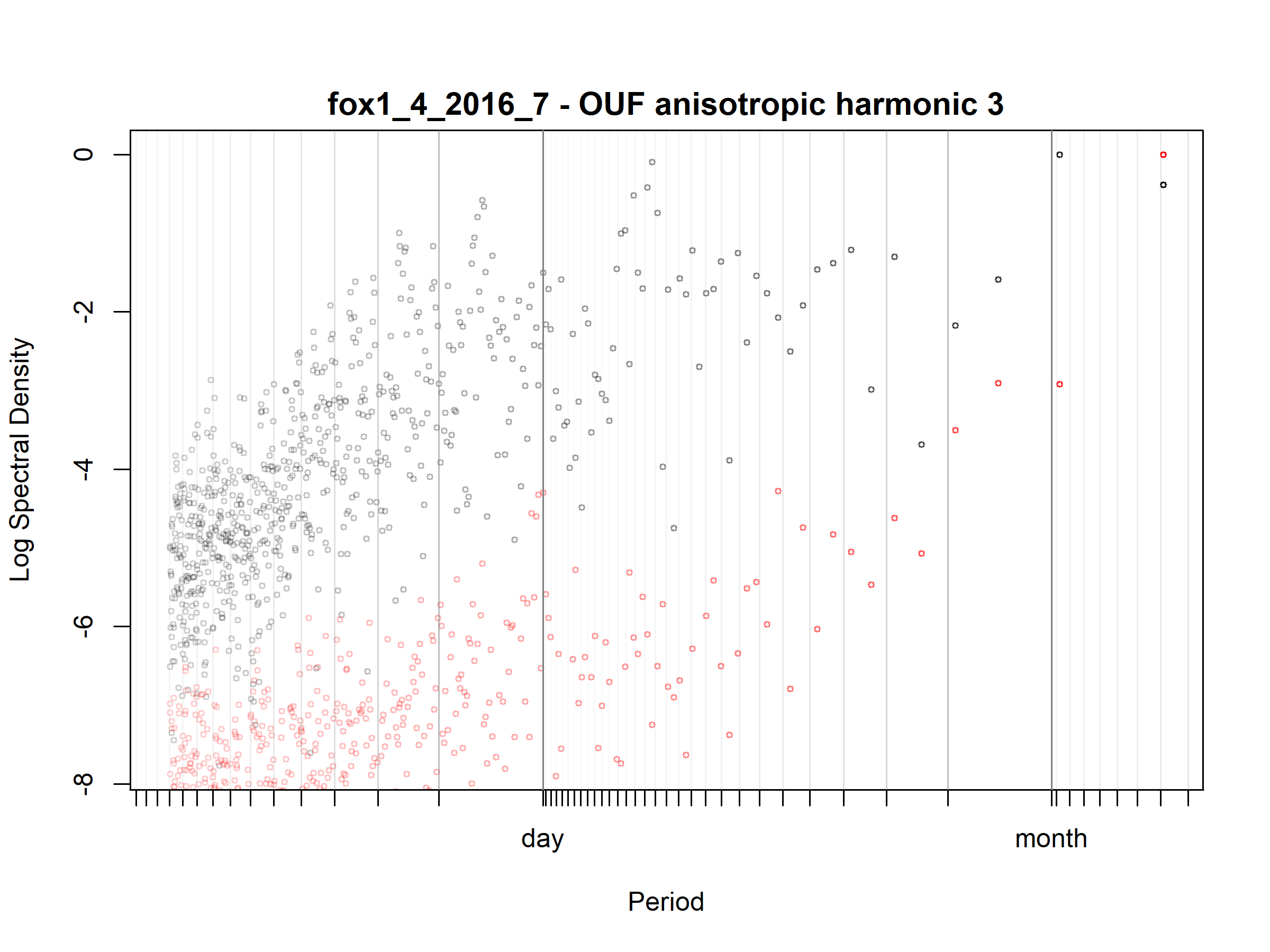

Figure S3.65. Fox; fox1_4_2016_7; diagnostic periodogram.

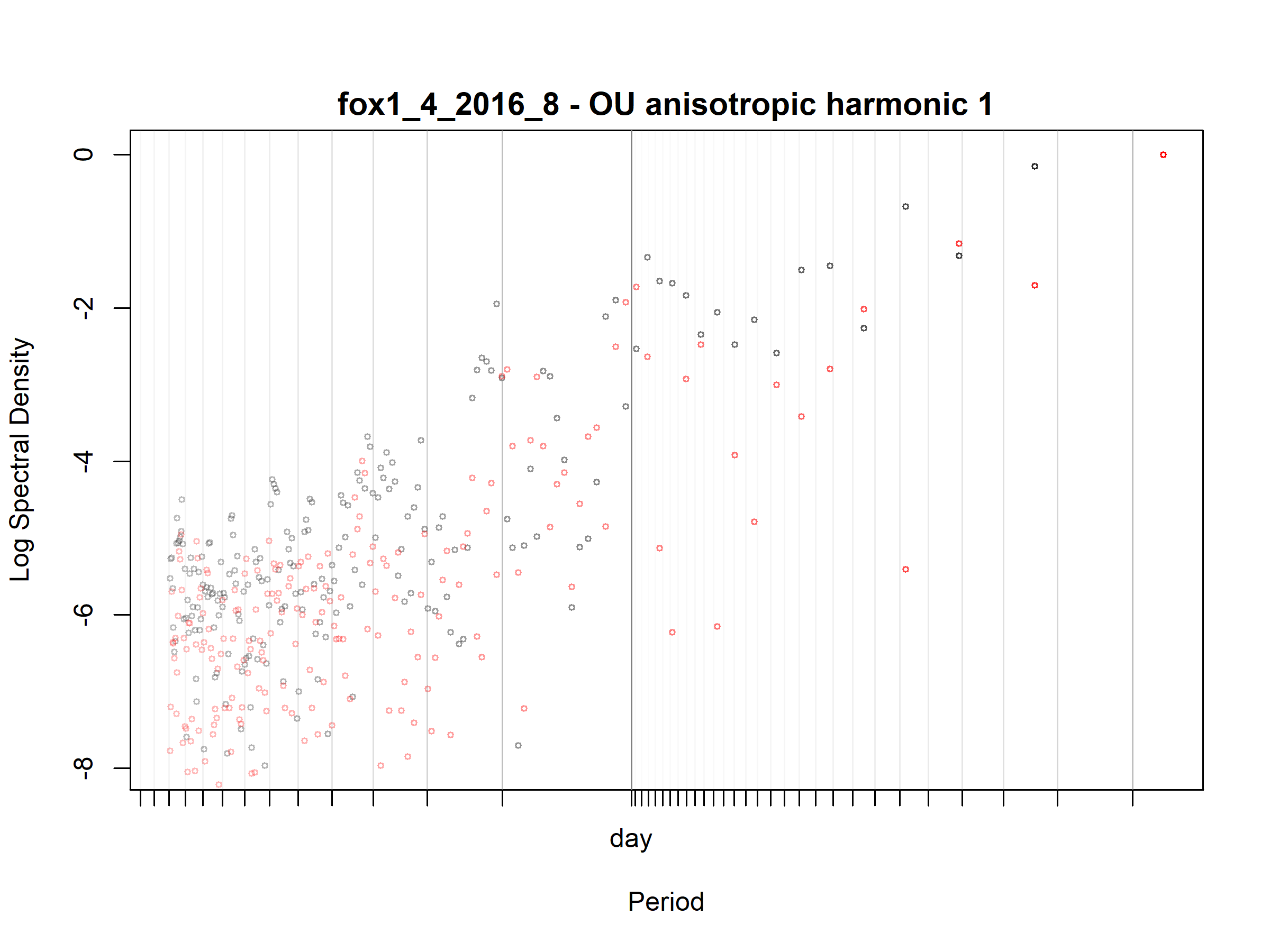

Figure S3.66. Fox; fox1_4_2016_8; diagnostic periodogram.

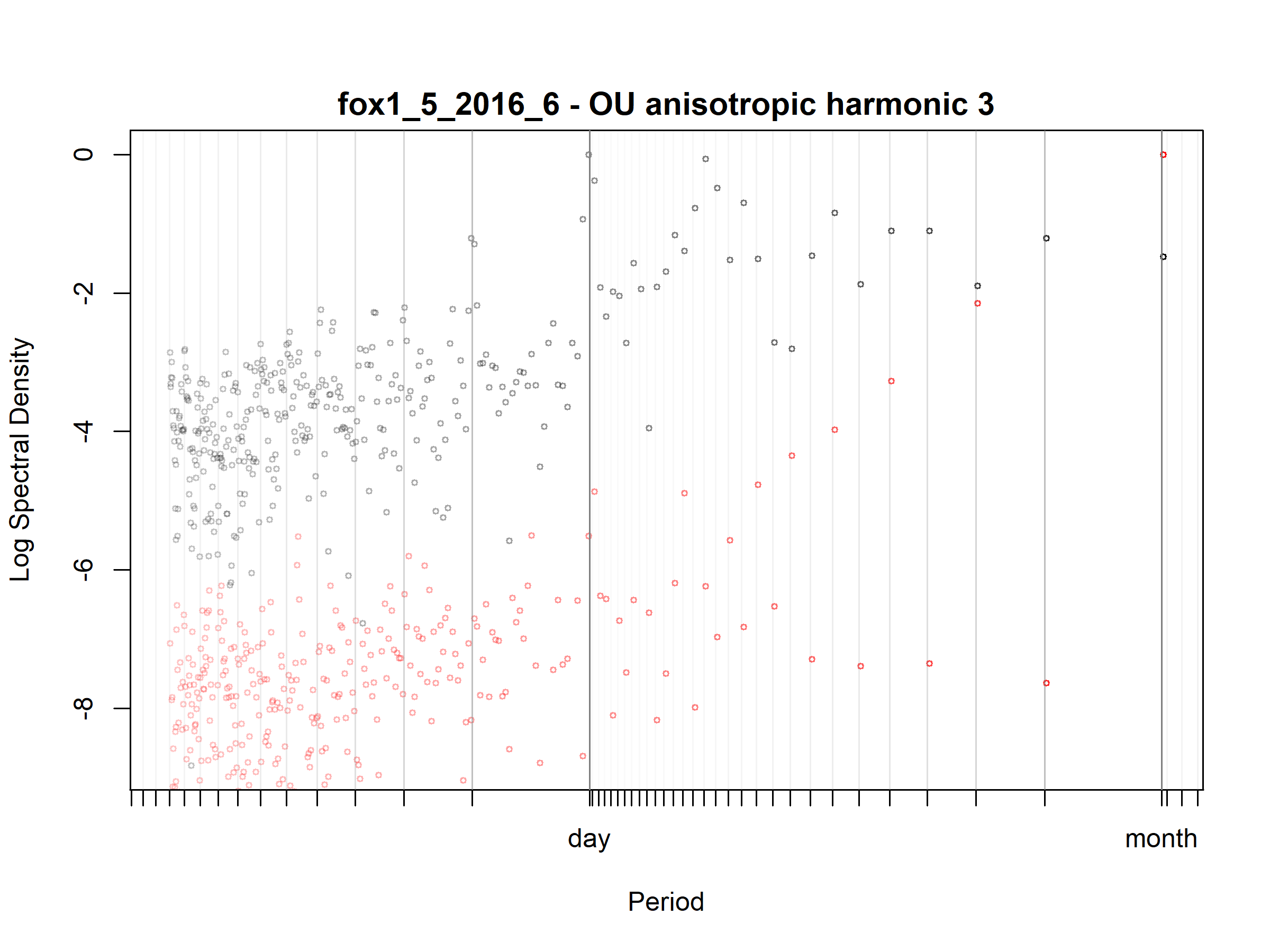

Figure S3.67. Fox; fox1_5_2016_6; diagnostic periodogram.

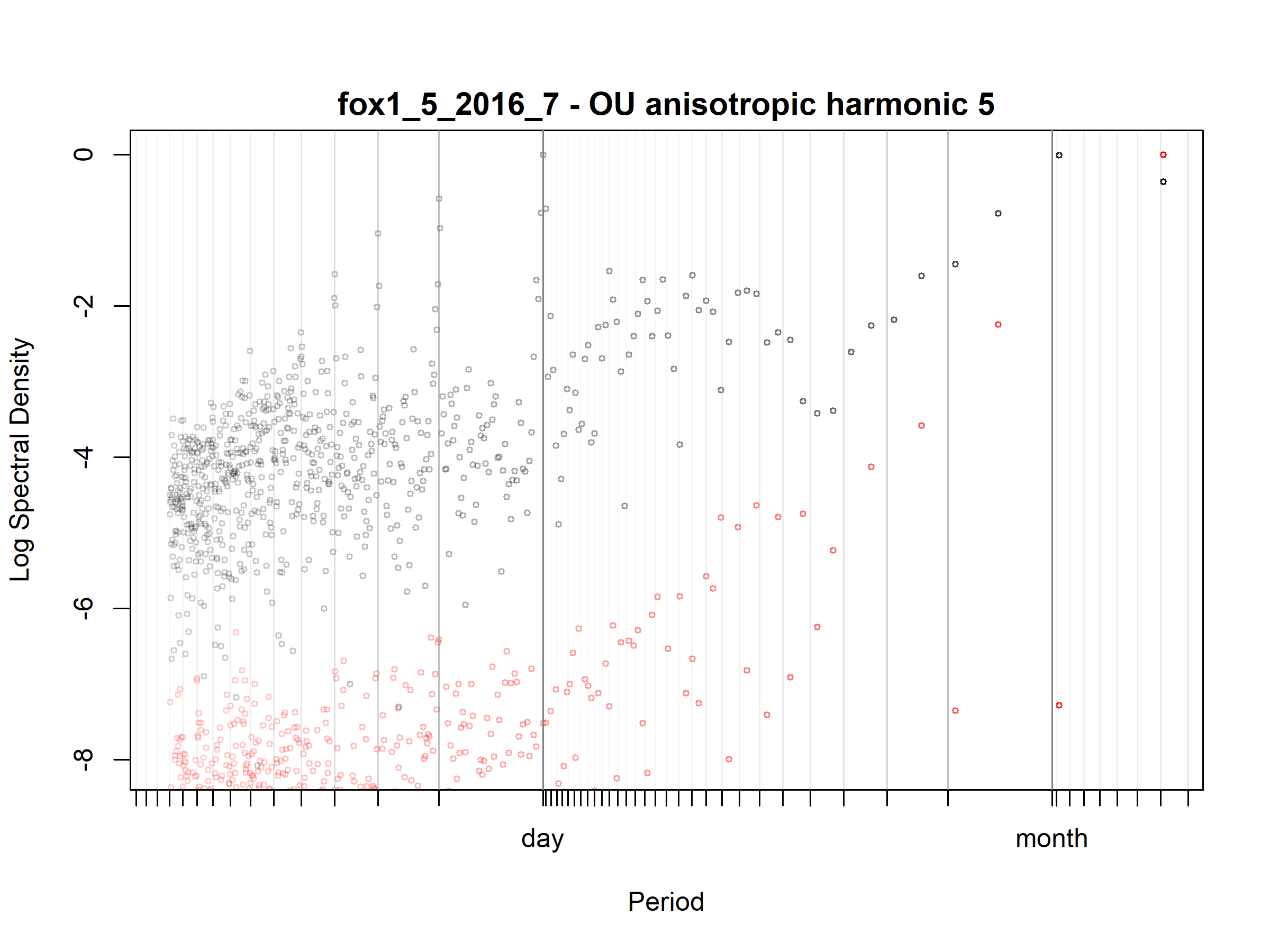

Figure S3.68. Fox; fox1_5_2016_7; diagnostic periodogram.

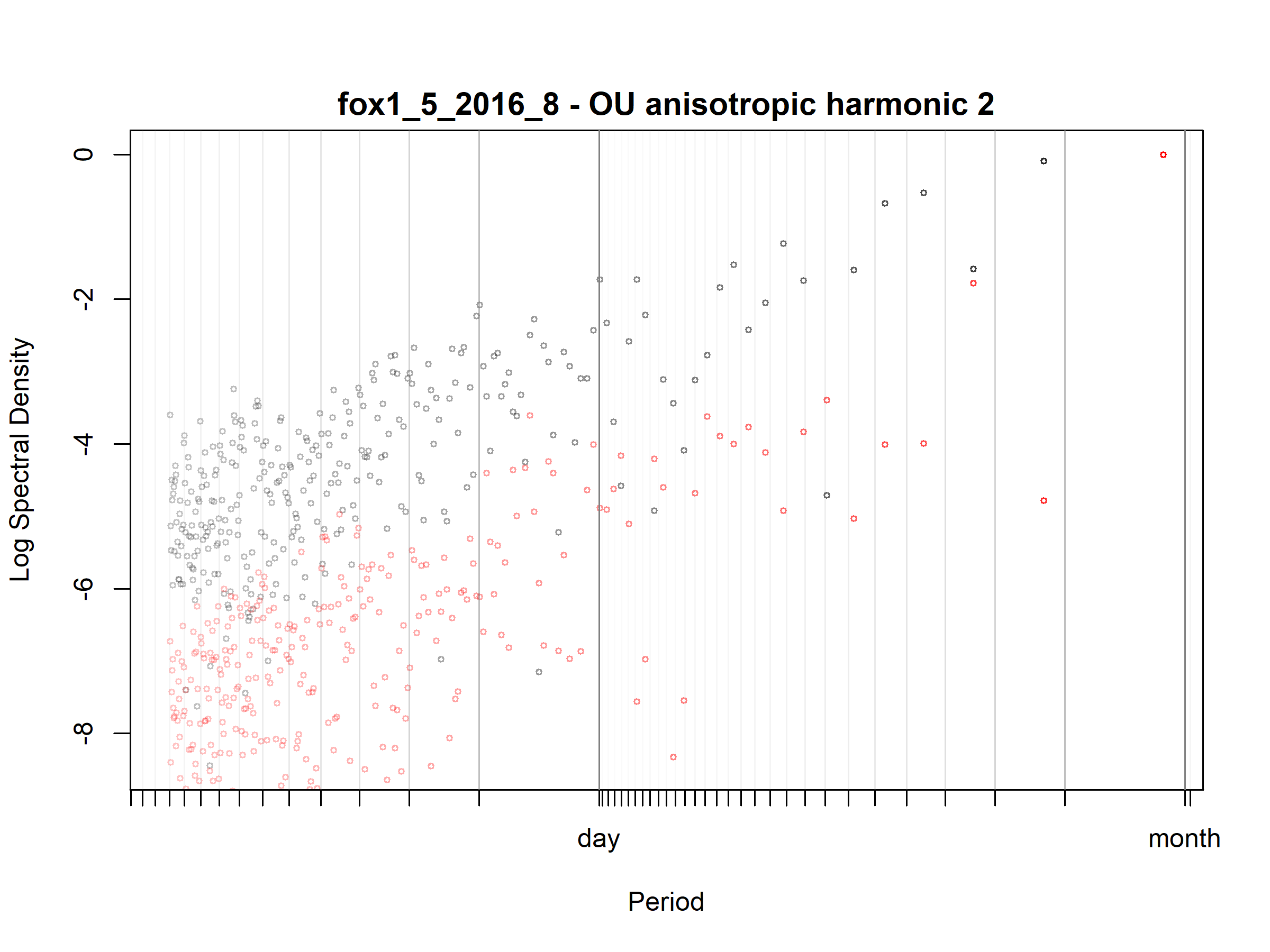

Figure S3.69. Fox; fox1_5_2016_8; diagnostic periodogram.

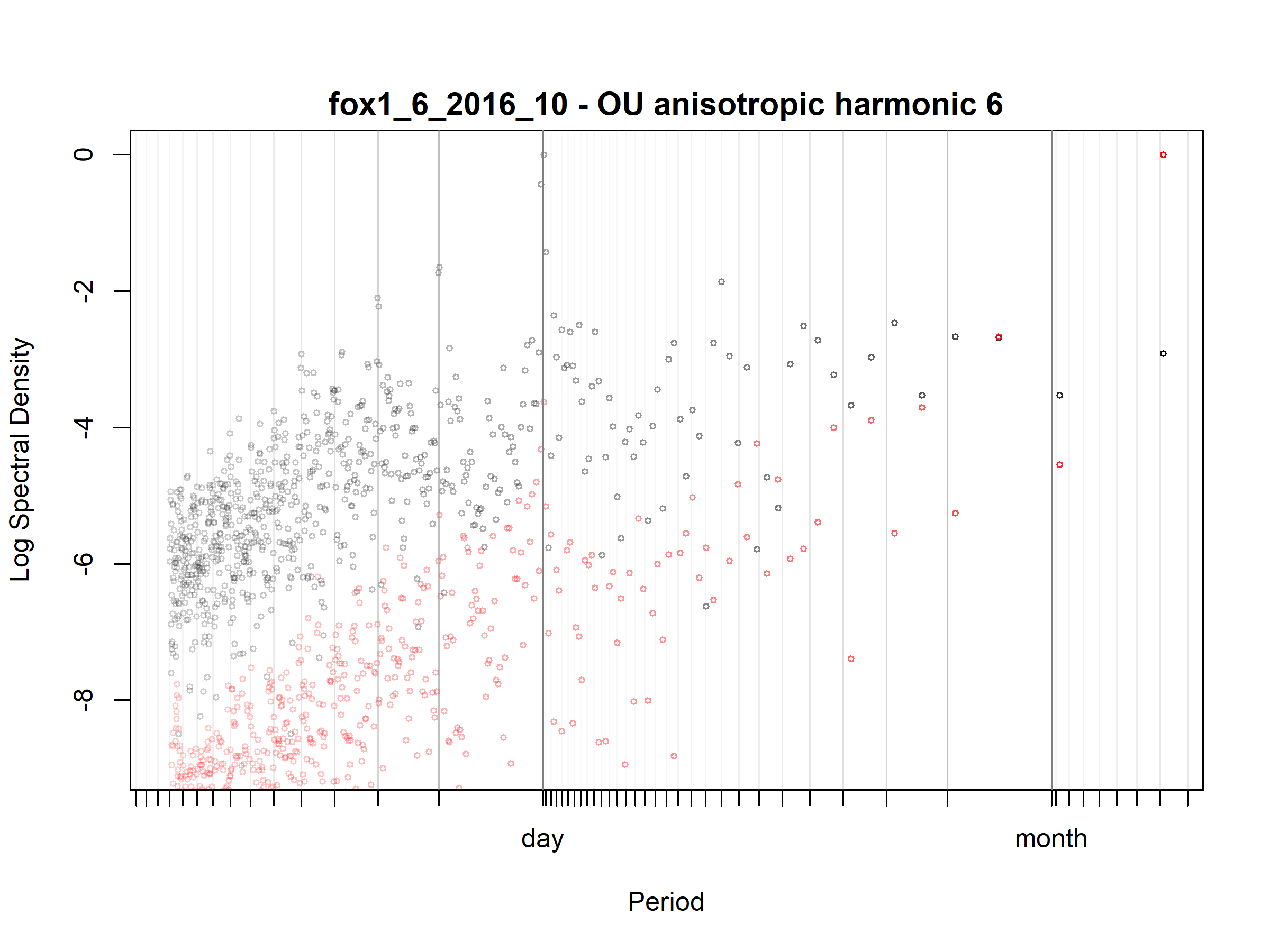

Figure S3.70. Fox; fox1_6_2016_10; diagnostic periodogram.

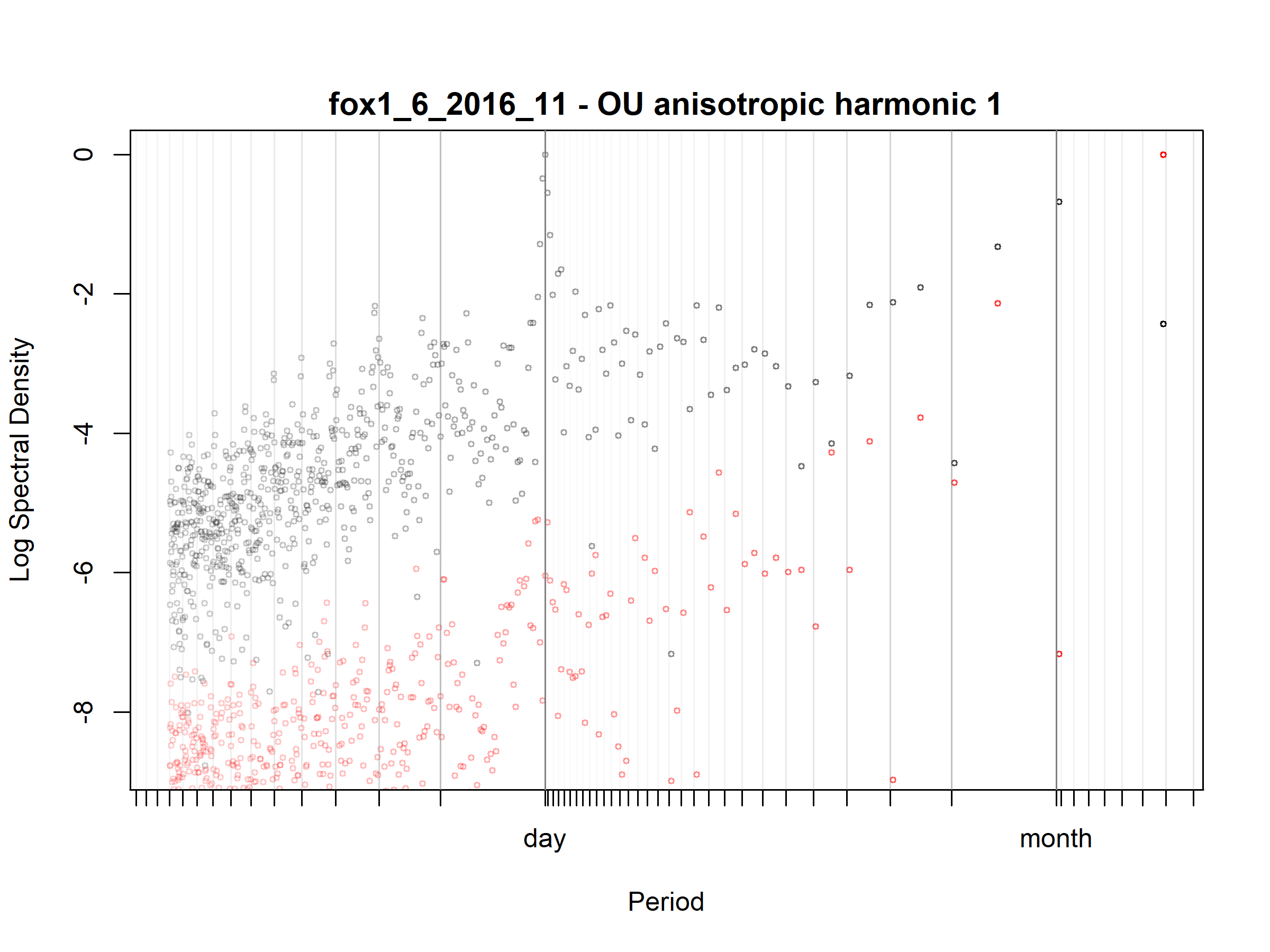

Figure S3.71. Fox; fox1_6_2016_11; diagnostic periodogram.

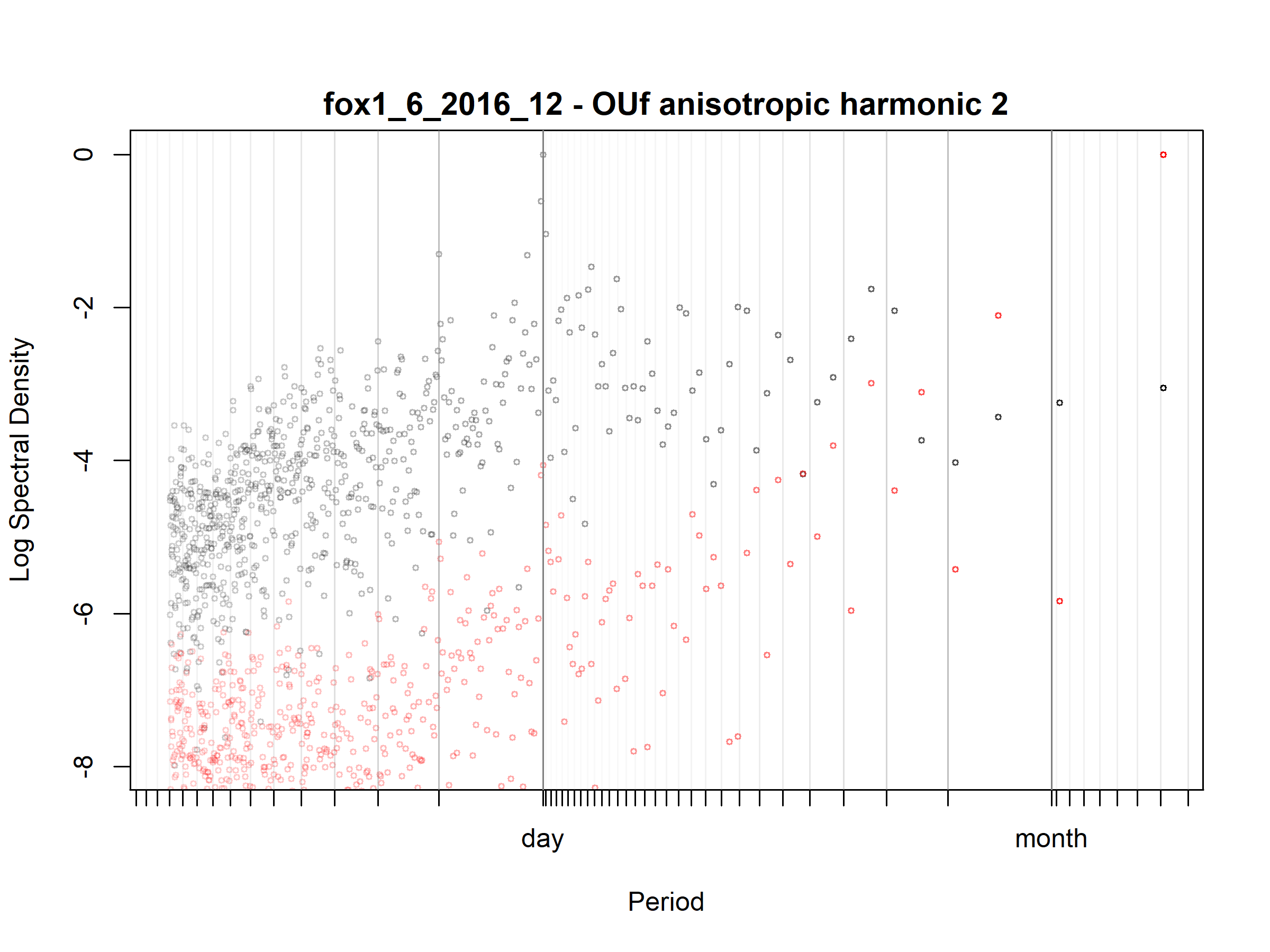

Figure S3.72. Fox; fox1_6_2016_12; diagnostic periodogram.

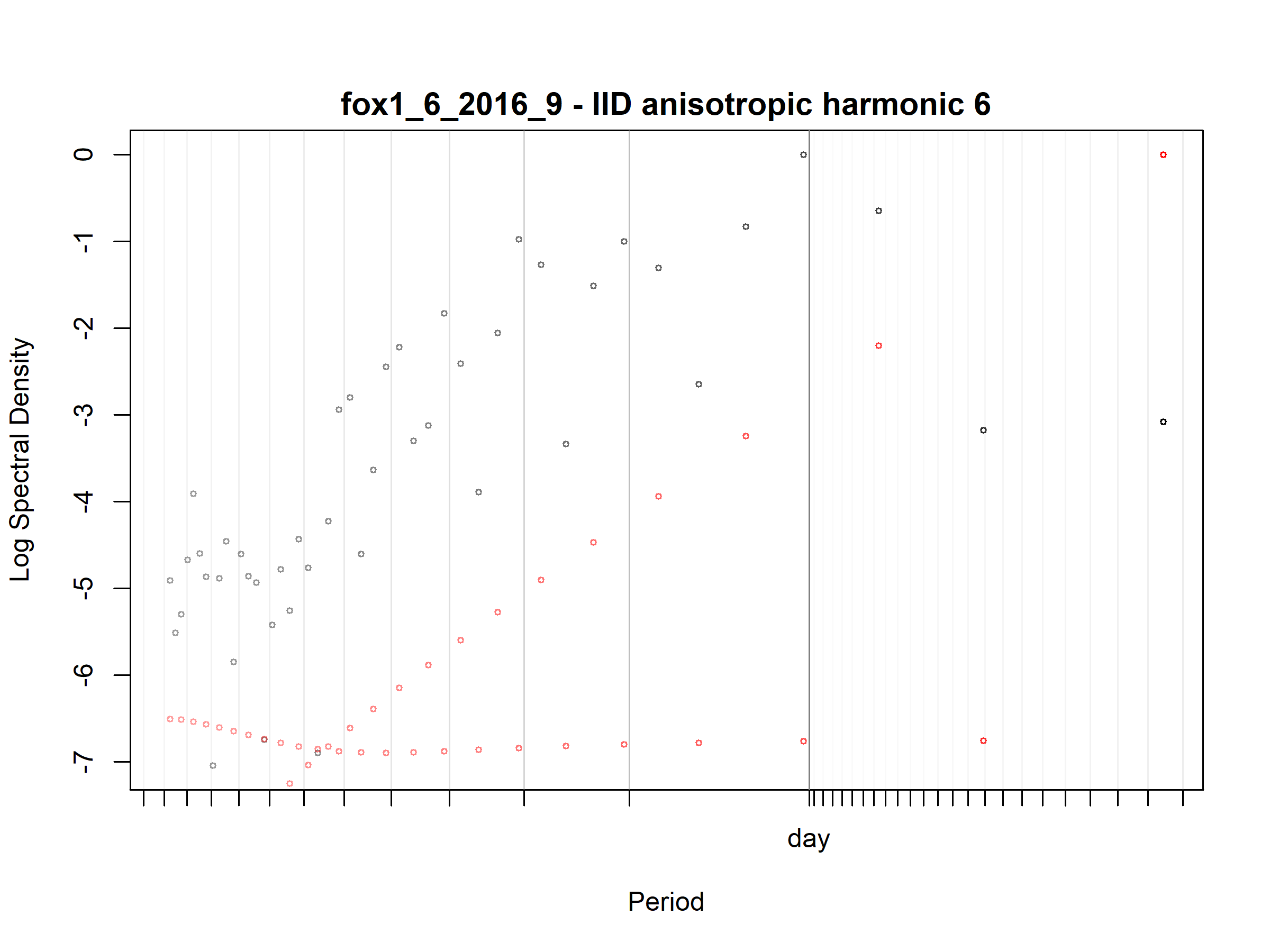

Figure S3.73. Fox; fox1_6_2016_9; diagnostic periodogram.

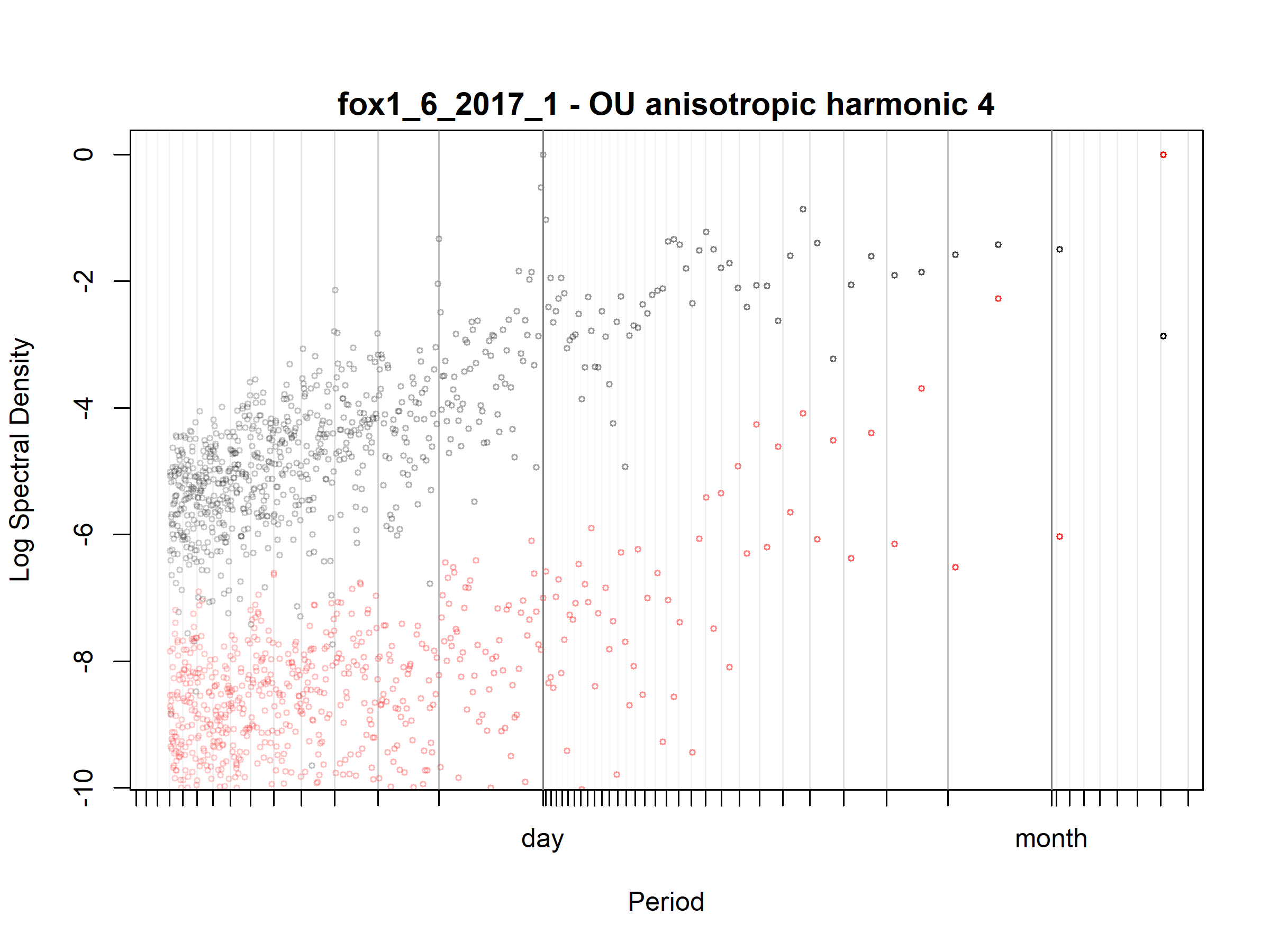

Figure S3.74. Fox; fox1_6_2017_1; diagnostic periodogram.

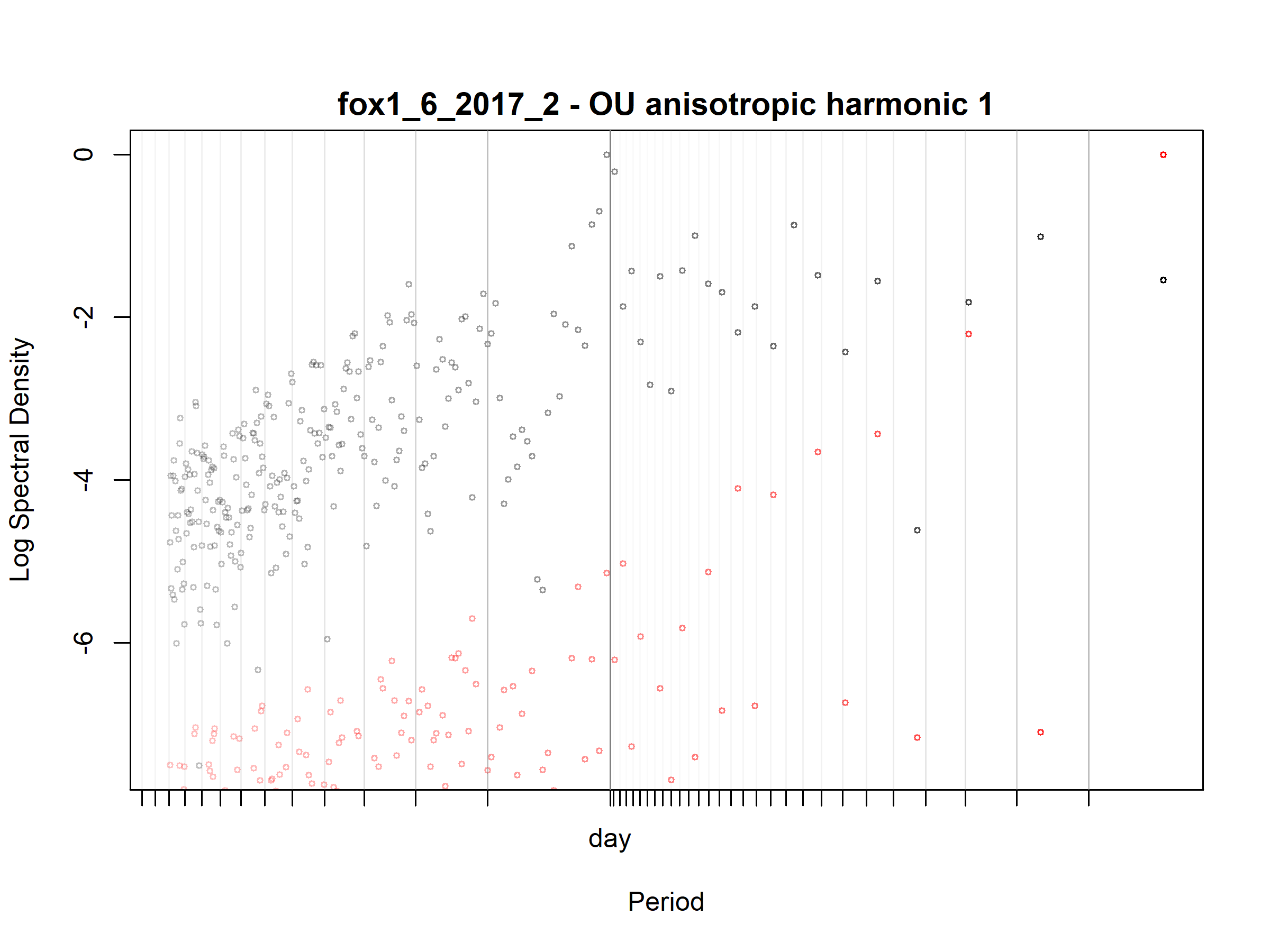

Figure S3.75. Fox; fox1_6_2017_2; diagnostic periodogram.

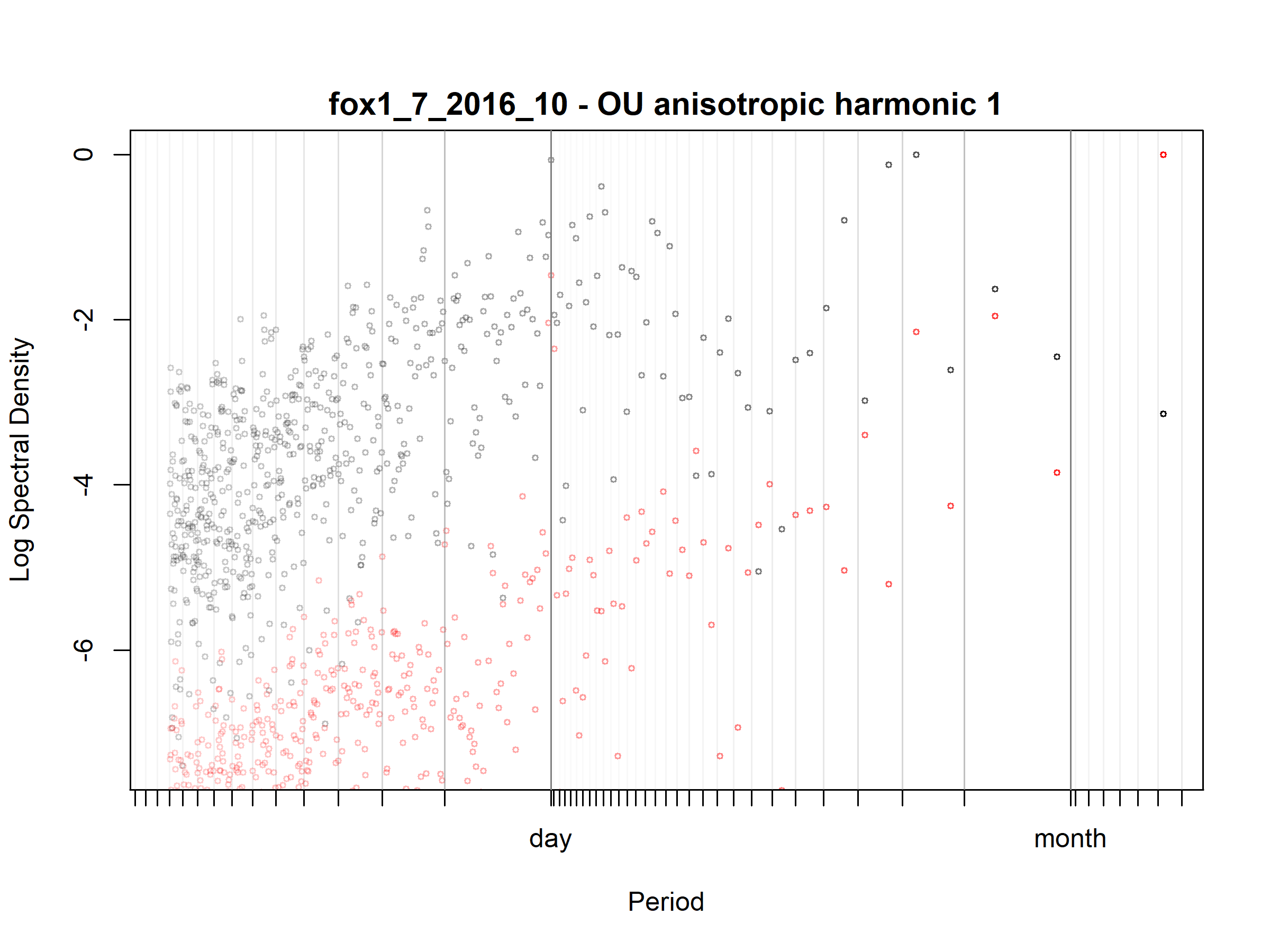

Figure S3.76. Fox; fox1_7_2016_10; diagnostic periodogram.

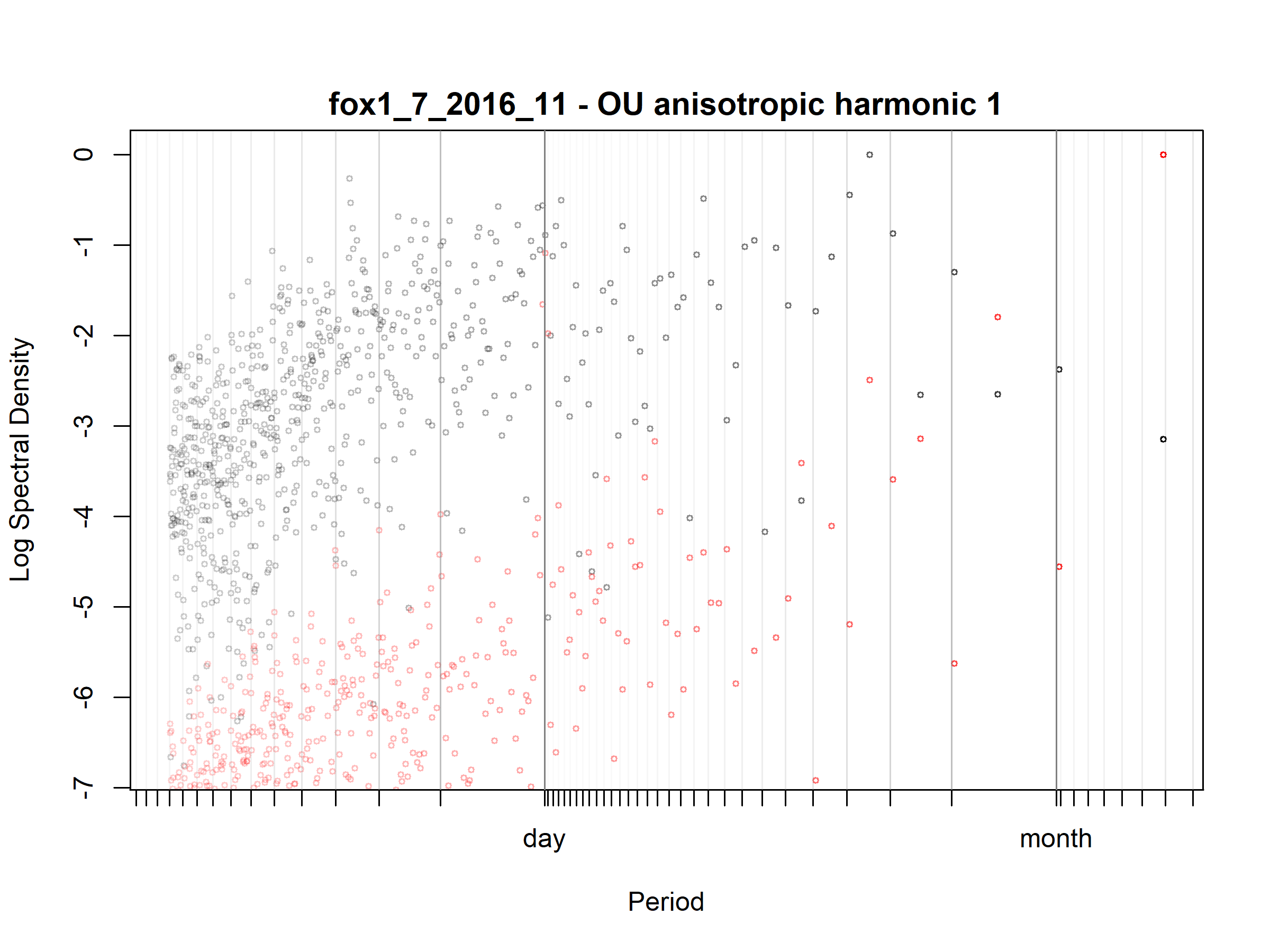

Figure S3.77. Fox; fox1_7_2016_11; diagnostic periodogram.

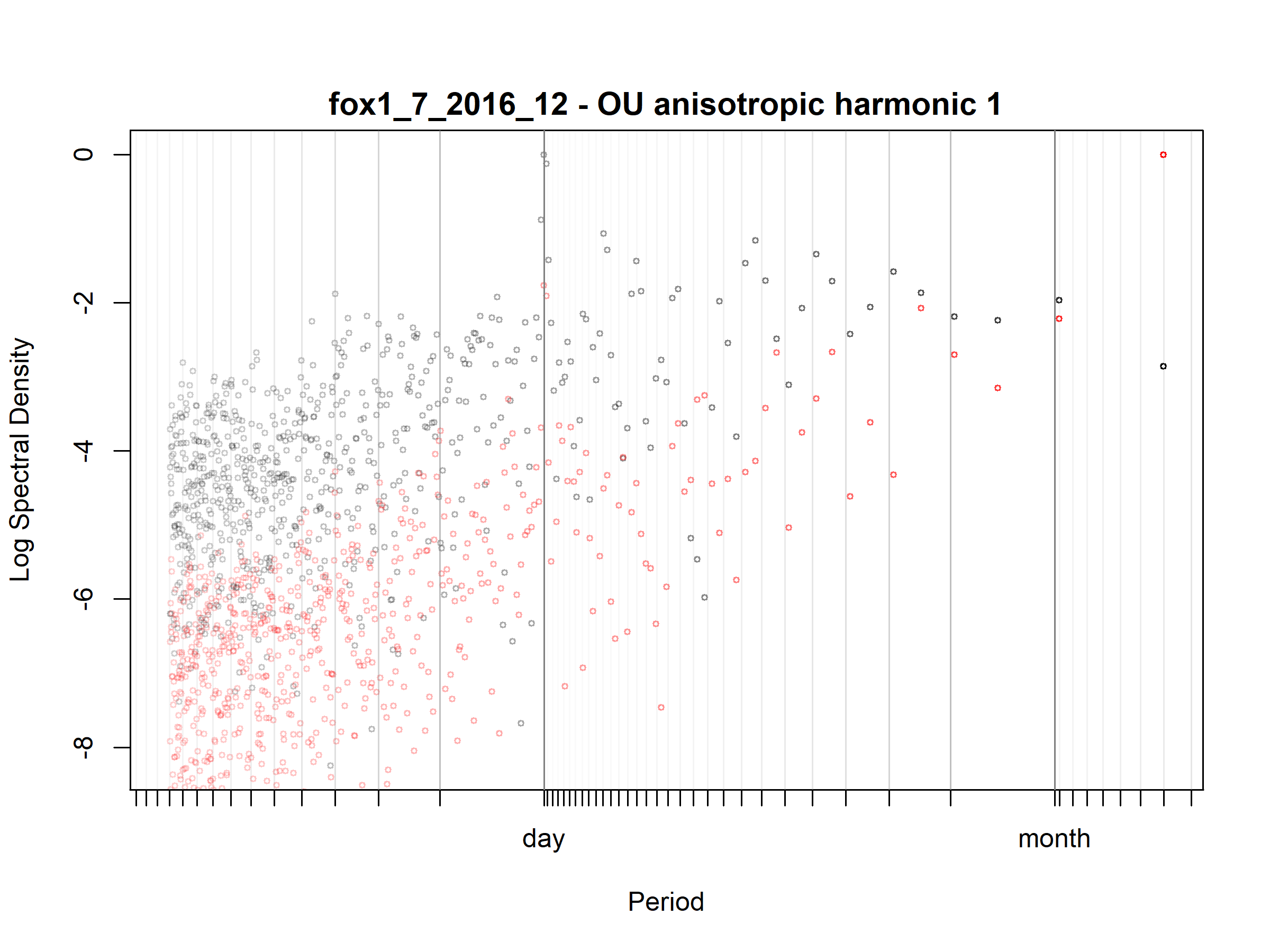

Figure S3.78. Fox; fox1_7_2016_12; diagnostic periodogram.

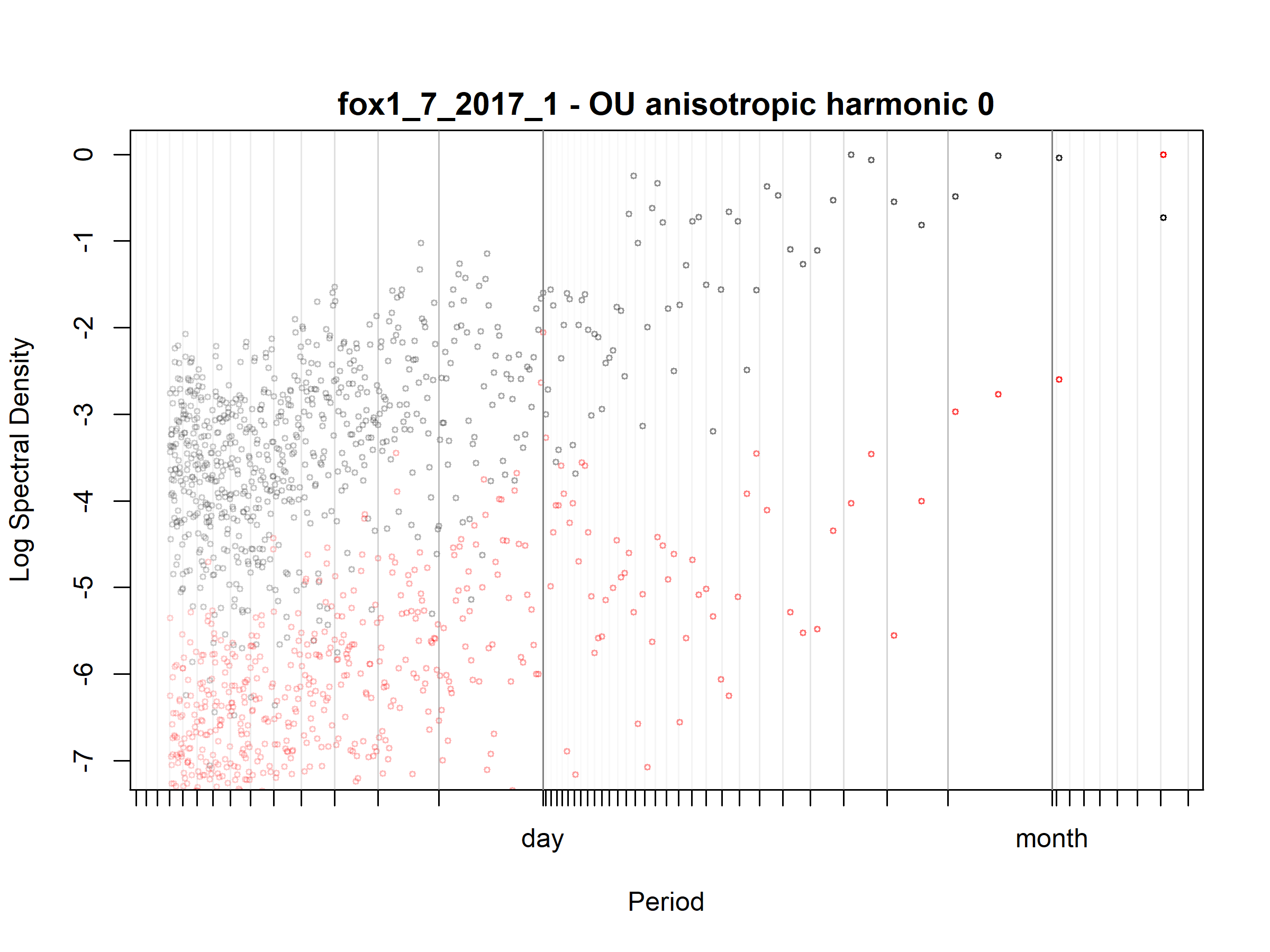

Figure S3.79. Fox; fox1_7_2017_1; diagnostic periodogram.

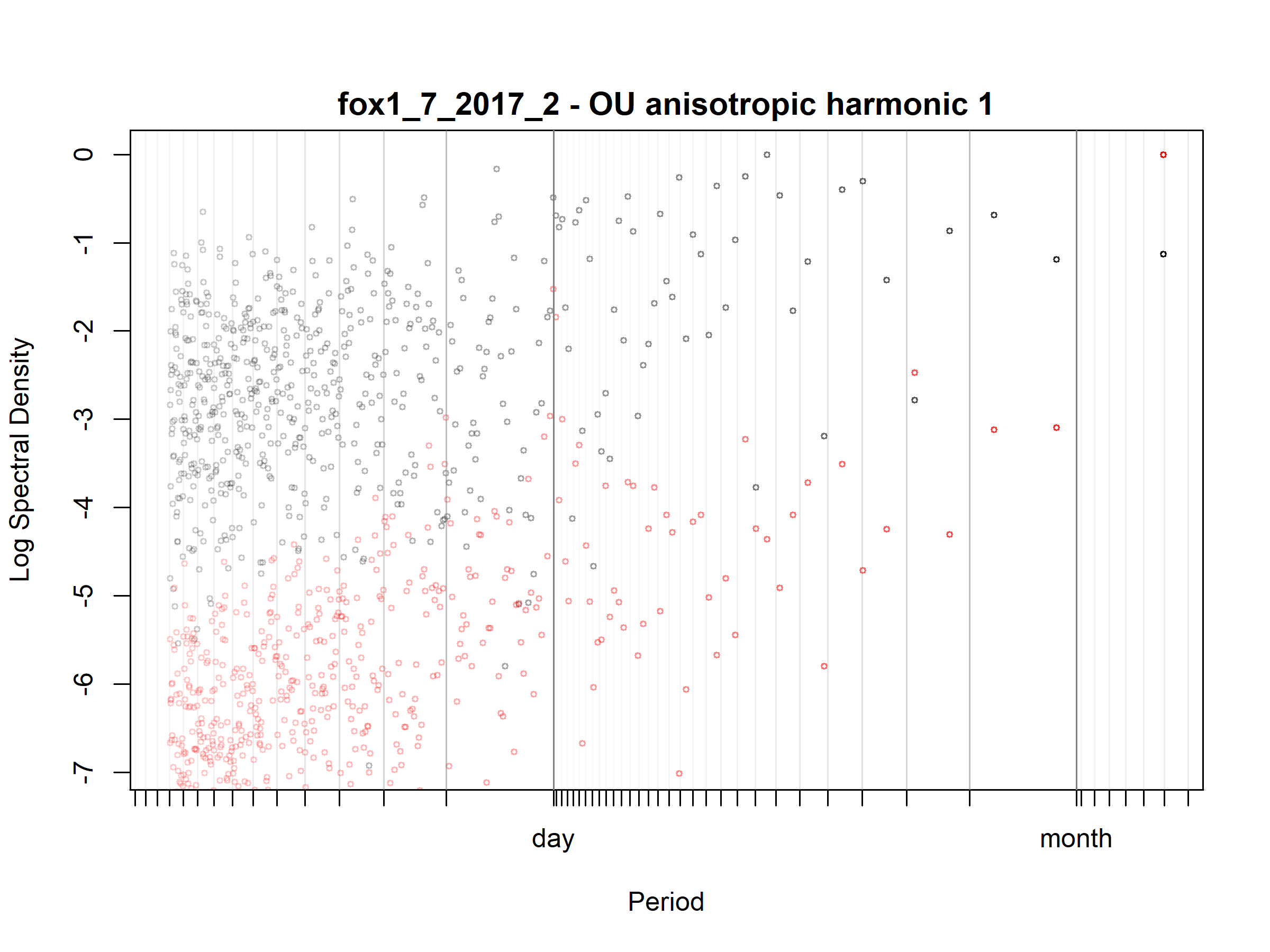

Figure S3.80. Fox; fox1_7_2017_2; diagnostic periodogram.

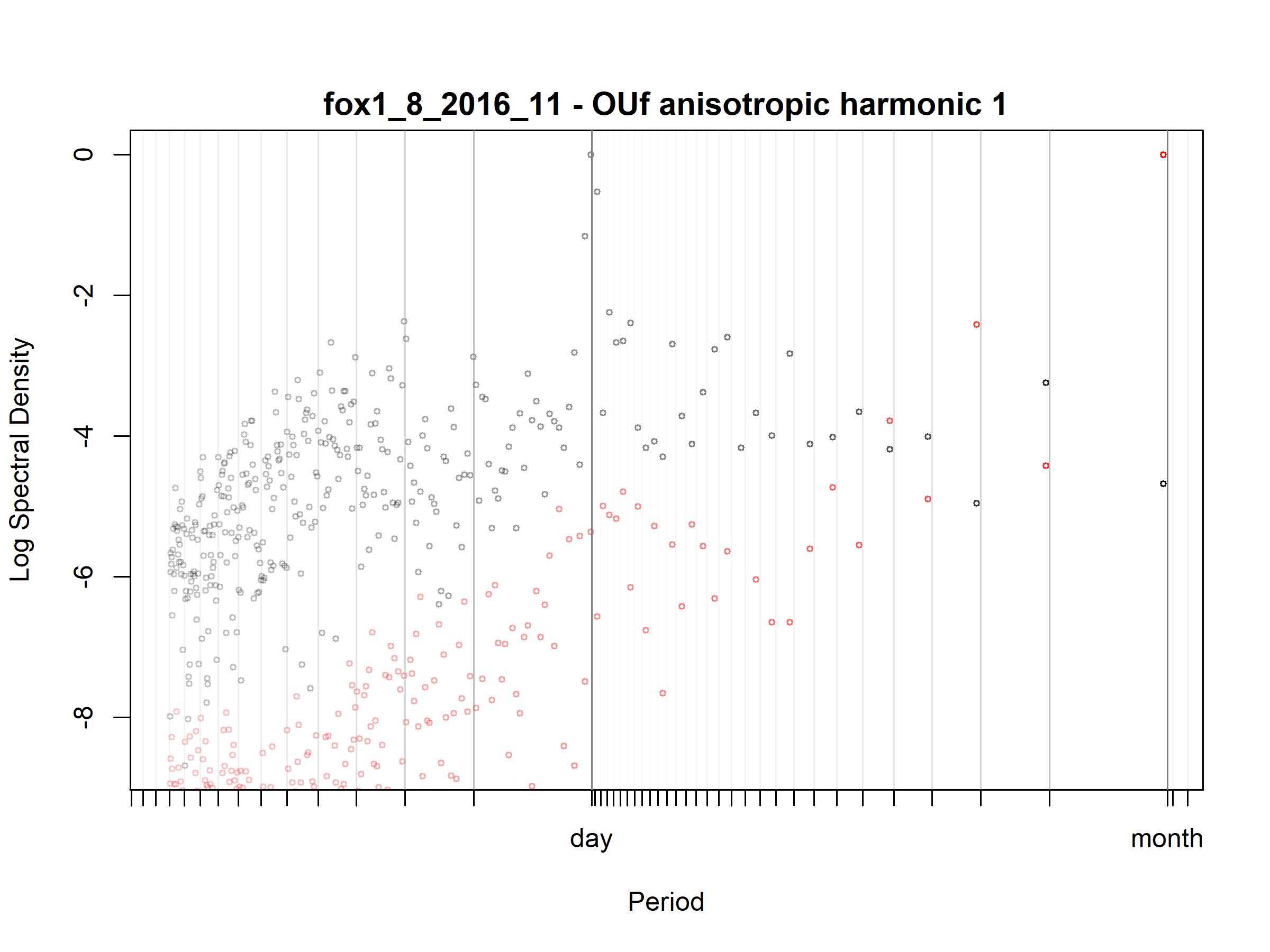

Figure S3.81. Fox; fox1_8_2016_11; diagnostic periodogram.

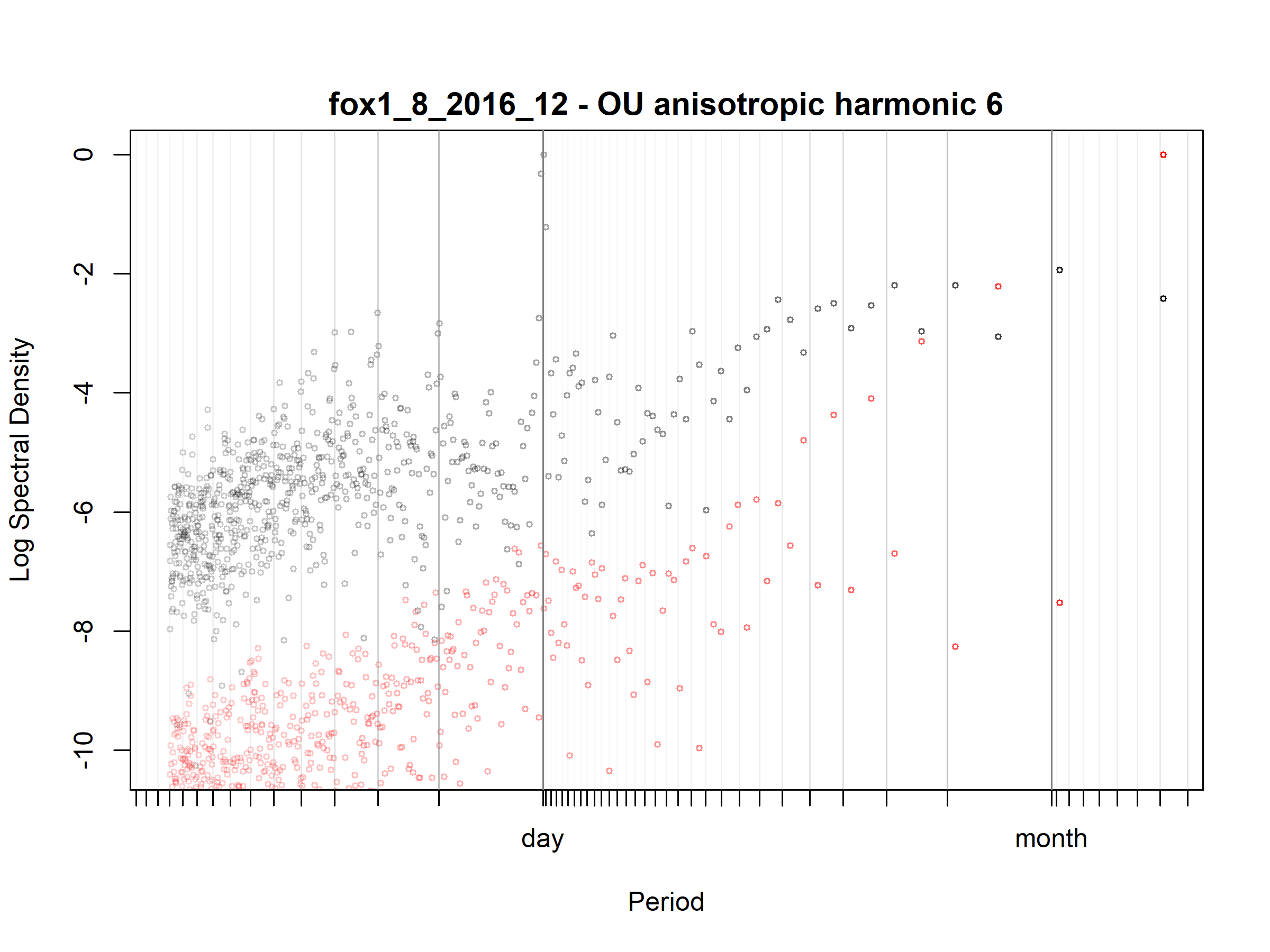

Figure S3.82. Fox; fox1_8_2016_12; diagnostic periodogram.

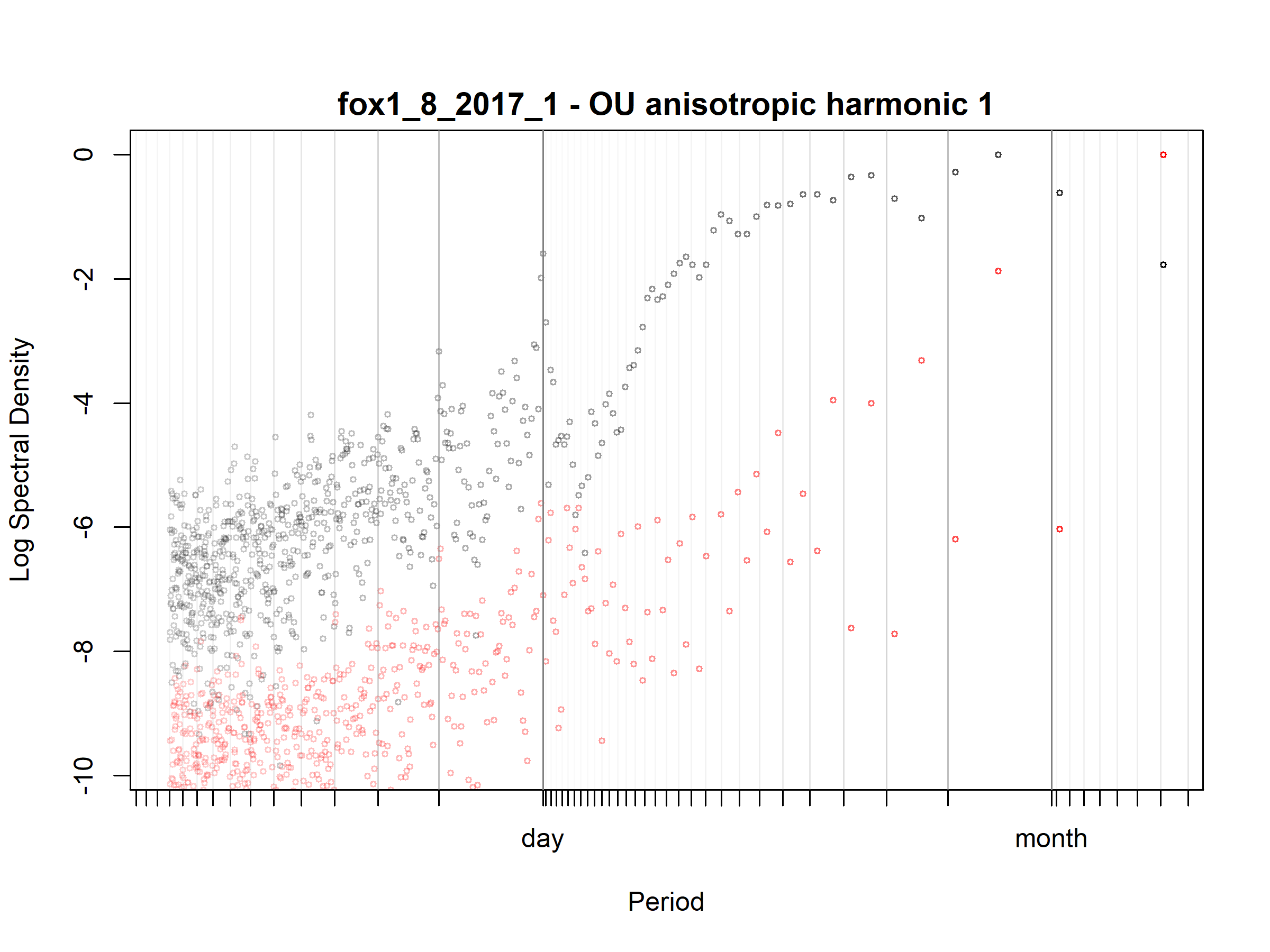

Figure S3.83. Fox; fox1_8_2017_1; diagnostic periodogram.

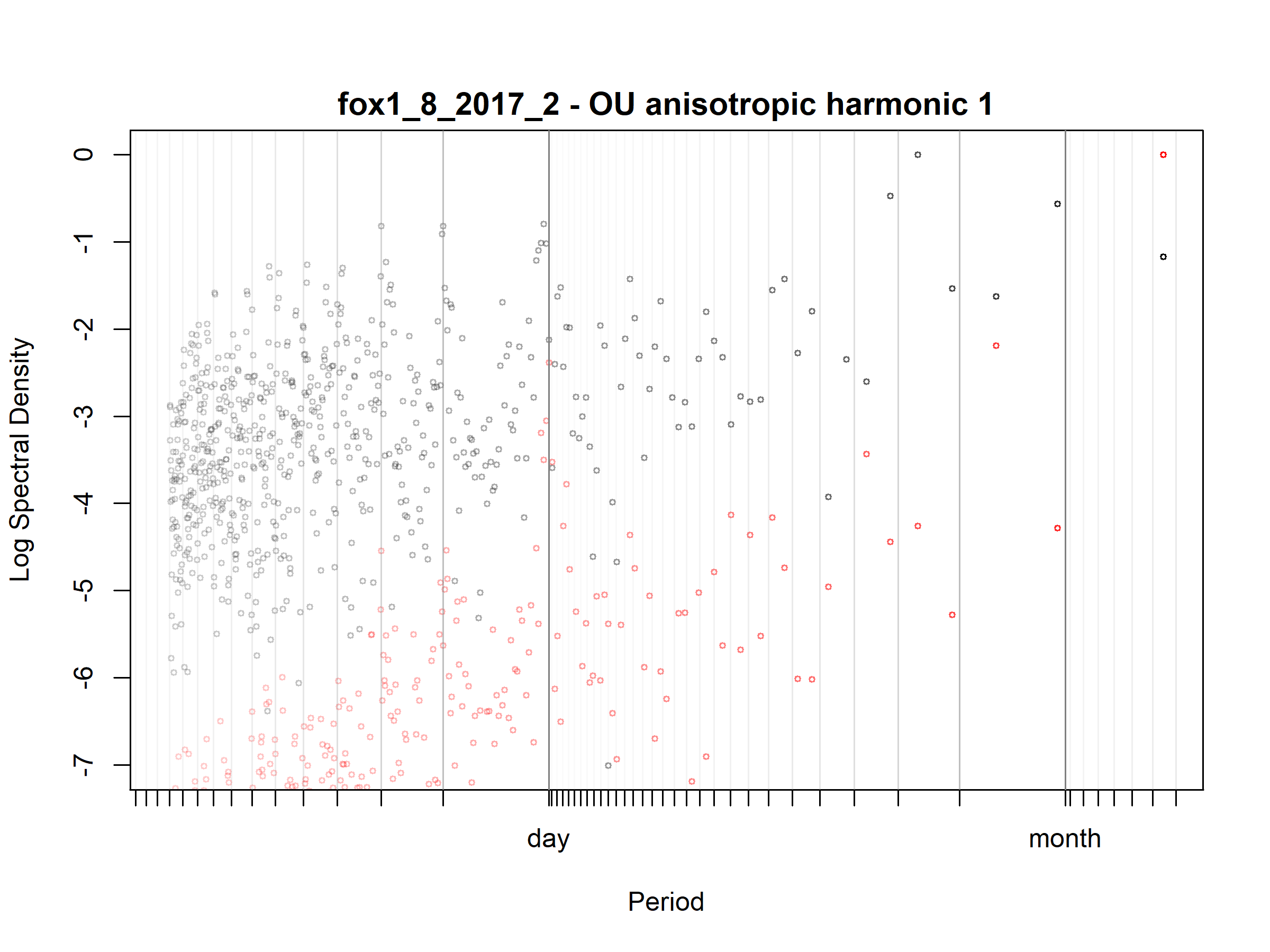

Figure S3.84. Fox; fox1_8_2017_2; diagnostic periodogram.

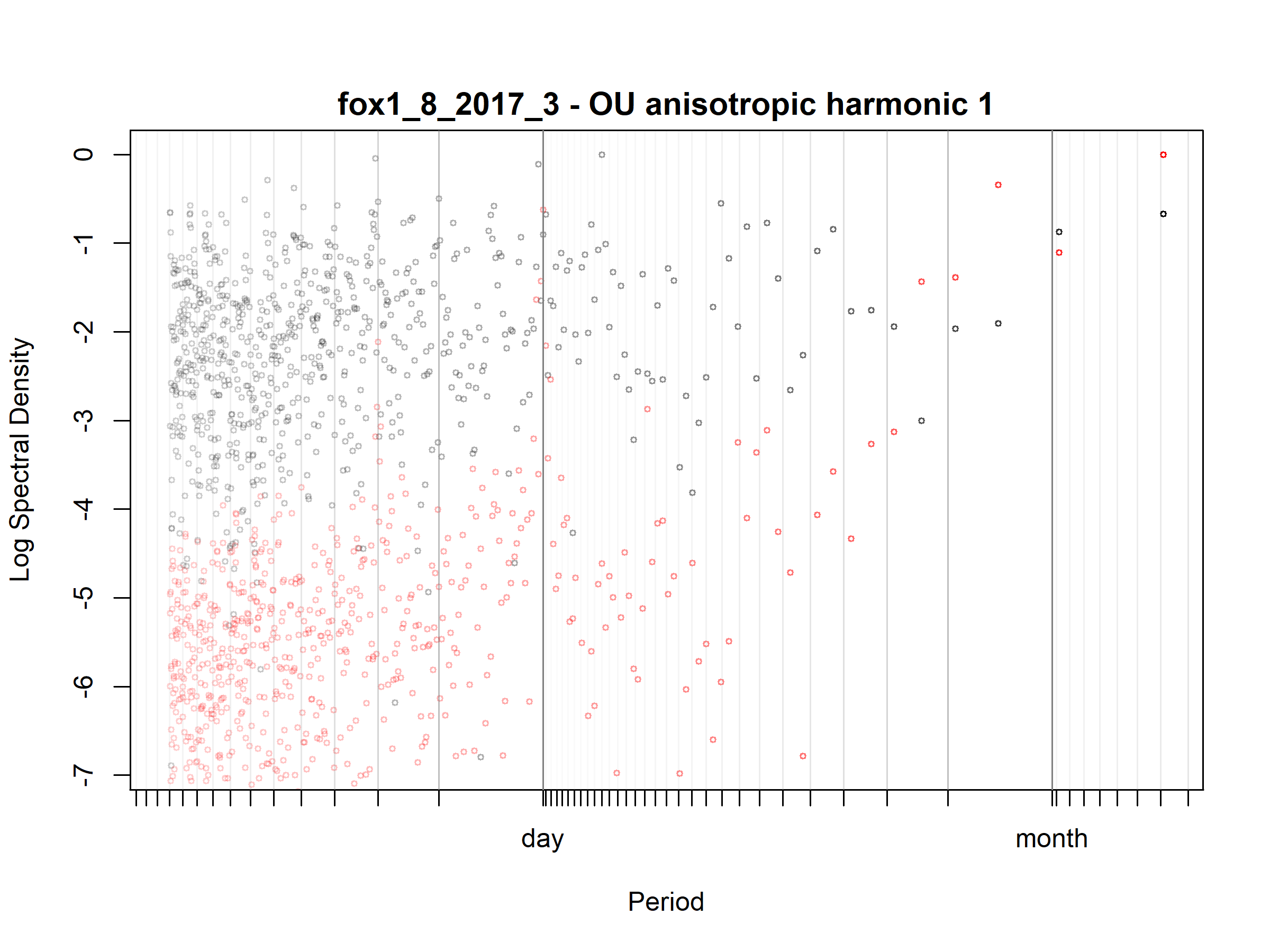

Figure S3.85. Fox; fox1_8_2017_3; diagnostic periodogram.

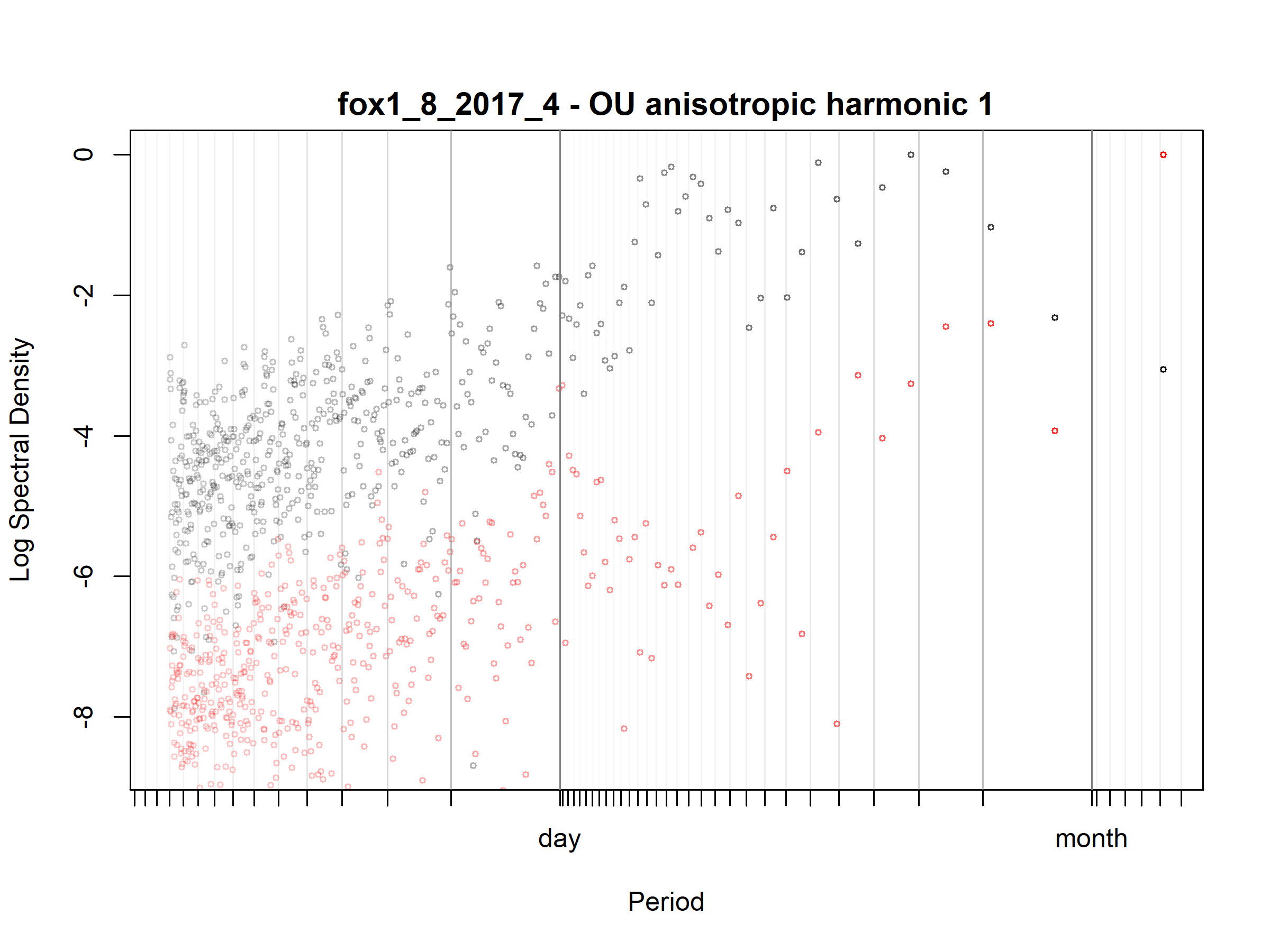

Figure S3.86. Fox; fox1_8_2017_4; diagnostic periodogram.

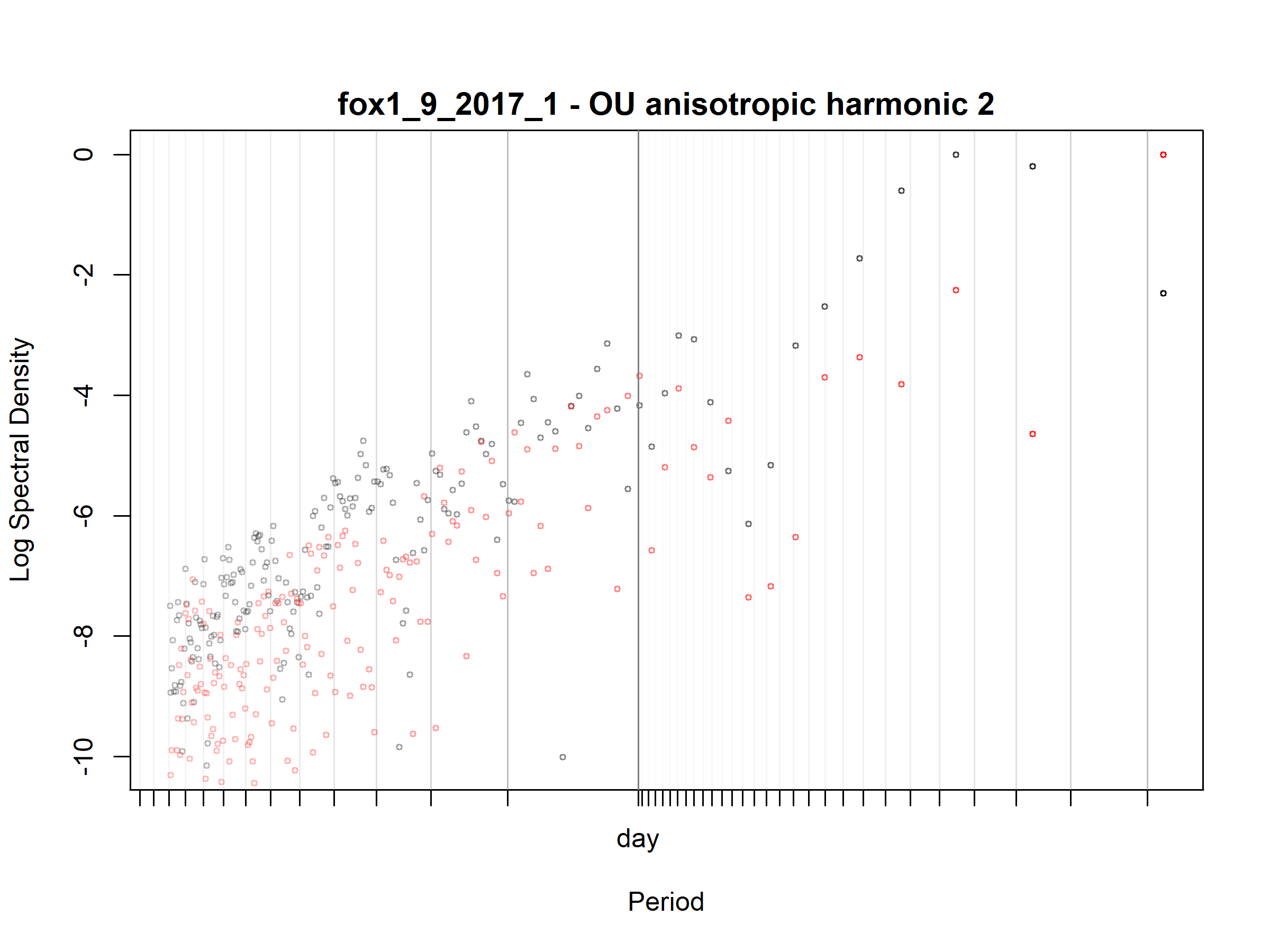

Figure S3.87. Fox; fox1_9_2017_1; diagnostic periodogram.

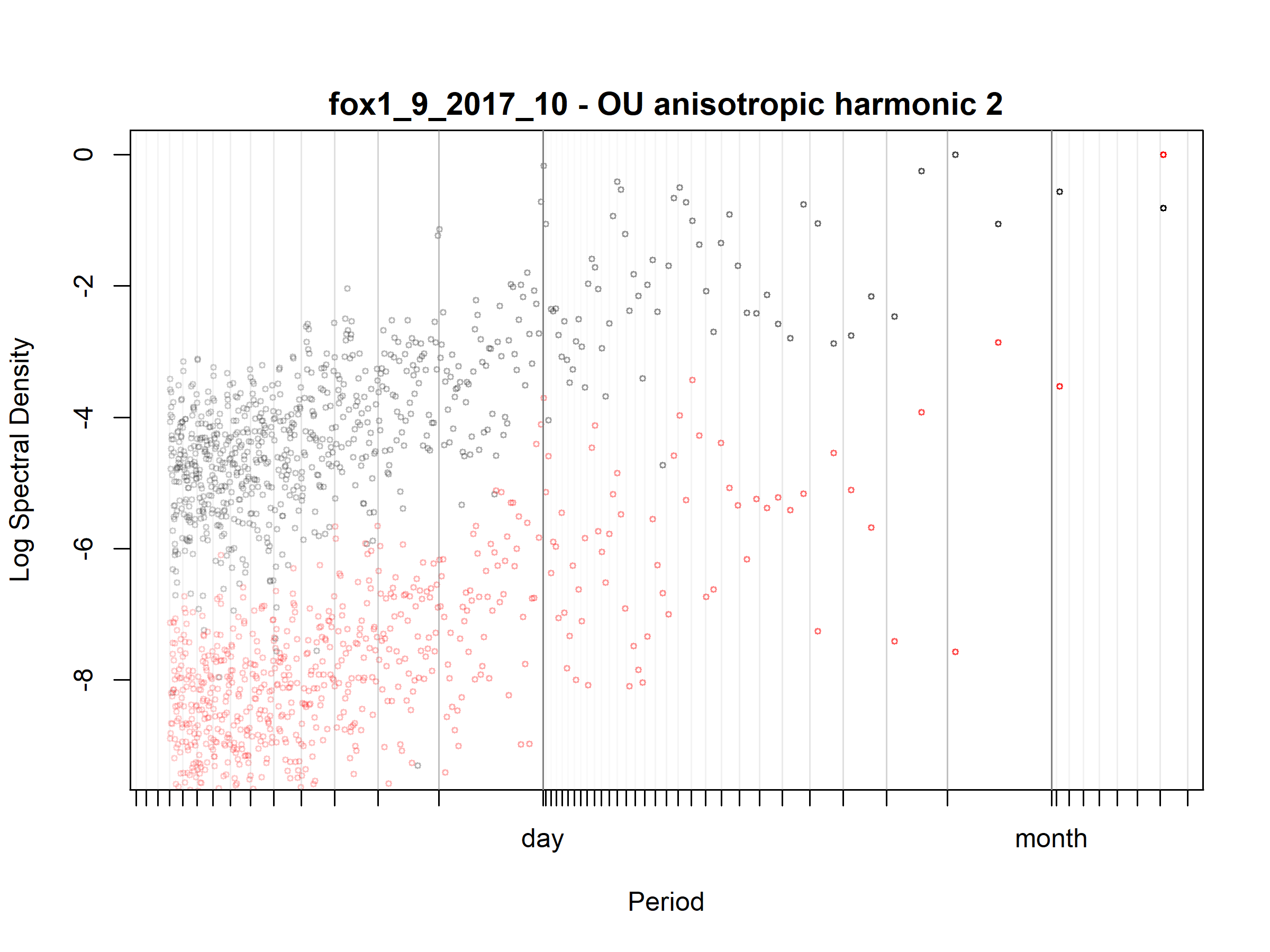

Figure S3.88. Fox; fox1_9_2017_10; diagnostic periodogram.

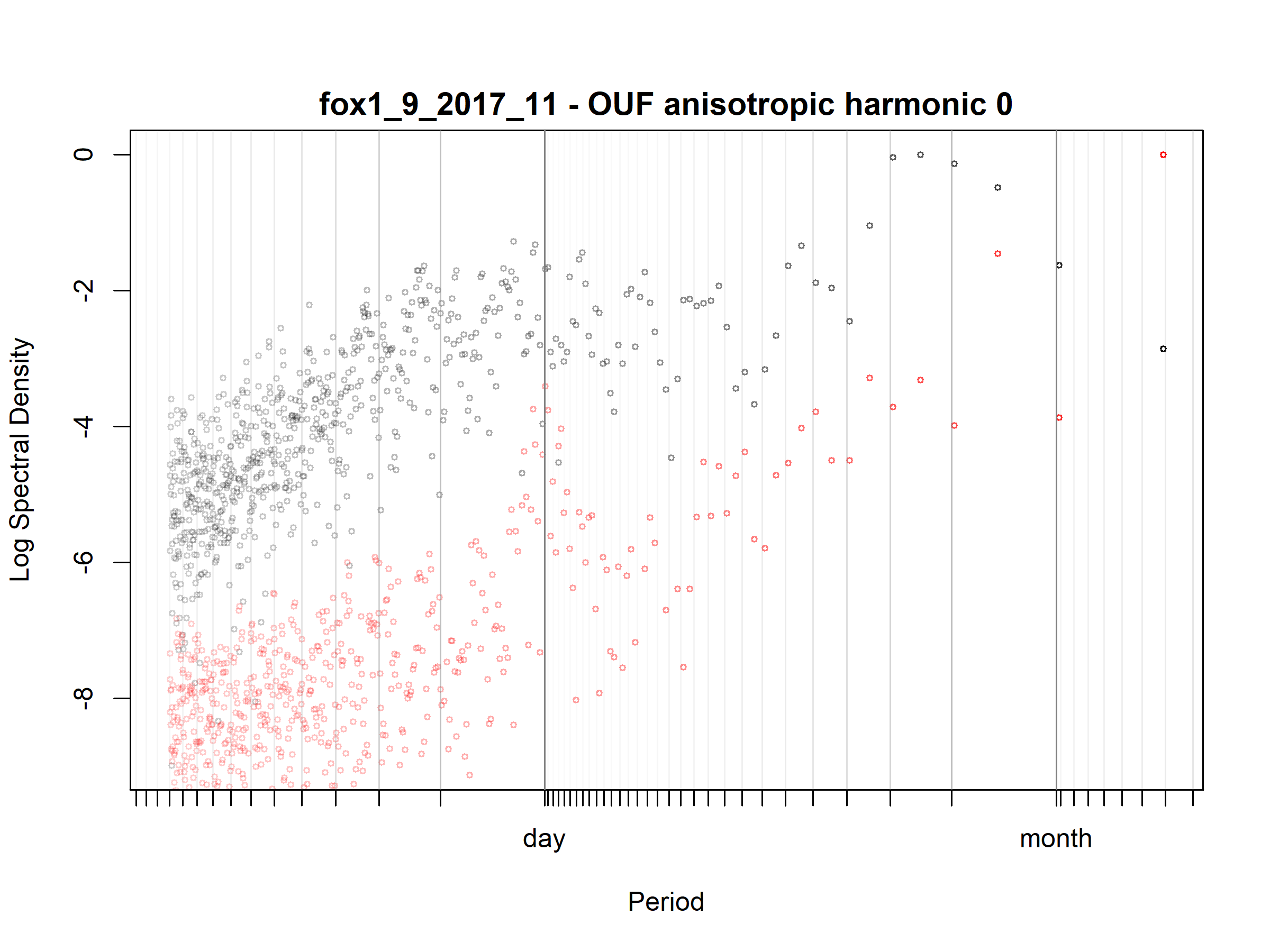

Figure S3.89. Fox; fox1_9_2017_11; diagnostic periodogram.

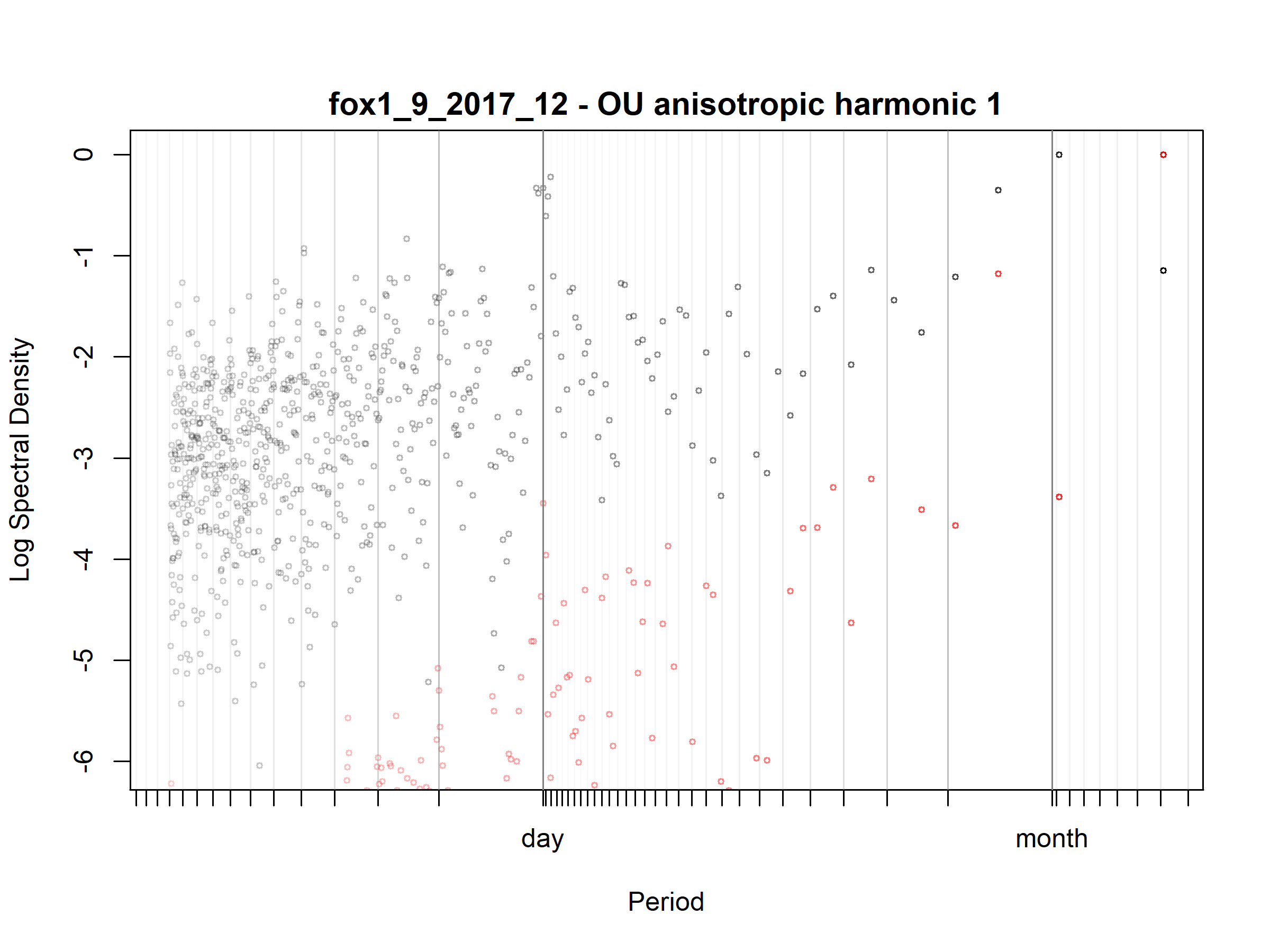

Figure S3.90. Fox; fox1_9_2017_12; diagnostic periodogram.

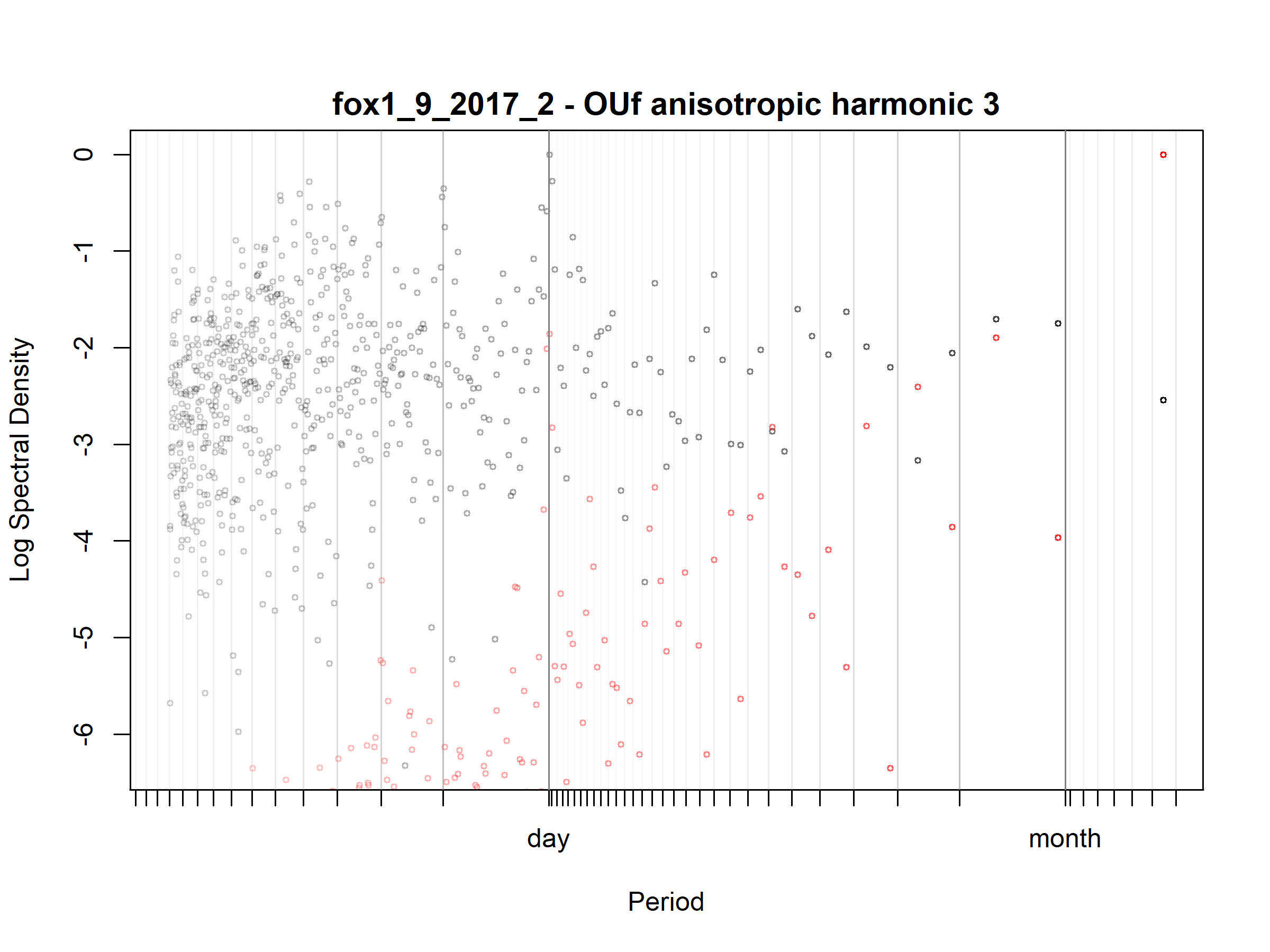

Figure S3.91. Fox; fox1_9_2017_2; diagnostic periodogram.

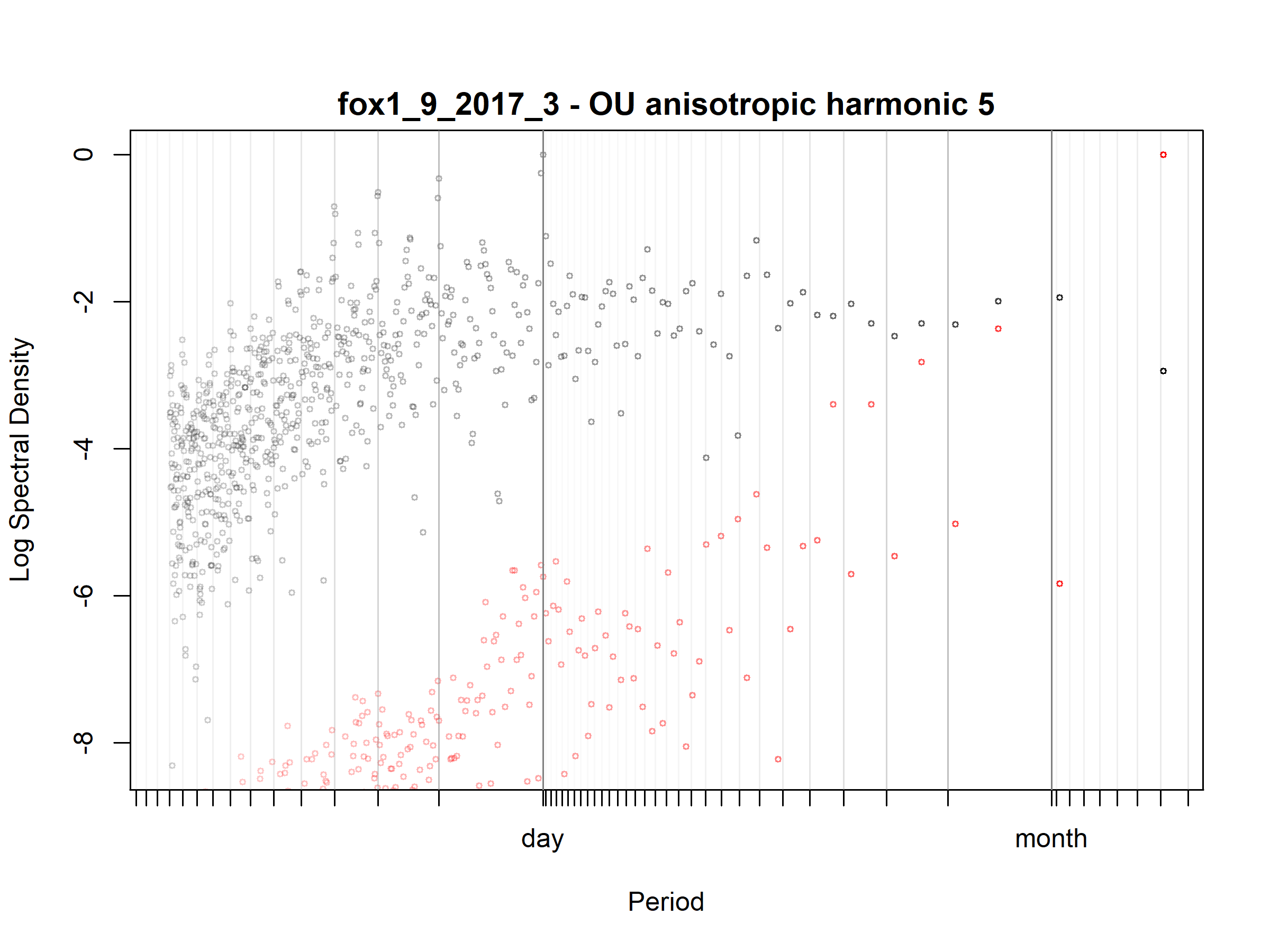

Figure S3.92. Fox; fox1_9_2017_3; diagnostic periodogram.

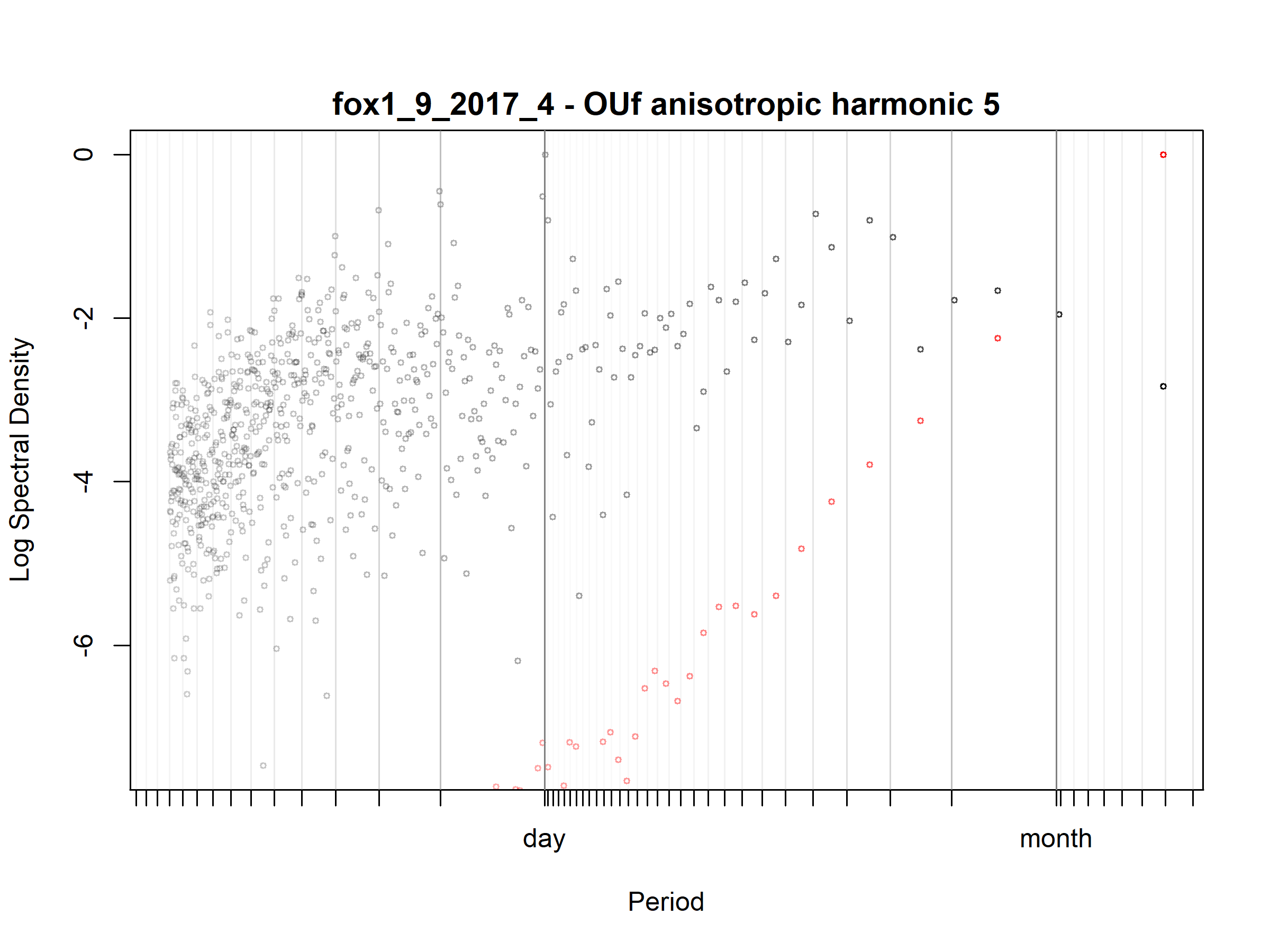

Figure S3.93. Fox; fox1_9_2017_4; diagnostic periodogram.

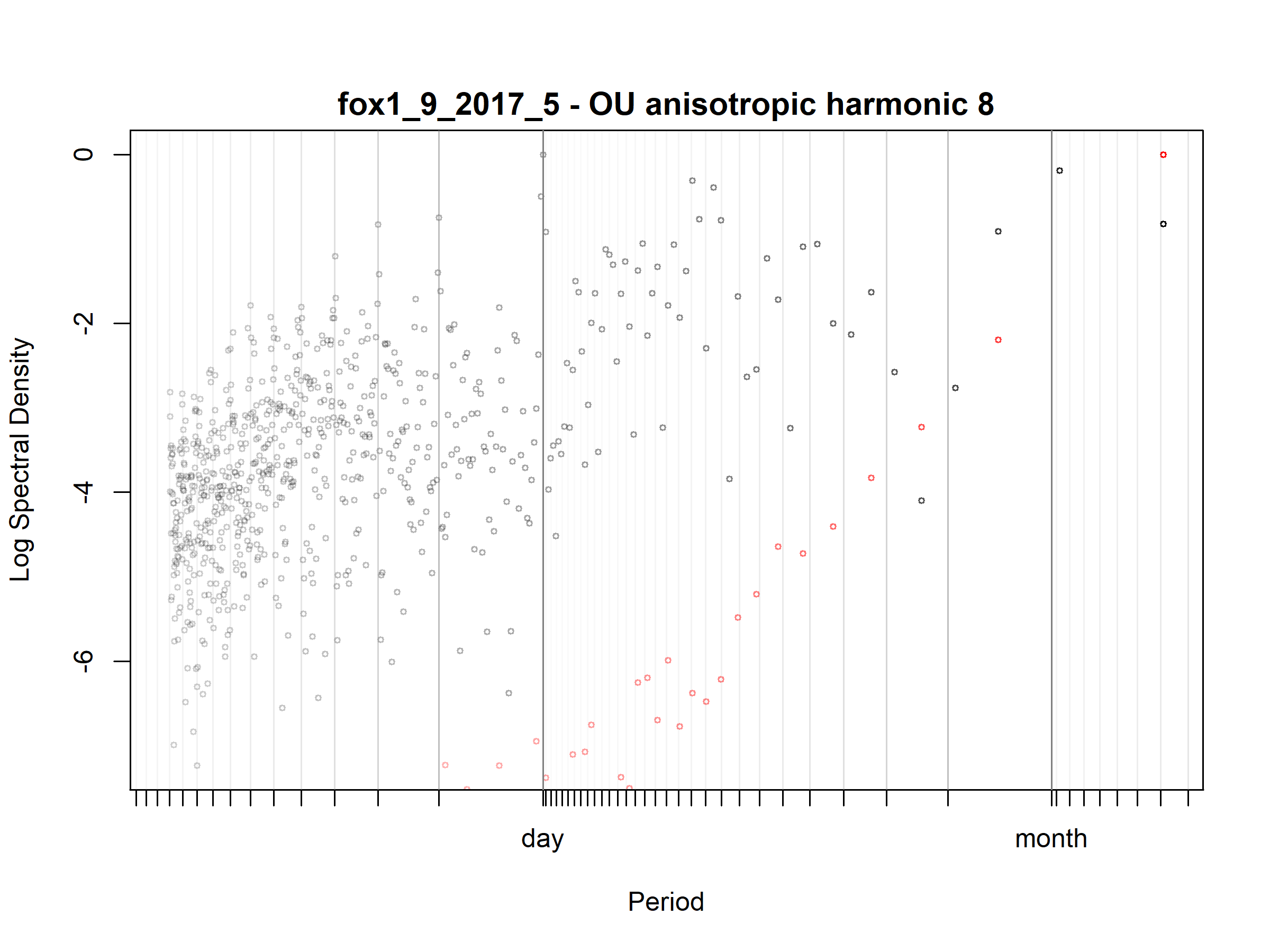

Figure S3.94. Fox; fox1_9_2017_5; diagnostic periodogram.

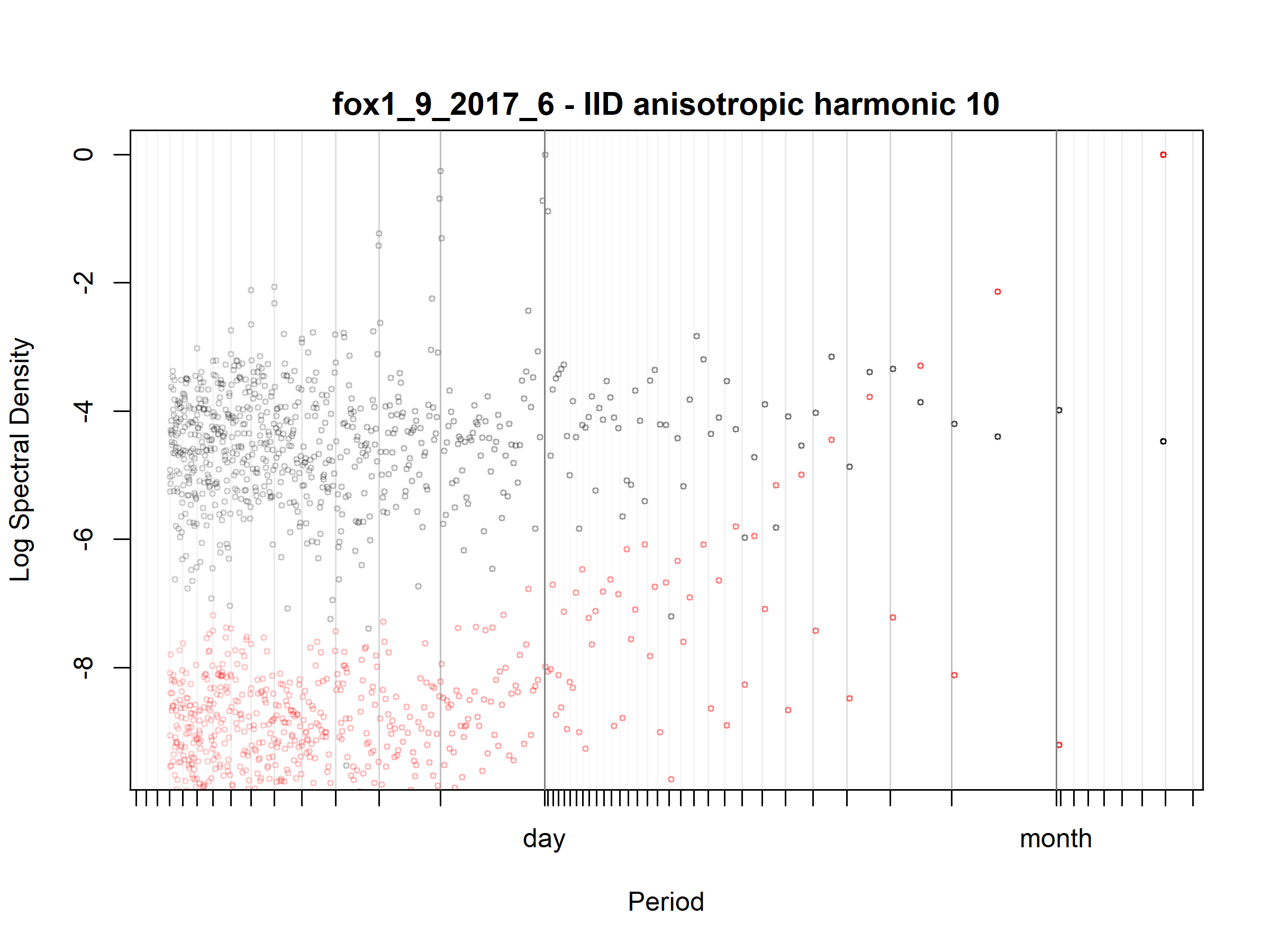

Figure S3.95. Fox; fox1_9_2017_6; diagnostic periodogram.

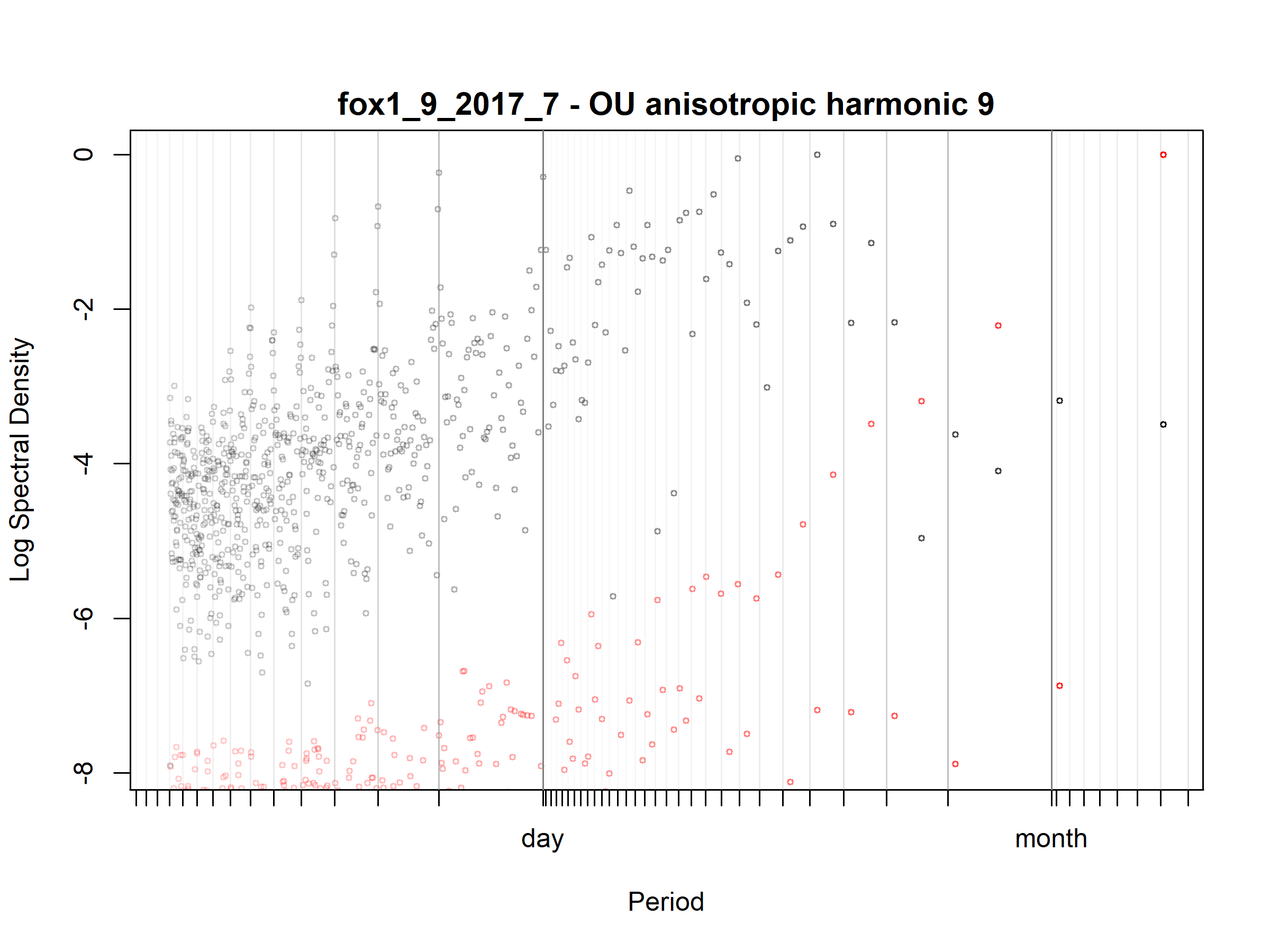

Figure S3.96. Fox; fox1_9_2017_7; diagnostic periodogram.

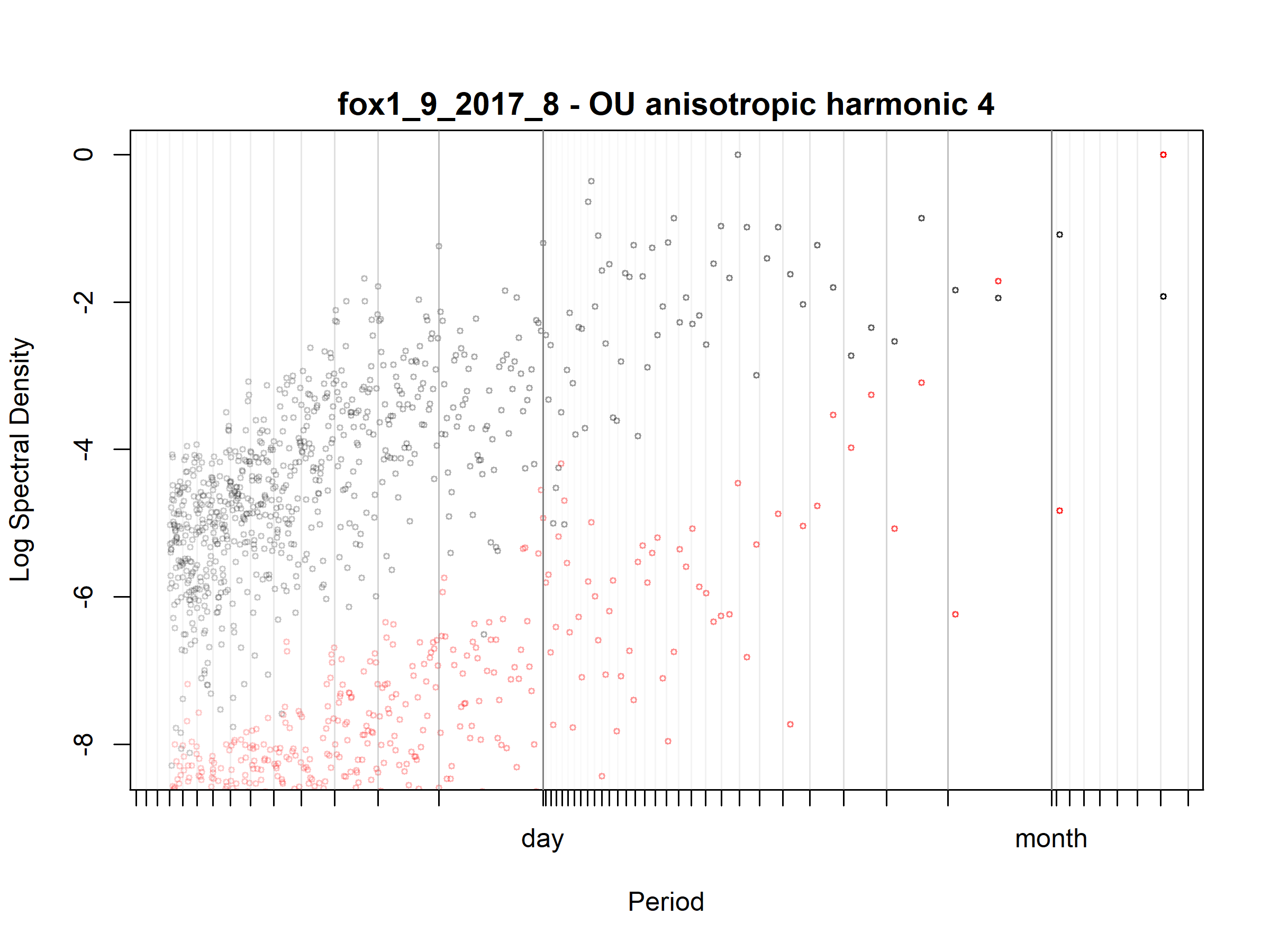

Figure S3.97. Fox; fox1_9_2017_8; diagnostic periodogram.

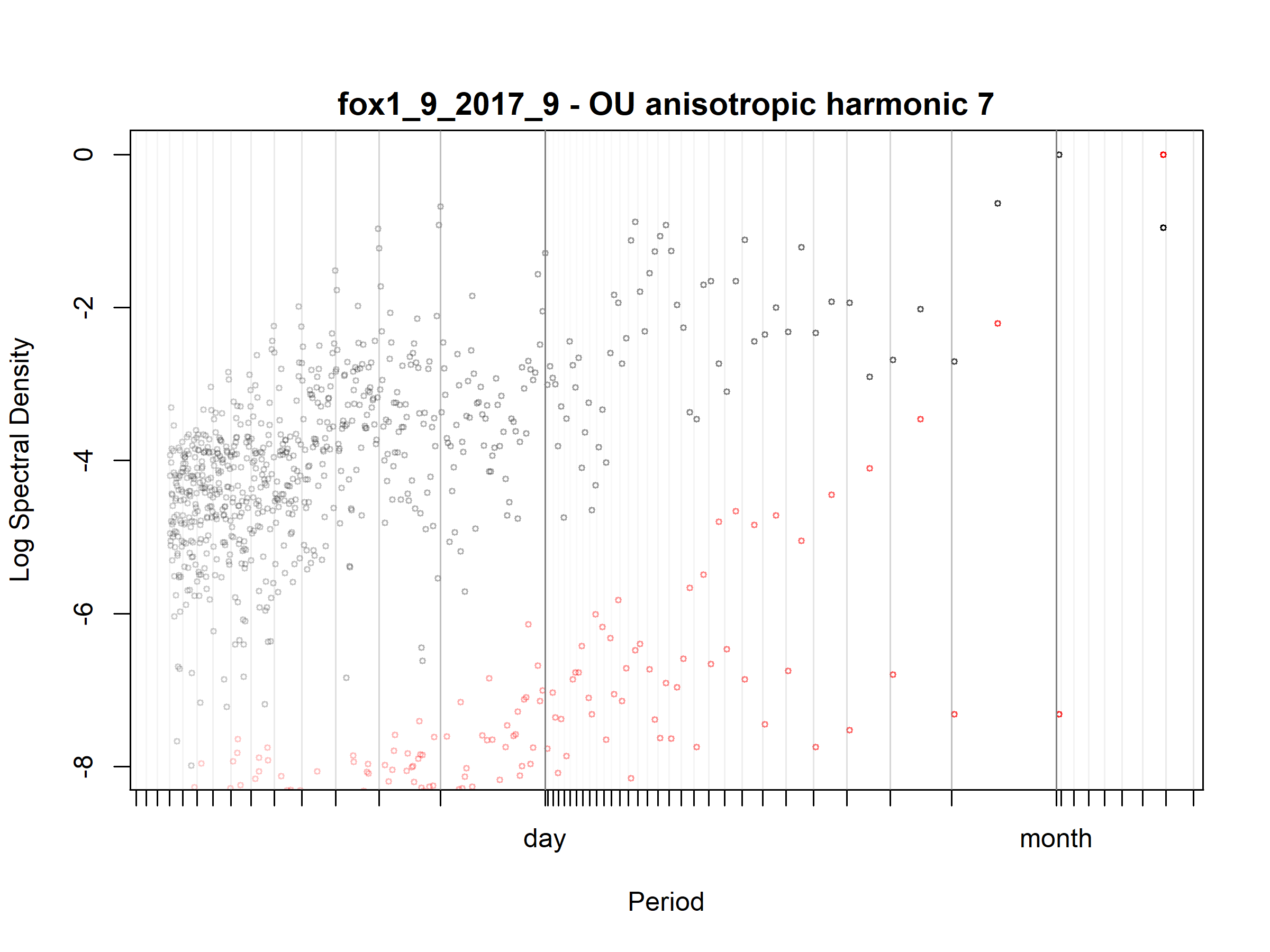

Figure S3.98. Fox; fox1_9_2017_9; diagnostic periodogram.

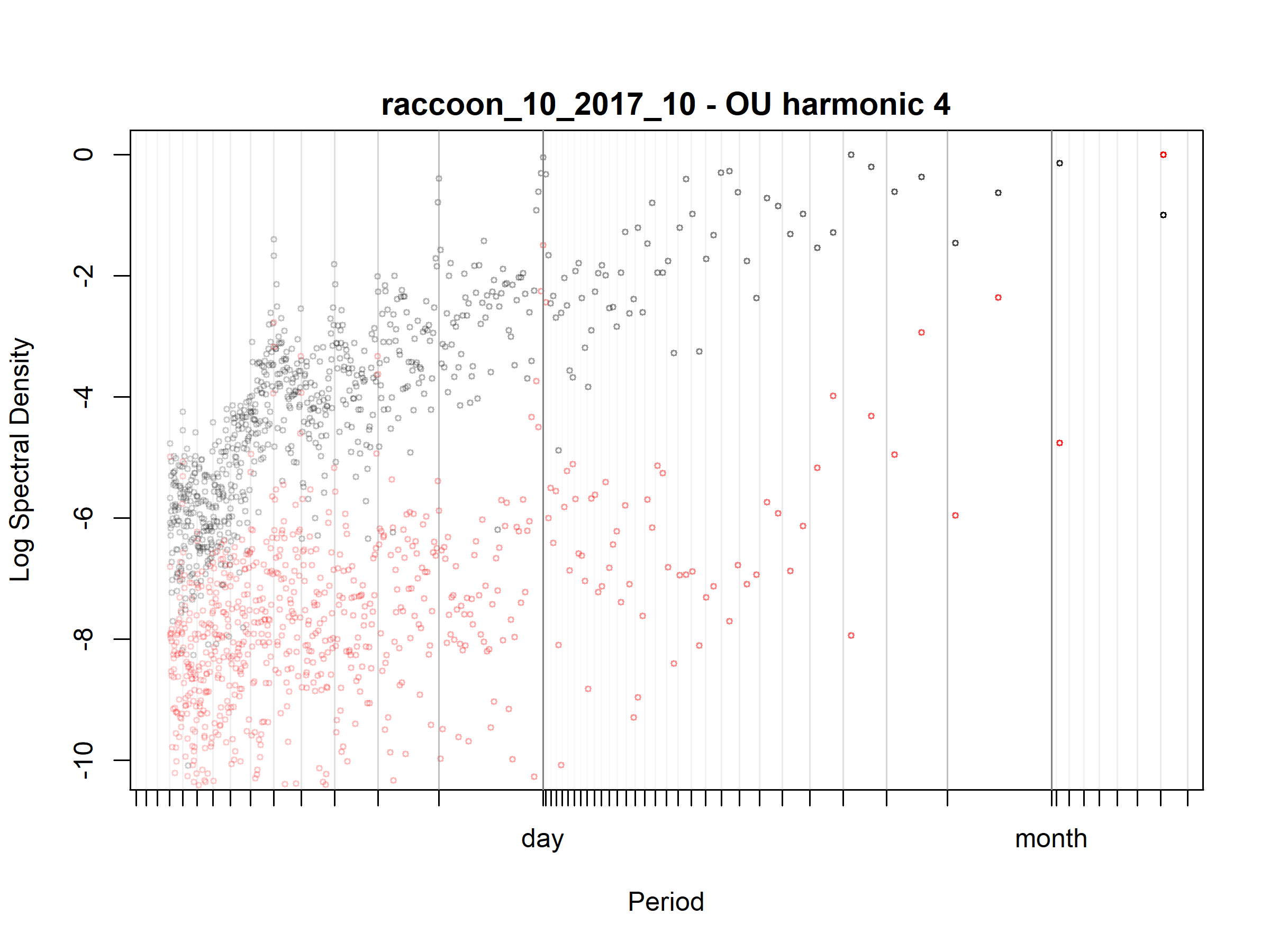

Figure S3.99. Raccoon; raccoon_10_2017_10; diagnostic periodogram.

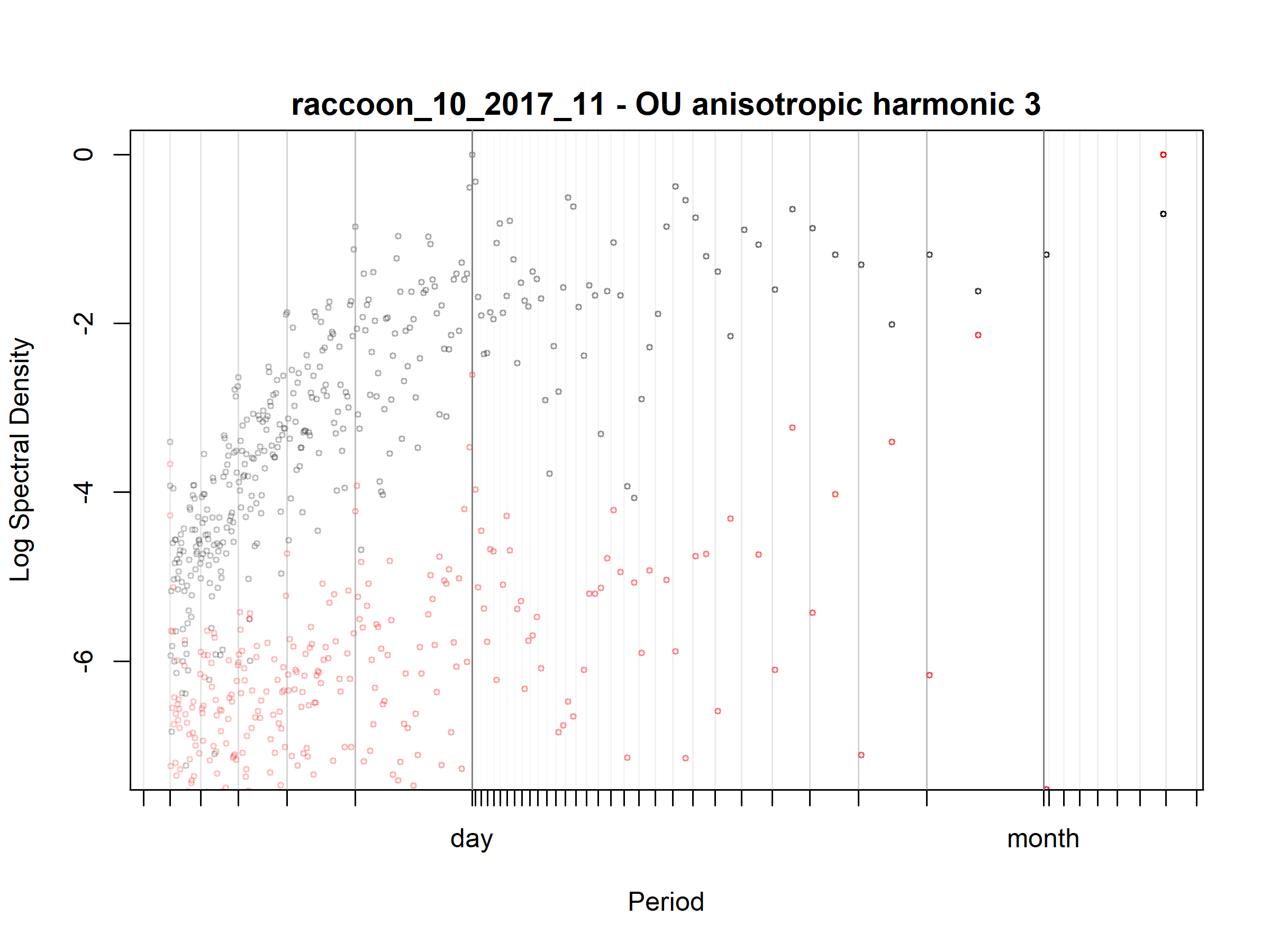

Figure S3.100. Raccoon; raccoon_10_2017_11; diagnostic periodogram.

Figure S3.101. Raccoon; raccoon_10_2017_12; diagnostic periodogram.

Figure S3.102. Raccoon; raccoon_10_2017_7; diagnostic periodogram.

Figure S3.103. Raccoon; raccoon_10_2017_8; diagnostic periodogram.

Figure S3.104. Raccoon; raccoon_10_2017_9; diagnostic periodogram.

Figure S3.105. Raccoon; raccoon_10_2018_1; diagnostic periodogram.

Figure S3.106. Raccoon; raccoon_10_2018_2; diagnostic periodogram.

Figure S3.107. Raccoon; raccoon_11_2017_10; diagnostic periodogram.

Figure S3.108. Raccoon; raccoon_11_2017_11; diagnostic periodogram.

Figure S3.109. Raccoon; raccoon_11_2017_12; diagnostic periodogram.

Figure S3.110. Raccoon; raccoon_11_2017_7; diagnostic periodogram.

Figure S3.111. Raccoon; raccoon_11_2017_8; diagnostic periodogram.

Figure S3.112. Raccoon; raccoon_11_2017_9; diagnostic periodogram.

Figure S3.113. Raccoon; raccoon_11_2018_1; diagnostic periodogram.

Figure S3.114. Raccoon; raccoon_11_2018_2; diagnostic periodogram.

Figure S3.115. Raccoon; raccoon_11_2018_3; diagnostic periodogram.

Figure S3.116. Raccoon; raccoon_11_2018_4; diagnostic periodogram.

Figure S3.117. Raccoon; raccoon_12_2017_10; diagnostic periodogram.

Figure S3.118. Raccoon; raccoon_12_2017_11; diagnostic periodogram.

Figure S3.119. Raccoon; raccoon_12_2017_12; diagnostic periodogram.

Figure S3.120. Raccoon; raccoon_12_2017_8; diagnostic periodogram.

Figure S3.121. Raccoon; raccoon_12_2017_9; diagnostic periodogram.

Figure S3.122. Raccoon; raccoon_12_2018_1; diagnostic periodogram.

Figure S3.123. Raccoon; raccoon_12_2018_2; diagnostic periodogram.

Figure S3.124. Raccoon; raccoon_12_2018_3; diagnostic periodogram.

Figure S3.125. Raccoon; raccoon_12_2018_4; diagnostic periodogram.

Figure S3.126. Raccoon; raccoon_12_2018_5; diagnostic periodogram.

Figure S3.127. Raccoon; raccoon_12_2018_6; diagnostic periodogram.

Figure S3.128. Raccoon; raccoon_13_2017_11; diagnostic periodogram.

Figure S3.129. Raccoon; raccoon_13_2017_12; diagnostic periodogram.

Figure S3.130. Raccoon; raccoon_13_2018_1; diagnostic periodogram.

Figure S3.131. Raccoon; raccoon_13_2018_2; diagnostic periodogram.

Figure S3.132. Raccoon; raccoon_13_2018_3; diagnostic periodogram.

Figure S3.133. Raccoon; raccoon_13_2018_4; diagnostic periodogram.

Figure S3.134. Raccoon; raccoon_13_2018_5; diagnostic periodogram.

Figure S3.135. Raccoon; raccoon_14_2018_2; diagnostic periodogram.

Figure S3.136. Raccoon; raccoon_14_2018_3; diagnostic periodogram.

Figure S3.137. Raccoon; raccoon_15_2017_12; diagnostic periodogram.

Figure S3.138. Raccoon; raccoon_15_2018_1; diagnostic periodogram.

Figure S3.139. Raccoon; raccoon_15_2018_2; diagnostic periodogram.

Figure S3.140. Raccoon; raccoon_15_2018_3; diagnostic periodogram.

Figure S3.141. Raccoon; raccoon_15_2018_4; diagnostic periodogram.

Figure S3.142. Raccoon; raccoon_15_2018_5; diagnostic periodogram.

Figure S3.143. Raccoon; raccoon_15_2018_6; diagnostic periodogram.

Figure S3.144. Raccoon; raccoon_15_2018_7; diagnostic periodogram.

Figure S3.145. Raccoon; raccoon_16_2018_10; diagnostic periodogram.

Figure S3.146. Raccoon; raccoon_16_2018_11; diagnostic periodogram.

Figure S3.147. Raccoon; raccoon_16_2018_12; diagnostic periodogram.

Figure S3.148. Raccoon; raccoon_16_2018_7; diagnostic periodogram.

Figure S3.149. Raccoon; raccoon_16_2018_8; diagnostic periodogram.

Figure S3.150. Raccoon; raccoon_16_2018_9; diagnostic periodogram.

Figure S3.151. Raccoon; raccoon_16_2019_1; diagnostic periodogram.

Figure S3.152. Raccoon; raccoon_16_2019_2; diagnostic periodogram.

Figure S3.153. Raccoon; raccoon_16_2019_3; diagnostic periodogram.

Figure S3.154. Raccoon; raccoon_17_2018_10; diagnostic periodogram.

Figure S3.155. Raccoon; raccoon_17_2018_11; diagnostic periodogram.

Figure S3.156. Raccoon; raccoon_17_2018_12; diagnostic periodogram.

Figure S3.157. Raccoon; raccoon_17_2018_7; diagnostic periodogram.

Figure S3.158. Raccoon; raccoon_17_2018_8; diagnostic periodogram.

Figure S3.159. Raccoon; raccoon_17_2018_9; diagnostic periodogram.

Figure S3.160. Raccoon; raccoon_17_2019_1; diagnostic periodogram.

Figure S3.161. Raccoon; raccoon_17_2019_2; diagnostic periodogram.

Figure S3.162. Raccoon; raccoon_17_2019_3; diagnostic periodogram.

Figure S3.163. Raccoon; raccoon_18_2018_10; diagnostic periodogram.

Figure S3.164. Raccoon; raccoon_18_2018_11; diagnostic periodogram.

Figure S3.165. Raccoon; raccoon_18_2018_12; diagnostic periodogram.

Figure S3.166. Raccoon; raccoon_18_2018_8; diagnostic periodogram.

Figure S3.167. Raccoon; raccoon_18_2018_9; diagnostic periodogram.

Figure S3.168. Raccoon; raccoon_18_2019_1; diagnostic periodogram.

Figure S3.169. Raccoon; raccoon_18_2019_2; diagnostic periodogram.

Figure S3.170. Raccoon; raccoon_18_2019_3; diagnostic periodogram.

Figure S3.171. Raccoon; raccoon_19_2018_10; diagnostic periodogram.

Figure S3.172. Raccoon; raccoon_19_2018_11; diagnostic periodogram.

Figure S3.173. Raccoon; raccoon_19_2018_12; diagnostic periodogram.

Figure S3.174. Raccoon; raccoon_19_2019_1; diagnostic periodogram.

Figure S3.175. Raccoon; raccoon_19_2019_2; diagnostic periodogram.

Figure S3.176. Raccoon; raccoon_1_2016_10; diagnostic periodogram.

Figure S3.177. Raccoon; raccoon_1_2016_11; diagnostic periodogram.

Figure S3.178. Raccoon; raccoon_1_2016_12; diagnostic periodogram.

Figure S3.179. Raccoon; raccoon_1_2016_9; diagnostic periodogram.

Figure S3.180. Raccoon; raccoon_1_2017_1; diagnostic periodogram.

Figure S3.181. Raccoon; raccoon_1_2017_2; diagnostic periodogram.

Figure S3.182. Raccoon; raccoon_1_2017_3; diagnostic periodogram.

Figure S3.183. Raccoon; raccoon_1_2017_4; diagnostic periodogram.

Figure S3.184. Raccoon; raccoon_1_2017_5; diagnostic periodogram.

Figure S3.185. Raccoon; raccoon_2_2016_10; diagnostic periodogram.

Figure S3.186. Raccoon; raccoon_2_2016_11; diagnostic periodogram.

Figure S3.187. Raccoon; raccoon_2_2016_12; diagnostic periodogram.

Figure S3.188. Raccoon; raccoon_2_2016_9; diagnostic periodogram.

Figure S3.189. Raccoon; raccoon_2_2017_1; diagnostic periodogram.

Figure S3.190. Raccoon; raccoon_2_2017_2; diagnostic periodogram.

Figure S3.191. Raccoon; raccoon_2_2017_3; diagnostic periodogram.

Figure S3.192. Raccoon; raccoon_2_2017_4; diagnostic periodogram.

Figure S3.193. Raccoon; raccoon_2_2017_5; diagnostic periodogram.

Figure S3.194. Raccoon; raccoon_2_2017_6; diagnostic periodogram.

Figure S3.195. Raccoon; raccoon_2_2017_7; diagnostic periodogram.

Figure S3.196. Raccoon; raccoon_3_2016_10; diagnostic periodogram.

Figure S3.197. Raccoon; raccoon_3_2016_11; diagnostic periodogram.

Figure S3.198. Raccoon; raccoon_3_2016_12; diagnostic periodogram.

Figure S3.199. Raccoon; raccoon_3_2016_9; diagnostic periodogram.

Figure S3.200. Raccoon; raccoon_4_2016_10; diagnostic periodogram.

Figure S3.201. Raccoon; raccoon_4_2016_11; diagnostic periodogram.

Figure S3.202. Raccoon; raccoon_4_2016_12; diagnostic periodogram.

Figure S3.203. Raccoon; raccoon_4_2016_9; diagnostic periodogram.

Figure S3.204. Raccoon; raccoon_4_2017_1; diagnostic periodogram.

Figure S3.205. Raccoon; raccoon_5_2016_10; diagnostic periodogram.

Figure S3.206. Raccoon; raccoon_5_2016_11; diagnostic periodogram.

Figure S3.207. Raccoon; raccoon_5_2016_12; diagnostic periodogram.

Figure S3.208. Raccoon; raccoon_5_2016_9; diagnostic periodogram.

Figure S3.209. Raccoon; raccoon_5_2017_1; diagnostic periodogram.

Figure S3.210. Raccoon; raccoon_5_2017_2; diagnostic periodogram.

Figure S3.211. Raccoon; raccoon_5_2017_3; diagnostic periodogram.

Figure S3.212. Raccoon; raccoon_6_2016_10; diagnostic periodogram.

Figure S3.213. Raccoon; raccoon_6_2016_11; diagnostic periodogram.

Figure S3.214. Raccoon; raccoon_6_2016_12; diagnostic periodogram.

Figure S3.215. Raccoon; raccoon_6_2017_1; diagnostic periodogram.

Figure S3.216. Raccoon; raccoon_6_2017_2; diagnostic periodogram.

Figure S3.217. Raccoon; raccoon_6_2017_3; diagnostic periodogram.

Figure S3.218. Raccoon; raccoon_6_2017_4; diagnostic periodogram.

Figure S3.219. Raccoon; raccoon_6_2017_5; diagnostic periodogram.

Figure S3.220. Raccoon; raccoon_6_2017_6; diagnostic periodogram.

Figure S3.221. Raccoon; raccoon_7_2016_10; diagnostic periodogram.

Figure S3.222. Raccoon; raccoon_7_2016_11; diagnostic periodogram.

Figure S3.223. Raccoon; raccoon_7_2016_12; diagnostic periodogram.

Figure S3.224. Raccoon; raccoon_7_2016_9; diagnostic periodogram.

Figure S3.225. Raccoon; raccoon_7_2017_1; diagnostic periodogram.

Figure S3.226. Raccoon; raccoon_7_2017_2; diagnostic periodogram.

Figure S3.227. Raccoon; raccoon_7_2017_3; diagnostic periodogram.

Figure S3.228. Raccoon; raccoon_7_2017_4; diagnostic periodogram.

Figure S3.229. Raccoon; raccoon_7_2017_5; diagnostic periodogram.

Figure S3.230. Raccoon; raccoon_8_2016_11; diagnostic periodogram.

Figure S3.231. Raccoon; raccoon_8_2016_12; diagnostic periodogram.

Figure S3.232. Raccoon; raccoon_8_2017_1; diagnostic periodogram.

Figure S3.233. Raccoon; raccoon_8_2017_2; diagnostic periodogram.

Figure S3.234. Raccoon; raccoon_8_2017_3; diagnostic periodogram.

Figure S3.235. Raccoon; raccoon_8_2017_4; diagnostic periodogram.

Figure S3.236. Wild boar; boar_10_2014_10; diagnostic periodogram.

Figure S3.237. Wild boar; boar_10_2014_11; diagnostic periodogram.

Figure S3.238. Wild boar; boar_10_2014_12; diagnostic periodogram.

Figure S3.239. Wild boar; boar_10_2014_8; diagnostic periodogram.

Figure S3.240. Wild boar; boar_10_2014_9; diagnostic periodogram.

Figure S3.241. Wild boar; boar_11_2015_3; diagnostic periodogram.

Figure S3.242. Wild boar; boar_1_2013_10; diagnostic periodogram.

Figure S3.243. Wild boar; boar_1_2013_11; diagnostic periodogram.

Figure S3.244. Wild boar; boar_1_2013_12; diagnostic periodogram.

Figure S3.245. Wild boar; boar_1_2013_7; diagnostic periodogram.

Figure S3.246. Wild boar; boar_1_2013_8; diagnostic periodogram.

Figure S3.247. Wild boar; boar_1_2013_9; diagnostic periodogram.

Figure S3.248. Wild boar; boar_2_2013_10; diagnostic periodogram.

Figure S3.249. Wild boar; boar_2_2013_11; diagnostic periodogram.

Figure S3.250. Wild boar; boar_2_2013_12; diagnostic periodogram.

Figure S3.251. Wild boar; boar_2_2013_7; diagnostic periodogram.

Figure S3.252. Wild boar; boar_2_2013_8; diagnostic periodogram.

Figure S3.253. Wild boar; boar_2_2013_9; diagnostic periodogram.

Figure S3.254. Wild boar; boar_2_2014_1; diagnostic periodogram.

Figure S3.255. Wild boar; boar_2_2014_2; diagnostic periodogram.

Figure S3.256. Wild boar; boar_3_2013_11; diagnostic periodogram.

Figure S3.257. Wild boar; boar_3_2013_12; diagnostic periodogram.

Figure S3.258. Wild boar; boar_3_2014_1; diagnostic periodogram.

Figure S3.259. Wild boar; boar_3_2014_2; diagnostic periodogram.

Figure S3.260. Wild boar; boar_3_2014_3; diagnostic periodogram.

Figure S3.261. Wild boar; boar_4_2014_10; diagnostic periodogram.

Figure S3.262. Wild boar; boar_4_2014_2; diagnostic periodogram.

Figure S3.263. Wild boar; boar_4_2014_3; diagnostic periodogram.

Figure S3.264. Wild boar; boar_4_2014_4; diagnostic periodogram.

Figure S3.265. Wild boar; boar_4_2014_5; diagnostic periodogram.

Figure S3.266. Wild boar; boar_4_2014_6; diagnostic periodogram.

Figure S3.267. Wild boar; boar_4_2014_7; diagnostic periodogram.

Figure S3.268. Wild boar; boar_4_2014_8; diagnostic periodogram.

Figure S3.269. Wild boar; boar_4_2014_9; diagnostic periodogram.

Figure S3.270. Wild boar; boar_5_2014_10; diagnostic periodogram.

Figure S3.271. Wild boar; boar_5_2014_4; diagnostic periodogram.

Figure S3.272. Wild boar; boar_5_2014_5; diagnostic periodogram.

Figure S3.273. Wild boar; boar_5_2014_6; diagnostic periodogram.

Figure S3.274. Wild boar; boar_5_2014_7; diagnostic periodogram.

Figure S3.275. Wild boar; boar_5_2014_8; diagnostic periodogram.

Figure S3.276. Wild boar; boar_5_2014_9; diagnostic periodogram.

Figure S3.277. Wild boar; boar_6_2014_10; diagnostic periodogram.

Figure S3.278. Wild boar; boar_6_2014_11; diagnostic periodogram.

Figure S3.279. Wild boar; boar_6_2014_5; diagnostic periodogram.

Figure S3.280. Wild boar; boar_6_2014_6; diagnostic periodogram.

Figure S3.281. Wild boar; boar_6_2014_7; diagnostic periodogram.

Figure S3.282. Wild boar; boar_6_2014_8; diagnostic periodogram.

Figure S3.283. Wild boar; boar_6_2014_9; diagnostic periodogram.

Figure S3.284. Wild boar; boar_7_2014_10; diagnostic periodogram.

Figure S3.285. Wild boar; boar_7_2014_11; diagnostic periodogram.

Figure S3.286. Wild boar; boar_7_2014_12; diagnostic periodogram.

Figure S3.287. Wild boar; boar_7_2014_5; diagnostic periodogram.

Figure S3.288. Wild boar; boar_7_2014_6; diagnostic periodogram.

Figure S3.289. Wild boar; boar_7_2014_7; diagnostic periodogram.

Figure S3.290. Wild boar; boar_7_2014_8; diagnostic periodogram.

Figure S3.291. Wild boar; boar_7_2014_9; diagnostic periodogram.

Figure S3.292. Wild boar; boar_7_2015_1; diagnostic periodogram.

Figure S3.293. Wild boar; boar_7_2015_2; diagnostic periodogram.

Figure S3.294. Wild boar; boar_7_2015_3; diagnostic periodogram.

Figure S3.295. Wild boar; boar_8_2014_10; diagnostic periodogram.

Figure S3.296. Wild boar; boar_8_2014_11; diagnostic periodogram.

Figure S3.297. Wild boar; boar_8_2014_12; diagnostic periodogram.

Figure S3.298. Wild boar; boar_8_2014_6; diagnostic periodogram.

Figure S3.299. Wild boar; boar_8_2014_7; diagnostic periodogram.

Figure S3.300. Wild boar; boar_8_2014_8; diagnostic periodogram.

Figure S3.301. Wild boar; boar_8_2014_9; diagnostic periodogram.

Figure S3.302. Wild boar; boar_9_2014_10; diagnostic periodogram.

Figure S3.303. Wild boar; boar_9_2014_11; diagnostic periodogram.

Figure S3.304. Wild boar; boar_9_2014_12; diagnostic periodogram.

Figure S3.305. Wild boar; boar_9_2014_6; diagnostic periodogram.

Figure S3.306. Wild boar; boar_9_2014_7; diagnostic periodogram.

Figure S3.307. Wild boar; boar_9_2014_8; diagnostic periodogram.

Figure S3.308. Wild boar; boar_9_2014_9; diagnostic periodogram.
