## Supplementary S4 for "Movement strategies reveal the success of mammals in urban areas"

**Supplementary material S4: temporal niche shift model**

**Table S4.1. Species-specific GAMM summaries for movement-derived activity along the urbanisation gradient.** Movement-derived activity was modelled as scaled step-derived speed. Models included a cyclic smooth for local time of day, a linear effect of the urbanisation gradient, a smooth interaction between local time of day and the urbanisation gradient, and a random intercept for individual identity. For parametric terms, estimates, standard errors and t-values are shown. For smooth terms, estimated degrees of freedom (edf), reference degrees of freedom, F-values and p-values are shown.

| **Species** | **Model component** | **Term** | **Estimate** | **SE** | **edf** | **Ref. df** | **t / F** | **p-value** |
| --- | --- | --- | --- | --- | --- | --- | --- | --- |
| Fox | Parametric | Intercept | 0.261 | 0.220 | — | — | 1.18 | 0.237 |
| Fox | Parametric | Urbanisation gradient | 0.125 | 0.008 | — | — | 15.40 | <0.001 |
| Fox | Smooth | Time of day | — | — | 15.67 | 18.00 | 30432.11 | 0.055 |
| Fox | Smooth | Time of day × urbanisation | — | — | 16.42 | 17.54 | 93.54 | <0.001 |
| Fox | Random effect | Individual identity | — | — | 16.97 | 17.00 | 1372.55 | <0.001 |
| Raccoon | Parametric | Intercept | −0.187 | 0.086 | — | — | −2.17 | 0.030 |
| Raccoon | Parametric | Urbanisation gradient | −0.082 | 0.016 | — | — | −5.24 | <0.001 |
| Raccoon | Smooth | Time of day | — | — | 13.08 | 18.00 | 1356.95 | <0.001 |
| Raccoon | Smooth | Time of day × urbanisation | — | — | 14.12 | 15.88 | 63.98 | <0.001 |
| Raccoon | Random effect | Individual identity | — | — | 31.73 | 32.00 | 275.73 | <0.001 |
| Wild boar | Parametric | Intercept | 0.001 | 0.010 | — | — | 0.07 | 0.942 |
| Wild boar | Parametric | Urbanisation gradient | −0.004 | 0.004 | — | — | −0.85 | 0.394 |
| Wild boar | Smooth | Time of day | — | — | 2.86 | 18.00 | 1.00 | 0.887 |
| Wild boar | Smooth | Time of day × urbanisation | — | — | 4.97 | 6.40 | 1.21 | 0.237 |
| Wild boar | Random effect | Individual identity | — | — | 6.17 | 10.00 | 1.55 | 0.006 |

Model fit: fox, n = 138,717, adjusted R² = 0.266, deviance explained = 26.6%; raccoon, n = 46,956, adjusted R² = 0.242, deviance explained = 24.3%; wild boar, n = 64,183, adjusted R² = 0.001, deviance explained = 0.1%.

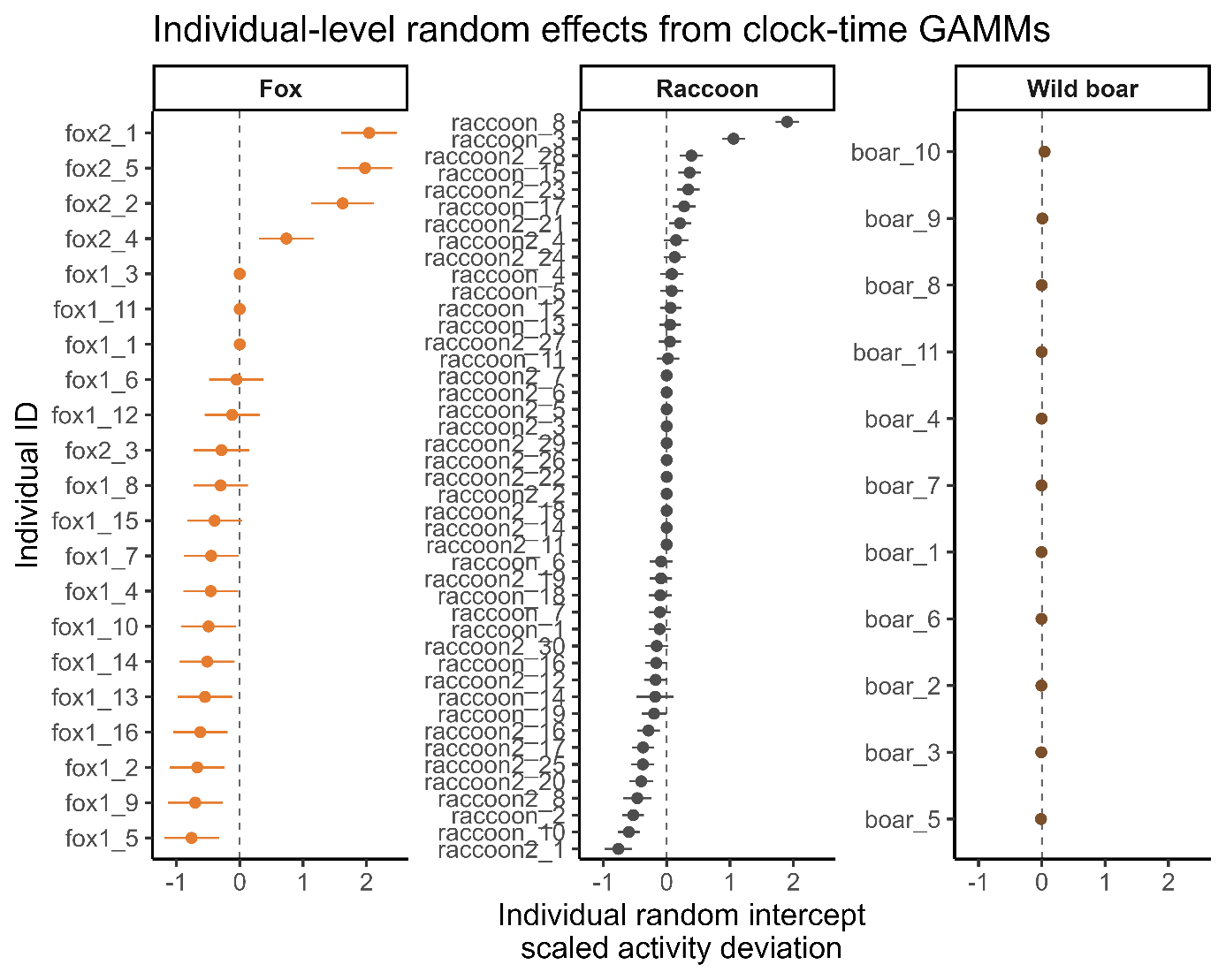

**Figure S4.2 Random effects plot for GAMM models (see S4.1)**

**Table S4.3. Species-level clock-time versus solar-time comparison.** Model comparison based on five-fold cross-validation. ΔlogMSE is calculated as log(MSE solar-relative model) − log(MSE clock-time model); positive values indicate better predictive performance of fixed clock-time models.

| **comparison** | **solar_reference** | **species** | **n_folds** | **total_observations** | **mean_individuals_per_fold** | **mean_dates_per_fold** | **mean_mse_clock** | **mean_mse_solar** | **mean_delta_log_mse** | **median_delta_log_mse** | **se_delta_log_mse** | **lower_95** | **upper_95** | **interpretation** |
| --- | --- | --- | --- | --- | --- | --- | --- | --- | --- | --- | --- | --- | --- | --- |
| clock_vs_first_light | first-light time | fox | 5 | 138,717 | 17.4 | 419 | 1.13 | 1.17 | 0.04 | 0.04 | 0 | 0.03 | 0.05 | clock-time clearly better |
| clock_vs_first_light | first-light time | raccoon | 5 | 46,956 | 32.8 | 506 | 0.90 | 0.91 | 0.01 | 0.01 | 0 | 0.01 | 0.01 | clock-time clearly better |
| clock_vs_first_light | first-light time | boar | 5 | 64,183 | 11.0 | 252 | 1.01 | 1.01 | 0.01 | 0.01 | 0 | 0.00 | 0.01 | clock-time clearly better |
| clock_vs_sunrise | sunrise time | fox | 5 | 138,717 | 17.4 | 419 | 1.13 | 1.17 | 0.04 | 0.04 | 0 | 0.03 | 0.04 | clock-time clearly better |
| clock_vs_sunrise | sunrise time | raccoon | 5 | 46,956 | 32.8 | 506 | 0.90 | 0.91 | 0.01 | 0.01 | 0 | 0.01 | 0.01 | clock-time clearly better |
| clock_vs_sunrise | sunrise time | boar | 5 | 64,183 | 11.0 | 252 | 1.01 | 1.01 | 0.01 | 0.01 | 0 | 0.00 | 0.01 | clock-time clearly better |

**Table S4.4. Urbanisation-bin clock-time versus solar-time comparison.** Bin-level comparison of fixed clock-time and solar-relative models across the urbanisation gradient within each species.

| **comparison** | **solar_reference** |  | **species** | **urban_bin** |  | **n_fold_bins** | **total_observations** | **mean_individuals_per_fold_bin** | **mean_dates_per_fold_bin** | **mean_mse_clock** | **mean_mse_solar** | **mean_delta_log_mse** | **median_delta_log_mse** | **se_delta_log_mse** | | **lower_95** | **upper_95** | | **urban_min** | **urban_median** | **urban_max** | **interpretation** |
| --- | --- | --- | --- | --- | --- | --- | --- | --- | --- | --- | --- | --- | --- | --- | --- | --- | --- | --- | --- | --- | --- | --- |
| clock_vs_first_light | first-light time |  | fox | Low |  | 5 | 53,921 | 9.4 | 211.8 | 1.54 | 1.59 | 0.03 | 0.03 | 0.00 | 0.03 | | | 0.04 | -5.41 | 0.05 | 1.27 | clock-time clearly better |
| clock_vs_first_light | first-light time |  | fox | Intermediate |  | 5 | 42,121 | 4.0 | 138.2 | 0.79 | 0.83 | 0.04 | 0.05 | 0.00 | 0.04 | | | 0.05 | 1.35 | 1.37 | 1.71 | clock-time clearly better |
| clock_vs_first_light | first-light time |  | fox | High |  | 5 | 42,675 | 5.0 | 127.8 | 0.92 | 0.98 | 0.06 | 0.06 | 0.00 | 0.05 | | | 0.07 | 1.96 | 1.99 | 2.60 | clock-time clearly better |
| clock_vs_first_light | first-light time |  | raccoon | Low |  | 5 | 16,477 | 16.0 | 98.6 | 1.19 | 1.22 | 0.02 | 0.02 | 0.00 | 0.01 | | | 0.02 | -2.72 | -2.01 | -1.57 | clock-time clearly better |
| clock_vs_first_light | first-light time |  | raccoon | Intermediate |  | 5 | 15,617 | 9.8 | 233.6 | 0.66 | 0.65 | 0.00 | 0.00 | 0.00 | -0.01 | | | 0.00 | -1.47 | -0.65 | 1.29 | first-light time weakly better / uncertain |
| clock_vs_first_light | first-light time |  | raccoon | High |  | 5 | 14,862 | 11.0 | 263.0 | 0.83 | 0.84 | 0.01 | 0.01 | 0.00 | 0.00 | | | 0.01 | 1.44 | 1.46 | 2.27 | clock-time clearly better |
| clock_vs_first_light | first-light time |  | boar | Low |  | 5 | 22,068 | 6.0 | 108.8 | 2.95 | 2.95 | -0.02 | -0.02 | 0.01 | -0.04 | | | 0.00 | -5.83 | -3.04 | -2.51 | first-light time clearly better |
| clock_vs_first_light | first-light time |  | boar | Intermediate |  | 5 | 21,353 | 6.0 | 102.0 | 0.00 | 0.00 | 0.04 | 0.03 | 0.01 | 0.02 | | | 0.05 | -2.47 | -1.51 | -1.35 | clock-time clearly better |
| clock_vs_first_light | first-light time |  | boar | High |  | 5 | 20,762 | 6.0 | 95.0 | 0.01 | 0.01 | 0.03 | 0.03 | 0.01 | 0.02 | | | 0.05 | -1.28 | -0.63 | 0.81 | clock-time clearly better |
| clock_vs_sunrise | sunrise time |  | fox | Low |  | 5 | 53,921 | 9.4 | 211.8 | 1.54 | 1.58 | 0.03 | 0.03 | 0.00 | 0.02 | | | 0.03 | -5.41 | 0.05 | 1.27 | clock-time clearly better |
| clock_vs_sunrise | sunrise time |  | fox | Intermediate |  | 5 | 42,121 | 4.0 | 138.2 | 0.79 | 0.83 | 0.04 | 0.04 | 0.00 | 0.03 | | | 0.05 | 1.35 | 1.37 | 1.71 | clock-time clearly better |
| clock_vs_sunrise | sunrise time |  | fox | High |  | 5 | 42,675 | 5.0 | 127.8 | 0.92 | 0.97 | 0.06 | 0.05 | 0.00 | 0.05 | | | 0.06 | 1.96 | 1.99 | 2.60 | clock-time clearly better |
| clock_vs_sunrise | sunrise time |  | raccoon | Low |  | 5 | 16,477 | 16.0 | 98.6 | 1.19 | 1.21 | 0.02 | 0.02 | 0.00 | 0.01 | | | 0.02 | -2.72 | -2.01 | -1.57 | clock-time clearly better |
| clock_vs_sunrise | sunrise time |  | raccoon | Intermediate |  | 5 | 15,617 | 9.8 | 233.6 | 0.66 | 0.65 | 0.00 | 0.00 | 0.00 | 0.00 | | | 0.00 | -1.47 | -0.65 | 1.29 | sunrise time weakly better / uncertain |
| clock_vs_sunrise | sunrise time |  | raccoon | High |  | 5 | 14,862 | 11.0 | 263.0 | 0.83 | 0.84 | 0.01 | 0.01 | 0.00 | 0.00 | | | 0.01 | 1.44 | 1.46 | 2.27 | clock-time clearly better |
| clock_vs_sunrise | sunrise time |  | boar | Low |  | 5 | 22,068 | 6.0 | 108.8 | 2.95 | 2.95 | -0.02 | -0.02 | 0.01 | -0.04 | | | 0.00 | -5.83 | -3.04 | -2.51 | sunrise time clearly better |
| clock_vs_sunrise | sunrise time |  | boar | Intermediate |  | 5 | 21,353 | 6.0 | 102.0 | 0.00 | 0.00 | 0.03 | 0.03 | 0.01 | 0.02 | | | 0.05 | -2.47 | -1.51 | -1.35 | clock-time clearly better |
| clock_vs_sunrise | sunrise time |  | boar | High |  | 5 | 20,762 | 6.0 | 95.0 | 0.01 | 0.01 | 0.03 | 0.03 | 0.01 | 0.02 | | | 0.04 | -1.28 | -0.63 | 0.81 | clock-time clearly better |

**Table S4.5. Representative urbanisation values used for prediction.** Low, intermediate, and high urbanisation values used for visualising activity curves. These correspond to the 5th, 50th, and 95th percentiles of the species-specific urbanisation-gradient values.

| **species** | **low** | **intermediate** | **high** |
| --- | --- | --- | --- |
| fox | -0.87 | 1.37 | 2.57 |
| raccoon | -2.31 | -0.65 | 1.80 |
| boar | -3.25 | -1.51 | 0.81 |

**S4.6 Additional Analyses Solar vs. clock-time**

Methods:

To assess whether urbanisation-mediated activity patterns aligned more closely with fixed clock time, corresponding to daily human activity schedules, or with solar-relative time, we fitted equivalent GAMMs using clock time, time since civil dawn, or time since sunrise as the cyclic temporal predictor. Solar times were calculated for each sampling date using suncalc (Thieurmel & Elmarhraoui, 2022). We compared clock-time and solar-relative models within species using five-fold cross-validation, with folds assigned at the individual-date level. Predictive performance was measured as the log-difference in mean squared error, calculated as log(MSEsolar-relative) − log(MSEclock-time), so that positive values indicate lower prediction error for the fixed clock-time model.

Results:

At the species level, fixed clock-time models consistently showed slightly lower prediction error than models based on time since first light or sunrise. For the first-light comparison, ΔlogMSE values were positive for all species, although effect sizes were small: foxes showed the clearest predictive advantage of clock time (0.040, 95% interval = 0.035–0.045), followed by raccoons (0.010, 0.008–0.011) and wild boar (0.009, 0.003–0.015). The sunrise comparison yielded the same qualitative result (fox: 0.038, 0.033–0.043; raccoon: 0.009, 0.007–0.010; wild boar: 0.007, 0.001–0.012). Local comparisons along the urbanisation gradient further suggested species-specific patterns in this clock-time advantage. In foxes, the predictive advantage of fixed clock time increased with urbanisation (Figure S4.6), consistent with the interpretation that urban fox activity is increasingly organised around predictable human daily schedules rather than light conditions alone. In raccoons, the clock-time advantage was strongest at lower levels of urbanisation but declined towards more urban conditions, suggesting weaker alignment with fixed daily schedules in highly urbanised environments. Wild boar showed a positive clock-time advantage across much of the observed gradient, although this pattern declined under more urban conditions and should be interpreted cautiously given the weak diel structure detected in the main activity models. Overall, these supplementary analyses support the use of local clock time in the main diel activity analyses and indicate that the relevance of fixed anthropogenic schedules versus solar-relative timing varies among species and along the urbanisation gradient.

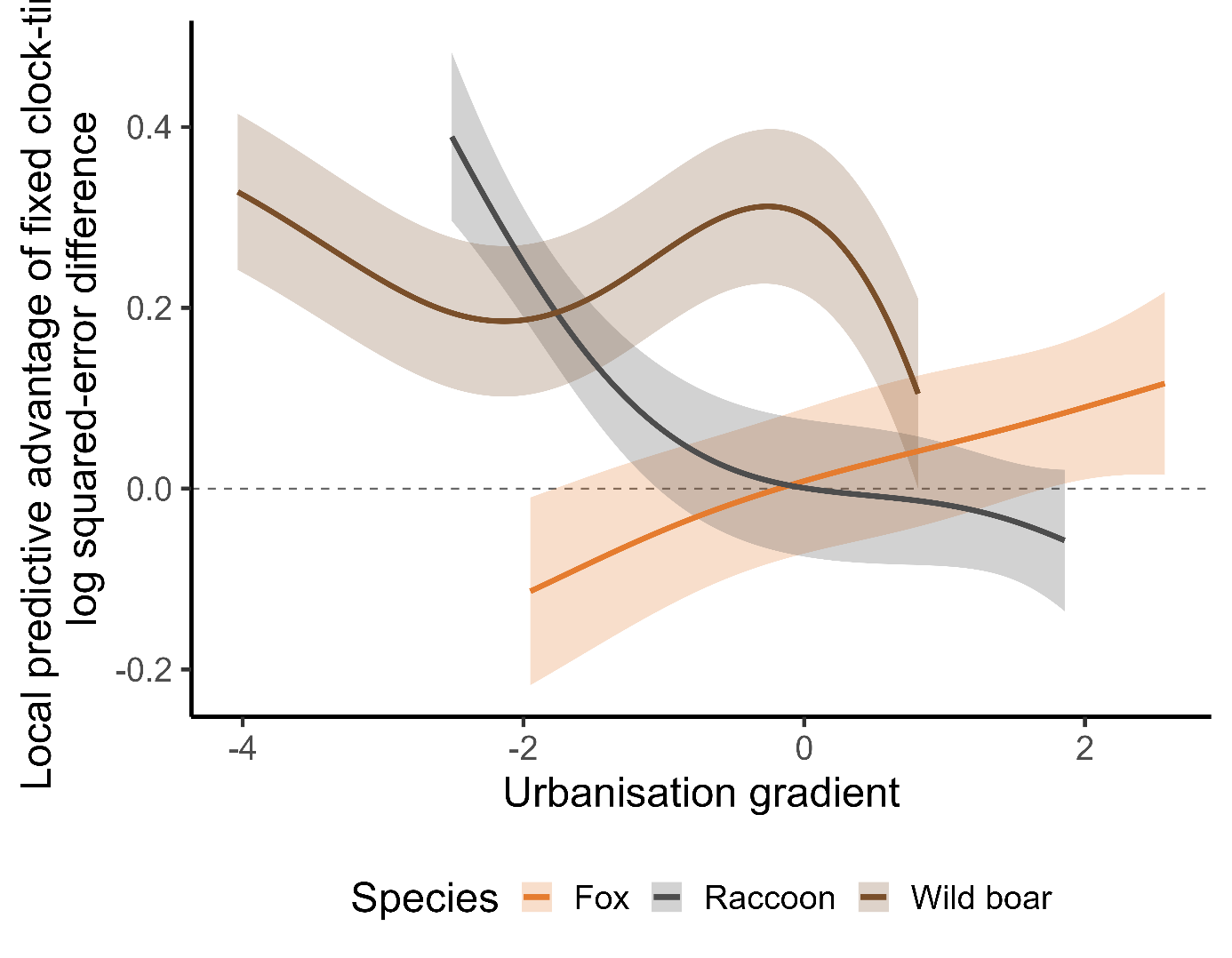

**Figure S4.7: Predictive advantage of fixed-clock time over sunlight times**
