## Supplementary S5 for "Movement strategies reveal the success of mammals in urban areas"

**Supplementary S5 two-part Bayesian regression approach**

**Table S5.1 Coefficient estimates from the hurdle-beta model of periodic movement.** Estimates are posterior means with 89% credible intervals in brackets. The first model estimates the probability that a periodic component was detected using a Bernoulli model. The second model estimates positive periodic strength conditional on periodicity being detected using a beta regression. Wild boar is the reference species. Urbanisation refers to the global urbanisation PC1, where higher values indicate more urbanised conditions.

|  | Periodicity occurrence | Positive periodic strength |
| --- | --- | --- |
| b_Intercept | 0.91 [-0.24, 2.05] | -0.42 [-0.80, -0.02] |
|  | [-0.24, 2.05] | [-0.80, -0.02] |
| b_speciesfox | 1.63 [0.30, 3.10] | 0.03 [-0.43, 0.47] |
|  | [0.30, 3.10] | [-0.43, 0.47] |
| b_speciesraccoon | 0.42 [-0.77, 1.72] | 0.02 [-0.41, 0.43] |
|  | [-0.77, 1.72] | [-0.41, 0.43] |
| b_urban_pc1_global | -0.41 [-0.95, 0.05] | 0.06 [-0.10, 0.22] |
|  | [-0.95, 0.05] | [-0.10, 0.22] |
| b_month_centered | -0.02 [-0.09, 0.05] | 0.01 [-0.01, 0.04] |
|  | [-0.09, 0.05] | [-0.01, 0.04] |
| b_month_centered2 | -0.04 [-0.07, -0.02] | -0.01 [-0.02, -0.01] |
|  | [-0.07, -0.02] | [-0.02, -0.01] |
| b_speciesfox × urban_pc1_global | 0.71 [0.11, 1.38] | 0.03 [-0.17, 0.23] |
|  | [0.11, 1.38] | [-0.17, 0.23] |
| b_speciesraccoon × urban_pc1_global | 0.44 [-0.13, 1.09] | -0.11 [-0.30, 0.08] |
|  | [-0.13, 1.09] | [-0.30, 0.08] |
| sd_ID__Intercept | 1.18 [0.66, 1.83] | 0.36 [0.24, 0.50] |
|  | [0.66, 1.83] | [0.24, 0.50] |
| Num.Obs. | 295 | 212 |
| R2 | 0.229 | 0.239 |
| R2 Marg. | 0.111 | 0.091 |
| ELPD | -165.1 | 78.7 |
| ELPD s.e. | 9.0 | 15.3 |
| LOOIC | 330.2 | -157.4 |
| LOOIC s.e. | 18.1 | 30.5 |
| WAIC | 328.0 | -159.6 |
| RMSE | 0.38 | 0.15 |

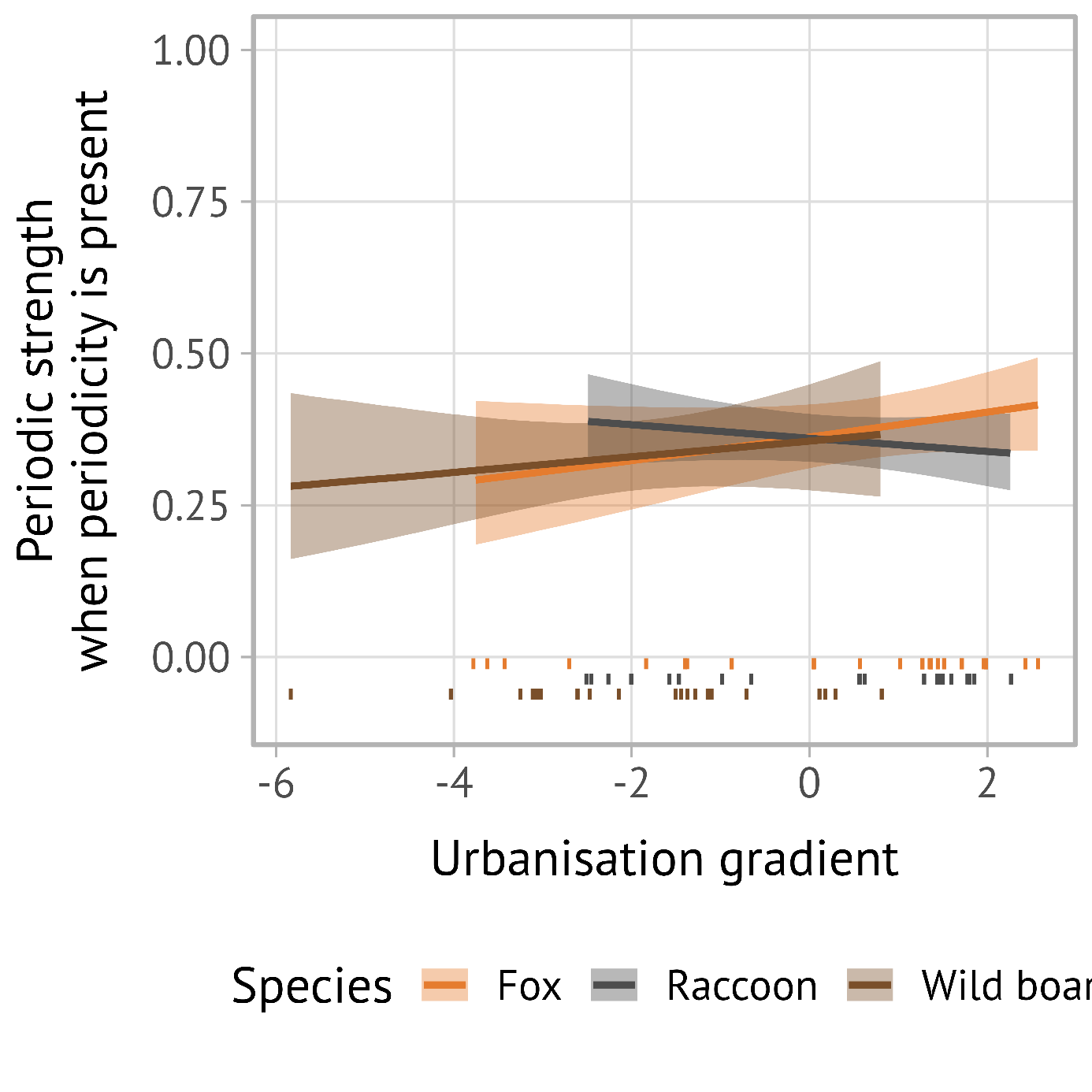

Figure S5.2 Conditional on periodicity being present, positive periodic strength showed little evidence of systematic variation with urbanisation or species (Figure SXX). Among observations with detected periodicity, the urbanisation slope for wild boar was weak and uncertain (β = 0.06 [−0.10, 0.22]). Foxes and raccoons did not show clearly different slopes from wild boar (fox × urbanisation: β = 0.03 [−0.17, 0.23]; raccoon × urbanisation: β = −0.11 [−0.30, 0.08]). Species differences in positive periodic strength were also small and uncertain (fox: β = 0.02 [−0.43, 0.47]; raccoon: β = 0.02 [−0.41, 0.43]). The conditional positive-strength model showed a weak linear month effect (β = 0.01 [−0.01, 0.04]) and a negative quadratic month effect (β = −0.01 [−0.02, −0.01]), suggesting slightly reduced periodic strength toward the seasonal margins. Among-individual variation in positive periodic strength was lower than in the periodicity-occurrence model but remained evident (SD = 0.37 [0.24, 0.50]).
