## Supplementary S6 for "Movement strategies reveal the success of mammals in urban areas"

**Table S6.1. Fixed effects from iSSF models**.

| model | term | estimate | std.error | conf.low | conf.high | statistic | p.value |
| --- | --- | --- | --- | --- | --- | --- | --- |
| Fox | (Intercept) | -26.40 | 0.73 | -27.82 | -24.97 | -36.33 | <0.001 |
| Fox | sl_ | 0.00 | 0.00 | 0.00 | 0.00 | 227.43 | <0.001 |
| Fox | ta_ | 0.00 | 0.00 | 0.00 | 0.00 | -0.22 | 0.82 |
| Fox | tod_binnight:prox_roads_sc | 0.16 | 0.01 | 0.14 | 0.18 | 18.84 | <0.001 |
| Fox | tod_binmorning:prox_roads_sc | -0.22 | 0.02 | -0.25 | -0.19 | -14.27 | <0.001 |
| Fox | tod_binafternoon:prox_roads_sc | -0.08 | 0.02 | -0.12 | -0.05 | -4.40 | <0.001 |
| Fox | tod_binevening:prox_roads_sc | 0.02 | 0.01 | 0.00 | 0.04 | 2.22 | 0.03 |
| Fox | tod_binnight:prox_paths_sc | 0.11 | 0.00 | 0.10 | 0.12 | 27.96 | <0.001 |
| Fox | tod_binmorning:prox_paths_sc | 0.07 | 0.01 | 0.05 | 0.08 | 8.75 | <0.001 |
| Fox | tod_binafternoon:prox_paths_sc | 0.12 | 0.01 | 0.11 | 0.14 | 13.11 | <0.001 |
| Fox | tod_binevening:prox_paths_sc | 0.10 | 0.00 | 0.09 | 0.11 | 23.14 | <0.001 |
| Fox | tod_binnight:treecover_sc | -0.11 | 0.00 | -0.12 | -0.11 | -37.23 | <0.001 |
| Fox | tod_binmorning:treecover_sc | -0.10 | 0.01 | -0.12 | -0.09 | -16.36 | <0.001 |
| Fox | tod_binafternoon:treecover_sc | -0.08 | 0.01 | -0.09 | -0.06 | -9.36 | <0.001 |
| Fox | tod_binevening:treecover_sc | -0.11 | 0.00 | -0.12 | -0.11 | -33.50 | <0.001 |
| Fox | tod_binnight:prox_roads_sc:human_mod_sc | 0.05 | 0.00 | 0.05 | 0.06 | 15.31 | <0.001 |
| Fox | tod_binmorning:prox_roads_sc:human_mod_sc | -0.01 | 0.01 | -0.02 | 0.00 | -1.47 | 0.14 |
| Fox | tod_binafternoon:prox_roads_sc:human_mod_sc | -0.02 | 0.01 | -0.03 | 0.00 | -2.44 | 0.01 |
| Fox | tod_binevening:prox_roads_sc:human_mod_sc | -0.01 | 0.00 | -0.01 | 0.00 | -2.11 | 0.04 |
| Fox | tod_binnight:prox_paths_sc:human_mod_sc | -0.02 | 0.00 | -0.03 | -0.02 | -7.32 | <0.001 |
| Fox | tod_binmorning:prox_paths_sc:human_mod_sc | -0.03 | 0.01 | -0.04 | -0.02 | -6.04 | <0.001 |
| Fox | tod_binafternoon:prox_paths_sc:human_mod_sc | 0.02 | 0.01 | 0.01 | 0.03 | 3.11 | <0.001 |
| Fox | tod_binevening:prox_paths_sc:human_mod_sc | -0.02 | 0.00 | -0.03 | -0.01 | -6.69 | <0.001 |
| Fox | tod_binnight:treecover_sc:human_mod_sc | 0.07 | 0.00 | 0.06 | 0.07 | 26.99 | <0.001 |
| Fox | tod_binmorning:treecover_sc:human_mod_sc | 0.12 | 0.00 | 0.12 | 0.13 | 26.86 | <0.001 |
| Fox | tod_binafternoon:treecover_sc:human_mod_sc | 0.07 | 0.01 | 0.06 | 0.08 | 12.86 | <0.001 |
| Fox | tod_binevening:treecover_sc:human_mod_sc | 0.07 | 0.00 | 0.07 | 0.08 | 25.77 | <0.001 |
| Raccoon | (Intercept) | -0.21 | 4.40 | -8.84 | 8.43 | -0.05 | 0.96 |
| Raccoon | sl_ | 0.00 | 0.00 | 0.00 | 0.00 | 24.09 | <0.001 |
| Raccoon | ta_ | 0.00 | 0.00 | 0.00 | 0.01 | 3.80 | <0.001 |
| Raccoon | tod_binnight:prox_roads_sc | -0.33 | 0.01 | -0.35 | -0.32 | -45.17 | <0.001 |
| Raccoon | tod_binmorning:prox_roads_sc | -0.29 | 0.02 | -0.33 | -0.24 | -13.29 | <0.001 |
| Raccoon | tod_binafternoon:prox_roads_sc | -0.14 | 0.03 | -0.20 | -0.08 | -4.65 | <0.001 |
| Raccoon | tod_binevening:prox_roads_sc | -0.25 | 0.01 | -0.27 | -0.23 | -29.42 | <0.001 |
| Raccoon | tod_binnight:prox_paths_sc | 0.03 | 0.01 | 0.02 | 0.05 | 5.19 | <0.001 |
| Raccoon | tod_binmorning:prox_paths_sc | -0.06 | 0.02 | -0.09 | -0.03 | -3.84 | <0.001 |
| Raccoon | tod_binafternoon:prox_paths_sc | 0.17 | 0.02 | 0.13 | 0.21 | 8.01 | <0.001 |
| Raccoon | tod_binevening:prox_paths_sc | 0.00 | 0.01 | -0.01 | 0.02 | 0.65 | 0.52 |
| Raccoon | tod_binnight:treecover_sc | -0.14 | 0.00 | -0.15 | -0.13 | -35.44 | <0.001 |
| Raccoon | tod_binmorning:treecover_sc | -0.19 | 0.01 | -0.21 | -0.17 | -18.58 | <0.001 |
| Raccoon | tod_binafternoon:treecover_sc | -0.31 | 0.01 | -0.34 | -0.29 | -21.92 | <0.001 |
| Raccoon | tod_binevening:treecover_sc | -0.04 | 0.00 | -0.05 | -0.03 | -7.85 | <0.001 |
| Raccoon | tod_binnight:prox_roads_sc:human_mod_sc | -0.39 | 0.01 | -0.41 | -0.38 | -51.54 | <0.001 |
| Raccoon | tod_binmorning:prox_roads_sc:human_mod_sc | -0.33 | 0.02 | -0.36 | -0.29 | -19.30 | <0.001 |
| Raccoon | tod_binafternoon:prox_roads_sc:human_mod_sc | -0.21 | 0.02 | -0.26 | -0.17 | -9.74 | <0.001 |
| Raccoon | tod_binevening:prox_roads_sc:human_mod_sc | -0.44 | 0.01 | -0.45 | -0.42 | -50.39 | <0.001 |
| Raccoon | tod_binnight:prox_paths_sc:human_mod_sc | -0.06 | 0.01 | -0.07 | -0.04 | -8.73 | <0.001 |
| Raccoon | tod_binmorning:prox_paths_sc:human_mod_sc | 0.01 | 0.01 | -0.02 | 0.04 | 0.82 | 0.41 |
| Raccoon | tod_binafternoon:prox_paths_sc:human_mod_sc | 0.12 | 0.02 | 0.08 | 0.15 | 6.86 | <0.001 |
| Raccoon | tod_binevening:prox_paths_sc:human_mod_sc | 0.04 | 0.01 | 0.03 | 0.06 | 6.11 | <0.001 |
| Raccoon | tod_binnight:treecover_sc:human_mod_sc | 0.15 | 0.00 | 0.14 | 0.16 | 36.56 | <0.001 |
| Raccoon | tod_binmorning:treecover_sc:human_mod_sc | 0.02 | 0.01 | 0.01 | 0.04 | 2.71 | 0.01 |
| Raccoon | tod_binafternoon:treecover_sc:human_mod_sc | -0.09 | 0.01 | -0.11 | -0.07 | -8.15 | <0.001 |
| Raccoon | tod_binevening:treecover_sc:human_mod_sc | 0.12 | 0.00 | 0.11 | 0.13 | 26.40 | <0.001 |
| Wild boar | (Intercept) | -21.76 | 0.90 | -23.53 | -19.98 | -24.06 | <0.001 |
| Wild boar | sl_ | 0.00 | 0.00 | 0.00 | 0.00 | 5.32 | <0.001 |
| Wild boar | ta_ | 0.00 | 0.00 | 0.00 | 0.00 | -1.71 | 0.09 |
| Wild boar | tod_binnight:prox_roads_sc | -0.17 | 0.02 | -0.21 | -0.14 | -9.39 | <0.001 |
| Wild boar | tod_binmorning:prox_roads_sc | 0.16 | 0.02 | 0.12 | 0.20 | 8.15 | <0.001 |
| Wild boar | tod_binafternoon:prox_roads_sc | 0.15 | 0.02 | 0.11 | 0.19 | 7.23 | <0.001 |
| Wild boar | tod_binevening:prox_roads_sc | -0.02 | 0.02 | -0.05 | 0.02 | -1.00 | 0.32 |
| Wild boar | tod_binnight:prox_paths_sc | -0.06 | 0.01 | -0.07 | -0.04 | -7.17 | <0.001 |
| Wild boar | tod_binmorning:prox_paths_sc | -0.33 | 0.01 | -0.34 | -0.31 | -41.07 | <0.001 |
| Wild boar | tod_binafternoon:prox_paths_sc | -0.29 | 0.01 | -0.31 | -0.27 | -35.76 | <0.001 |
| Wild boar | tod_binevening:prox_paths_sc | 0.18 | 0.01 | 0.16 | 0.19 | 22.23 | <0.001 |
| Wild boar | tod_binnight:treecover_sc | 0.02 | 0.01 | 0.01 | 0.04 | 3.35 | <0.001 |
| Wild boar | tod_binmorning:treecover_sc | -0.03 | 0.01 | -0.05 | -0.02 | -4.77 | <0.001 |
| Wild boar | tod_binafternoon:treecover_sc | -0.01 | 0.01 | -0.02 | 0.00 | -1.38 | 0.17 |
| Wild boar | tod_binevening:treecover_sc | 0.05 | 0.01 | 0.03 | 0.06 | 6.62 | <0.001 |
| Wild boar | tod_binnight:prox_roads_sc:human_mod_sc | -0.17 | 0.02 | -0.20 | -0.14 | -10.75 | <0.001 |
| Wild boar | tod_binmorning:prox_roads_sc:human_mod_sc | 0.06 | 0.02 | 0.03 | 0.09 | 3.57 | <0.001 |
| Wild boar | tod_binafternoon:prox_roads_sc:human_mod_sc | 0.04 | 0.02 | 0.01 | 0.07 | 2.44 | 0.01 |
| Wild boar | tod_binevening:prox_roads_sc:human_mod_sc | 0.02 | 0.02 | -0.01 | 0.05 | 1.39 | 0.17 |
| Wild boar | tod_binnight:prox_paths_sc:human_mod_sc | 0.13 | 0.01 | 0.12 | 0.15 | 17.33 | <0.001 |
| Wild boar | tod_binmorning:prox_paths_sc:human_mod_sc | 0.06 | 0.01 | 0.05 | 0.08 | 8.16 | <0.001 |
| Wild boar | tod_binafternoon:prox_paths_sc:human_mod_sc | 0.07 | 0.01 | 0.06 | 0.09 | 9.41 | <0.001 |
| Wild boar | tod_binevening:prox_paths_sc:human_mod_sc | 0.04 | 0.01 | 0.02 | 0.05 | 4.70 | <0.001 |
| Wild boar | tod_binnight:treecover_sc:human_mod_sc | 0.10 | 0.01 | 0.09 | 0.12 | 14.83 | <0.001 |
| Wild boar | tod_binmorning:treecover_sc:human_mod_sc | 0.23 | 0.01 | 0.22 | 0.25 | 33.23 | <0.001 |
| Wild boar | tod_binafternoon:treecover_sc:human_mod_sc | 0.28 | 0.01 | 0.26 | 0.29 | 37.82 | <0.001 |
| Wild boar | tod_binevening:treecover_sc:human_mod_sc | 0.02 | 0.01 | 0.00 | 0.03 | 2.64 | 0.01 |

**Table S6.2. Random effects (variance components) from iSSF models.**

| model | group | term | estimate |
| --- | --- | --- | --- |
| Fox | step_id1_ | sd__(Intercept) | 1,000.000 |
| Fox | id | sd__(Intercept) | 0.053 |
| Raccoon | step_id1_ | sd__(Intercept) | 1,000.000 |
| Raccoon | id | sd__(Intercept) | 0.921 |
| Wild boar | step_id1_ | sd__(Intercept) | 1,000.000 |
| Wild boar | id | sd__(Intercept) | 0.030 |
